# *NPM1::ALK* fusion-derived circular RNAs drive aggressiveness in ALK-positive anaplastic large cell lymphoma

**DOI:** 10.64898/2026.11.09.765349

**Authors:** Loelia Babin, Lola Colras, Sandra Dailhau, Steffen Fuchs, Stéphane Pyronnet, Laurence Lamant, Fabienne Meggetto

**Affiliations:** Inserm, UMR1037 CRCT, Toulouse, France; Université Toulouse III-Paul Sabatier, UMR1037 CRCT, UMR5071 CNRS, Toulouse, France; Equipe labellisée FRM- 2024, Toulouse, France

## Abstract

Circular RNAs (circRNAs) derived from the *NPM1::ALK* fusion have been identified in ALK-positive anaplastic large cell lymphoma (ALK+ ALCL), but their functional contribution to disease progression remains poorly understood. Divergent RT-PCR and long-read Nanopore sequencing identified eight *NPM1::ALK*-derived circRNA variants across ALK(+) ALCL cell-lines, a CRISPR/Cas9-engineered model, and a patient-derived xenograft. The abundance of f-circA23N2 and f-circA26N3, recurrently detected, steadily increased in crizotinib-resistant cell models. Both, highly stable, were predominantly cytoplasmic. Their overexpression in SU-DHL-1 did not affect *NPM1::ALK* expression, cell proliferation, or cell-cycle, in the absence or presence of crizotinib. circRNA overexpression delayed subcutaneous tumor-growth without altering response to crizotinib. Xenograft analysis disclosed altered immune response pathways and changes in inflammatory and myeloid-recruiting cytokines. RNA-seq deconvolution identified remodeling of the myeloid compartment. Intravenous f-circA23N2 and f-circA26N3 shortened survival with systemic lymphoma dissemination. *NPM1::ALK*-derived circRNAs appear to be non-cell-autonomous modulators of cytokine signaling and myeloid remodeling contributing to lymphoma dissemination.

**Statement of significance:**

- Stable, cytoplasmic circular RNAS f-circA23N2 and f-circA26N3 derived from *NPM1::ALK,* are upregulated in crizotinib-resistant ALCL cells.
- Their overexpression remodels the cytokine milieu and myeloid compartment, leading to increased lymphoma dissemination *in vivo*.

## Introduction

Anaplastic large cell lymphoma (ALCL) is a rare and aggressive subtype of peripheral T-cell lymphoma, characterized in approximately 80% of pediatric cases by the presence of the *NPM1::ALK* fusion gene, resulting from the chromosomal translocation t(2;5)(p23;q35) (1). This fusion gene encodes the anaplastic lymphoma kinase (ALK), constitutively active and driving oncogenic signaling, including activation of the STAT3, PI3K-AKT, and MAPK pathways. ALK plays a critical role in ALCL pathogenesis (2).

The current standard of care for ALK(+) ALCL includes multi-agent chemotherapy. In patients resistant to chemotherapy, the ALK-targeting therapy crizotinib is prescribed (3). However, despite initial responses, crizotinib resistance frequently emerges, limiting long-term effectiveness (4). Understanding the molecular mechanisms underlying this resistance is therefore critical to improve treatment strategies.

Emerging evidence indicates that fusion genes not only encode chimeric proteins driving oncogenesis, but also give rise to fusion-derived circular RNAs (f-circRNAs), generated through back-splicing of oncogenic fusion transcripts, identified as a novel class of regulatory molecules in cancer. f-circRNAs biological activity promotes proliferation and exhibits proto-oncogenic functions in various malignancies (5).

In acute myeloid leukemia and acute promyelocytic leukemia (5), f-circM9_1 and f-circPR derive respectively from the *MLL::AF9* and *PML::RAR*α fusions. Both contribute to leukemogenesis independently of their linear mRNA and fusion protein counterparts. Similarly, in non-small cell lung cancer cell lines, f-circEA-2a and f-circEA-4a, both cytoplasmic, derive from *EML4::ALK* fusions and have been shown to promote cell migration and invasion *in vitro*(6). f-circEA1 affects tumor proliferation, migration, invasion, apoptosis and cell cycle, and is involved in resistance to crizotinib (7). *In vivo*, knockdown of f-circEA1 inhibits tumor growth and reduces expression of the parental oncogene *EML4::ALK*(7). These findings support a model in which f-circRNAs, in cooperation with canonical oncogenic drivers, actively sustain tumor development and drug resistance, revealing new potential therapeutic vulnerabilities.

Several f-circRNAs have been identified, using a CRISPR/Cas9-based approach to induce the *NPM1::ALK* translocation in murine and human cells (8). To date, the role of f-circNPM1::ALK in ALCL tumorigenesis remains to be investigated.

Here, two f-circNPM1::ALK were identified, overexpressed in crizotinib-resistant human cell lines. These two f-circRNAs promote tumor dissemination *in vivo* by regulating key mediators of inflammation and cell invasion, including IL-1α, CXCL5, and TIMP2, highlighting the critical role of circRNAs in the aggressiveness of NPM1-ALK(+) ALCL.

## Results

Circular RNA variants derived from the *NPM1::ALK* fusion gene were detected by RT-PCR with divergent primers, as previously described (8). PCR was performed on ALK(+) ALCL cell lines (including ALKima) and a PDX, in both crizotinib-sensitive (parental) and -resistant (CR) forms (**Figure 1A**). As previously reported (8), numerous qRT-PCR artifacts with microhomologies at the junctions were detected, likely resulting from the abundant expression of *NPM1::ALK* mRNA in the cells (8), and notably in the overexpressing M1 clone (9). Purified PCR products were submitted to long-read Nanopore sequencing. The involvement of each exon in the backsplice junction (BSJ) was first examined (**Figure 1B**). Exons ALK21 and ALK27 were rarely involved possibly because they are flanked by unusually short intronic regions (108 and 681 nucleotides, respectively), which may account for their limited participation in BSJ formation (10). BSJs spanning all *NPM::ALK* exon combinations were detected across all samples (≥100 supporting reads; **Figure 1C**). Then, circular RNAs consistency was analyzed across all samples, resulting in the identification of eight candidates (A23N2, A23N4, A24N3, A24N4, A25N4 A26N2, A26N3 A26N4) (**Figure 1C**). Of note A23N2 generated from *NPM1* exon 2 and *ALK* exon 23 was previously reported in ALKIma (8). The average expression of these eight circRNAs was evaluated in crizotinib-resistant cells using qRT-PCR. (**Figure 1D**). A26N3 (from *NPM1* exon 3 and *ALK* exon 26, **Figure S1B**) showed the highest detection levels in crizotinib-resistant cell lines. Given the prior identification of A23N2 in ALKima, both f-circA23N2 and f-circA26N3 were selected for further investigation.

**Figure 1.**
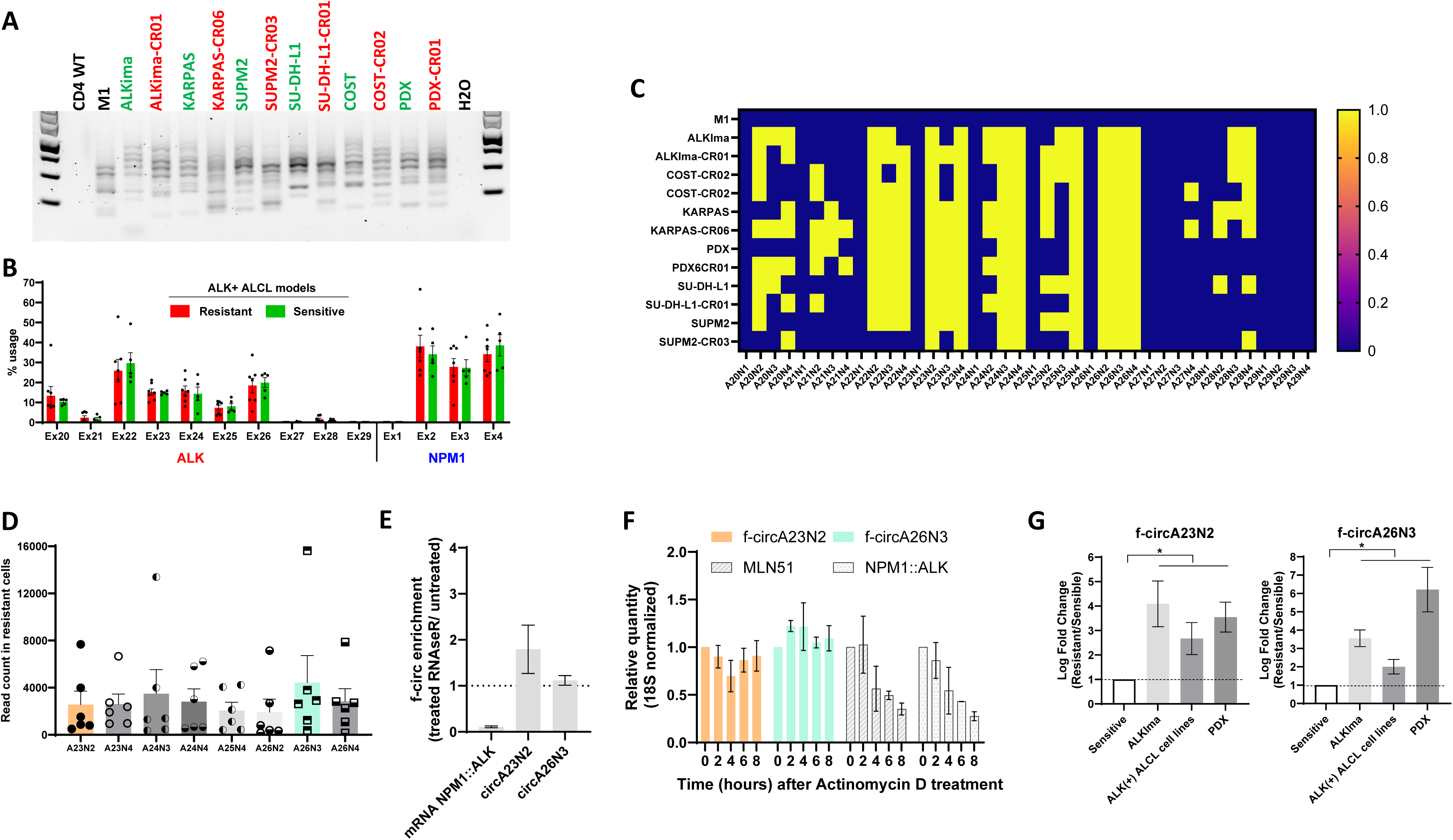
Identification and characterization of *NPM1::ALK*-fusion derived circular RNAs in six crizotinib-sensitive ALK-positive ALCL cell lines and their matched crizotinib-resistant counterparts. **(A)** PCR amplification of *NPM1::ALK* fusion-derived circular RNAs (f-circRNA) using divergent primers targeting *NPM1* exon 4 and *ALK* exon 20. Green: parental/sensitive; red: resistant. Normal CD4 (CD4 WT) and the M1 cell line (established via lentiviral transduction of cDNA *NPM1::ALK* lacking the intronic sequences) were used as negative controls. **(B)** Frequency of exon usage in *NPM1::ALK* f-circRNA across the six paired ALK+ ALCL models Percentages indicate the proportion of Nanopore long reads containing each exon among the PCR products shown in A. As expected, the first exon of *NPM1* and the last exon of *ALK* (exon 29) were never involved in BSJs. **(C)** Heatmap of individual exonic back-splice junctions (BSJs) detected in the six paired ALK+ ALCL cell lines. Yellow and blue respectively indicate the presence and absence of each BSJ. BSJs were identified by Nanopore long-read sequencing of the PCR products shown in A. **(D)** Read counts for selected exonic BSJs identified by Nanopore long-read sequencing of the PCR products shown in A. **(E)** RT-qPCR analysis of *NPM1::ALK* mRNA, f-circ-A23N2, and f-circ-A26N3 following RNase R treatment. Both f-circRNAs were highly stable while the linear *NPM1::ALK* mRNA transcript was almost completely degraded **(F)** RT-qPCR quantification of *MLN5*1, *NPM1::ALK* mRNA, f-circ-A23N2, and f-circ-A26N3 following treatment with actinomycin D for 0, 2, 4, 6, and 8 h. *MLN51* and *NPM-ALK* mRNAs were used as positive controls for mRNA degradation. Expression levels were normalized to 18S rRNA **(G)** Relative enrichment of f-circ-A23N2 and f-circ-A26N3 in crizotinib-resistant versus matched crizotinib-sensitive ALK(+) ALCL cell lines, measured by RT-qPCR and normalized to *MLN51mRNA*.

Both circRNAs showed no degradation after RNase R treatment (**Figure 1E**). Stability was also assessed following transcriptional inhibition with actinomycin D (**Figure 1F**). Expression of both circRNAs remained stable over time. These results confirm the high stability of f-circA23N2 and f-circA26N3. Analyses in qRT-PCR found that f-circA23N2 and f-circA26N3 were predominantly located in the cytoplasm (**Figure S1C**). Their expression levels in crizotinib-sensitive and -resistant cells by qRT-PCR using BSJs-specific divergent primers (**Table S1)** showed that both were consistently upregulated in crizotinib-resistant cells relative to matched parental cells in cell lines, ALKima and PDX (**Figure 1G)**.

The functional contribution of f-circA23N2 and f-circA26N3 to tumorigenesis was next investigated through stable overexpression (OE) using lentiviral vectors. The linear sequences of both circRNAs were cloned into the pcDNA3.1(+) CircRNA Mini Vector (Addgene #60648), which contains minimal intronic sequences to facilitate circularization. CircRNA sequences, flanked by minimal introns, were then subcloned into a lentiviral vector. Overexpression of both circRNAs was confirmed by qRT-PCR (**Figure 2A**), and did not affect *NPM1::ALK* mRNA nor protein levels (**Figure 2B)**. To assess the functional relevance of f-circRNAs *in vivo*, cells overexpressing f-circA23N2 or f-circA26N3 were injected subcutaneously into NSG mice. There was a significant delay in tumor growth, that however did not affect the response to crizotinib (**Figure 2C**). Importantly, this reduced tumorigenicity was not associated with impaired proliferation *in vitro* (**Figure S2**), in the absence or presence of crizotinib. These results suggest that the tumor-suppressive effect observed *in vivo* may involve non-cell autonomous mechanisms, potentially related to interactions with the tumor microenvironment. In addition, during continuous treatment with crizotinib, resistant tumors eventually emerged (**Figure 2D),** indicating that f-circA23N2 and f-circA26N3 play a role in the development of resistance to crizotinib.

**Figure 2:**
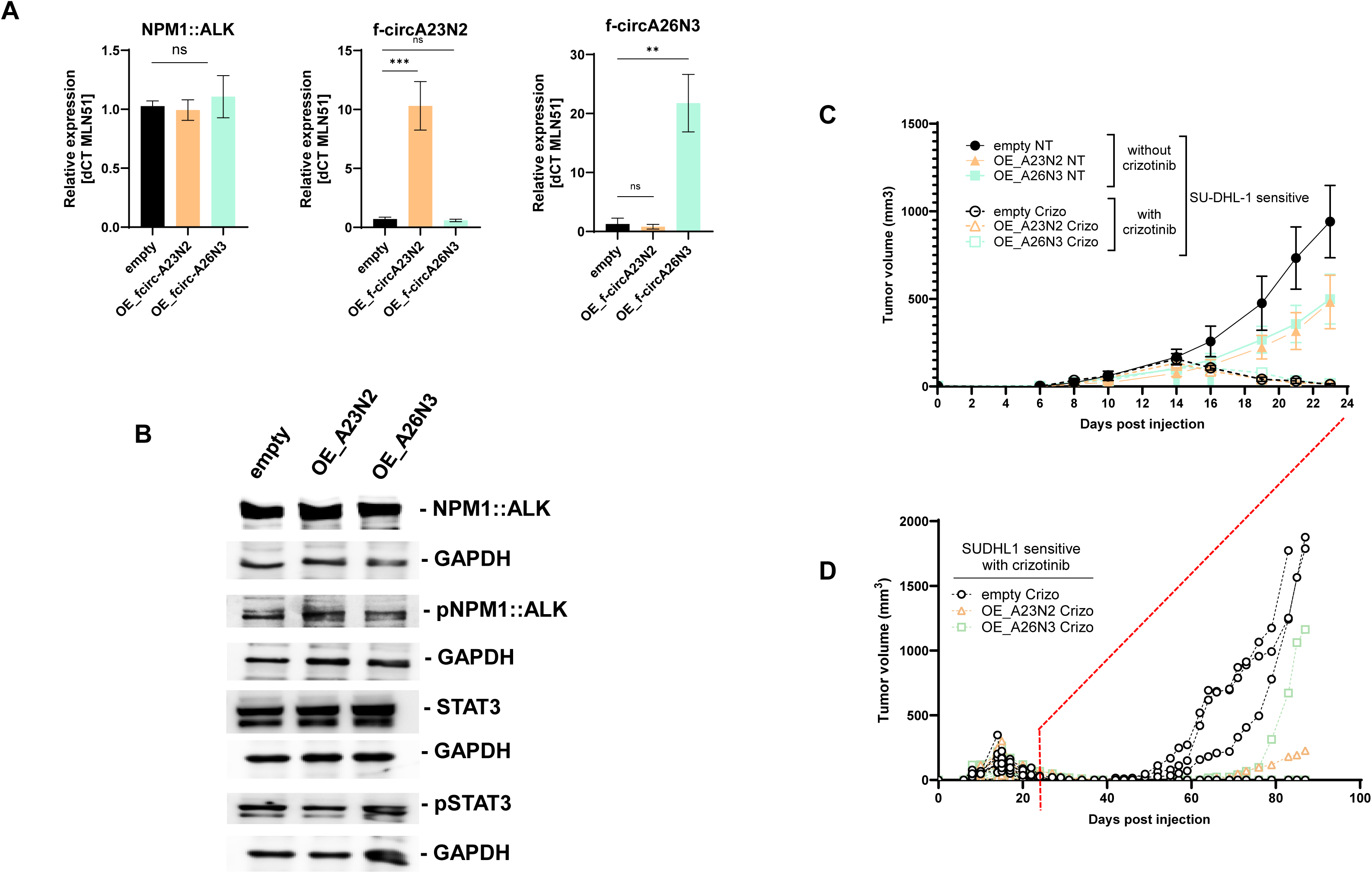
Effects of *NPM1::ALK* fusion-derived f-circRNAs overexpression on NPM1::ALK signaling and tumor growth in SU-DHL-1 cells overexpressing f-circRNAs. SU-DHL-1 transduced cells with empty vector as control (empty) or with overexpressing vectors encoding f-circ-A23N2 (OE-A23N2) or f-circ-A26N3 (OE-A26N3). (**A**) RT-qPCR quantification of *NPM1::ALK* mRNA, f-circ-A23N2, and f-circ-A26N3 expression levels normalized to *MLN51 mRNA.* Circular RNA expression is increased by 10- to 20-fold. **(B)** Western blot analysis of total and phosphorylated NPM1::ALK and STAT3 proteins. GAPDH served as a loading control. Tumor growth following subcutaneous injection of SU-DHL-1 transduced cells with empty or overexpressing vectors. **(C–D)**. Mice received vehicle (NT) or crizotinib (Crizo, 100 mg/kg/day). Tumor volumes, measured with a caliper, are shown from day 0 to day 24 for all treatment groups (**C**). To assess long-term tumor response, follow-up was extended to day 100 in the crizotinib-treated groups only (**D**).

To investigate the molecular mechanisms underlying the *in vivo* phenotype and distinguish tumor-cell intrinsic effects from host microenvironmental responses, xenograft tumors generated from control (OE-empty vector), f-circA23N2 (OE-A23N2) or f-circA26N3 (OE-A26N3)-overexpressing SU-DHL-1 cells were submitted to RNA-seq. Reads were assigned to the human or murine genome (hg38 and mm10), corresponding to tumor-cell and host-derived transcriptomes, respectively. RNA-seq analysis of human-derived SU-DHL-1 tumor cells revealed differential gene expression (DESeq2 analysis, **Table S2**) in f-circ-overexpressing tumors, with enrichment of immune response–related pathways by gene set enrichment analysis (GSEA, **Figure S3A–B**). This suggests that f-circ may indirectly modulate the tumor microenvironment. Consistent with this hypothesis, human cytokine array analysis of 80 cytokines identified 10 commonly altered factors in tumors overexpressing f-circA23N2 or f-circA26N3 (**Figure 3A**). ENA-78/CXCL5, G-CSF, GRO/CXCL1, GRO-α, and IL-1α were increased, whereas TGF-β2, TIMP-1, and TIMP-2 were decreased. For 3 of the 10 commonly altered cytokines identified, antibodies suitable for Western blot validation were available, that confirmed increased CXCL5 and IL-1α expression together with reduced TIMP2 levels (**Figure 3C**), supporting a pro-inflammatory, pro-invasive tumor microenvironment. This cytokine profile, characterized by increased inflammatory and myeloid-recruiting signals, prompted examination of the myeloid compartment in lymphoid-deficient NSG mice tumor models (**Figure S4)**. The ImmuCC tool was used for deconvolution of murine RNA-seq reads. This revealed significant remodeling of myeloid cell populations across all tumors, including an increased M2-like macrophage signature **(Figure 3C**).

**Figure 3.**
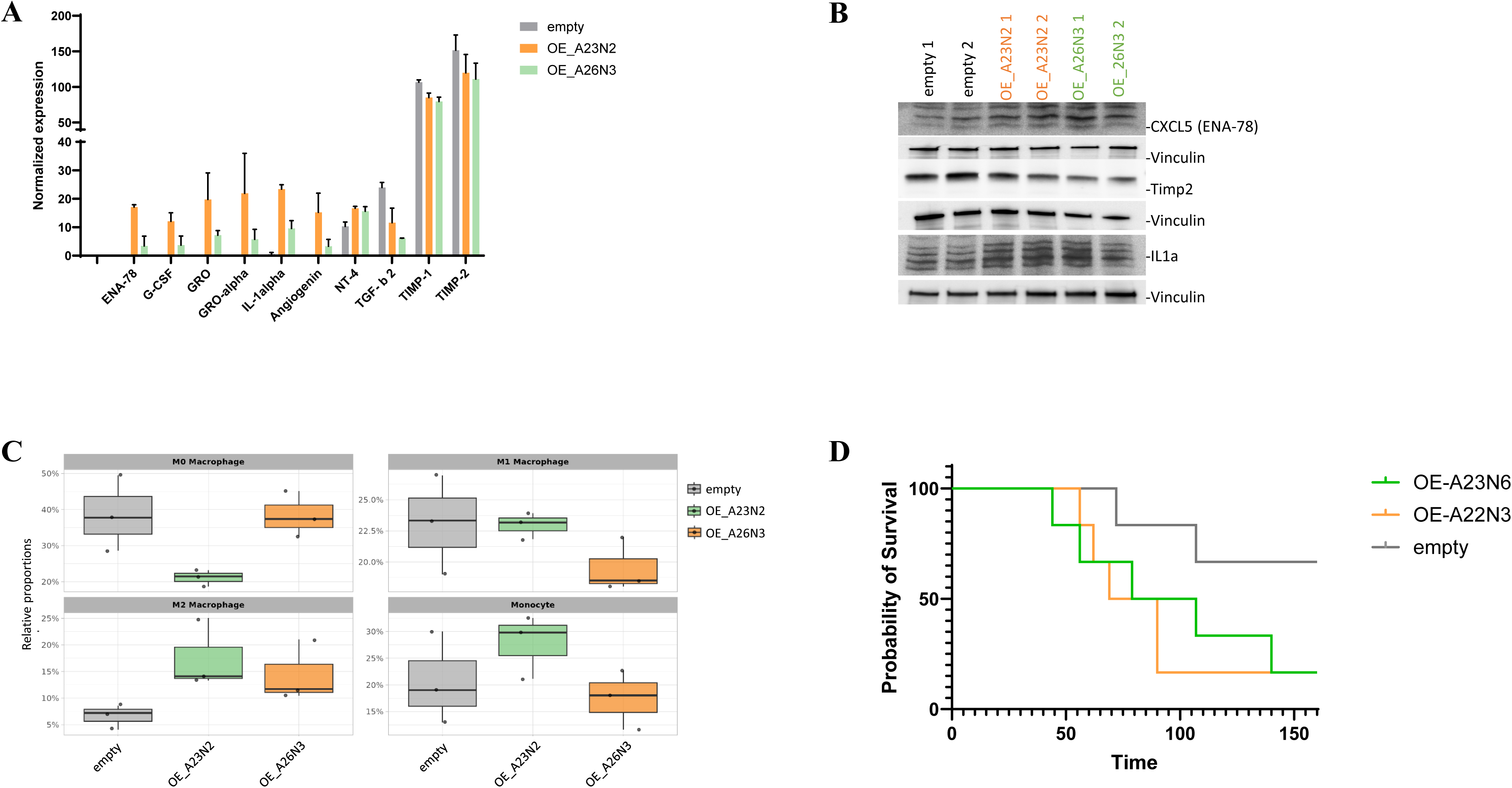
Impact of *NPM1::ALK* fusion-derived f-circRNA overexpression on cytokine expression, myeloid cell composition, and in vivo tumor progression. SU-DHL-1 transduced cells with empty or overexpressing vectors encoding f-circ-A23N2 (OE-A23N2) or f-circ-A26N3 (OE-A26N3). **(A)** Normalized expression levels of differentially expressed cytokines **(B)** Western blot analysis of CXCL5, TIMP2, and IL-1α in tumors obtained from NSG mice subcutaneously injected with transduced SU-DHL-1 cells **(C)** ImmuCC deconvolution of murine RNA-seq reads showing the relative abundance of inferred myeloid cell sub-populations in tumors from the indicated groups. **(D)** Kaplan–Meier survival curves of NSG mice intravenously injected with transduced SU-DHL-1 cells

Collectively, these findings indicate that f-circ expression is associated with coordinated remodeling of the tumor microenvironment, characterized by altered cytokine signaling and myeloid compartment changes, including an M2-like macrophage signature, that may contribute to favor tumor cell migration and dissemination.

Because f-circA23N2 and f-circA26N3 did not affect cell proliferation *in vitro* but altered the expression of cytokines involved in tumor-stroma interactions, it was hypothesized that they would promote lymphoma dissemination *in vivo*. To assess the impact of f-circA23N2 and f-circA26N3 expression on lymphoma dissemination *in vivo*, NSG mice were injected intravenously with control (OE-empty vector), f-circA23N2 (OE-A23N2) or f-circA26N3 (OE-A26N3)-overexpressing SU-DHL-1 cells, and disease progression was monitored over time. Mice injected with f-circA23N2- or f-circA26N3-overexpressing cells exhibited significantly shorter survival than control mice (**Figure 3D**). The liver, spleen, kidneys, and ovaries represented the predominant sites of lymphoma dissemination, with comparable organ tropism between the two f-circ-overexpressing groups (data not shown).

## Discussion

Long-read Nanopore sequencing revealed the overexpression of two fusion circular RNAs derived from the *NPM1::ALK* translocation, across distinct cellular models of resistance to the ALK-targeted agent crizotinib, including cell lines, a CRISPR-engineered translocation model, and a patient-derived xenograft (PDX) model. To investigate their potential role in ALCL, stable overexpression models were established for both circRNAs. Unexpectedly, this overexpression was not associated with any detectable phenotypic changes *in vitro*. Given that circRNAs can be released in the extracellular environment via exosomes and subsequently affect neighboring cells (11), their biological effects may depend on the tissue microenvironment and therefore differ between *in vitro* and *in vivo* settings. Consistent with this possibility, subcutaneous xenografts of cell models overexpressing the f-circRNAs exhibited reduced tumor growth in mice. Further RNA-seq analyses revealed no apparent changes in signaling pathways associated with cell proliferation or apoptosis. Instead, substantial transcriptional changes were observed in genes involved in immune responses, including increased expression of CXCL5 and IL-1α and decreased expression of TIMP2. These findings suggest that the f-circRNAs may influence tumor growth through mechanisms beyond direct regulation of tumor cell proliferation or apoptosis, potentially involving interactions with the tumor microenvironment (TME).

CXCL5 and IL-1α have been implicated in TME remodeling and tumor dissemination. CXCL5 overexpression has been associated with poor prognosis across several solid tumors, as well as increased tumor dissemination and angiogenesis (12). Similarly, IL-1α can promote metastatic progression, as demonstrated in melanoma, where it modulates macrophage activity within the TME (13). Conversely, TIMP2 has been associated with reduced tumor dissemination through the inhibition of matrix metalloproteinases involved in extracellular matrix remodeling (14,15). Given these transcriptional changes and their potential implications for the TME, its cellular composition was subsequently assessed using the ImuCC tool. Because the xenografts were established in immunodeficient NSG mice, the analysis was restricted to the immune cell populations supported by the tool in this model, principally macrophages and monocytes (16,17). Notably, circRNA overexpression models exhibited an increased proportion of M2-like macrophages, a population generally associated with immunosuppressive and pro-tumoral functions (18).

Taken together, these findings suggest that f-circRNAs may influence tumor progression primarily through modulation of TME, rather than through direct effects on tumor cell proliferation, with potential consequences for cell migration and dissemination. This possibility would reconcile the reduced local tumor growth observed in the subcutaneous model with the increased dissemination observed *in vivo*. Indeed, previous studies in melanoma have shown that TME alterations can enhance metastatic potential despite a reduction in primary tumor burden (19). Because subcutaneous xenografts grew rapidly, the humane endpoint was reached before metastatic progression could be adequately assessed. To further investigate the dissemination potential and aggressiveness of the different cell models, intravenous injection experiments were therefore performed. Animals injected with cells overexpressing the f-circRNAs exhibited markedly shorter survival and accelerated dissemination, although the distribution of metastatic sites was similar to that observed in control animals. Collectively, these findings implicate *NPM1::ALK*-derived circRNAs as potential non-cell-autonomous modulators of cytokine-associated signaling and myeloid cell remodeling that may promote lymphoma dissemination.

## Methods

### Cell line culture

The ALCL cell lines KARPAS-299, SU-DHL-1 SUPM2, PIO and COST, were obtained from the DSMZ (German Collection of Microorganisms and Cell Culture, Leibnitz, Germany) or established locally. ALKima, a CRISPR-engineered ALK(+) ALCL cell line was also used (20) as well as patient-derived xenografts (PDX), established and provided by Dr. D. Sibon (21). The crizotinib-resistant KARPAS-299-CR06 and SUPM2-CR03 cell line was provided by C. Gambacorti-Passerini (22). The resistant COST-CR02, SUDHL1-CR01, ALKIma-CR01 and PDX-CR01 cells were obtained by exposing to increasing concentrations of crizotinib, ranging from 25 nM to 200 nM. At each step, the dose was increased only when treated cells achieved a proliferation rate comparable to that of untreated wild-type (WT) cells. All NPM1::ALK(+) ALCL cell lines and PDX cells were cultured in RPMI-1640 medium (Invitrogen, Waltham, MA #61870044) supplemented with 20% FBS (Pan Biotech, Aidenbach, Germany#500105M1M). For crizotinib-resistant cell lines, crizotinib was added to the medium twice a week at a concentration of 100 nM for ALKIma-CR01, SUDHL1-CR01 and PDX-CR01, 200 nM for COST-CR02, 300 nM for SUPM2-CR03 and 600 nM for KARPAS-299-CR06. The cancer cell lines employed in this study had been genotyped in advance to verify expected mutations. Cells were cryopreserved and maintained at early passages. Following thawing, each line was passaged at least once, and *Mycoplasma* contamination was assessed (Lonza, LT07-710) before implantation into mice.

### Crizotinib treatment

Cells were seeded at 2 × 10^5^ cells/mL and treated at 24 h with different doses of crizotinib (Selleckchem, Houston, TX, PF-02341066#S1068).

### Xenograft tumor assay

Mice were housed under pathogen-free conditions in an animal room at constant temperature (20 °C–22 °C) with a 12 h light/dark cycle and free access to food and water. A total of 3 × 10^6^ cells were injected subcutaneously into one flank of 5-week-old female NSG mice (Janvier Labs, Le Genest Saint Isle, France). Mouse body weight and tumor volumes were assessed 3 times a week with calipers via the following formula: length × width2× π/6. At the end of the experiment, the mice (8 per group) were humanely sacrificed. For crizotinib-treated mice, they were treated with 100 mg/kg/day crizotinib, starting when the tumor size reached 300 mm^3^. For animal studies no randomization nor blinding were used.

For caudal intravenous injection, 100 000 cells were injected in the blood circulation. Mouse body weight was recorded and the clinical score was determined once a week. Mice were sacrificed when endpoint was reached.

All animal procedures were performed following the principle guidelines of INSERM. The protocol was approved by the Midi-Pyrénées Ethics Committee on Animal Experimentation and conducted in accordance with institutional guidelines and the Directive 2010/63/EU under protocol number DAP-APAFiS-2021100714596279-, CEEA122 agreement: 2021100714596279_v9). All experiments were conducted in compliance with all relevant guidelines and regulations, including the Animal Research: Reporting of In Vivo Experiments (ARRIVE) guidelines.

### CircRNA validation

The validation of circRNA circularity, and stability were performed as previously reported. In brief circularity and exonuclease resistance were assessed via RNase R treatment, whereas transcript stability was evaluated following exposure to actinomycin D. Relative expression levels were determined via RT qPCR via the 2-ΔΔCt method.

### Western blot analysis

Whole-cell extracts were prepared with protein lysis buffer (50 mM Tris-HCl pH 7.4, 1%Triton X-100, 0.1% SDS, 150 mM NaCl, 1 mM EDTA, and 1 mM DTT), supplemented with «Complete cocktail protease inhibitor tablets » (Roche, Vienna, Austria, #11697498001) and «Halt Protease and Phosphatase Inhibitor Cocktail » (Thermo Fisher Scientific #78440). Typically, 30 μg of protein were loaded on 10% acrylamide gel (w/v) Tris-HCl SDS PAGE. After transfer, membranes were stained with anti-Phospho-ALK (Tyr1604 #3341 RRID: AB_331047), anti-ALK (31F12 #3791, RRID: AB_1950402), anti-P-STAT3 (Tyr705 #9131, RRID: AB_331586), anti-STAT3 (#4904, RRID: AB_331269), anti TIMP2 (#5738, RRID:AB_10694774), anti-IL-1α (#34819) from Cell Signaling, anti-CXCL5 (Abcam, Cambridge, UK #AB305100) and anti-GAPDH (Santa-Cruz,Dallas, TX #sc-32233, RRID: AB_627679) antibodies were used as loading controls. Secondary antibodies were HRP-conjugated, and blot developed with ECL prime (BIO-RAD, Hercules, CA, # 1705060).

### Cell cycle analysis

Cell cycle analysis was performed via an EdU Click-iT kit (Invitrogen, Carlsbad, CA #C10337) following manufacturer instructions.

### RNA extraction, reverse transcription, PCR and quantitative PCR

Total RNA was extracted from 5 to 10x10^6^ cells via TRIzol reagent (Invitrogen # 15596018) followed by a Direct-zol™ RNA Miniprep Plus Kit (Zymo, Orange, CA #R2072) following the manufacturer instructions. Typically, 1 μg of total RNA was reverse transcribed via ProtoScript® II First Strand cDNA Synthesis Kit (New England Biolabs, NEB, Ipswitch, MA, #E6560L) following manufacturer instructions. cDNA was next diluted with 5 in water, and 2 μL were used for PCR and quantitative PCR analysis. PCR was performed as previously described(8). qPCR was performed via the Master Mix Select SYBR™ (Applied Biosystems, Waltham, MA, #4472908) in a 10 μL total reaction. The PCR program was as follows: 50°C for 2 min, 95°C for 2 min, and 40 cycles of 95°C for 15 s and 60°C for 1 min. All primer sequences are listed in Supplementary Table S1.

### Detection of NPM1–ALK fusion circRNAs by full-length Nanopore sequencing

Fusion circRNAs (f-circRNAs) derived from the NPM1–ALK fusion gene were detected in full length using an adapted Oxford Nanopore sequencing strategy. In brief, a cDNA library enriched for NPM1–ALK f-circRNAs was generated and integrated into our previously established Nanopore workflow for full-length circRNA profiling (23,24).

Total RNA was isolated from ALK(+) ALCL cell lines using TRIzol reagent according to the manufacturer’s instructions, as described previously (25). First-strand cDNA was synthesized from 1 µg of total RNA using the ProtoScript II First Strand cDNA Synthesis Kit (New England Biolabs) with random hexamer primers, following the manufacturer’s protocol.

To specifically amplify circular NPM1–ALK fusion transcripts, outward-facing primers were designed to flank the fusion junction between NPM1 exon 4 and ALK exon 20. Primers were positioned to leave approximately 10 nucleotides on each side of the fusion breakpoint to avoid microhomology effects and accommodate nucleotide variations at the breakpoint, thereby ensuring capture of all transcript isoforms (primer sequences: f-circ-ONT-Forw AAGCACCAGGAGCTGCAAG, f-circ-ONT-Rew AATATGCACTGGCCCTGAAC). PCR amplification was carried out using LongAmp Taq DNA Polymerase under cycling conditions and cleaned up using Agencourt AMPure XP beads (Beckman Coulter, Miami, FL).

The purified libraries were then assessed for their quality and Nanopore ligation-based sequencing with Native Barcoding was performed.

### Protein extraction from tumors

Small tumor fragments (∼20 mg) were lysed in protein lysis buffer (see Western blot section for buffer composition) using Precellys system with steel beads (#P000925-LYSK0-A). Program used : 20s on, 20s off, 20s on.

### Cytokines array

Cytokine quantification was performed from tumor protein extracts with Human Cytokine Antibody Array C5 Kit (Raybiotech, Peachtree Corners, GA) following manufacturer instructions.

### RNA dataset generation and analysis

Ribo zero full RNA sequencing library was prepared as previously described. Pooled library prep samples were subsequently sequenced on a NovaSeq 6000 Illumina (San Diego, CA), corresponding to 1x30 million 100-base reads per sample after demultiplexing. The reads were aligned with hisat2 on the hg38 human genome and expression was quantified via featureCounts. Differential expression analysis was performed via the DESeq2 tool. Gene set enrichment analysis was performed using GSEA.

### Over expression model cell lines

Sequences of f-circ for overexpression models were cloned and inserted into the cDNA3.1(+) CircRNA Mini Vector backbone (Addgene, Wartertown, MA, #60648) via EcoRI + NotI digestion. Minimal intronic sequence and circular RNA sequence were subcloned in pCW57.1 lentivector by Gibson assembly (sequences are listed in supplementary table 2).

The cells were transfected with psPAX2 (Addgene #12260), pMD2. G (Addegene #12259) and the plasmid of interest using Lipofectamine 2000 and 24 μg total DNA (with a ratio of 1:1:1). Lentiviruses were harvested 48 h after transfection and used directly on ALCL ALK(+) cells. A total of 10^6^ cells were transduced with 1 mL of lentiviral supernatant supplemented with 10 μg/mL polybren (Sigma, St Louis, MO, #TR-1003-G). Cells were spun-inoculated for 2 hours at 1000 rpm and 37°C, after which 1 mL of fresh medium was added. The transduced cells were then selected with puromycin (1 μg/mL).

## Supporting information

Supplemental Informations

## Acknowledgements.

The study was supported by the INSERM, Labex Toucan, Association Eva pour la vie, Féderation Grandir sans Cancer, RACCE et le Lions Club de Lourdes, Enfants Cancers Santé; Association Ellye, the Fondation les batisseurs d’étoiles et le marcheur au grand Coeur, the Fond Amgen France pour la science et l’humain, AAP3-2023, the French Ministry of Health and the French National Cancer lnstitute (CircOma, PRT-K 2022-184), Fondation ARC et Fondation pour la Recherche Médicale supported by its sponsor Nagui and his production company, Banijay Production (EQU202403018019). L.B was supported by Fondation de France, Fondation Toulouse Cancer Santé, Fondation l’Oréal for Women in Science, Prolific Graine de Chercheur and Association Ellye. L.C and S.D. were supported by the French Ministry of Health and the French National Cancer lnstitute (CircOma, PRT-K 2022-184) and the Fond Amgen France pour la science et l’humain, AAP3-2023. S.F. S.F. were supported by a grant from the French Ministry of Health and the French National Cancer lnstitute (CircOma, PRT-K 2022-184). S.F. was supported by a grant from the Deutsche Forschungsgemeinschaft (DFG, grant no. 439441203). S.F. is a participant in the BIH-Charité Clinician Scientist Program funded by the Charité - Universitätsmedizin Berlin and the Berlin Institute of Health. The authors thank the staff of the animal facility of UMS US006/CREFRE (Toulouse, France) and of the facilities of the Cancer Research Center of Toulouse (INSERM U1037) for their technical assistance.

Medical writing for this manuscript was assisted by MPIYP (MC Béné), Paris, France.

## Author contributions

L.B. and F.M. designed the research study. L.B. and L.C. performed the experiments. L.B. and F.M. analyzed the data. L.B and F.M wrote the paper. LL and SP helped with discussions. All authors have read and agreed to the submitted version of the manuscript.

