## Supplemental Table S1 for "*NPM1::ALK* fusion-derived circular RNAs drive aggressiveness in ALK-positive anaplastic large cell lymphoma"

**Antibodies**

| **Target** | **Provider** | **Ref** |
|---|---|---|
| ALK | Cell signaling technology | 3633 |
| STAT3 | Cell signaling technology | 9132 |
| pALK | Cell signaling technology | 3341 |
| pSTAT3 | Cell signaling technology | 9131 |
| GAPDH | Santa Cruz | sc-25778 |
| CXCL5 | Abcam | ab305100 |
| TIMP2 | Cell signaling technology | 5738 |
| Il1-alpha | Cell signaling technology | 34819 |

| **Primers** | **Sequence** |
|---|---|
| mRNA-NPM-ALK-F | ACCACCAGTGGTCTTAAGGTTG |
| mRNA-NPM-ALK-B | CGGAGCTTGCTCAGCTTGTA |
| circ-A26N3-F | GAGACATCTACAGGGCGAGC |
| circ-A26N3-B | CAGCCCCTAAACTGACCATG |
| circ-A23N2-F | CATTGTTCGCTGCATTGGGG |
| circ-A23N2-B | GCCTTTAGTTCACAACCGGG |
| MLN51-F | TAATCCCAGTTACCCTTATGCTCCA |
| MLN51-B | GTTATAGTAGGTCACTCCTCCATATACCTGT |
| circNPM-ALK-div-F | GCAGAGCCCTGAGTACAAGC |
| circNPM-ALK-div-B | AATATGCACTGGCCCTGAAC |
