## Supplemental Table S2 for "*NPM1::ALK* fusion-derived circular RNAs drive aggressiveness in ALK-positive anaplastic large cell lymphoma"

| A26N3_vs_CTL | | | | | |
|---|---|---|---|---|---|
| baseMean | log2FoldCha | lfcSE | pvalue | padj | gene_name |
| 26,6028342 | 7,38945094 | 1,26121952 | 4,66E-09 | 4,91E-07 | ANKRD44-IT1 |
| 30,0983051 | 7,35120593 | 1,25921787 | 5,29E-09 | 5,44E-07 | CXCR2P1 |
| 12,4843674 | 7,0938066 | 1,38058972 | 2,77E-07 | 1,72E-05 | ZNF14 |
| 10,8863141 | 7,01283116 | 1,39229831 | 4,73E-07 | 2,63E-05 | CXCR1 |
| 23,4057792 | 6,97040759 | 1,29058567 | 6,63E-08 | 4,83E-06 | DEFA1 |
| 22,7936235 | 6,9103588 | 1,2831136 | 7,22E-08 | 5,17E-06 | DEFA1B |
| 10,2670072 | 6,85215397 | 1,3596179 | 4,66E-07 | 2,61E-05 | ZSCAN18 |
| 17,966136 | 6,82635656 | 1,34058735 | 3,54E-07 | 2,11E-05 | LILRA1 |
| 9,58527933 | 6,76322837 | 1,43180372 | 2,32E-06 | 0,0001015 | RIN2 |
| 17,0683891 | 6,74191795 | 1,33175447 | 4,14E-07 | 2,36E-05 | ZNF841 |
| 17,58282 | 6,65898575 | 1,35696387 | 9,24E-07 | 4,64E-05 | MIR3667HG |
| 7,37496107 | 6,55818937 | 1,41132419 | 3,37E-06 | 0,00013741 | RHOBTB1 |
| 7,74144968 | 6,54332769 | 1,50112379 | 1,31E-05 | 0,00043765 | TRDC |
| 7,5233307 | 6,43759021 | 1,50108503 | 1,80E-05 | 0,00056507 | ZNF808 |
| 7,27837234 | 6,41640075 | 1,56311968 | 4,05E-05 | 0,00108573 | GUCY1A1 |
| 14,9186564 | 6,36075776 | 1,31512661 | 1,32E-06 | 6,34E-05 | FAM20C |
| 7,36730332 | 6,33983542 | 1,42359335 | 8,45E-06 | 0,00030568 | RIPK3 |
| 23,6424069 | 6,25704651 | 1,13248508 | 3,29E-08 | 2,67E-06 | LILRA5 |
| 6,32699139 | 6,25485579 | 1,48616647 | 2,57E-05 | 0,00076067 | S1PR5 |
| 11,9159202 | 6,2140934 | 1,32581292 | 2,77E-06 | 0,00011507 | DEFA3 |
| 5,3557092 | 6,14420389 | 1,70432211 | 0,00031207 | 0,00553384 | TCL1A |
| 5,35314679 | 6,07157006 | 1,66543972 | 0,00026674 | 0,00492873 | BTLA |
| 21,2170782 | 6,02953609 | 1,17146517 | 2,65E-07 | 1,65E-05 | CD52 |
| 5,18035615 | 6,02074236 | 1,55705694 | 0,0001103 | 0,00247561 | ZNF567 |
| 6,12345108 | 6,00031167 | 1,50522281 | 6,71E-05 | 0,00163617 | GIMAP7 |
| 5,0387662 | 5,96490775 | 1,4978713 | 6,83E-05 | 0,00165805 | ZNF600 |
| 5,43836045 | 5,94375112 | 1,45304642 | 4,30E-05 | 0,00114206 | ARHGEF34P |
| 11,0476049 | 5,943129 | 1,34905617 | 1,06E-05 | 0,00036591 | TRAT1 |
| 4,72201286 | 5,92918907 | 2,92261082 | 0,04248577 | 0,1815842 | PDZK1IP1 |
| 5,87892624 | 5,91018552 | 1,43845056 | 3,98E-05 | 0,00107084 | ARHGEF5 |
| 4,44123105 | 5,88760819 | 1,58247291 | 0,00019882 | 0,0039212 | ZNF10 |
| 4,24684091 | 5,83586051 | 1,56543666 | 0,00019305 | 0,00384087 | ZNF345 |
| 18,2056015 | 5,83099828 | 1,16008194 | 5,00E-07 | 2,76E-05 | P2RY2 |
| 5,88688433 | 5,82511331 | 1,44439979 | 5,51E-05 | 0,00140112 | LOC102723604 |
| 9,93149369 | 5,76086696 | 1,39465993 | 3,62E-05 | 0,00098852 | TBC1D9 |
| 4,04528986 | 5,75799406 | 1,83194645 | 0,00167168 | 0,0198306 | ZNF441 |
| 4,08745264 | 5,74119274 | 1,60033754 | 0,00033388 | 0,00585478 | ZNF585B |
| 10,6711953 | 5,72606208 | 1,29200839 | 9,34E-06 | 0,00033095 | LPCAT2 |
| 9,59478011 | 5,69571883 | 1,35819333 | 2,75E-05 | 0,0008017 | FFAR2 |
| 45,6605087 | 5,68023872 | 0,7646469 | 1,10E-13 | 4,95E-11 | TYROBP |
| 16,8853312 | 5,6762732 | 1,15303772 | 8,53E-07 | 4,34E-05 | CSF2RA |
| 24,1420852 | 5,66402468 | 1,02635637 | 3,42E-08 | 2,74E-06 | MX1 |
| 3,67410891 | 5,62411415 | 1,5818081 | 0,00037727 | 0,00642505 | CHRNA2 |
| 4,47207123 | 5,60732183 | 1,63423989 | 0,00060103 | 0,00911731 | ZNF320 |
| 29,1878039 | 5,58130968 | 0,90141138 | 5,95E-10 | 8,04E-08 | NELL2 |
| 3,49961714 | 5,56565934 | 1,65589665 | 0,00077628 | 0,0111575 | TMEM40 |

| | | | | | |
|---|---|---|---|---|---|
| 4,52757219 | 5,54959574 | 1,42871038 | 0,00010261 | 0,00233629 | S100Z |
| 7,30874536 | 5,54327007 | 1,47653799 | 0,00017387 | 0,00353365 | AHSP |
| 3,49314655 | 5,52881738 | 1,83717639 | 0,00261755 | 0,02737797 | CCDC149 |
| 3,33500172 | 5,4952393 | 1,63150394 | 0,00075659 | 0,01093622 | CEACAM3 |
| 4,48651975 | 5,49292479 | 1,65262619 | 0,00088814 | 0,01236536 | ZNF845 |
| 4,16957796 | 5,48409205 | 1,5508661 | 0,00040601 | 0,00684066 | LOC100507006 |
| 3,41873635 | 5,4829198 | 1,5572142 | 0,00042996 | 0,00714239 | AMIGO2 |
| 7,91398691 | 5,46939153 | 1,4325195 | 0,00013453 | 0,00286199 | PLCL1 |
| 10,947005 | 5,46433599 | 1,29623621 | 2,49E-05 | 0,00074248 | ARRDC4 |
| 34,8332919 | 5,43443269 | 0,8188528 | 3,21E-11 | 6,98E-09 | PF4 |
| 14,1110468 | 5,42780734 | 1,19837116 | 5,92E-06 | 0,00022303 | IGHD |
| 14,8186717 | 5,42636477 | 1,23880501 | 1,19E-05 | 0,000404 | ZNF831 |
| 3,90900184 | 5,41986678 | 1,74785251 | 0,00192952 | 0,02191884 | ZNF571-AS1 |
| 3,19448127 | 5,40861576 | 1,74307674 | 0,00191626 | 0,02182104 | ESPN |
| 14,4474642 | 5,39974356 | 1,73883535 | 0,00190035 | 0,02171674 | CXCL5 |
| 6,6092085 | 5,39174701 | 1,42250911 | 0,00015046 | 0,00314189 | ZNF493 |
| 7,68821436 | 5,37423462 | 1,37040351 | 8,79E-05 | 0,00205437 | GIMAP4 |
| 3,15460868 | 5,35690094 | 1,58726673 | 0,00073837 | 0,01070547 | TSPAN2 |
| 3,33970339 | 5,34935477 | 1,61756489 | 0,00094287 | 0,01291524 | PNPLA4 |
| 2,81065374 | 5,34610362 | 1,87913915 | 0,00444149 | 0,03990384 | DEFA4 |
| 4,32669684 | 5,33487372 | 1,71690387 | 0,00188828 | 0,0216373 | LRRC18 |
| 3,6260218 | 5,32948976 | 1,56072312 | 0,00063839 | 0,00955698 | SERPINB9-AS1 |
| 2,89292788 | 5,31483574 | 1,72248582 | 0,00203168 | 0,02271336 | ARHGEF35-AS1 |
| 5,92573205 | 5,31320453 | 1,60567705 | 0,00093626 | 0,01285555 | FCRL2 |
| 3,36062375 | 5,27708085 | 1,65980738 | 0,00147614 | 0,01805436 | MLYCD |
| 6,49974871 | 5,26265202 | 1,47420016 | 0,00035721 | 0,00616668 | ZNF469 |
| 61,5673077 | 5,24917246 | 0,64656324 | 4,72E-16 | 5,25E-13 | PLXND1 |
| 3,39360075 | 5,23542808 | 1,62300899 | 0,00125641 | 0,01596546 | VENTX |
| 6,29877675 | 5,23318107 | 1,53968375 | 0,00067665 | 0,01001799 | ZNF350 |
| 2,85856404 | 5,23289887 | 1,67419596 | 0,00177431 | 0,02072331 | LANCL3 |
| 2,68615225 | 5,22084462 | 1,74022079 | 0,00269887 | 0,02805785 | TRAJ8 |
| 17,8095463 | 5,21954765 | 1,09154156 | 1,74E-06 | 7,98E-05 | AFF3 |
| 3,01743643 | 5,19533012 | 1,74109285 | 0,00284555 | 0,02911883 | ZNF540 |
| 4,10078605 | 5,18472193 | 1,9316578 | 0,00727299 | 0,05622275 | ZNF799 |
| 2,98828053 | 5,18257422 | 1,73295415 | 0,00278429 | 0,02865771 | KNDC1 |
| 35,1932898 | 5,17511212 | 1,21910241 | 2,19E-05 | 0,00066425 | IGKC |
| 5,43344176 | 5,17271151 | 1,48830413 | 0,00050976 | 0,0081052 | RNASE6 |
| 5,19290026 | 5,16824738 | 1,47867747 | 0,00047374 | 0,00767431 | LOC388436 |
| 2,52197945 | 5,16744209 | 1,81350783 | 0,00437993 | 0,03948195 | POGLUT3 |
| 2,78801106 | 5,16294174 | 1,73100156 | 0,00285781 | 0,02921263 | ANTXRLP1 |
| 2,76031675 | 5,15173345 | 1,8676015 | 0,00580716 | 0,04825505 | CLC |
| 2,93611258 | 5,14211641 | 1,6688298 | 0,00206119 | 0,02290065 | MRPL23-AS1 |
| 22,2113289 | 5,12817215 | 0,86773958 | 3,43E-09 | 3,77E-07 | LTF |
| 2,43193173 | 5,12200171 | 1,81271999 | 0,00471938 | 0,04182451 | ZNF667-AS1 |
| 3,45646314 | 5,09626016 | 1,58516891 | 0,00130461 | 0,01639457 | KCNJ2 |
| 3,51384728 | 5,07732667 | 1,55749537 | 0,0011144 | 0,01455174 | PIWIL4 |
| 11,0017028 | 5,07694691 | 1,21842229 | 3,09E-05 | 0,00088016 | LINC00989 |
| 3,50287543 | 5,07052255 | 1,62827487 | 0,00184541 | 0,02130383 | IGHG3 |

| | | | | | |
|---|---|---|---|---|---|
| 9,18398045 | 5,05928232 | 1,19922467 | 2,46E-05 | 0,000733 | DSC2 |
| 21,5222585 | 5,05329468 | 1,00520553 | 4,98E-07 | 2,75E-05 | WWC3 |
| 3,85444031 | 5,05090182 | 1,7092197 | 0,00312576 | 0,03086766 | HID1 |
| 2,57969021 | 5,04924424 | 1,9278419 | 0,00881576 | 0,06388472 | SIRPG-AS1 |
| 2,43280281 | 5,04539873 | 1,98038815 | 0,01084414 | 0,07312255 | ELANE |
| 2,99267709 | 5,04414557 | 1,58034269 | 0,00141387 | 0,01749627 | TRGC2 |
| 3,55210601 | 5,0323515 | 1,77475419 | 0,00457511 | 0,04069848 | CLEC1B |
| 15,7310724 | 5,01703395 | 1,51935962 | 0,00095974 | 0,01306421 | SIGLEC5 |
| 2,34733726 | 5,01397756 | 1,75031726 | 0,00417522 | 0,03825137 | SOX13 |
| 2,56088496 | 5,0026628 | 1,88010076 | 0,00779441 | 0,05870939 | CR2 |
| 15,4297703 | 4,99103794 | 1,07060506 | 3,13E-06 | 0,00012888 | IGHG1 |
| 127,839999 | 4,98221355 | 0,63064767 | 2,79E-15 | 2,20E-12 | CYBB |
| 2,54504218 | 4,97263546 | 2,14466324 | 0,02041625 | 0,11190493 | ZNF667 |
| 4,94943716 | 4,96852116 | 1,44130703 | 0,00056636 | 0,00876737 | CYSLTR1 |
| 4,98957223 | 4,95834332 | 1,61651885 | 0,00216004 | 0,02365171 | ZNF701 |
| 6,52671148 | 4,95350078 | 1,4912031 | 0,00089434 | 0,01239242 | LY75 |
| 3,17810891 | 4,9351396 | 1,7578768 | 0,00499365 | 0,04340136 | KLRC4-KLRK1 |
| 2,76428287 | 4,92619176 | 1,63193253 | 0,00253925 | 0,02669174 | LY6G6E |
| 2,23928019 | 4,91922057 | 1,87874261 | 0,00883541 | 0,06395362 | TRPC2 |
| 3,20966612 | 4,91836922 | 1,7076123 | 0,00397346 | 0,0368252 | MSR1 |
| 4,5416146 | 4,9173645 | 1,69782023 | 0,00377614 | 0,03544594 | FCRL3 |
| 10,6809816 | 4,90868229 | 1,17974541 | 3,17E-05 | 0,00089636 | ZNF160 |
| 2,56007421 | 4,90812002 | 1,8155248 | 0,00686307 | 0,0540519 | ZNF888 |
| 60,1545178 | 4,89315912 | 0,54324213 | 2,11E-19 | 4,99E-16 | NRIP1 |
| 2,66589808 | 4,88803033 | 1,90948523 | 0,01047119 | 0,07144803 | FCRL5 |
| 29,1951395 | 4,88801138 | 0,79602685 | 8,23E-10 | 1,09E-07 | ABCB1 |
| 14,7430945 | 4,87995766 | 1,01916524 | 1,68E-06 | 7,84E-05 | MME |
| 24,433054 | 4,86713275 | 0,82007653 | 2,94E-09 | 3,31E-07 | CD28 |
| 25,0509663 | 4,85793207 | 0,8314632 | 5,14E-09 | 5,34E-07 | SIGLEC14 |
| 10,6405626 | 4,85760177 | 1,27292718 | 0,00013558 | 0,002876 | SELP |
| 279,112569 | 4,85651116 | 0,57537021 | 3,16E-17 | 5,43E-14 | XIST |
| 4,38954602 | 4,85298779 | 1,74646769 | 0,00545694 | 0,04621789 | CABP5 |
| 13,3252334 | 4,84847348 | 1,17659684 | 3,78E-05 | 0,00102802 | GRAP |
| 65,3099555 | 4,84794377 | 0,51476541 | 4,61E-21 | 1,25E-17 | CLEC7A |
| 26,6745615 | 4,83697968 | 0,84214989 | 9,27E-09 | 8,78E-07 | MNDA |
| 3,03161748 | 4,83400801 | 1,69145777 | 0,00426462 | 0,03877934 | CA4 |
| 2,5449939 | 4,83250969 | 1,82838614 | 0,00821637 | 0,06089385 | HDGFL3 |
| 2,55780605 | 4,83068069 | 1,8863872 | 0,01044282 | 0,07130589 | LOC105373780 |
| 5,09194421 | 4,82254831 | 1,57832248 | 0,00224693 | 0,02441942 | CD8B |
| 1,94319383 | 4,81925097 | 2,13851378 | 0,0242244 | 0,12499313 | LINC00965 |
| 2,28403976 | 4,81823742 | 2,07934726 | 0,02049352 | 0,11210116 | ZNF563 |
| 3,94426266 | 4,80384287 | 1,53511103 | 0,00175216 | 0,0205025 | GRAPL-AS1 |
| 2,52215382 | 4,80340374 | 1,90515409 | 0,01169328 | 0,07716087 | LILRA4 |
| 2,47161731 | 4,79814198 | 1,90823829 | 0,01192231 | 0,07827276 | ZNF571 |
| 21,6578728 | 4,7954697 | 0,88794144 | 6,64E-08 | 4,83E-06 | ARAP3 |
| 2,39256234 | 4,79140128 | 1,95567771 | 0,01428581 | 0,08841893 | TRAJ38 |
| 2,38241689 | 4,78975092 | 1,70688541 | 0,00501397 | 0,04353799 | ZFP30 |
| 2,2931384 | 4,78708406 | 1,80897403 | 0,00813781 | 0,06048764 | TMEM176B |

| | | | | | |
|---|---|---|---|---|---|
| 2,30484974 | 4,77186033 | 1,86624367 | 0,01055995 | 0,07174321 | AGR2 |
| 3,10649876 | 4,76972329 | 1,59148106 | 0,0027262 | 0,02825102 | MIR4298 |
| 4,43462783 | 4,76605359 | 1,47716541 | 0,0012532 | 0,01596071 | QPCT |
| 2,78361647 | 4,76573254 | 1,74497976 | 0,00631213 | 0,05112664 | SAXO5 |
| 2,48978902 | 4,76536534 | 1,9986338 | 0,01711112 | 0,09945292 | ZNF420 |
| 49,6929676 | 4,75189455 | 0,66042014 | 6,23E-13 | 2,36E-10 | CD69 |
| 193,161192 | 4,75144462 | 0,73258938 | 8,82E-11 | 1,70E-08 | TNFAIP2 |
| 43,8498002 | 4,73251702 | 0,9141854 | 2,26E-07 | 1,44E-05 | SNX9 |
| 3,95604586 | 4,7300398 | 1,66567176 | 0,00451533 | 0,04031832 | TARP |
| 17,0869804 | 4,71596051 | 1,48319602 | 0,00147482 | 0,01804982 | TP53I11 |
| 2,26030284 | 4,71216716 | 1,93638154 | 0,01495401 | 0,09133142 | ZNF891 |
| 16,5959708 | 4,71189124 | 0,96785542 | 1,13E-06 | 5,50E-05 | CD1D |
| 1,96964368 | 4,70950389 | 1,88323761 | 0,01239311 | 0,08028565 | SLC16A10 |
| 7,32556805 | 4,70853842 | 1,36619564 | 0,00056798 | 0,00877731 | LINC02520 |
| 1,8221504 | 4,70838876 | 2,12153775 | 0,0264644 | 0,13260934 | SALRNA2 |
| 17,3313436 | 4,70786954 | 0,9885621 | 1,91E-06 | 8,66E-05 | ARSI |
| 1,80455686 | 4,70491434 | 2,30423526 | 0,04116591 | 0,17783402 | VSTM1 |
| 18,1409653 | 4,69906391 | 0,91183956 | 2,56E-07 | 1,61E-05 | TLR4 |
| 17,3990357 | 4,69646412 | 0,99267091 | 2,23E-06 | 9,85E-05 | CXCR2 |
| 24,7476276 | 4,69479841 | 0,74576692 | 3,07E-10 | 4,76E-08 | IRAG2 |
| 9,04803315 | 4,69161498 | 1,32928847 | 0,00041647 | 0,00697355 | TPTEP1 |
| 3,92966306 | 4,68886129 | 1,64417244 | 0,00434716 | 0,03928013 | PCYT1B |
| 167,137796 | 4,68746603 | 0,67578685 | 4,02E-12 | 1,14E-09 | F13A1 |
| 25,0049782 | 4,68627922 | 0,80779728 | 6,58E-09 | 6,62E-07 | MS4A6A |
| 29,3464939 | 4,67902758 | 0,79821583 | 4,58E-09 | 4,87E-07 | FCMR |
| 12,8705838 | 4,67177565 | 0,98716393 | 2,22E-06 | 9,83E-05 | JCHAIN |
| 2,05371579 | 4,6431883 | 2,06486777 | 0,02453405 | 0,126113 | LOC90246 |
| 54,5422274 | 4,64158566 | 0,60386044 | 1,51E-14 | 8,70E-12 | IRS2 |
| 1,72157657 | 4,63717198 | 2,20449188 | 0,03542116 | 0,1606685 | ALDH1A1 |
| 4,079711 | 4,63625757 | 1,65668627 | 0,00513386 | 0,04425406 | CCNJL |
| 2,49357293 | 4,62949058 | 1,90635322 | 0,01516336 | 0,09219342 | ZNF717 |
| 3,864398 | 4,61981209 | 1,59936771 | 0,00387054 | 0,03609392 | GCSAML |
| 2,04279792 | 4,61899304 | 2,13030818 | 0,03014137 | 0,14438099 | RPH3A |
| 4,05675715 | 4,61372882 | 1,54676892 | 0,00285609 | 0,02921081 | LGALS12 |
| 10,8279194 | 4,60962618 | 1,17035296 | 8,19E-05 | 0,00194277 | CLEC17A |
| 8,37852856 | 4,59781168 | 1,39402038 | 0,00097293 | 0,01319634 | IGLC2 |
| 1,89213908 | 4,59709636 | 2,33498733 | 0,04897738 | 0,19997776 | VAV3-AS1 |
| 10,7529123 | 4,59597526 | 1,10396149 | 3,14E-05 | 0,00089042 | JUP |
| 3,74772646 | 4,59595086 | 1,52524926 | 0,0025847 | 0,0270642 | ILDR2 |
| 3,75874693 | 4,59537562 | 1,58345266 | 0,00370642 | 0,03497715 | GIMAP1 |
| 4,86786663 | 4,58959518 | 1,58145705 | 0,00370634 | 0,03497715 | C13orf46 |
| 11,4559417 | 4,58692198 | 1,15564974 | 7,21E-05 | 0,00174538 | LILRB4 |
| 19,8190953 | 4,58504267 | 0,81594216 | 1,92E-08 | 1,66E-06 | FGD4 |
| 1,96208665 | 4,57928246 | 1,89770682 | 0,01581917 | 0,09468981 | CDKL5 |
| 4,06459913 | 4,5735815 | 1,53866038 | 0,00295439 | 0,02971657 | ANKRD36BP2 |
| 3,87807806 | 4,57203762 | 1,67775598 | 0,00642838 | 0,05170547 | HTATIP2 |
| 2,38431688 | 4,56914522 | 1,88994925 | 0,01562315 | 0,09411196 | TRAJ22 |
| 10,9044664 | 4,56891602 | 1,26097801 | 0,00029086 | 0,00525121 | IL24 |

| | | | | | |
|---|---|---|---|---|---|
| 30,0876615 | 4,56734055 | 0,71001134 | 1,25E-10 | 2,32E-08 | IGHA1 |
| 3,30052147 | 4,56646636 | 1,66708292 | 0,00615897 | 0,05033861 | FBXO40 |
| 1,74358607 | 4,56376664 | 1,94644385 | 0,01904398 | 0,10673314 | LOC101929098 |
| 185,243184 | 4,5482158 | 0,62146821 | 2,51E-13 | 1,01E-10 | COTL1 |
| 3,56515201 | 4,54210866 | 1,85354991 | 0,01426613 | 0,08832767 | PDK4 |
| 3,64547989 | 4,54185247 | 1,52644243 | 0,00292559 | 0,02958095 | ZFP14 |
| 20,2030734 | 4,54013312 | 0,92292238 | 8,69E-07 | 4,41E-05 | XK |
| 15,8204665 | 4,53450686 | 1,47080072 | 0,00204912 | 0,02283356 | PF4V1 |
| 18,763978 | 4,53305225 | 0,91046429 | 6,40E-07 | 3,42E-05 | PHACTR2 |
| 4,13781418 | 4,52408002 | 1,67665321 | 0,00696988 | 0,05463009 | SHISAL2A |
| 37,1761933 | 4,52347216 | 0,69593309 | 8,04E-11 | 1,58E-08 | P2RX1 |
| 1,96917576 | 4,52325728 | 2,18382102 | 0,03833467 | 0,1696679 | IGHV3-23 |
| 8,89631398 | 4,51992722 | 1,43105405 | 0,00158605 | 0,01908826 | TTC9 |
| 1,90412871 | 4,51885784 | 1,82006572 | 0,01303545 | 0,08281771 | ETNK2 |
| 35,3108527 | 4,50947895 | 0,68798041 | 5,58E-11 | 1,15E-08 | CD7 |
| 1,75834182 | 4,50420701 | 2,1646265 | 0,03745001 | 0,16716009 | ECHDC3 |
| 2,87014869 | 4,49850172 | 1,67277437 | 0,00716136 | 0,05564688 | ZNF528 |
| 10,7724466 | 4,49695828 | 1,04932772 | 1,82E-05 | 0,00057066 | PCED1B-AS1 |
| 2,5943172 | 4,48370786 | 2,04329609 | 0,02821022 | 0,13849651 | ZNF793 |
| 67,5363835 | 4,48233389 | 0,76391543 | 4,42E-09 | 4,73E-07 | ELF4 |
| 13,373614 | 4,46557959 | 1,07330024 | 3,17E-05 | 0,00089636 | CD244 |
| 2,12944174 | 4,46346264 | 1,85042663 | 0,01585979 | 0,09479174 | CD200 |
| 1,91902668 | 4,45863774 | 2,10495087 | 0,03416092 | 0,15676903 | LARGE1 |
| 3,8431184 | 4,45609277 | 1,4900902 | 0,00278535 | 0,02865771 | CD79B |
| 1,6330762 | 4,45552014 | 2,03439247 | 0,02851709 | 0,13956849 | LINC00391 |
| 17,5678626 | 4,45372145 | 0,91137784 | 1,02E-06 | 5,04E-05 | GPRASP1 |
| 30,5255803 | 4,44865421 | 0,7460001 | 2,47E-09 | 2,85E-07 | TES |
| 3,54494902 | 4,44772346 | 1,48567153 | 0,00275575 | 0,0284771 | FAM43A |
| 7,46038628 | 4,44185801 | 1,27431213 | 0,00049087 | 0,00791005 | COL6A2 |
| 4,65316579 | 4,44155768 | 1,49054757 | 0,00288421 | 0,02937149 | NEURL1 |
| 20,6102805 | 4,44069609 | 0,83556219 | 1,07E-07 | 7,41E-06 | ADAMTS10 |
| 1,90049902 | 4,44063587 | 1,88823448 | 0,01868585 | 0,1055073 | LOC105375112 |
| 3,01271584 | 4,44055499 | 1,77364809 | 0,01229272 | 0,07979094 | MIR3670-4 |
| 33,3964397 | 4,43957896 | 0,72518956 | 9,24E-10 | 1,19E-07 | CD3E |
| 43,7305298 | 4,43738337 | 0,86895799 | 3,28E-07 | 1,98E-05 | LRRC25 |
| 4,53915042 | 4,4353054 | 1,69074399 | 0,00870861 | 0,06334882 | ZNF615 |
| 1,91967195 | 4,43378128 | 1,83500772 | 0,01568261 | 0,09426005 | CFAP46 |
| 1,9162712 | 4,42622906 | 1,74254588 | 0,01108229 | 0,0742635 | ZNF486 |
| 5,77551846 | 4,42459662 | 1,3698455 | 0,00123792 | 0,0158154 | GP9 |
| 1,84913713 | 4,42385343 | 2,00511206 | 0,02736388 | 0,13584218 | NMUR1 |
| 10,0079216 | 4,4211192 | 1,16116058 | 0,00014038 | 0,0029611 | ACSM1 |
| 1,9843444 | 4,42069301 | 1,81705172 | 0,01497871 | 0,09140321 | CCL4 |
| 2,36253373 | 4,41902001 | 1,75853168 | 0,01197422 | 0,07850455 | VSIG1 |
| 16,1820066 | 4,41170094 | 0,9615212 | 4,47E-06 | 0,0001751 | IGHA2 |
| 16,1234573 | 4,40769037 | 0,8628553 | 3,25E-07 | 1,97E-05 | DOK2 |
| 16,333096 | 4,39032374 | 0,98039798 | 7,53E-06 | 0,00027549 | LY9 |
| 1,80350311 | 4,38367495 | 2,14682131 | 0,04115773 | 0,17783402 | TRAJ18 |
| 21,2351808 | 4,37966457 | 0,87983391 | 6,43E-07 | 3,43E-05 | MS4A1 |

| | | | | | |
|---|---|---|---|---|---|
| 1,44622021 | 4,37205532 | 2,05010092 | 0,03295715 | NA | FCN2 |
| 10,8277441 | 4,37068736 | 1,0950107 | 6,57E-05 | 0,00160948 | RNF175 |
| 1,61251402 | 4,36454954 | 2,1567644 | 0,04300554 | NA | LOC101928381 |
| 4,51576212 | 4,36425368 | 1,44668521 | 0,0025552 | 0,02682962 | ZNF253 |
| 6,57814022 | 4,35838321 | 1,46232899 | 0,00287835 | 0,02932755 | ZNF141 |
| 69,5909647 | 4,3583696 | 0,45650977 | 1,33E-21 | 4,20E-18 | LRRK2 |
| 6,35015254 | 4,35563135 | 1,22659621 | 0,00038378 | 0,00652434 | GUCY1B1 |
| 30,157293 | 4,35372688 | 0,76129371 | 1,07E-08 | 9,80E-07 | EPB41L3 |
| 117,615809 | 4,35194362 | 0,67074119 | 8,68E-11 | 1,69E-08 | SLC4A1 |
| 3,46471552 | 4,34920359 | 1,66088708 | 0,00882907 | 0,06393218 | ZNF28 |
| 1,49683524 | 4,33837398 | 2,1932845 | 0,04792576 | NA | CBR3 |
| 30,0616263 | 4,32755421 | 0,69495538 | 4,75E-10 | 6,77E-08 | SNX19 |
| 367,577413 | 4,32684418 | 0,63996267 | 1,37E-11 | 3,32E-09 | LYZ |
| 4,42152949 | 4,315945 | 1,45637716 | 0,00304182 | 0,03032425 | BEGAIN |
| 50,6366336 | 4,31504951 | 0,57386564 | 5,51E-14 | 2,61E-11 | TC2N |
| 6,38733931 | 4,31194817 | 1,48283881 | 0,00363864 | 0,03459551 | S1PR3 |
| 2,01199052 | 4,30081331 | 2,10114823 | 0,04066917 | 0,17625078 | LINC02692 |
| 7,14364487 | 4,29117924 | 1,28979379 | 0,00087778 | 0,0122662 | ZNF100 |
| 32,3424811 | 4,28827891 | 0,71796755 | 2,33E-09 | 2,71E-07 | FCGR3B |
| 168,743955 | 4,28704239 | 0,61583316 | 3,37E-12 | 9,81E-10 | DOCK8 |
| 87,5732272 | 4,28676518 | 0,65624975 | 6,48E-11 | 1,30E-08 | CHST15 |
| 3,31709333 | 4,28610056 | 1,54072837 | 0,00540473 | 0,04584623 | SLAMF6 |
| 1,83388533 | 4,28350335 | 1,87791622 | 0,02254919 | 0,11901066 | SYNGR1 |
| 1,91596776 | 4,2732531 | 1,87401464 | 0,02259188 | 0,11920271 | PCDH9 |
| 37,7851513 | 4,26815478 | 0,67585289 | 2,70E-10 | 4,40E-08 | LILRB2 |
| 3,37448039 | 4,26730613 | 1,76166543 | 0,01542204 | 0,09316741 | ZNF738 |
| 50,9764519 | 4,26607464 | 0,6132608 | 3,49E-12 | 1,00E-09 | IGHM |
| 1,51873352 | 4,26559659 | 1,99087397 | 0,03214725 | NA | MS4A14 |
| 55,0362225 | 4,26416147 | 0,78467751 | 5,50E-08 | 4,11E-06 | HBB |
| 198,037879 | 4,25505322 | 0,73741462 | 7,92E-09 | 7,74E-07 | PPBP |
| 63,8898494 | 4,25458009 | 0,72172976 | 3,75E-09 | 4,10E-07 | GYPC |
| 2,32957374 | 4,25057077 | 1,92401975 | 0,02715978 | 0,13526987 | KCNN1 |
| 1,57846389 | 4,2494861 | 1,97508219 | 0,03143289 | NA | TRAJ13 |
| 1,49537453 | 4,24938969 | 2,10461577 | 0,0434788 | NA | ZNF222 |
| 1,63478682 | 4,24429738 | 2,12985961 | 0,04628782 | 0,19286761 | TRAJ11 |
| 3,21412666 | 4,24353923 | 1,7621837 | 0,01603517 | 0,09564473 | TRAJ7 |
| 4,3186748 | 4,23651289 | 1,6532363 | 0,01039044 | 0,07107648 | MIR6817 |
| 18,8422783 | 4,23568556 | 0,85627539 | 7,55E-07 | 3,90E-05 | GNG11 |
| 1,99673917 | 4,23345817 | 1,78808761 | 0,01790437 | 0,10241815 | C1orf167 |
| 1,3932648 | 4,21892405 | 1,99178052 | 0,03416091 | NA | RGSL1 |
| 3,17764031 | 4,21388792 | 1,92840705 | 0,02887674 | 0,14055523 | CLEC12B |
| 20,5657157 | 4,21111286 | 0,83816328 | 5,05E-07 | 2,77E-05 | PLEKHG3 |
| 2,97663483 | 4,20459394 | 1,70325789 | 0,0135658 | 0,08527524 | HLA-DQB2 |
| 6,5907731 | 4,20269017 | 1,30575766 | 0,00128825 | 0,01623921 | DAPK2 |
| 62,0192045 | 4,20246417 | 0,49628439 | 2,50E-17 | 4,73E-14 | LFNG |
| 8,90864677 | 4,20146556 | 1,03776059 | 5,15E-05 | 0,00133008 | MS4A7 |
| 14,2176901 | 4,19914621 | 0,9774033 | 1,74E-05 | 0,00054877 | SIRPB2 |
| 71,4323172 | 4,19813438 | 0,78481808 | 8,84E-08 | 6,19E-06 | GRAP2 |

| | | | | | |
|---|---|---|---|---|---|
| 139,829307 | 4,19661135 | 0,38344201 | 7,05E-28 | 6,53E-24 | ANKRD44 |
| 2,69316804 | 4,1957038 | 2,05499424 | 0,04118023 | 0,17785236 | LOC100134868 |
| 3,08667548 | 4,19158843 | 1,94646741 | 0,03128461 | 0,14809511 | SYDE2 |
| 5,26916977 | 4,18567715 | 1,36221515 | 0,00212132 | 0,02337656 | ZBP1 |
| 5,61565026 | 4,18114592 | 1,38021104 | 0,00245078 | 0,02597832 | IGHG2 |
| 3,59336776 | 4,18027033 | 1,57903575 | 0,00811238 | 0,06037398 | SEC14L5 |
| 2,50233035 | 4,1798767 | 1,97804017 | 0,03458875 | 0,15800427 | CCR3 |
| 1,75326618 | 4,17840996 | 2,07980089 | 0,04453255 | 0,18774516 | GOLGA8CP |
| 1,7749549 | 4,1767368 | 2,06459631 | 0,04307023 | 0,18358758 | CLEC1A |
| 7,6035577 | 4,17658006 | 1,27944736 | 0,00109711 | 0,01440583 | ZNF765 |
| 30,1688632 | 4,17030869 | 0,748379 | 2,51E-08 | 2,10E-06 | NUAK2 |
| 211,652708 | 4,16490232 | 0,66256673 | 3,26E-10 | 4,97E-08 | VSIR |
| 2,97472423 | 4,1642318 | 1,6735285 | 0,01283574 | 0,08197731 | LINC01278 |
| 13,9088289 | 4,15669338 | 0,85733621 | 1,24E-06 | 6,02E-05 | AATK |
| 1,59485069 | 4,1524023 | 2,04866778 | 0,04267476 | NA | TSPYL5 |
| 60,8740772 | 4,1514523 | 0,53810671 | 1,21E-14 | 7,39E-12 | ABCG1 |
| 22,9047187 | 4,14860892 | 0,7332062 | 1,53E-08 | 1,35E-06 | CD300LF |
| 8,97342518 | 4,14581767 | 1,11870956 | 0,00021065 | 0,00409626 | MATK |
| 11,5042094 | 4,14128335 | 0,99575181 | 3,20E-05 | 0,00090153 | TNNT3 |
| 16,0705403 | 4,14067861 | 0,90339278 | 4,57E-06 | 0,00017804 | FBP1 |
| 3,53822105 | 4,13002517 | 1,75806166 | 0,01881418 | 0,10578875 | CEACAM4 |
| 2,80730802 | 4,12006103 | 1,9690308 | 0,03639998 | 0,16370906 | MIR4420 |
| 9,08717271 | 4,11637633 | 1,10708974 | 0,00020065 | 0,00393829 | GAB1 |
| 1,49024233 | 4,11174767 | 2,06739274 | 0,04671703 | NA | CCL4L2 |
| 3,59338941 | 4,10818 | 1,61133292 | 0,01078609 | 0,07288698 | VNN1 |
| 3,2744634 | 4,10599702 | 1,63672559 | 0,01211882 | 0,07917823 | ZNF585A |
| 160,810889 | 4,10453478 | 0,65899662 | 4,71E-10 | 6,77E-08 | RASA3 |
| 21,9477883 | 4,09982785 | 0,85565877 | 1,66E-06 | 7,76E-05 | RGCC |
| 118,130137 | 4,09943491 | 0,5834055 | 2,11E-12 | 6,56E-10 | CAVIN2 |
| 20,9332379 | 4,09862297 | 0,80537362 | 3,60E-07 | 2,13E-05 | CKAP4 |
| 4,18243869 | 4,09585037 | 1,59300149 | 0,01013606 | 0,06994323 | C5AR2 |
| 2,68360246 | 4,09335005 | 1,67360104 | 0,01445176 | 0,08912703 | EPHB1 |
| 10,4638028 | 4,09233559 | 1,16415766 | 0,0004393 | 0,00724668 | PADI4 |
| 14,6090275 | 4,07748902 | 0,9255192 | 1,05E-05 | 0,00036591 | IFI44L |
| 1,36910754 | 4,07610408 | 2,03631489 | 0,04531634 | NA | ARMCX5 |
| 1,35794353 | 4,07513497 | 2,04155447 | 0,04592367 | NA | IGKV3-11 |
| 2,3247531 | 4,07444623 | 1,85940711 | 0,02843291 | 0,1392646 | CHRM3-AS2 |
| 3,08544231 | 4,07256297 | 1,83833919 | 0,02673612 | 0,13358174 | SALL2 |
| 2,50752876 | 4,07246637 | 1,93846867 | 0,03565257 | 0,16130616 | SLC4A4 |
| 1,86248864 | 4,07213232 | 2,0176844 | 0,04356829 | 0,18483308 | ZNF829 |
| 1,54897868 | 4,07117958 | 2,03881897 | 0,0458434 | NA | MIR24-2 |
| 5,07535863 | 4,07087239 | 1,40421566 | 0,00374311 | 0,03525309 | PLCB1 |
| 37,6501806 | 4,06940989 | 0,63345826 | 1,33E-10 | 2,44E-08 | ZNF331 |
| 43,1216451 | 4,06764369 | 0,53716759 | 3,66E-14 | 1,84E-11 | CYRIA |
| 2,88825747 | 4,05756304 | 1,67235937 | 0,01525573 | 0,09246376 | ZNF71 |
| 42,475832 | 4,05505568 | 0,63908129 | 2,22E-10 | 3,75E-08 | RCSD1 |
| 3,00712823 | 4,05435222 | 1,75227816 | 0,02068086 | 0,112816 | LDLRAD4-AS1 |
| 4,38926803 | 4,05200896 | 1,44667527 | 0,00509585 | 0,04400669 | ZNF558 |

| | | | | | |
|---|---|---|---|---|---|
| 20,3715143 | 4,04643487 | 0,70462674 | 9,32E-09 | 8,78E-07 | RAB37 |
| 22,3573437 | 4,04529568 | 0,7817875 | 2,29E-07 | 1,46E-05 | FCGR3A |
| 146,28344 | 4,04005392 | 0,57491303 | 2,11E-12 | 6,56E-10 | FGR |
| 63,5426922 | 4,03505576 | 0,52498369 | 1,52E-14 | 8,70E-12 | IRAK3 |
| 3,46782621 | 4,03343876 | 1,72234469 | 0,01918944 | 0,10726107 | ZNF844 |
| 7,74872294 | 4,0300684 | 1,1964632 | 0,00075628 | 0,01093622 | HPSE |
| 3,58604696 | 4,02755149 | 1,75593722 | 0,0218091 | 0,11663537 | P2RY1 |
| 21,182304 | 4,02527094 | 0,80745207 | 6,19E-07 | 3,32E-05 | CA2 |
| 19,2047124 | 4,02356925 | 0,81735898 | 8,54E-07 | 4,34E-05 | CD86 |
| 1,51514656 | 4,0225939 | 2,00745282 | 0,04508829 | NA | PLA2R1 |
| 29,4760304 | 4,02099776 | 0,72709686 | 3,20E-08 | 2,62E-06 | CPVL |
| 2,60642333 | 4,01325806 | 1,95876092 | 0,04047425 | 0,17572586 | COL5A3 |
| 68,5193962 | 4,00973792 | 0,48574249 | 1,52E-16 | 1,92E-13 | SYK |
| 2,27553504 | 4,00145205 | 1,89877755 | 0,03508437 | 0,15942156 | KIR2DL3 |
| 6,45997498 | 3,99531855 | 1,2511892 | 0,00140697 | 0,01742233 | TBX21 |
| 11,087234 | 3,98731492 | 0,95585731 | 3,03E-05 | 0,00086375 | PELATON |
| 75,7226034 | 3,98427357 | 0,49698188 | 1,08E-15 | 1,08E-12 | PLEK |
| 3,3007027 | 3,98232272 | 1,63697057 | 0,01498508 | 0,09140321 | EIF5A2 |
| 16,3663487 | 3,97057682 | 0,91403377 | 1,40E-05 | 0,00046116 | ANKRD33B |
| 5,53613039 | 3,96941113 | 1,33341228 | 0,00291196 | 0,0294795 | ADGRE3 |
| 2,98192166 | 3,96329702 | 1,59440552 | 0,0129278 | 0,08238694 | ZNF529-AS1 |
| 2,91682598 | 3,96014029 | 1,7285838 | 0,02196493 | 0,11710295 | ZC3HAV1L |
| 111,458816 | 3,95695533 | 0,68808895 | 8,89E-09 | 8,54E-07 | PIK3R5 |
| 2,20686619 | 3,9543281 | 1,84933483 | 0,03249699 | 0,15163392 | LOC101928850 |
| 20,6983161 | 3,95361879 | 0,72481001 | 4,91E-08 | 3,73E-06 | SLC15A3 |
| 18,7954339 | 3,95166844 | 0,76055927 | 2,04E-07 | 1,31E-05 | SH2D1A |
| 15,1805593 | 3,95123987 | 0,82571458 | 1,71E-06 | 7,88E-05 | CAMK2A |
| 209,343295 | 3,94913158 | 0,55702088 | 1,34E-12 | 4,62E-10 | CXCR4 |
| 12,4580708 | 3,94358515 | 0,89777472 | 1,12E-05 | 0,00038457 | KIAA0040 |
| 90,6657446 | 3,94096655 | 0,4522473 | 2,93E-18 | 6,16E-15 | RIPOR2 |
| 2,63315008 | 3,93738917 | 1,88365545 | 0,03659161 | 0,16444223 | LINC01732 |
| 2,29931439 | 3,93513608 | 1,82866072 | 0,03140346 | 0,14832374 | ARK2C |
| 57,9892363 | 3,93448966 | 0,79243663 | 6,87E-07 | 3,59E-05 | IKZF3 |
| 13,6399068 | 3,93342389 | 0,89044412 | 9,99E-06 | 0,00034896 | BEND2 |
| 136,202007 | 3,93063915 | 0,61888045 | 2,14E-10 | 3,67E-08 | PECAM1 |
| 38,0997009 | 3,92792316 | 1,04073329 | 0,00016053 | 0,00330869 | CD4 |
| 25,1287419 | 3,92653137 | 0,72300546 | 5,61E-08 | 4,15E-06 | LCK |
| 93,9684607 | 3,92610555 | 0,65922374 | 2,59E-09 | 2,97E-07 | CSF2RB |
| 2,92391509 | 3,9234936 | 1,6629429 | 0,01830611 | 0,10410879 | TRG-AS1 |
| 4,3744176 | 3,92078266 | 1,50411425 | 0,0091418 | 0,06534642 | TBXA2R |
| 3,04807443 | 3,91630024 | 1,51893408 | 0,00992816 | 0,06903373 | LINC01762 |
| 1,8234882 | 3,91064962 | 1,81925869 | 0,03158812 | 0,14893568 | ASAH2 |
| 1,7520969 | 3,90472609 | 1,93204633 | 0,04327653 | 0,18396656 | GOLGA8EP |
| 16,9857197 | 3,90160719 | 0,82906527 | 2,53E-06 | 0,0001078 | GMPR |
| 40,1999179 | 3,90070725 | 0,63411536 | 7,68E-10 | 1,02E-07 | TLR2 |
| 6,94238297 | 3,89205773 | 1,19750852 | 0,00115352 | 0,01493897 | PTGES |
| 16,8080044 | 3,89132777 | 0,8302017 | 2,77E-06 | 0,00011507 | ABCA5 |
| 15,0400616 | 3,89004658 | 0,92091652 | 2,40E-05 | 0,00071934 | LILRA2 |

| | | | | | |
|---|---|---|---|---|---|
| 58,4645522 | 3,88934655 | 0,56195928 | 4,48E-12 | 1,21E-09 | CD84 |
| 80,6933948 | 3,88478088 | 0,47938491 | 5,33E-16 | 5,61E-13 | DMTN |
| 56,1966535 | 3,88442633 | 0,5250294 | 1,38E-13 | 5,92E-11 | ADA2 |
| 2,74476602 | 3,87637859 | 1,597994 | 0,01527561 | 0,09251917 | ZNF418 |
| 157,29656 | 3,87176453 | 0,62178696 | 4,76E-10 | 6,77E-08 | RASSF2 |
| 7,46309995 | 3,86074939 | 1,12137733 | 0,00057559 | 0,0088327 | CCR4 |
| 4,78688619 | 3,85495786 | 1,41020716 | 0,0062644 | 0,05093627 | KIF5C |
| 3,05670548 | 3,84952288 | 1,54060691 | 0,01246479 | 0,08057609 | IGFBP3 |
| 4,31738776 | 3,83589747 | 1,46666246 | 0,00891251 | 0,06427333 | GAPT |
| 23,2865947 | 3,83387418 | 0,74710536 | 2,87E-07 | 1,77E-05 | EPSTI1 |
| 2,29768751 | 3,83035361 | 1,91340281 | 0,04530041 | 0,1898403 | CCIN |
| 1,83387993 | 3,82331689 | 1,89194333 | 0,04329621 | 0,18400891 | LINC00243 |
| 60,2876389 | 3,81623528 | 0,51550616 | 1,33E-13 | 5,86E-11 | RASGRP2 |
| 10,5183954 | 3,80986653 | 0,97638382 | 9,54E-05 | 0,00219314 | CD9 |
| 33,6397644 | 3,8091987 | 0,65897509 | 7,45E-09 | 7,38E-07 | ARHGAP25 |
| 4,5672997 | 3,80910962 | 1,46368211 | 0,00925695 | 0,06585498 | RNF217 |
| 6,11454083 | 3,80618705 | 1,33964931 | 0,00449467 | 0,04020977 | PACSIN1 |
| 4,77745209 | 3,80501658 | 1,38127991 | 0,00587453 | 0,04862295 | ZNF525 |
| 5,63021636 | 3,79766447 | 1,5956086 | 0,01730922 | 0,10009407 | LINC00861 |
| 4,81258778 | 3,79287895 | 1,48021042 | 0,0103954 | 0,0710847 | TTC24 |
| 2,57706093 | 3,77880785 | 1,62769454 | 0,02025607 | 0,11131718 | TPM2 |
| 2,7692078 | 3,77767286 | 1,67128361 | 0,02380002 | 0,12368052 | FUT3 |
| 81,963728 | 3,77697017 | 0,74721503 | 4,31E-07 | 2,43E-05 | CD37 |
| 12,6109299 | 3,77332771 | 0,84549504 | 8,09E-06 | 0,00029314 | INSR |
| 11,5423916 | 3,76976495 | 1,05500028 | 0,0003526 | 0,00612063 | CCR2 |
| 2,03016677 | 3,76950744 | 1,76435203 | 0,0326401 | 0,15211108 | IGKV3-20 |
| 6,81669068 | 3,76923549 | 1,12866382 | 0,00083913 | 0,01182211 | MIR4742 |
| 3,73100026 | 3,76555079 | 1,57596253 | 0,01687749 | 0,09868931 | PAOX |
| 79,5391881 | 3,75657001 | 0,44744326 | 4,63E-17 | 7,31E-14 | SESN3 |
| 13,2861319 | 3,75551486 | 0,99287853 | 0,00015529 | 0,0032156 | GASK1B |
| 2,43688951 | 3,75259762 | 1,88383834 | 0,04637154 | 0,19308891 | ZNF573 |
| 101,765496 | 3,74800779 | 0,70063184 | 8,82E-08 | 6,19E-06 | CXCL8 |
| 64,1957572 | 3,74537067 | 0,5201616 | 6,00E-13 | 2,32E-10 | SH3BGRL2 |
| 4,32286998 | 3,74107195 | 1,59357535 | 0,01889498 | 0,10611809 | RBPMS2 |
| 27,2456179 | 3,73667119 | 0,62913586 | 2,86E-09 | 3,24E-07 | TENM1 |
| 81,4174201 | 3,73559858 | 0,68457498 | 4,85E-08 | 3,70E-06 | PTGS1 |
| 12,1299414 | 3,72660481 | 0,95120237 | 8,94E-05 | 0,00207729 | PLEKHG1 |
| 16,23263 | 3,72475795 | 0,80022608 | 3,25E-06 | 0,00013321 | H1-0 |
| 13,700645 | 3,72407817 | 0,91537664 | 4,73E-05 | 0,00123734 | ZNF43 |
| 44,4460758 | 3,71294553 | 0,54348823 | 8,39E-12 | 2,09E-09 | CD247 |
| 3,97773838 | 3,70739876 | 1,76005341 | 0,03516852 | 0,15969483 | ADAM12 |
| 10,6123084 | 3,70506089 | 0,89618813 | 3,56E-05 | 0,00097655 | FCRL1 |
| 5,1321574 | 3,69243076 | 1,34289786 | 0,00596682 | 0,04921457 | ZNF440 |
| 145,540802 | 3,68874433 | 0,57864761 | 1,83E-10 | 3,18E-08 | INPP5D |
| 19,7749901 | 3,68222478 | 0,77097022 | 1,79E-06 | 8,13E-05 | RASGEF1B |
| 67,2899039 | 3,67629703 | 0,69294364 | 1,12E-07 | 7,65E-06 | MPEG1 |
| 20,3096393 | 3,67334456 | 0,72536394 | 4,10E-07 | 2,35E-05 | ICOSLG |
| 2,44319894 | 3,67262559 | 1,7365627 | 0,03443999 | 0,15759105 | IFNLR1 |

| | | | | | |
|---|---|---|---|---|---|
| 124,231287 | 3,6713333 | 0,64811549 | 1,47E-08 | 1,31E-06 | ARHGDIB |
| 5,01413325 | 3,670903 | 1,50802077 | 0,01492259 | 0,09119843 | TSIX |
| 42,2928455 | 3,66846356 | 0,5719812 | 1,42E-10 | 2,59E-08 | SCML1 |
| 56,6690527 | 3,66315332 | 0,53047543 | 5,01E-12 | 1,33E-09 | HCK |
| 64,5955909 | 3,66217077 | 0,52887829 | 4,38E-12 | 1,20E-09 | NFAM1 |
| 13,8331453 | 3,6574715 | 1,01851455 | 0,00032943 | 0,00580377 | SELENBP1 |
| 287,336218 | 3,65732177 | 0,5539782 | 4,06E-11 | 8,63E-09 | SPOCK2 |
| 8,93322811 | 3,65320396 | 1,0053346 | 0,00027926 | 0,00510021 | LRATD2 |
| 2,20602415 | 3,65036434 | 1,77087754 | 0,0392715 | 0,17276354 | LOC283194 |
| 46,0147226 | 3,64815388 | 0,55573569 | 5,22E-11 | 1,09E-08 | WDFY4 |
| 9,23555534 | 3,64189943 | 1,06766376 | 0,00064703 | 0,00963976 | PIK3CD-AS1 |
| 66,4747399 | 3,63974298 | 0,4372822 | 8,54E-17 | 1,24E-13 | SNCA |
| 9,17925798 | 3,63661098 | 1,05053564 | 0,00053683 | 0,00842932 | ZNF383 |
| 27,291533 | 3,63566907 | 0,73172494 | 6,74E-07 | 3,57E-05 | ZNF91 |
| 54,9058585 | 3,6214979 | 0,53774626 | 1,64E-11 | 3,89E-09 | KCNA3 |
| 138,445649 | 3,62061514 | 0,64671256 | 2,16E-08 | 1,84E-06 | LSP1 |
| 400,504886 | 3,61847582 | 0,57332447 | 2,77E-10 | 4,47E-08 | ALAS2 |
| 3,50453789 | 3,61803085 | 1,53540595 | 0,01845303 | 0,10462449 | MFSD2A |
| 14,4532737 | 3,61502679 | 1,05506457 | 0,00061174 | 0,00922287 | LOC728989 |
| 17,8479954 | 3,61291751 | 0,84501767 | 1,91E-05 | 0,00058947 | CLEC12A |
| 14,2707555 | 3,61214095 | 0,85015668 | 2,15E-05 | 0,00065802 | PTGER2 |
| 14,3610823 | 3,60394953 | 0,89644245 | 5,81E-05 | 0,00145484 | ZNF44 |
| 10,8835338 | 3,60323733 | 0,89796216 | 6,00E-05 | 0,00149272 | ARHGAP6 |
| 4,22074127 | 3,60304224 | 1,58476354 | 0,02299294 | 0,12091423 | TPM3P9 |
| 6,42093045 | 3,5988028 | 1,17989695 | 0,00228766 | 0,02472003 | FUT5 |
| 8,03164228 | 3,59177166 | 1,05774989 | 0,0006846 | 0,01009617 | ZNF280D |
| 2,65561896 | 3,58816185 | 1,6678886 | 0,03145096 | 0,14847397 | CES4A |
| 21,1230607 | 3,58237125 | 0,69774103 | 2,83E-07 | 1,75E-05 | VNN2 |
| 11,3186948 | 3,57742502 | 0,95214428 | 0,0001718 | 0,00350293 | ULK2 |
| 21,3618257 | 3,5740263 | 0,66250424 | 6,86E-08 | 4,98E-06 | JAML |
| 5,82810363 | 3,57148532 | 1,42911889 | 0,01245155 | 0,08051803 | KLRF1 |
| 11,6072679 | 3,56757554 | 0,85668368 | 3,12E-05 | 0,0008882 | CD22 |
| 88,2367721 | 3,56685126 | 0,45132729 | 2,72E-15 | 2,20E-12 | TMC8 |
| 3,27258144 | 3,5624928 | 1,5737453 | 0,02359235 | 0,122905 | ITGB8 |
| 31,1479913 | 3,55928564 | 0,61542697 | 7,32E-09 | 7,29E-07 | TESPA1 |
| 2,0699166 | 3,55423537 | 1,76018657 | 0,0434625 | 0,18454982 | AVPR1A |
| 14,1484872 | 3,55066995 | 0,98310833 | 0,00030422 | 0,00541506 | ZNF529 |
| 15,9843203 | 3,54860793 | 0,73991364 | 1,62E-06 | 7,62E-05 | ZNF736 |
| 11,0022972 | 3,54837201 | 0,92278808 | 0,00012042 | 0,00264934 | P2RX7 |
| 35,9455115 | 3,54737392 | 0,91946685 | 0,00011428 | 0,00254393 | LILRB1 |
| 123,350828 | 3,54681222 | 0,82503633 | 1,72E-05 | 0,00054382 | HBA2 |
| 89,1829241 | 3,52808795 | 0,44807015 | 3,44E-15 | 2,60E-12 | SASH3 |
| 3,41305609 | 3,52119972 | 1,54130057 | 0,02233841 | 0,11816188 | TRAJ23 |
| 12,2948058 | 3,51795057 | 0,83981268 | 2,80E-05 | 0,00081571 | BANK1 |
| 24,5858789 | 3,5174278 | 0,65242781 | 7,00E-08 | 5,05E-06 | LOC102723996 |
| 17,6402918 | 3,51499704 | 0,7244722 | 1,22E-06 | 5,94E-05 | AUTS2 |
| 5,66912697 | 3,51383518 | 1,18228524 | 0,00295799 | 0,02971657 | MSRB3 |
| 3,52575557 | 3,51348529 | 1,68432517 | 0,03697954 | 0,16564627 | CACNA1I |

| | | | | | |
|---|---|---|---|---|---|
| 2,8255656 | 3,49839547 | 1,61718257 | 0,03052075 | 0,14579227 | SPATA46 |
| 70,480477 | 3,49132128 | 0,44469478 | 4,13E-15 | 2,97E-12 | ARRB1 |
| 10,2357552 | 3,47223907 | 0,96347749 | 0,00031352 | 0,00554411 | CX3CR1 |
| 56,172082 | 3,46736276 | 0,49168657 | 1,76E-12 | 5,75E-10 | RAB31 |
| 8,65797916 | 3,46717631 | 0,9166641 | 0,00015533 | 0,0032156 | ZBTB16 |
| 5,03091865 | 3,46614698 | 1,2541251 | 0,00571331 | 0,04774803 | CNR1 |
| 14,927851 | 3,46580493 | 0,78058816 | 9,00E-06 | 0,00032218 | PLEKHA1 |
| 7,00619853 | 3,46328922 | 1,0249455 | 0,0007275 | 0,01058041 | FCER2 |
| 4,15671962 | 3,4595382 | 1,47676849 | 0,0191478 | 0,10712371 | C10orf105 |
| 58,4477524 | 3,45141489 | 0,51293856 | 1,71E-11 | 4,00E-09 | GNLY |
| 6,30504082 | 3,44651019 | 1,12834711 | 0,00225453 | 0,02447391 | CD38 |
| 6,96410732 | 3,44197845 | 1,09703399 | 0,00170377 | 0,02007313 | ANKRD63 |
| 2,55961116 | 3,43534648 | 1,65506803 | 0,03792608 | 0,16860887 | LINC00877 |
| 19,6048633 | 3,43003953 | 0,75145311 | 5,01E-06 | 0,0001929 | CEACAM21 |
| 2,99709895 | 3,42908355 | 1,54329543 | 0,02628851 | 0,13206016 | ZNF461 |
| 20,4479776 | 3,42776092 | 0,82547682 | 3,29E-05 | 0,00092473 | KLRD1 |
| 7,98683024 | 3,4274826 | 1,07597532 | 0,00144521 | 0,01779105 | ZNF483 |
| 66,8884048 | 3,42712822 | 0,48836648 | 2,26E-12 | 6,89E-10 | PDE3B |
| 12,7135441 | 3,42512507 | 0,99200956 | 0,00055498 | 0,00866538 | PCED1B |
| 28,7267001 | 3,42423137 | 0,58289559 | 4,24E-09 | 4,56E-07 | SLC40A1 |
| 2,92781889 | 3,42105356 | 1,72758591 | 0,04767534 | 0,19665846 | SLCO4C1 |
| 43,1481364 | 3,42004457 | 0,54000893 | 2,40E-10 | 3,98E-08 | PRKAG2 |
| 5,50627413 | 3,41172948 | 1,32671803 | 0,01012427 | 0,06988737 | LOC102724624 |
| 13,7510287 | 3,40858625 | 0,84897366 | 5,95E-05 | 0,00148615 | ABLIM3 |
| 63,3690869 | 3,40504257 | 0,48710957 | 2,74E-12 | 8,13E-10 | ALOX5 |
| 58,8387368 | 3,40268328 | 0,7839526 | 1,42E-05 | 0,00046473 | CCND2 |
| 4,80356148 | 3,39452489 | 1,14369898 | 0,00299722 | 0,03000549 | RTN1 |
| 4,83711043 | 3,39382653 | 1,50319544 | 0,02396179 | 0,12397621 | ZNF329 |
| 8,17441128 | 3,39286806 | 1,11674629 | 0,00238018 | 0,02540062 | SIGLEC1 |
| 50,2051284 | 3,39186555 | 0,45963851 | 1,59E-13 | 6,68E-11 | TLE4 |
| 2,99509926 | 3,38249778 | 1,52106121 | 0,02616359 | 0,13158499 | ZNF566 |
| 5,12867451 | 3,38008263 | 1,21966744 | 0,00558304 | 0,04694374 | CXorf65 |
| 103,006241 | 3,37508023 | 0,67963539 | 6,83E-07 | 3,59E-05 | FCN1 |
| 8,92431786 | 3,37077806 | 1,13223511 | 0,00290988 | 0,0294795 | LCN2 |
| 75,3731464 | 3,36530506 | 0,71629196 | 2,62E-06 | 0,00011108 | P2RY8 |
| 47,4215034 | 3,36183295 | 0,50772168 | 3,56E-11 | 7,65E-09 | SRC |
| 13,0492829 | 3,36166734 | 0,77154742 | 1,32E-05 | 0,0004407 | TGFBR3 |
| 63,2809588 | 3,34683597 | 0,47249825 | 1,41E-12 | 4,76E-10 | AHR |
| 10,6307109 | 3,34402463 | 0,93447695 | 0,00034557 | 0,00602636 | ADM |
| 4,84620405 | 3,34374772 | 1,50060978 | 0,02586323 | 0,13059998 | TXLNGY |
| 12,4970016 | 3,34325447 | 0,85391521 | 9,03E-05 | 0,00209388 | TFEC |
| 2,04810611 | 3,33237097 | 1,70021257 | 0,04999893 | 0,20227277 | TNFRSF11A |
| 34,8115246 | 3,33219235 | 0,55362986 | 1,76E-09 | 2,08E-07 | DAPK1 |
| 3,68574658 | 3,32443122 | 1,61073089 | 0,03902462 | 0,17203749 | P2RY6 |
| 148,594904 | 3,32259131 | 0,59518834 | 2,37E-08 | 2,00E-06 | SLC11A1 |
| 3,24664846 | 3,3219674 | 1,49137008 | 0,02591654 | 0,13065893 | ACSL6 |
| 19,8748072 | 3,31961709 | 1,32379856 | 0,01215386 | 0,0793524 | LBH |
| 70,8122624 | 3,31927587 | 0,48205163 | 5,75E-12 | 1,51E-09 | RAP1GAP2 |

| | | | | | |
|---|---|---|---|---|---|
| 13,4974791 | 3,31713347 | 0,83682046 | 7,37E-05 | 0,00177895 | OAS3 |
| 3,42881237 | 3,31574951 | 1,4513123 | 0,022333 | 0,11816188 | ZIK1 |
| 12,2987624 | 3,30013621 | 0,89822596 | 0,00023873 | 0,00450793 | APOBEC3A |
| 26,059203 | 3,29933935 | 0,69737414 | 2,23E-06 | 9,85E-05 | CD36 |
| 3,59797418 | 3,29666668 | 1,45500381 | 0,02346704 | 0,12256757 | TFPI |
| 3,35977286 | 3,296076 | 1,53879217 | 0,03219436 | 0,15082232 | ARHGAP42 |
| 6,36372129 | 3,29213599 | 1,10586492 | 0,00291105 | 0,0294795 | FCGR2B |
| 58,795756 | 3,28439428 | 0,49852051 | 4,45E-11 | 9,35E-09 | NCF1 |
| 2,51663405 | 3,27922656 | 1,57634885 | 0,03750105 | 0,16726952 | TMEM95 |
| 3,1531454 | 3,27460784 | 1,60197207 | 0,04094264 | 0,1771092 | ZNF761 |
| 5,0986858 | 3,26382241 | 1,26368512 | 0,00980074 | 0,06858965 | LY6G6F |
| 55,7792318 | 3,26374206 | 0,51079874 | 1,66E-10 | 2,97E-08 | ADAM28 |
| 16,3010783 | 3,24626157 | 0,7831107 | 3,39E-05 | 0,00094551 | RGS1 |
| 6,43470539 | 3,24547463 | 1,24829886 | 0,0093246 | 0,06617811 | MUC5AC |
| 3,85749796 | 3,24262378 | 1,43482712 | 0,02382495 | 0,12371385 | CA6 |
| 3,64471112 | 3,23774962 | 1,59577789 | 0,04246363 | 0,18155502 | CCL28 |
| 10,1155013 | 3,2337101 | 0,88387048 | 0,00025362 | 0,00473244 | AIF1 |
| 34,4382926 | 3,22108448 | 0,57503054 | 2,12E-08 | 1,82E-06 | PELI2 |
| 3,35827827 | 3,22071923 | 1,45297371 | 0,02664772 | 0,13331611 | EPHA2 |
| 5,94713547 | 3,21676924 | 1,33125301 | 0,01567711 | 0,09425696 | FAM30A |
| 5,71053139 | 3,21384554 | 1,48658418 | 0,03062593 | 0,14607339 | THEMIS |
| 57,2608858 | 3,21039571 | 0,51026897 | 3,14E-10 | 4,83E-08 | SELL |
| 10,5635002 | 3,20809961 | 0,93580671 | 0,00060768 | 0,00917176 | ATP10A |
| 5,33610971 | 3,20727159 | 1,16181934 | 0,00577024 | 0,04805403 | ZNF718 |
| 6,92062181 | 3,20541568 | 1,13494002 | 0,00473834 | 0,04191405 | BEX4 |
| 56,2059457 | 3,20536828 | 0,4948103 | 9,30E-11 | 1,78E-08 | C16orf54 |
| 8,07588056 | 3,20173825 | 1,03427547 | 0,00196393 | 0,02221127 | SH3RF3 |
| 2,86415324 | 3,19704965 | 1,54110411 | 0,03803114 | 0,16891719 | DLGAP3 |
| 9,69740186 | 3,19391934 | 0,89630094 | 0,00036602 | 0,00627874 | UBASH3A |
| 2,72431009 | 3,18505292 | 1,5904355 | 0,04521708 | 0,18958135 | PTGDS |
| 29,694369 | 3,18291987 | 0,63792958 | 6,06E-07 | 3,26E-05 | CAMK4 |
| 191,295189 | 3,18122655 | 0,53675894 | 3,09E-09 | 3,46E-07 | NRGN |
| 221,462272 | 3,1802239 | 0,64883231 | 9,51E-07 | 4,74E-05 | TSC22D3 |
| 32,4297327 | 3,17997033 | 0,55539428 | 1,03E-08 | 9,47E-07 | CHFR |
| 13,6700541 | 3,17849295 | 0,7616834 | 3,01E-05 | 0,00086056 | ZNF136 |
| 66,2140258 | 3,17831636 | 0,47502256 | 2,22E-11 | 5,12E-09 | ANPEP |
| 7,26577086 | 3,16277772 | 1,16988858 | 0,00686163 | 0,0540519 | ZNF737 |
| 24,9289152 | 3,16180289 | 0,65314846 | 1,29E-06 | 6,22E-05 | ALDH2 |
| 11,7527748 | 3,16037569 | 0,90362998 | 0,00046978 | 0,00763631 | B3GNT5 |
| 26,5421438 | 3,14702147 | 0,51788858 | 1,23E-09 | 1,54E-07 | CD14 |
| 4,87764694 | 3,14429189 | 1,21211689 | 0,00948527 | 0,06708052 | TRIQK |
| 4,74062471 | 3,13672216 | 1,38617442 | 0,02364416 | 0,12307324 | ITM2A |
| 5,12423528 | 3,13335274 | 1,427906 | 0,02820889 | 0,13849651 | HOXB2 |
| 4,17525979 | 3,12228565 | 1,52583016 | 0,04072819 | 0,17634282 | MIR5684 |
| 3,29990939 | 3,11966843 | 1,53542475 | 0,04217441 | 0,18062063 | AOC4P |
| 4,4173674 | 3,11729235 | 1,28576855 | 0,01533124 | 0,09279667 | OR2A1 |
| 28,9506712 | 3,11422481 | 0,63102035 | 8,01E-07 | 4,10E-05 | DHRS3 |
| 251,185517 | 3,10342435 | 0,51127801 | 1,28E-09 | 1,59E-07 | CTSS |

| | | | | | |
|---|---|---|---|---|---|
| 5,63711591 | 3,10170798 | 1,14943111 | 0,00696585 | 0,05463009 | LRRC32 |
| 47,9511377 | 3,09908605 | 0,57061488 | 5,60E-08 | 4,15E-06 | SERPINB9 |
| 22,4050526 | 3,09353233 | 0,67860418 | 5,15E-06 | 0,00019757 | MEFV |
| 19,0588529 | 3,09140245 | 0,71738642 | 1,64E-05 | 0,00052442 | CD8A |
| 9,61274414 | 3,0896671 | 0,80166413 | 0,00011617 | 0,00257387 | CCR6 |
| 482,784499 | 3,08670839 | 0,5375082 | 9,32E-09 | 8,78E-07 | LAPTM5 |
| 17,7486621 | 3,08129406 | 0,83338911 | 0,0002179 | 0,00420274 | HLA-DMB |
| 7,91375181 | 3,06214268 | 1,06985743 | 0,00420715 | 0,03845582 | SLC14A1 |
| 5,71607337 | 3,04925352 | 1,08779712 | 0,00506068 | 0,04382298 | ALS2CL |
| 77,7713999 | 3,04369527 | 0,44556113 | 8,42E-12 | 2,09E-09 | BACH2 |
| 35,8852218 | 3,04345485 | 0,53387237 | 1,19E-08 | 1,07E-06 | CSF1 |
| 22,1235861 | 3,03851576 | 0,71966579 | 2,42E-05 | 0,00072341 | SECTM1 |
| 11,3200791 | 3,03634906 | 0,8406907 | 0,00030416 | 0,00541506 | NID1 |
| 33,0923618 | 3,02195414 | 0,59978301 | 4,69E-07 | 2,62E-05 | HCP5 |
| 4,95203973 | 3,01669099 | 1,09834543 | 0,00602206 | 0,04948969 | TLE1 |
| 4,27978751 | 3,01637212 | 1,498497 | 0,04412181 | 0,18633493 | SLC24A3 |
| 69,3616978 | 3,01504132 | 0,78610482 | 0,00012535 | 0,0027199 | HBA1 |
| 5,50381257 | 3,01266663 | 1,15269324 | 0,00895967 | 0,06448303 | GBGT1 |
| 4,98525967 | 3,01180892 | 1,26639456 | 0,0173948 | 0,10037398 | PTGS2 |
| 13,9686605 | 3,00870614 | 0,90233363 | 0,00085496 | 0,01200046 | HEMGN |
| 48,6747043 | 2,99841264 | 0,87961091 | 0,00065251 | 0,00971367 | HLA-DPB1 |
| 10,1948984 | 2,99836066 | 1,04862992 | 0,0042456 | 0,03868556 | GIMAP2 |
| 20,4593984 | 2,99799378 | 0,58944078 | 3,65E-07 | 2,15E-05 | SKAP1 |
| 55,3085866 | 2,99756715 | 0,47213372 | 2,17E-10 | 3,69E-08 | HRH2 |
| 111,436925 | 2,99567423 | 0,39533579 | 3,52E-14 | 1,84E-11 | CCL5 |
| 18,4196901 | 2,99186026 | 0,64286559 | 3,26E-06 | 0,00013337 | CLEC2B |
| 39,2863084 | 2,98994593 | 0,60900718 | 9,13E-07 | 4,61E-05 | GBP4 |
| 11,6105981 | 2,9842491 | 0,7246802 | 3,82E-05 | 0,00103292 | CRYAB |
| 5,55075645 | 2,97950108 | 1,11176701 | 0,0073629 | 0,05663147 | ITGA9-AS1 |
| 33,0880497 | 2,97080051 | 0,67773043 | 1,17E-05 | 0,00039895 | SH3TC1 |
| 7,29177096 | 2,96908883 | 0,96093411 | 0,00200295 | 0,02250042 | MIR29B2 |
| 7,65525332 | 2,96588293 | 0,94564791 | 0,00171065 | 0,02012884 | HCAR3 |
| 5,23261295 | 2,96581013 | 1,20407836 | 0,01377272 | 0,08626073 | CCDC144NL |
| 10,5665257 | 2,96105022 | 0,88923289 | 0,00086879 | 0,01216762 | CD2 |
| 3,42852353 | 2,96015153 | 1,37072655 | 0,03080758 | 0,14664407 | SPTA1 |
| 11,5135571 | 2,95893819 | 0,97416255 | 0,00238615 | 0,02542691 | LINGO3 |
| 3,22350801 | 2,95836901 | 1,49198665 | 0,04738544 | 0,19580255 | ZNF234 |
| 10,0874862 | 2,95819468 | 0,88448014 | 0,00082414 | 0,01168931 | BLK |
| 22,5198497 | 2,95247434 | 0,65649085 | 6,88E-06 | 0,00025526 | TNFSF8 |
| 10,6503326 | 2,95081743 | 1,03880066 | 0,00450287 | 0,04024473 | ZNF433 |
| 27,5351948 | 2,94181772 | 0,553286 | 1,05E-07 | 7,34E-06 | CD79A |
| 22,5899933 | 2,94079623 | 0,64102735 | 4,48E-06 | 0,00017526 | ARHGAP18 |
| 10,6880822 | 2,9398121 | 0,87888746 | 0,00082305 | 0,01168262 | C11orf21 |
| 43,3635215 | 2,93974135 | 0,44184986 | 2,87E-11 | 6,38E-09 | NFATC2 |
| 7,6433086 | 2,93348594 | 0,96000963 | 0,00224548 | 0,02441765 | CDC42BPG |
| 10,6219507 | 2,93155326 | 0,91389118 | 0,00133768 | 0,01675068 | ERP27 |
| 14,7037607 | 2,92933814 | 0,84406887 | 0,0005195 | 0,00821856 | SLC16A5 |
| 24,7253045 | 2,92416496 | 0,62848569 | 3,28E-06 | 0,00013387 | TLR8 |

| | | | | | |
|---|---|---|---|---|---|
| 6,41015859 | 2,89462917 | 1,14332077 | 0,01134873 | 0,0755823 | PID1 |
| 45,373572 | 2,88603214 | 0,46850816 | 7,27E-10 | 9,76E-08 | NCF1B |
| 36,8343988 | 2,87323569 | 0,56616414 | 3,88E-07 | 2,26E-05 | APBA2 |
| 3,00160862 | 2,86914775 | 1,39900844 | 0,04028216 | 0,17512995 | PDE9A |
| 6,31125921 | 2,85800391 | 1,02137062 | 0,00513876 | 0,04427131 | RLBP1 |
| 7,48786978 | 2,85417961 | 0,93245069 | 0,0022064 | 0,02408963 | LINC00304 |
| 27,1645923 | 2,85134213 | 0,54038809 | 1,32E-07 | 8,80E-06 | SWAP70 |
| 7,13480941 | 2,83367112 | 0,91069279 | 0,00186105 | 0,02140598 | POU2AF1 |
| 49,8740965 | 2,82685019 | 0,42512064 | 2,94E-11 | 6,47E-09 | NCF1C |
| 54,0445514 | 2,82536242 | 0,54433311 | 2,10E-07 | 1,35E-05 | GNG2 |
| 15,1751573 | 2,82472718 | 0,82288202 | 0,00059755 | 0,00908861 | ACSS1 |
| 5,39714299 | 2,81204335 | 1,21256055 | 0,02038988 | 0,11179276 | ZNF468 |
| 40,9503168 | 2,80995638 | 0,60514929 | 3,43E-06 | 0,00013915 | CARD11 |
| 36,8966806 | 2,80249827 | 1,0897824 | 0,01012261 | 0,06988737 | ANXA6 |
| 13,4099431 | 2,80200031 | 0,79648323 | 0,00043487 | 0,00719245 | P2RY10 |
| 8,56070897 | 2,79913072 | 0,9476748 | 0,00314007 | 0,0309928 | CCDC141 |
| 15,2561074 | 2,79685597 | 0,69732113 | 6,05E-05 | 0,00150025 | TSPAN32 |
| 46,2066515 | 2,77305619 | 0,47850889 | 6,82E-09 | 6,83E-07 | SPN |
| 80,4545605 | 2,76541868 | 0,43592739 | 2,24E-10 | 3,75E-08 | GAS7 |
| 6,9060265 | 2,76212248 | 1,09449806 | 0,01161458 | 0,07683896 | TNFRSF4 |
| 62,922624 | 2,75295376 | 0,47204653 | 5,48E-09 | 5,60E-07 | C5AR1 |
| 10,5016763 | 2,7503701 | 0,82297568 | 0,00083182 | 0,01177183 | PEX11G |
| 7,50138429 | 2,74394716 | 1,00689395 | 0,00642703 | 0,05170547 | PHACTR2-AS1 |
| 35,5857888 | 2,73466335 | 0,60094091 | 5,35E-06 | 0,00020362 | FLI1 |
| 21,1705489 | 2,73334519 | 0,6300333 | 1,44E-05 | 0,00046818 | FUT4 |
| 10,9432463 | 2,73060322 | 0,77045852 | 0,00039391 | 0,0066666 | MTCL2 |
| 13,6297072 | 2,72844854 | 0,70263982 | 0,00010312 | 0,00234221 | SLC2A6 |
| 6,40690556 | 2,72613311 | 1,07582957 | 0,01127744 | 0,07529304 | MMRN1 |
| 8,37939261 | 2,72467725 | 1,11843531 | 0,01484449 | 0,09092669 | GIMAP6 |
| 3,88763612 | 2,71698826 | 1,21189624 | 0,02496581 | 0,12746303 | RNF182 |
| 5,78702107 | 2,71200344 | 1,06521894 | 0,01089781 | 0,07337986 | MIR6770-1 |
| 6,83908278 | 2,70815521 | 0,99993815 | 0,00676242 | 0,05342455 | ZC3H12C |
| 20,9121683 | 2,70383757 | 0,57684256 | 2,77E-06 | 0,00011507 | UNC119 |
| 7,85331438 | 2,69966153 | 1,08459967 | 0,01280721 | 0,08189429 | GRAPL |
| 11,9367868 | 2,69899893 | 0,84566809 | 0,00141508 | 0,01749977 | IKZF2 |
| 24,8065752 | 2,69822089 | 0,50863387 | 1,13E-07 | 7,65E-06 | SNX29P2 |
| 9,53004627 | 2,69710976 | 0,82610744 | 0,00109525 | 0,01440118 | CYP4F3 |
| 5,57681457 | 2,69691915 | 1,08367996 | 0,01282228 | 0,08193528 | GZMK |
| 4,66990283 | 2,69504426 | 1,34258196 | 0,04471143 | 0,18816391 | GPR4 |
| 12,4017036 | 2,69481298 | 0,83436884 | 0,00123893 | 0,0158177 | B3GNT7 |
| 24,6828948 | 2,68978006 | 0,5664287 | 2,05E-06 | 9,21E-05 | CASP10 |
| 6,3429975 | 2,66052617 | 1,17788631 | 0,02390024 | 0,12397567 | VNN3P |
| 3,97738239 | 2,65925874 | 1,27805351 | 0,03746046 | 0,16716729 | PLAU |
| 8,04485693 | 2,65683687 | 0,89896796 | 0,00312234 | 0,03086766 | KLRC3 |
| 65,7020453 | 2,65662235 | 0,35709318 | 1,01E-13 | 4,66E-11 | HSPA7 |
| 6,19944349 | 2,65376087 | 1,14808936 | 0,02080754 | 0,11324699 | ST6GALNAC1 |
| 9,729829 | 2,64236246 | 0,95168557 | 0,00549463 | 0,04643319 | INPP4B |
| 38,0839736 | 2,63910733 | 0,55742493 | 2,20E-06 | 9,76E-05 | PADI2 |

| | | | | | |
|---|---|---|---|---|---|
| 21,6266437 | 2,63224998 | 0,72467888 | 0,00028091 | 0,00512079 | SLC24A4 |
| 11,7553777 | 2,62842428 | 0,68328613 | 0,0001197 | 0,00263967 | BST1 |
| 69,5088187 | 2,62821861 | 0,4463613 | 3,91E-09 | 4,22E-07 | ARHGAP9 |
| 9,64722954 | 2,6132732 | 0,91215404 | 0,00417079 | 0,03825137 | HSD11B2 |
| 67,7382234 | 2,60478005 | 0,40391776 | 1,13E-10 | 2,13E-08 | STAB1 |
| 6,52791047 | 2,59771289 | 1,00211721 | 0,00953575 | 0,0673231 | PCDHGA1 |
| 6,37194296 | 2,5965455 | 0,94942309 | 0,00624056 | 0,05078605 | MTAP |
| 19,7218142 | 2,59517884 | 0,55464662 | 2,88E-06 | 0,00011911 | ST8SIA4 |
| 30,2647303 | 2,59514001 | 0,45453243 | 1,13E-08 | 1,03E-06 | SLC8A1 |
| 12,8570588 | 2,59013827 | 0,77245303 | 0,00079902 | 0,01141864 | VIPR1 |
| 7,90819946 | 2,58905999 | 0,96114234 | 0,00706569 | 0,05510716 | ZAR1L |
| 95,1453972 | 2,58529265 | 0,40388376 | 1,54E-10 | 2,78E-08 | PRKCB |
| 15,6482443 | 2,57575753 | 0,65987015 | 9,48E-05 | 0,00218834 | TRBC1 |
| 3,88743993 | 2,57566786 | 1,2569105 | 0,04044203 | 0,17562626 | MESTIT1 |
| 5,57268685 | 2,57276764 | 1,02441775 | 0,01202384 | 0,07874805 | TSBP1 |
| 7,83997506 | 2,56257345 | 0,85433326 | 0,00270422 | 0,02807577 | PABPN1L |
| 3,91300071 | 2,5618327 | 1,26902478 | 0,04351412 | 0,18464468 | MIR6746 |
| 5,64507464 | 2,56143159 | 1,2472818 | 0,04001336 | 0,17456601 | RHBDL3 |
| 81,5990975 | 2,55816309 | 0,64838672 | 7,97E-05 | 0,00190098 | CR1 |
| 3,29209654 | 2,53419942 | 1,24486829 | 0,04177879 | 0,17937293 | FCGR1BP |
| 7,85275923 | 2,52917791 | 1,06667147 | 0,01773554 | 0,10174141 | LOC101927131 |
| 38,6930267 | 2,52849361 | 0,49482273 | 3,22E-07 | 1,96E-05 | ANK1 |
| 5,69775058 | 2,52798958 | 1,09816917 | 0,02133496 | 0,11509571 | CDC42EP5 |
| 7,23213732 | 2,51026751 | 0,92147865 | 0,00644627 | 0,05171486 | PTGIR |
| 42,0245785 | 2,5088594 | 0,5651144 | 9,01E-06 | 0,00032218 | APBB1IP |
| 3,81873385 | 2,50477648 | 1,24470394 | 0,04418335 | 0,18652235 | ADGRE4P |
| 23,6262122 | 2,49928736 | 0,61220163 | 4,46E-05 | 0,00117431 | SPTB |
| 22,8116747 | 2,4957982 | 0,65310468 | 0,00013268 | 0,00283338 | ANO9 |
| 5,69838179 | 2,48887549 | 0,9597401 | 0,00950651 | 0,06714175 | PDLIM4 |
| 11,5105521 | 2,48321433 | 0,70264402 | 0,00040917 | 0,00688759 | LINC01619 |
| 7,20299072 | 2,48220604 | 1,06306567 | 0,01954599 | 0,10869791 | GIMAP8 |
| 14,2305756 | 2,48010473 | 0,75841183 | 0,00107498 | 0,01425279 | MMP9 |
| 6,32675356 | 2,47522012 | 0,97787747 | 0,01136675 | 0,07562245 | CTSF |
| 19,5950953 | 2,46651904 | 0,67041359 | 0,00023406 | 0,00445388 | FCGR2C |
| 7,03566375 | 2,46571279 | 1,04433608 | 0,01822407 | 0,10386056 | THRB-AS2 |
| 4,56108115 | 2,45950722 | 1,05201832 | 0,01939275 | 0,10801598 | TAX1BP3 |
| 3,47026774 | 2,45751625 | 1,21786095 | 0,04360214 | 0,18493522 | SNORD116-16 |
| 8,26002315 | 2,45390858 | 0,84976826 | 0,00388022 | 0,03616635 | MS4A4E |
| 13,8844468 | 2,44672869 | 0,85305786 | 0,00412833 | 0,03793689 | EHD3 |
| 14,3513317 | 2,44524965 | 0,72962625 | 0,00080412 | 0,01148282 | GPD2 |
| 13,4218605 | 2,44280846 | 0,64603668 | 0,00015606 | 0,00322712 | CORO2A |
| 4,21854329 | 2,44260262 | 1,05147306 | 0,02017758 | 0,11106148 | LPAR5 |
| 11,2844883 | 2,44247625 | 0,80341277 | 0,00236479 | 0,02526493 | TNF |
| 7,58464674 | 2,43358033 | 0,97435183 | 0,0125023 | 0,08073584 | CABP4 |
| 60,6009109 | 2,43344877 | 0,94188175 | 0,00977742 | 0,06854336 | SLA |
| 5,06725707 | 2,4332736 | 1,07680776 | 0,02383924 | 0,1237482 | DYNLT3 |
| 13,8641946 | 2,43179069 | 0,77882281 | 0,00179387 | 0,02089323 | GCNT2 |
| 22,3464179 | 2,42766294 | 0,55665802 | 1,29E-05 | 0,00043417 | SCRN1 |

| | | | | | |
|---|---|---|---|---|---|
| 13,5852513 | 2,42748632 | 0,73406639 | 0,00094334 | 0,01291524 | GLT1D1 |
| 17,1397724 | 2,41494995 | 0,72175954 | 0,0008201 | 0,01165823 | CD3G |
| 16,335391 | 2,4099487 | 0,56773091 | 2,19E-05 | 0,00066425 | LST1 |
| 66,2526256 | 2,40666982 | 0,39345148 | 9,55E-10 | 1,22E-07 | PTPN6 |
| 4,19995457 | 2,4015804 | 1,19736913 | 0,04488674 | 0,18869184 | IL15 |
| 25,5599451 | 2,39866293 | 0,56801183 | 2,41E-05 | 0,00072202 | P2RY13 |
| 661,048959 | 2,38896147 | 0,6237709 | 0,00012821 | 0,00276296 | CD74 |
| 12,6002045 | 2,38494913 | 0,87820909 | 0,00661366 | 0,05264734 | TNFRSF10C |
| 4,80195178 | 2,38393993 | 1,09357009 | 0,02926037 | 0,14163095 | ORM1 |
| 8,56837483 | 2,38281502 | 0,82529347 | 0,00388646 | 0,03620661 | FOXN3-AS2 |
| 124,848535 | 2,37076326 | 0,55298251 | 1,81E-05 | 0,00056768 | FYB1 |
| 25,630128 | 2,36987048 | 0,49511531 | 1,70E-06 | 7,87E-05 | PLBD1 |
| 8,23402843 | 2,36513319 | 0,88639509 | 0,00762451 | 0,05798364 | SERPINF2 |
| 28,4770313 | 2,36331583 | 0,61944834 | 0,00013608 | 0,00288319 | IL2RG |
| 11,9890266 | 2,36244281 | 0,66701386 | 0,00039738 | 0,00671319 | EMID1 |
| 52,1519531 | 2,36217227 | 0,42035936 | 1,92E-08 | 1,66E-06 | FPR1 |
| 9,82900188 | 2,35973123 | 0,88609751 | 0,00774335 | 0,05849564 | CBFA2T3 |
| 4,45954234 | 2,35951255 | 1,1584819 | 0,0416773 | 0,17914045 | LIMS2 |
| 5,08497711 | 2,35828226 | 1,13993597 | 0,03856628 | 0,17041397 | ARHGAP32 |
| 11,1107402 | 2,35639466 | 0,87682288 | 0,00720055 | 0,05585964 | CHN2 |
| 109,344509 | 2,35050051 | 0,57252178 | 4,03E-05 | 0,0010842 | PLXDC2 |
| 22,8400889 | 2,33058503 | 0,584328 | 6,65E-05 | 0,0016265 | CPNE3 |
| 12,2357555 | 2,32953054 | 0,99667705 | 0,01942373 | 0,10811927 | HLA-DOA |
| 14,2925336 | 2,32767586 | 0,73523056 | 0,00154597 | 0,01869094 | F2R |
| 36,198574 | 2,32640564 | 0,5006838 | 3,38E-06 | 0,00013741 | PEAK3 |
| 52,7475724 | 2,32608872 | 0,39949302 | 5,79E-09 | 5,89E-07 | GBP5 |
| 8,77059246 | 2,3232591 | 0,79035112 | 0,00328713 | 0,0320762 | NPC1L1 |
| 12,9848502 | 2,32209511 | 0,72158528 | 0,00129067 | 0,01624799 | SNPH |
| 14,426198 | 2,3182386 | 0,71897629 | 0,00126254 | 0,01602531 | GNA15 |
| 186,725402 | 2,3170829 | 0,44293384 | 1,68E-07 | 1,10E-05 | ATP2A3 |
| 8,82428497 | 2,30977806 | 0,88825144 | 0,00931246 | 0,06611673 | ACAN |
| 157,159288 | 2,30885306 | 0,89184057 | 0,00962934 | 0,06783199 | HLA-DRA |
| 6,17544558 | 2,30806848 | 0,89208918 | 0,0096742 | 0,06807197 | MIR6801 |
| 6,35288364 | 2,2894565 | 1,08954025 | 0,03561419 | 0,16120959 | MIR4761 |
| 141,412239 | 2,28335532 | 0,53325191 | 1,85E-05 | 0,0005775 | ITGAL |
| 28,9550023 | 2,2634907 | 0,58607415 | 0,00011241 | 0,00251849 | TRAC |
| 5,14477643 | 2,2594507 | 1,10060805 | 0,04008118 | 0,17470962 | MIR4730 |
| 10,3913294 | 2,25940766 | 0,72967293 | 0,00195842 | 0,02216225 | TNR |
| 28,6644894 | 2,25811664 | 0,56991093 | 7,43E-05 | 0,00178748 | ADGRE1 |
| 8,34288132 | 2,25730897 | 0,80958698 | 0,00529981 | 0,04517843 | SARDH |
| 4,82122684 | 2,24764565 | 1,05706948 | 0,03347838 | 0,15457405 | SNX29P1 |
| 37,0348083 | 2,24443935 | 0,45905441 | 1,01E-06 | 4,99E-05 | XYLT1 |
| 6,93141229 | 2,2302295 | 1,13341515 | 0,0491017 | 0,20027017 | SCN11A |
| 10,0638276 | 2,22421654 | 0,79108048 | 0,00492929 | 0,04296045 | DNAJC15 |
| 37,5337044 | 2,22215927 | 0,49240777 | 6,40E-06 | 0,00023918 | VAV3 |
| 80,5982639 | 2,21856085 | 0,4024886 | 3,55E-08 | 2,83E-06 | IL16 |
| 5,27478032 | 2,21852963 | 1,04003752 | 0,03291451 | 0,15282831 | CD19 |
| 5,07086832 | 2,21743384 | 1,00611401 | 0,02752724 | 0,13614715 | LOC105373289 |

| | | | | | |
|---|---|---|---|---|---|
| 7,61792873 | 2,21211423 | 0,93741794 | 0,01828503 | 0,10402017 | SH2D1B |
| 13,9932305 | 2,20985583 | 0,7139121 | 0,00196536 | 0,02221423 | SLAMF7 |
| 128,56594 | 2,20730593 | 0,57586597 | 0,00012658 | 0,00273716 | FYN |
| 41,7940883 | 2,19868712 | 0,39484784 | 2,57E-08 | 2,14E-06 | FGD2 |
| 6,8349181 | 2,19761548 | 1,07477113 | 0,0408816 | 0,17688562 | UPB1 |
| 25,729537 | 2,19415543 | 0,58472688 | 0,00017512 | 0,00354755 | CXCL16 |
| 83,4836536 | 2,18819408 | 0,33567802 | 7,09E-11 | 1,41E-08 | ITGA2B |
| 8,19493777 | 2,18179501 | 0,82452167 | 0,00814175 | 0,06048764 | KLRK1-AS1 |
| 43,7080139 | 2,17991721 | 0,51319812 | 2,16E-05 | 0,00065908 | MGAM |
| 8,23314008 | 2,17957493 | 0,79799275 | 0,00630809 | 0,05111576 | IFIT1B |
| 20,7464564 | 2,17759373 | 0,56476601 | 0,00011538 | 0,00256529 | TREML1 |
| 11,3248427 | 2,17182027 | 0,66378405 | 0,00106836 | 0,01418554 | GFOD3P |
| 112,484791 | 2,16885576 | 0,3532628 | 8,28E-10 | 1,09E-07 | THBS1 |
| 18,1932928 | 2,16590857 | 0,68330898 | 0,00152578 | 0,01851781 | FAM157C |
| 13,2269772 | 2,16484802 | 0,78492061 | 0,00581478 | 0,04827621 | EPHA4 |
| 40,3216363 | 2,15588851 | 0,44264015 | 1,11E-06 | 5,46E-05 | PRAM1 |
| 143,361453 | 2,15109536 | 0,48012423 | 7,45E-06 | 0,00027334 | SPI1 |
| 8,19307626 | 2,13917722 | 0,78668156 | 0,00654318 | 0,05230397 | SULT1A1 |
| 7,5519772 | 2,13180326 | 0,96675062 | 0,02744552 | 0,13612452 | SGCG |
| 39,8922548 | 2,12507969 | 0,45798638 | 3,48E-06 | 0,00014113 | DPEP2 |
| 76,0930886 | 2,12256751 | 0,3550609 | 2,26E-09 | 2,64E-07 | BMF |
| 27,0068537 | 2,11966534 | 0,54081549 | 8,88E-05 | 0,00206605 | PSTPIP2 |
| 25,6023151 | 2,11358116 | 0,50921952 | 3,32E-05 | 0,00093079 | PARP15 |
| 7,69962024 | 2,08938984 | 1,05999829 | 0,0487095 | 0,1992719 | BNC2 |
| 34,3500109 | 2,08363327 | 0,47114582 | 9,76E-06 | 0,00034252 | TMCC3 |
| 361,075947 | 2,08353333 | 0,51318927 | 4,91E-05 | 0,00127558 | HSPA6 |
| 10,9876622 | 2,07913495 | 0,77830353 | 0,00755428 | 0,05758848 | FRAT1 |
| 7,77987639 | 2,07824529 | 0,82462636 | 0,01172793 | 0,07726469 | LPA |
| 387,829034 | 2,07690426 | 0,40866026 | 3,73E-07 | 2,19E-05 | TUBB1 |
| 4,03190718 | 2,06889003 | 1,0501395 | 0,0488258 | 0,19961818 | LOC101929578 |
| 41,1469492 | 2,06612235 | 0,41614291 | 6,87E-07 | 3,59E-05 | TBC1D10C |
| 15,1643033 | 2,05677835 | 0,68422895 | 0,00264727 | 0,0275972 | PDLIM1 |
| 13,6569059 | 2,05162068 | 0,63689238 | 0,00127611 | 0,01614335 | NLRP6 |
| 26,1863687 | 2,05139302 | 0,61773716 | 0,00089754 | 0,0124213 | TRBC2 |
| 13,8928988 | 2,04542021 | 0,68624059 | 0,00287669 | 0,02932755 | LYL1 |
| 10,7577842 | 2,04165343 | 0,7465055 | 0,00623902 | 0,05078605 | SH3TC2 |
| 13,216188 | 2,03976035 | 0,80899248 | 0,01169026 | 0,07716087 | BCL11A |
| 37,5951034 | 2,03536576 | 0,50723534 | 6,00E-05 | 0,00149272 | CELF2-AS1 |
| 6,02501022 | 2,02209773 | 0,92422246 | 0,02867757 | 0,1399918 | LINC03048 |
| 11,2378261 | 2,01238625 | 0,92479951 | 0,02955344 | 0,1425657 | TAL1 |
| 13,6777341 | 2,01154825 | 0,8212348 | 0,01430868 | 0,08841893 | NRROS |
| 9,70034234 | 2,00710678 | 0,73541879 | 0,00634877 | 0,05131358 | RAB4A |
| 93,6155973 | 1,99753008 | 0,38660462 | 2,38E-07 | 1,51E-05 | PIK3CG |
| 6,23064961 | 1,99686659 | 1,00073387 | 0,04599903 | 0,19204493 | NCAM1 |
| 51,5482529 | 1,993098 | 0,39716757 | 5,21E-07 | 2,85E-05 | LOC400499 |
| 15,1876911 | 1,98621833 | 0,6522708 | 0,00232617 | 0,02502175 | SDK2 |
| 23,1837946 | 1,9775271 | 0,59461083 | 0,00088181 | 0,01229529 | DTX4 |
| 398,993471 | 1,97223781 | 0,23881741 | 1,48E-16 | 1,92E-13 | FOS |

| | | | | | |
|---|---|---|---|---|---|
| 6,0763055 | 1,9715206 | 0,94940323 | 0,03783948 | 0,16834253 | NTNG2 |
| 7,48784119 | 1,97058634 | 0,93855569 | 0,03576453 | 0,161774 | CDH15 |
| 9,20736852 | 1,9626236 | 0,7607272 | 0,00988201 | 0,06891909 | SLAMF1 |
| 48,1230371 | 1,96189376 | 0,38616953 | 3,77E-07 | 2,21E-05 | S100A4 |
| 11,0181182 | 1,96084755 | 0,72894424 | 0,00714557 | 0,05556803 | CTDSPL |
| 28,9916994 | 1,94655616 | 0,5013535 | 0,00010334 | 0,00234458 | ADORA2A-AS1 |
| 33,2504171 | 1,94179972 | 0,50571941 | 0,0001232 | 0,00268252 | SCML4 |
| 8,62066918 | 1,93664436 | 0,70861442 | 0,006276 | 0,05098679 | MIR4505 |
| 12,9588762 | 1,92385361 | 0,78912975 | 0,01477109 | 0,09056506 | P2RY12 |
| 111,346762 | 1,91974068 | 0,37652666 | 3,42E-07 | 2,06E-05 | ITGAM |
| 72,2088005 | 1,91710795 | 0,42770458 | 7,38E-06 | 0,00027233 | HLA-F |
| 10,4212768 | 1,91268011 | 0,7737494 | 0,01343733 | 0,08466458 | AFAP1 |
| 8,83072797 | 1,89961768 | 0,95139501 | 0,04586152 | 0,19163998 | ADGRG5 |
| 7,09869445 | 1,89725064 | 0,85404106 | 0,02631724 | 0,13208183 | MIR4655 |
| 14,5228752 | 1,8969994 | 0,72871905 | 0,00923588 | 0,06580486 | MIR23A |
| 23,8045658 | 1,89475772 | 0,53411833 | 0,00038898 | 0,00659955 | ACRBP |
| 20,2292257 | 1,88963419 | 0,56580289 | 0,00083857 | 0,01182211 | PTP4A3 |
| 74,8700551 | 1,88805742 | 0,29934849 | 2,84E-10 | 4,56E-08 | S1PR1 |
| 29,3654422 | 1,88717567 | 0,50275821 | 0,00017428 | 0,00353439 | PTK2 |
| 10,9449121 | 1,88552022 | 0,78508274 | 0,01631982 | 0,09661683 | HCAR2 |
| 19,269286 | 1,88533687 | 0,61172853 | 0,00205627 | 0,0228662 | LTB |
| 22,6475925 | 1,87730387 | 0,67980355 | 0,00575297 | 0,04793127 | OAS2 |
| 63,4974637 | 1,86513753 | 0,35213913 | 1,18E-07 | 7,96E-06 | KLHL6 |
| 10,3968538 | 1,86333612 | 0,7552706 | 0,01362083 | 0,0855359 | GZMM |
| 42,3857449 | 1,8613913 | 0,36949901 | 4,71E-07 | 2,62E-05 | WAKMAR2 |
| 8,93541723 | 1,86121518 | 0,7665234 | 0,01517708 | 0,09224721 | TLR7 |
| 7,13598517 | 1,85789889 | 0,88075646 | 0,03490701 | 0,15888275 | DEPDC1-AS1 |
| 7,49865238 | 1,83945048 | 0,77876396 | 0,01817601 | 0,10374308 | LOC105374382 |
| 14,7108546 | 1,83365104 | 0,69591814 | 0,00841705 | 0,06184816 | CFD |
| 13,3377951 | 1,82773492 | 0,72951426 | 0,01223083 | 0,07966248 | FAS-AS1 |
| 10,432918 | 1,82512065 | 0,69030046 | 0,00819445 | 0,06077898 | VSIG2 |
| 7,27126867 | 1,82040183 | 0,91422333 | 0,04645886 | 0,19336739 | MYL4 |
| 71,9572716 | 1,81452926 | 0,39723267 | 4,93E-06 | 0,00019059 | YPEL3 |
| 21,8166643 | 1,80964954 | 0,57174588 | 0,00155014 | 0,01872939 | KIF13A |
| 8,02056061 | 1,80943389 | 0,84677675 | 0,03261029 | 0,15201264 | ARHGAP24 |
| 319,208881 | 1,80730739 | 0,43078839 | 2,72E-05 | 0,000798 | DENND3 |
| 13,6778748 | 1,80701348 | 0,62743113 | 0,00397651 | 0,03682801 | PDE2A |
| 6,43096858 | 1,79962333 | 0,90541953 | 0,04685454 | 0,19427204 | AMIGO1 |
| 7,76679109 | 1,79795614 | 0,7824689 | 0,02157324 | 0,11583067 | LINC00954 |
| 5,97175894 | 1,79653739 | 0,84336214 | 0,03315438 | 0,15370129 | B4GALNT3 |
| 151,140431 | 1,78249216 | 0,47841598 | 0,00019468 | 0,00386522 | FOXN3 |
| 13,0951108 | 1,78073061 | 0,57546188 | 0,00197187 | 0,02226116 | CLEC18A |
| 6,93426809 | 1,77163892 | 0,88015898 | 0,04412911 | 0,18633493 | MIR3178 |
| 12,4494804 | 1,77128261 | 0,72030999 | 0,01393029 | 0,08693106 | FRG1CP |
| 10,6256127 | 1,77094174 | 0,78669198 | 0,02437764 | 0,12552313 | NTN5 |
| 19,6987933 | 1,76827917 | 0,55300041 | 0,00138572 | 0,01719292 | CA1 |
| 118,769761 | 1,7643388 | 0,32299108 | 4,69E-08 | 3,60E-06 | KLF3 |
| 192,336253 | 1,76088035 | 0,26354839 | 2,37E-11 | 5,39E-09 | RAB11FIP1 |

| | | | | | |
|---|---|---|---|---|---|
| 15,3275604 | 1,76057245 | 0,63437564 | 0,00551536 | 0,04656673 | PPM1L |
| 138,688835 | 1,75994217 | 0,25178305 | 2,75E-12 | 8,13E-10 | TNS1 |
| 79,5303822 | 1,74927119 | 0,31865215 | 4,03E-08 | 3,16E-06 | DENND2D |
| 36,8866294 | 1,74625713 | 0,43842205 | 6,80E-05 | 0,00165678 | CFP |
| 8,38532833 | 1,74078398 | 0,87633858 | 0,04698576 | 0,19461855 | CYBRD1 |
| 103,854239 | 1,73663872 | 0,35706188 | 1,15E-06 | 5,62E-05 | TRIM22 |
| 15,1983059 | 1,73400798 | 0,63962047 | 0,00670816 | 0,05312898 | EGF |
| 32,3569848 | 1,72665558 | 0,51213267 | 0,0007476 | 0,01083103 | SORT1 |
| 78,0872598 | 1,72614523 | 0,34022975 | 3,91E-07 | 2,27E-05 | ZAP70 |
| 8,56912031 | 1,72520491 | 0,69268071 | 0,01275202 | 0,08173473 | LEF1-AS1 |
| 11,4297049 | 1,72298821 | 0,81420909 | 0,03433209 | 0,15724945 | EPB42 |
| 26,0355591 | 1,72254916 | 0,42979227 | 6,13E-05 | 0,00151542 | LAMC2 |
| 97,296249 | 1,71896288 | 0,39061235 | 1,08E-05 | 0,00037122 | DOCK11 |
| 16,0657738 | 1,71421598 | 0,63355366 | 0,00681574 | 0,05377467 | RHOBTB3 |
| 9,64069352 | 1,71383656 | 0,75642438 | 0,02346928 | 0,12256757 | LINC01215 |
| 24,2848435 | 1,7125821 | 0,45883292 | 0,00018961 | 0,00378436 | ZC3H12D |
| 16,2594444 | 1,71207035 | 0,63331711 | 0,00686469 | 0,0540519 | CAVIN3 |
| 16,6603915 | 1,71196796 | 0,5544496 | 0,00201719 | 0,02259759 | REPS2 |
| 15,4818225 | 1,70799352 | 0,69627878 | 0,01416581 | 0,08794373 | ST3GAL5 |
| 11,8152574 | 1,7052272 | 0,6397474 | 0,00768809 | 0,05816328 | BAHCC1 |
| 40,281051 | 1,69805669 | 0,39367099 | 1,61E-05 | 0,00051558 | MYL9 |
| 13,5163893 | 1,6956081 | 0,60452955 | 0,00503417 | 0,04367334 | CALD1 |
| 52,6462047 | 1,69483441 | 0,41132389 | 3,78E-05 | 0,00102802 | HK3 |
| 7,70669022 | 1,69317095 | 0,71038987 | 0,01715172 | 0,09947233 | SOX6 |
| 59,7775449 | 1,69051049 | 0,40518652 | 3,02E-05 | 0,00086231 | FCAR |
| 24,326038 | 1,68509697 | 0,50995207 | 0,00095172 | 0,0129831 | TMEM71 |
| 15,2409773 | 1,66121606 | 0,61616593 | 0,00701665 | 0,05479241 | RNF207 |
| 23,7554412 | 1,6575165 | 0,48034756 | 0,00055922 | 0,00870142 | LGALS3BP |
| 26,976758 | 1,65559284 | 0,5381027 | 0,0020929 | 0,02314491 | CXCR5 |
| 19,0514083 | 1,6533294 | 0,60310241 | 0,00611828 | 0,05007985 | TREML2 |
| 57,137348 | 1,65273571 | 0,35935222 | 4,24E-06 | 0,00016752 | ARHGAP31 |
| 8,02448566 | 1,64762208 | 0,77938257 | 0,03451446 | 0,15770813 | RRAS2 |
| 11,3657037 | 1,6458397 | 0,62687222 | 0,00865272 | 0,06299274 | FA2H |
| 8,9206742 | 1,64582451 | 0,77435122 | 0,03355127 | 0,15479724 | ST3GAL6 |
| 7,0724723 | 1,64185947 | 0,78669887 | 0,03688596 | 0,16534453 | LINC00428 |
| 6,95875286 | 1,6360054 | 0,81608832 | 0,04499603 | 0,18898332 | CDC20B |
| 42,0178032 | 1,635302 | 0,48370178 | 0,00072273 | 0,01054346 | CDH23 |
| 12,6452716 | 1,62895747 | 0,61176913 | 0,00775166 | 0,05850382 | ZNF549 |
| 17,6713381 | 1,61902158 | 0,57346107 | 0,00475401 | 0,04199375 | PKHD1L1 |
| 9,58126065 | 1,61789585 | 0,72621836 | 0,02589141 | 0,13065893 | MIR5194 |
| 20,8967614 | 1,61647881 | 0,46450073 | 0,00050135 | 0,00800984 | MUC5B |
| 20,7540452 | 1,61545885 | 0,57333996 | 0,00483798 | 0,04251715 | MFSD6L |
| 12,4076972 | 1,6137314 | 0,63682399 | 0,01127589 | 0,07529304 | TMOD2 |
| 7,47642007 | 1,61221386 | 0,79747593 | 0,04321296 | 0,18382023 | TSPEAR |
| 13,2231991 | 1,6077749 | 0,60972123 | 0,00836671 | 0,06161923 | FBN2 |
| 13,573999 | 1,60144819 | 0,67124067 | 0,01704186 | 0,09930678 | PEAR1 |
| 200,546685 | 1,59971485 | 0,25396155 | 2,99E-10 | 4,75E-08 | PIK3CD |
| 14,8513047 | 1,59753119 | 0,74323224 | 0,03159974 | 0,14895333 | LAMA5 |

| | | | | | |
|---|---|---|---|---|---|
| 24,1670736 | 1,59652122 | 0,50294397 | 0,00150171 | 0,01831975 | ABCC3 |
| 100,835668 | 1,59489462 | 0,3055632 | 1,79E-07 | 1,17E-05 | STAT4 |
| 12,9955061 | 1,5936652 | 0,70971364 | 0,02473576 | 0,12659598 | PROK2 |
| 67,9961812 | 1,59284783 | 0,35846465 | 8,85E-06 | 0,00031877 | CCDC88B |
| 8,51704674 | 1,58628888 | 0,77250255 | 0,04002977 | 0,17459734 | SNORD105 |
| 12,9257835 | 1,57388593 | 0,66316063 | 0,01762945 | 0,10132648 | SCT |
| 9,69272177 | 1,57115294 | 0,66872541 | 0,01879996 | 0,10574139 | RPS6KA2 |
| 280,787083 | 1,56361438 | 0,2483839 | 3,07E-10 | 4,76E-08 | IL10RA |
| 22,661285 | 1,56195031 | 0,49704106 | 0,00167513 | 0,01984669 | ZNF26 |
| 113,820697 | 1,5572469 | 0,37038604 | 2,62E-05 | 0,00077275 | TAGAP |
| 12,404651 | 1,5556504 | 0,64535657 | 0,01592943 | 0,09510909 | MPL |
| 15,9703965 | 1,55334348 | 0,75147399 | 0,03872829 | 0,17085056 | LOC105371409 |
| 7,85256028 | 1,55183299 | 0,77013287 | 0,04390258 | 0,18574885 | BEND4 |
| 143,387071 | 1,54933261 | 0,56888794 | 0,00646058 | 0,05179688 | S100A8 |
| 1066,97181 | 1,549126 | 0,3800573 | 4,58E-05 | 0,00120554 | HLA-B |
| 16,4245546 | 1,54372392 | 0,63926906 | 0,0157427 | 0,09448647 | LIN7A |
| 8,34574105 | 1,54219652 | 0,7629609 | 0,04324551 | 0,18387603 | MIR3183 |
| 7,05120283 | 1,53932988 | 0,77732221 | 0,04767002 | 0,19665846 | LINC01848 |
| 24,1922115 | 1,53722635 | 0,58409383 | 0,00849303 | 0,0622614 | TRPM2 |
| 430,338967 | 1,53674873 | 0,23871594 | 1,21E-10 | 2,27E-08 | CSF3R |
| 68,6222215 | 1,53641595 | 0,41800083 | 0,00023726 | 0,00449377 | HSH2D |
| 87,3606313 | 1,53211903 | 0,35102964 | 1,27E-05 | 0,00042945 | RAB11FIP4 |
| 15,0265982 | 1,51084603 | 0,59311042 | 0,01085517 | 0,07314486 | PYHIN1 |
| 14,5689435 | 1,50926426 | 0,5634294 | 0,00739063 | 0,05677248 | SSPOP |
| 147,805874 | 1,50829085 | 0,2546302 | 3,15E-09 | 3,51E-07 | SNN |
| 79,0544287 | 1,5058373 | 0,34692895 | 1,42E-05 | 0,00046473 | SCARF1 |
| 12,801277 | 1,49996698 | 0,63518683 | 0,01820324 | 0,10381722 | ADAP2 |
| 9,40006729 | 1,49935784 | 0,72967889 | 0,03989652 | 0,17421138 | PDZD4 |
| 26,3045288 | 1,49917328 | 0,49452778 | 0,00243322 | 0,02582105 | DCHS1 |
| 9,78819732 | 1,49488009 | 0,75605827 | 0,04801878 | 0,19734215 | CD109 |
| 9,69213336 | 1,49459636 | 0,73870449 | 0,04304564 | 0,18356243 | MIR27A |
| 186,703247 | 1,48801776 | 0,23534908 | 2,57E-10 | 4,23E-08 | CD6 |
| 174,59725 | 1,48749129 | 0,29203192 | 3,51E-07 | 2,10E-05 | PPP1R16B |
| 26,2012022 | 1,48731765 | 0,45851763 | 0,00117966 | 0,01521499 | N4BP3 |
| 60,714391 | 1,48650847 | 0,34315202 | 1,48E-05 | 0,00047972 | SH2D3C |
| 72,7388978 | 1,48609048 | 0,37851861 | 8,63E-05 | 0,00202446 | SH2D6 |
| 107,631938 | 1,48123621 | 0,31551278 | 2,67E-06 | 0,00011245 | SULF2 |
| 10,2934065 | 1,47990854 | 0,620928 | 0,01715439 | 0,09947233 | CPNE9 |
| 115,708538 | 1,4765483 | 0,27878799 | 1,18E-07 | 7,96E-06 | MTURN |
| 15,7705052 | 1,47556258 | 0,64056018 | 0,0212479 | 0,11478142 | HLA-F-AS1 |
| 16,5803418 | 1,47466918 | 0,50269013 | 0,00335104 | 0,03248228 | ITPKA |
| 46,3716795 | 1,47295863 | 0,40000769 | 0,00023112 | 0,00440821 | CLIP4 |
| 46526,2999 | 1,45997312 | 0,44825204 | 0,00112582 | 0,01465806 | MIR3648-1 |
| 10,7683785 | 1,43975076 | 0,62897192 | 0,02207621 | 0,11739853 | IHH |
| 15,2992456 | 1,43345801 | 0,6806861 | 0,03521289 | 0,159852 | RUNX1-IT1 |
| 181,913551 | 1,43001114 | 0,2693498 | 1,10E-07 | 7,55E-06 | LCP2 |
| 9,99460755 | 1,42137649 | 0,62811749 | 0,02364093 | 0,12307324 | CLEC18C |
| 37,2187591 | 1,42086507 | 0,37357816 | 0,00014273 | 0,00300393 | KCNK7 |

| | | | | | |
|---|---|---|---|---|---|
| 53,0802872 | 1,42000625 | 0,39709653 | 0,00034893 | 0,00606808 | GCA |
| 22,7956714 | 1,41525602 | 0,44619791 | 0,00151491 | 0,01844504 | FREY1 |
| 66,1841153 | 1,40480329 | 0,36657958 | 0,00012701 | 0,00274326 | LOC374443 |
| 87,5711029 | 1,40414963 | 0,27311865 | 2,73E-07 | 1,70E-05 | SLA2 |
| 146,985734 | 1,40025454 | 0,27127621 | 2,45E-07 | 1,55E-05 | CCR7 |
| 28,8881148 | 1,39814555 | 0,44885847 | 0,00184012 | 0,02126878 | PWWP2B |
| 24,1528857 | 1,39595902 | 0,54656231 | 0,01064715 | 0,072206 | SLC9A7 |
| 11,7137032 | 1,39344663 | 0,61298736 | 0,02301376 | 0,1209901 | CIMAP1A |
| 25,7631032 | 1,39000027 | 0,42731025 | 0,00114231 | 0,01481403 | ASTL |
| 15,5807106 | 1,3887478 | 0,62536385 | 0,02637104 | 0,1322466 | MIR616 |
| 61,2629085 | 1,38798766 | 0,35270323 | 8,31E-05 | 0,00196537 | LILRB3 |
| 18,2728128 | 1,38639648 | 0,53925122 | 0,01014153 | 0,06995549 | LASP1NB |
| 10,9578711 | 1,385224 | 0,67403178 | 0,0398663 | 0,17421138 | A2M-AS1 |
| 13,332548 | 1,38101542 | 0,64575338 | 0,03246717 | 0,15153214 | RAP1GAP |
| 24,7168068 | 1,37621411 | 0,47862183 | 0,00403556 | 0,03721095 | ZNF467 |
| 110,173233 | 1,37589073 | 0,5526985 | 0,01279569 | 0,08189429 | HLA-DPA1 |
| 75,5371327 | 1,36461859 | 0,2845757 | 1,62E-06 | 7,63E-05 | PTAFR |
| 11,6725197 | 1,36445716 | 0,6736824 | 0,04282923 | 0,18284563 | ELF3 |
| 97,386251 | 1,3505991 | 0,26872756 | 5,01E-07 | 2,76E-05 | POU2F2 |
| 24,0561737 | 1,34645205 | 0,52215704 | 0,00991917 | 0,06902563 | BTK |
| 149,491271 | 1,34502414 | 0,28537132 | 2,44E-06 | 0,00010508 | NCF2 |
| 29,5973349 | 1,33630683 | 0,41953024 | 0,00144631 | 0,01779303 | SNORA52 |
| 23,3524285 | 1,33079837 | 0,48589018 | 0,00616475 | 0,05036411 | CEND1 |
| 243,733578 | 1,32729022 | 0,23300046 | 1,22E-08 | 1,10E-06 | PTK2B |
| 26,6146959 | 1,32527586 | 0,42569979 | 0,00185089 | 0,0213392 | PYGM |
| 14,080272 | 1,32218329 | 0,60512089 | 0,02888956 | 0,14055523 | PDE7B |
| 27,3402772 | 1,31220957 | 0,40625528 | 0,00123785 | 0,0158154 | S100A16 |
| 23,8511506 | 1,31084277 | 0,56083786 | 0,01942392 | 0,10811927 | UNC5A |
| 15,7460677 | 1,30387915 | 0,58480052 | 0,02577311 | 0,13031882 | ADA |
| 521,762203 | 1,30322016 | 0,19510918 | 2,40E-11 | 5,40E-09 | TCF7 |
| 18,3521668 | 1,29882513 | 0,47511766 | 0,00626287 | 0,05093627 | ABCC6 |
| 24,6329717 | 1,28743696 | 0,52925737 | 0,01499355 | 0,0914254 | JDP2 |
| 30,0206197 | 1,28684757 | 0,39221795 | 0,00103458 | 0,01384391 | PDXP-DT |
| 9002,99224 | 1,28441253 | 0,37286078 | 0,00057159 | 0,00879266 | MIR3648-2 |
| 1032,34437 | 1,28388133 | 0,17404828 | 1,62E-13 | 6,68E-11 | TNFAIP3 |
| 38,5375344 | 1,28060643 | 0,3707737 | 0,00055259 | 0,00864283 | LOC105371934 |
| 547,104146 | 1,27793433 | 0,2029021 | 3,01E-10 | 4,75E-08 | IRF1 |
| 10,0273007 | 1,27628233 | 0,61588826 | 0,03824094 | 0,16949094 | MIR378I |
| 13,1802437 | 1,27572019 | 0,59367345 | 0,03164563 | 0,14905823 | IFITM10 |
| 126,503827 | 1,27226167 | 0,34935677 | 0,00027082 | 0,00499426 | ARL4C |
| 35,2141829 | 1,26891938 | 0,42898484 | 0,00309684 | 0,03074253 | MPP7 |
| 46,3984428 | 1,26735543 | 0,35730072 | 0,0003896 | 0,00659955 | CLIC3 |
| 82,6353206 | 1,26107732 | 0,43437969 | 0,00369408 | 0,03491294 | EBF4 |
| 10,5808212 | 1,26090787 | 0,59768768 | 0,03488908 | 0,15886602 | ORM2 |
| 26,550881 | 1,26057172 | 0,52017821 | 0,01537828 | 0,09299214 | SNX20 |
| 16,939495 | 1,24993374 | 0,58398914 | 0,03232752 | 0,15121607 | SLFN14 |
| 15,7680206 | 1,24876969 | 0,55307959 | 0,02395513 | 0,12397567 | CACTIN-AS1 |
| 14,9044155 | 1,23930711 | 0,6262261 | 0,04781499 | 0,19697527 | SSTR3 |

| | | | | | |
|---|---|---|---|---|---|
| 58,3240291 | 1,23786709 | 0,38906726 | 0,0014645 | 0,01795193 | JAG2 |
| 27,475402 | 1,22792356 | 0,53486996 | 0,02169062 | 0,11622038 | RNF43 |
| 25,4264489 | 1,22655437 | 0,45097555 | 0,00653253 | 0,05228513 | HSFX4 |
| 14,7713853 | 1,22357591 | 0,61543251 | 0,04679443 | 0,19413398 | PYCARD |
| 28,230682 | 1,22292434 | 0,43365084 | 0,00480137 | 0,04233301 | PATL2 |
| 14,3520765 | 1,2159028 | 0,59237736 | 0,04011319 | 0,17471952 | MIR4651 |
| 471,872867 | 1,2038459 | 0,54012885 | 0,02582639 | 0,13049472 | S100A9 |
| 30,1905682 | 1,20125764 | 0,47126522 | 0,01080306 | 0,07294956 | KLRB1 |
| 16,6153327 | 1,20074528 | 0,58081166 | 0,03870049 | 0,17084739 | RSAD2 |
| 22,4576846 | 1,19971129 | 0,48198633 | 0,01280676 | 0,08189429 | EPHA1 |
| 15,560244 | 1,18711903 | 0,56328259 | 0,03507411 | 0,15942156 | CFAP45 |
| 76,7933603 | 1,1819321 | 0,30869886 | 0,00012879 | 0,00277236 | TNFRSF25 |
| 45,9594325 | 1,17628883 | 0,3651765 | 0,00127677 | 0,01614335 | KLF4 |
| 19,4004363 | 1,17320625 | 0,48611341 | 0,01580267 | 0,09465092 | PMEL |
| 123,204875 | 1,17267813 | 0,32310766 | 0,00028411 | 0,00516394 | TGFBI |
| 61,6053332 | 1,17064244 | 0,3649807 | 0,00133936 | 0,01676065 | PHOSPHO1 |
| 85,8752959 | 1,16719705 | 0,23352584 | 5,79E-07 | 3,14E-05 | ENG |
| 210,547325 | 1,16637904 | 0,23636222 | 8,03E-07 | 4,10E-05 | KIAA0513 |
| 264,362614 | 1,16340952 | 0,20510213 | 1,41E-08 | 1,26E-06 | ITPKB |
| 39,1364139 | 1,16312199 | 0,37032399 | 0,00168474 | 0,01993556 | SIGLEC10 |
| 287,415597 | 1,16238968 | 0,22911743 | 3,91E-07 | 2,27E-05 | BTG2 |
| 21,5061428 | 1,16202357 | 0,45416287 | 0,0105093 | 0,07156625 | IFIT1 |
| 16,8775451 | 1,16157697 | 0,49276316 | 0,01840974 | 0,10447831 | CD72 |
| 46,2972598 | 1,14418579 | 0,42807877 | 0,00752123 | 0,0574754 | IFI44 |
| 20,5168187 | 1,14105999 | 0,47170376 | 0,01556246 | 0,09383599 | IL1B |
| 333,647374 | 1,13017356 | 0,20836345 | 5,83E-08 | 4,29E-06 | SLC43A2 |
| 25,507869 | 1,12693223 | 0,47157733 | 0,01686176 | 0,09868277 | TNFRSF13C |
| 109,719882 | 1,11952463 | 0,29538431 | 0,00015061 | 0,00314189 | PDE4B |
| 29,1571495 | 1,11166184 | 0,42011863 | 0,00814338 | 0,06048764 | PLPPR3 |
| 19,6733016 | 1,11042368 | 0,49715696 | 0,02551286 | 0,12961785 | MYO15A |
| 24,9144504 | 1,11025772 | 0,51509964 | 0,03112873 | 0,1477581 | ST14 |
| 35,7153397 | 1,1003236 | 0,38980154 | 0,00476094 | 0,04203533 | FRG2B |
| 34,0692045 | 1,09264389 | 0,40935918 | 0,00760419 | 0,05785321 | CSGALNACT1 |
| 28,1605501 | 1,09193531 | 0,45219062 | 0,01574525 | 0,09448647 | SLC25A34 |
| 18,2356289 | 1,08749282 | 0,52408764 | 0,03798459 | 0,16880498 | BPIFA2 |
| 63,3017824 | 1,08619065 | 0,34043106 | 0,00141961 | 0,01754442 | CARMIL2 |
| 26,7394789 | 1,08280408 | 0,44478426 | 0,01491453 | 0,09119843 | SCN1B |
| 35,6681303 | 1,08222881 | 0,37378318 | 0,0037875 | 0,03551355 | NAPSB |
| 139,880623 | 1,08032301 | 0,22937439 | 2,48E-06 | 0,00010635 | MYLIP |
| 26,0462964 | 1,07960488 | 0,43855476 | 0,01382674 | 0,08654178 | LINC02362 |
| 30,8511079 | 1,078725 | 0,40768523 | 0,00814558 | 0,06048764 | PLAGL1 |
| 16,1807355 | 1,07632198 | 0,50304131 | 0,03238473 | 0,15133402 | SRGAP3 |
| 21,7056703 | 1,06882523 | 0,44740484 | 0,01689686 | 0,09873551 | SUSD3 |
| 129,303336 | 1,06873915 | 0,2179549 | 9,41E-07 | 4,70E-05 | CD83 |
| 33,6860851 | 1,06767364 | 0,3771952 | 0,00464666 | 0,04127674 | CXCR3 |
| 19,472959 | 1,06736677 | 0,53360196 | 0,04546732 | 0,19041326 | CHRND |
| 31,9566722 | 1,06518196 | 0,40807034 | 0,00904655 | 0,06496005 | SPINT2 |
| 41,9403104 | 1,05681636 | 0,40989604 | 0,00993008 | 0,06903373 | LOC102723566 |

| | | | | | |
|---|---|---|---|---|---|
| 101,859801 | 1,05565101 | 0,24935888 | 2,30E-05 | 0,00069334 | SEMA4A |
| 60,9004205 | 1,05449597 | 0,40991462 | 0,01009736 | 0,06985025 | LGALS3 |
| 97,4755112 | 1,04922381 | 0,28898284 | 0,0002826 | 0,00514144 | CLEC2D |
| 17,553061 | 1,04066228 | 0,51141334 | 0,04186344 | 0,17961409 | ARHGAP26-AS1 |
| 895,825749 | 1,03949422 | 0,18826145 | 3,36E-08 | 2,71E-06 | LRP1 |
| 119,918943 | 1,03698939 | 0,31488768 | 0,00099053 | 0,01338695 | MPIG6B |
| 57,163964 | 1,03369009 | 0,38280815 | 0,00692807 | 0,05441508 | PILRA |
| 319,26809 | 1,03309506 | 0,23686088 | 1,29E-05 | 0,00043417 | SPARC |
| 22,4921836 | 1,02206962 | 0,44985699 | 0,02308724 | 0,12127533 | ITIH4 |
| 30,3833725 | 1,01818529 | 0,41113936 | 0,01326788 | 0,08384824 | DNAJB13 |
| 30,7711503 | 1,01452441 | 0,39641266 | 0,01048942 | 0,07154663 | LMTK3 |
| 22,5200299 | 1,00712738 | 0,4487264 | 0,02480584 | 0,12692029 | MIR637 |
| 36,2746239 | 1,00120814 | 0,38309753 | 0,00896319 | 0,06448384 | PZP |
| 68,4570729 | 0,99622277 | 0,28486386 | 0,00047019 | 0,00763643 | NCF4 |
| 27,91097 | 0,99199693 | 0,41378488 | 0,01651308 | 0,09742562 | IGFALS |
| 347,883924 | 0,99073621 | 0,18818978 | 1,41E-07 | 9,33E-06 | FOSB |
| 69,1651607 | 0,98793149 | 0,27176351 | 0,0002777 | 0,00508156 | ZNF185 |
| 18,0644031 | 0,98741665 | 0,49308648 | 0,04522859 | 0,18958135 | SNORD81 |
| 90,5459259 | 0,98700372 | 0,25705463 | 0,0001232 | 0,00268252 | SYTL1 |
| 86,6100096 | 0,98598558 | 0,26980363 | 0,00025773 | 0,0047822 | SNORA64 |
| 20,8716783 | 0,98234895 | 0,44604943 | 0,02764186 | 0,13646992 | SLC43A1 |
| 82,7634213 | 0,98008266 | 0,28843321 | 0,00067892 | 0,01003581 | CRISPLD2 |
| 57,4063761 | 0,97882612 | 0,28508053 | 0,00059583 | 0,00907094 | CAMKK1 |
| 89,728638 | 0,97851718 | 0,2366137 | 3,54E-05 | 0,00097403 | SNORD46 |
| 91,7748474 | 0,97499911 | 0,31390747 | 0,00189632 | 0,02169298 | PTGER4 |
| 22,5029428 | 0,97463998 | 0,45182688 | 0,03099715 | 0,14728705 | OBSCN-AS1 |
| 156,218765 | 0,97236353 | 0,27103065 | 0,00033367 | 0,00585478 | TCP11L2 |
| 255,916645 | 0,96505306 | 0,20918484 | 3,96E-06 | 0,00015813 | MAP4K2 |
| 63,115707 | 0,96388163 | 0,32830151 | 0,00332512 | 0,03236349 | BAIAP3 |
| 33,7297992 | 0,96349553 | 0,39167365 | 0,01389584 | 0,08680198 | RIMKLA |
| 36,2808703 | 0,96261618 | 0,40217229 | 0,01668658 | 0,09805887 | MFSD2B |
| 33,1591764 | 0,95196542 | 0,36604898 | 0,00930472 | 0,0661114 | HIF3A |
| 44,702067 | 0,94673491 | 0,3958088 | 0,01676143 | 0,09824751 | EPPK1 |
| 41,755463 | 0,94611584 | 0,37102814 | 0,01077279 | 0,07283299 | NHSL2 |
| 23,8824642 | 0,94560525 | 0,47795319 | 0,04787826 | 0,19709161 | SCARF2 |
| 25,0563644 | 0,94382267 | 0,44609642 | 0,03436719 | 0,15733403 | SNORD37 |
| 65,6625026 | 0,93975892 | 0,28561499 | 0,00100081 | 0,01346819 | GATA3 |
| 86,5125775 | 0,93907069 | 0,29799033 | 0,0016252 | 0,01944137 | FAM117A |
| 459,08466 | 0,92897656 | 0,17033867 | 4,93E-08 | 3,73E-06 | CORO1A |
| 31,8038774 | 0,92885201 | 0,41568406 | 0,02544925 | 0,12933795 | MTSS2 |
| 35,5135583 | 0,92552267 | 0,39369774 | 0,0187308 | 0,10554062 | HLA-DMA |
| 390,818502 | 0,91914799 | 0,18157223 | 4,15E-07 | 2,36E-05 | MYO15B |
| 225,758269 | 0,91609032 | 0,21236338 | 1,60E-05 | 0,00051558 | SYNE1 |
| 226,074943 | 0,91368785 | 0,21133511 | 1,54E-05 | 0,00049692 | STAT5A |
| 81,9528103 | 0,91186869 | 0,33178235 | 0,0059888 | 0,04930259 | MIR4784 |
| 39,6151799 | 0,9117349 | 0,40497111 | 0,02436288 | 0,12550233 | IQSEC2 |
| 25,9068636 | 0,91038005 | 0,43859294 | 0,03792325 | 0,16860887 | MIR661 |
| 445,443951 | 0,90806527 | 0,20789521 | 1,25E-05 | 0,00042567 | CELF2 |

| | | | | | |
|---|---|---|---|---|---|
| 78,0168808 | 0,90304912 | 0,27144818 | 0,00087855 | 0,01226791 | SCART1 |
| 47,2394391 | 0,90260901 | 0,40100409 | 0,02439362 | 0,12555815 | LGMN |
| 37,5444932 | 0,90260533 | 0,39106803 | 0,02099625 | 0,11392889 | MAPK8IP1 |
| 20,6582654 | 0,90147162 | 0,4274904 | 0,03496592 | 0,15907433 | ORAI3 |
| 65,7731023 | 0,90066627 | 0,29936623 | 0,00262475 | 0,02742286 | SMPD3 |
| 52,7759429 | 0,90053088 | 0,34889657 | 0,0098491 | 0,06874025 | SMIM32 |
| 105,304766 | 0,89381278 | 0,20879908 | 1,86E-05 | 0,00057875 | SAP25 |
| 32,416667 | 0,89215227 | 0,35636701 | 0,01229839 | 0,07980037 | PLCD1 |
| 104,108149 | 0,88976005 | 0,2047805 | 1,39E-05 | 0,00046002 | CTSW |
| 42,7545474 | 0,88688802 | 0,33927407 | 0,00894693 | 0,06444035 | CPNE2 |
| 31,7362278 | 0,88009223 | 0,39404718 | 0,02551802 | 0,12961785 | APOBEC3D |
| 44,410816 | 0,87527609 | 0,36060518 | 0,0152141 | 0,09235352 | OGFR-AS1 |
| 76,5686112 | 0,8743846 | 0,31750139 | 0,00588798 | 0,04867039 | GPX3 |
| 39,390781 | 0,87178896 | 0,36436122 | 0,01672718 | 0,09813796 | TMOD1 |
| 189,809343 | 0,86886851 | 0,24020907 | 0,00029788 | 0,00535255 | SMIM5 |
| 21,9167808 | 0,86480301 | 0,42230653 | 0,04057874 | 0,17605585 | C20orf181 |
| 50,106566 | 0,86084417 | 0,3512753 | 0,01426083 | 0,08832377 | PANX2 |
| 43,6396914 | 0,86029827 | 0,32991501 | 0,00911698 | 0,06525235 | NRSN2 |
| 33,9265378 | 0,86007019 | 0,35911433 | 0,01662146 | 0,09786557 | PRR25 |
| 33,2153728 | 0,85913646 | 0,40324568 | 0,03312595 | 0,15362159 | FGF8 |
| 140,040926 | 0,85900118 | 0,25880925 | 0,00090324 | 0,01245639 | MIEF2 |
| 24,138443 | 0,85624077 | 0,42705431 | 0,04496383 | 0,18889002 | ANGPTL1 |
| 50,7430801 | 0,85467636 | 0,33017183 | 0,00963724 | 0,06783714 | GRIN3B |
| 235,852519 | 0,85400367 | 0,20766989 | 3,92E-05 | 0,00105579 | TAMALIN |
| 446,599704 | 0,85363965 | 0,35141945 | 0,01513557 | 0,09210549 | ITGB3 |
| 30,1120784 | 0,85343127 | 0,37343309 | 0,02229138 | 0,11801825 | MBOAT4 |
| 75,8968557 | 0,85264835 | 0,34589682 | 0,01369992 | 0,08591853 | IFIT2 |
| 72,6466893 | 0,85184039 | 0,36177382 | 0,01854158 | 0,10488052 | ITGA9 |
| 80,6376265 | 0,8508682 | 0,3123124 | 0,00644158 | 0,05171486 | TMEM249 |
| 150,718293 | 0,85004242 | 0,26490231 | 0,00133248 | 0,01669664 | GP1BA |
| 80,9493681 | 0,8483611 | 0,36010919 | 0,01848067 | 0,10469228 | PDCD1 |
| 140,670406 | 0,84759803 | 0,22765096 | 0,00019669 | 0,00388872 | FES |
| 100,927789 | 0,84430555 | 0,30533588 | 0,00568932 | 0,0476106 | FCGR2A |
| 57,0285584 | 0,84399149 | 0,32237339 | 0,00884324 | 0,0639858 | MAP4K1 |
| 147,880295 | 0,83692573 | 0,2322237 | 0,0003134 | 0,00554411 | BIN1 |
| 90,5175144 | 0,83491103 | 0,25743715 | 0,0011821 | 0,01521873 | MIR5587 |
| 98,8560647 | 0,83409503 | 0,2351502 | 0,00038954 | 0,00659955 | SYTL2 |
| 53,7144242 | 0,83356254 | 0,39312385 | 0,03397603 | 0,15625767 | SLCO3A1 |
| 49,6343839 | 0,833003 | 0,29823559 | 0,00522049 | 0,04483746 | GRIN2C |
| 685,646371 | 0,832399 | 0,13440657 | 5,90E-10 | 8,03E-08 | ZFP36L2 |
| 619,105792 | 0,8309187 | 0,18091799 | 4,37E-06 | 0,00017204 | ITGB2 |
| 142,957213 | 0,83031605 | 0,19474967 | 2,01E-05 | 0,00061915 | TSPAN4 |
| 83,0228861 | 0,82981147 | 0,31234753 | 0,00789113 | 0,05924922 | PLIN4 |
| 39,5092798 | 0,82704544 | 0,39811289 | 0,03776333 | 0,16816191 | C17orf107 |
| 72,836529 | 0,82639047 | 0,30959191 | 0,00760119 | 0,05785321 | HACD4 |
| 71,8969343 | 0,8236498 | 0,26647664 | 0,00199558 | 0,02247522 | TBXAS1 |
| 27,922772 | 0,82326508 | 0,38926697 | 0,03443752 | 0,15759105 | RAB44 |
| 43,6320994 | 0,82058368 | 0,29968324 | 0,00617821 | 0,05040878 | SLED1 |

| | | | | | |
|---|---|---|---|---|---|
| 32,2697426 | 0,82050061 | 0,37035652 | 0,02673026 | 0,13358174 | GOLGA8K |
| 48,0343153 | 0,82018669 | 0,32758106 | 0,01228789 | 0,07978693 | PINK1 |
| 44,4469678 | 0,81797087 | 0,31711648 | 0,00989716 | 0,06893294 | PDF |
| 91,4596301 | 0,81513647 | 0,30369895 | 0,0072741 | 0,05622275 | LINC00926 |
| 161,593826 | 0,81297099 | 0,25364028 | 0,00134963 | 0,01685566 | NLRC3 |
| 28,3669859 | 0,81150542 | 0,37852118 | 0,0320422 | 0,15043931 | LTC4S |
| 34,8696853 | 0,81077144 | 0,40201717 | 0,04372075 | 0,18523069 | RAB40AL |
| 69,5562867 | 0,80868773 | 0,28972634 | 0,0052511 | 0,04496241 | CDHR5 |
| 54,8533736 | 0,80865986 | 0,35611592 | 0,02316048 | 0,12155878 | DERL3 |
| 90,9517651 | 0,80583865 | 0,30577202 | 0,00840325 | 0,06179472 | KCNQ1 |
| 71,49604 | 0,80418214 | 0,27788678 | 0,00380465 | 0,03561988 | CARNS1 |
| 33,1359413 | 0,80399988 | 0,36220477 | 0,02643652 | 0,13250475 | C16orf86 |
| 60,4270617 | 0,80376807 | 0,33341715 | 0,01592203 | 0,0950949 | SNORD23 |
| 92,0638988 | 0,80235577 | 0,23107703 | 0,00051613 | 0,00817893 | DUSP8 |
| 71,2498296 | 0,80161595 | 0,37936969 | 0,03459901 | 0,15801299 | MEF2C |
| 89,2106522 | 0,80046302 | 0,23118587 | 0,00053534 | 0,0084197 | AATBC |
| 52,8491999 | 0,8004277 | 0,34163787 | 0,01913386 | 0,1070783 | RAPSN |
| 119,060339 | 0,80041929 | 0,2492915 | 0,00132377 | 0,01659843 | CSNK1G2-AS1 |
| 84,1338095 | 0,79901439 | 0,27958386 | 0,00426495 | 0,03877934 | PITPNM3 |
| 136,390355 | 0,79195366 | 0,24577499 | 0,00127179 | 0,01611754 | RIN1 |
| 60,3821046 | 0,79175316 | 0,2819808 | 0,00498771 | 0,04336969 | MRS2P2 |
| 580,32963 | 0,79019527 | 0,14497638 | 5,02E-08 | 3,79E-06 | EEIG1 |
| 81,1484016 | 0,78929599 | 0,29172677 | 0,00681811 | 0,05377467 | TAGLN |
| 90,3257292 | 0,78852927 | 0,28252262 | 0,00525405 | 0,04496241 | TMPRSS9 |
| 202,757601 | 0,78773934 | 0,19091348 | 3,69E-05 | 0,00100568 | MAFF |
| 181,640617 | 0,78761169 | 0,3079526 | 0,0105405 | 0,07166253 | COL1A1 |
| 37,0766098 | 0,78706422 | 0,35142887 | 0,02511614 | 0,12798881 | PRDM12 |
| 79,0276541 | 0,7866515 | 0,28233077 | 0,00533174 | 0,04540139 | LSMEM2 |
| 138,361759 | 0,78367587 | 0,27618433 | 0,00454669 | 0,040498 | CIITA |
| 104,613719 | 0,7835627 | 0,26311589 | 0,00290123 | 0,02944967 | ACP4 |
| 65,3584639 | 0,78237999 | 0,33606303 | 0,01990811 | 0,11007639 | SPON2 |
| 71,1079444 | 0,78193933 | 0,24624219 | 0,00149584 | 0,01825989 | CACNA1C |
| 383,047249 | 0,78073545 | 0,14861636 | 1,49E-07 | 9,85E-06 | H2AB1 |
| 43,0292187 | 0,7784483 | 0,37244078 | 0,03660645 | 0,16444223 | GMFG |
| 30,4150426 | 0,77475064 | 0,38980104 | 0,04686103 | 0,19427204 | MIR630 |
| 34,9918814 | 0,77146006 | 0,33992721 | 0,02323879 | 0,12182235 | STAR |
| 55,776476 | 0,77077034 | 0,31437298 | 0,01421554 | 0,08816323 | LINC00896 |
| 60,1909029 | 0,76577244 | 0,27610924 | 0,00554671 | 0,04674798 | MBNL1-AS1 |
| 70,7146628 | 0,76419804 | 0,26732703 | 0,0042543 | 0,03873704 | RNF144B |
| 162,289376 | 0,76116989 | 0,22736503 | 0,00081459 | 0,01159738 | ZCCHC24 |
| 37,2861923 | 0,75944251 | 0,3392053 | 0,02516323 | 0,12819428 | MIR7111 |
| 248,436767 | 0,75941448 | 0,24680408 | 0,002091 | 0,02314491 | ADD3 |
| 61,10494 | 0,75887157 | 0,26002783 | 0,00351805 | 0,03372089 | CCDC163 |
| 505,740604 | 0,75710174 | 0,17420418 | 1,39E-05 | 0,00045999 | PIK3IP1 |
| 59,8041488 | 0,75546914 | 0,29090181 | 0,00940446 | 0,06664488 | ZXDA |
| 435,055072 | 0,75225655 | 0,17157223 | 1,16E-05 | 0,00039778 | MIR23AHG |
| 91,575754 | 0,75130016 | 0,21408339 | 0,00044915 | 0,00737067 | ENDOG |
| 89,0208818 | 0,74699332 | 0,29708165 | 0,01192223 | 0,07827276 | NKD1 |

| | | | | | |
|---|---|---|---|---|---|
| 94,0809151 | 0,74621583 | 0,21154752 | 0,00041964 | 0,00700794 | AJM1 |
| 151,505315 | 0,74551073 | 0,22580487 | 0,00096145 | 0,01306796 | TRANK1 |
| 44,6181943 | 0,74454254 | 0,36341016 | 0,04048496 | 0,17573201 | UST |
| 92,4021298 | 0,7430354 | 0,26812659 | 0,00558482 | 0,04694374 | SPEF2 |
| 745,188 | 0,74246958 | 0,14653712 | 4,05E-07 | 2,33E-05 | H2AB3 |
| 277,911751 | 0,74211061 | 0,18614548 | 6,70E-05 | 0,00163555 | SPATA2L |
| 71,4747624 | 0,74205352 | 0,28522411 | 0,00927763 | 0,06596843 | ALPK3 |
| 28,9086123 | 0,73856008 | 0,36140299 | 0,04099392 | 0,1772905 | SEMA3B |
| 157,905601 | 0,73847322 | 0,21759041 | 0,00068913 | 0,01015498 | PRKCH |
| 730,055831 | 0,7381664 | 0,14937562 | 7,75E-07 | 3,99E-05 | H2AB2 |
| 41,0427632 | 0,73769995 | 0,3216765 | 0,02183088 | 0,11663537 | SNORA14B |
| 62,9992882 | 0,7373636 | 0,31377308 | 0,01877392 | 0,10567479 | LOC100652777 |
| 49,0187374 | 0,73693205 | 0,28544418 | 0,00983142 | 0,06869287 | PLCH2 |
| 61,6575282 | 0,73545053 | 0,27388685 | 0,00724788 | 0,05609914 | DLEC1 |
| 205,167924 | 0,73382913 | 0,28794643 | 0,01081898 | 0,07303102 | LOC105372714 |
| 81,4996047 | 0,73313878 | 0,3317522 | 0,02711219 | 0,13506838 | GNG3 |
| 185,368577 | 0,73280777 | 0,19428628 | 0,00016208 | 0,00333662 | NOXO1 |
| 39,4578975 | 0,7309115 | 0,33467586 | 0,02896677 | 0,14071379 | GLRA3 |
| 482,041169 | 0,73036469 | 0,12286186 | 2,77E-09 | 3,16E-07 | TTC7A |
| 71,0835326 | 0,72982437 | 0,30460966 | 0,01657812 | 0,09771792 | LINC00106 |
| 62,3862715 | 0,72917589 | 0,3578448 | 0,04158119 | 0,17884921 | ADAP1 |
| 75,5871456 | 0,72910225 | 0,27116399 | 0,00717121 | 0,05570054 | LOC105372824 |
| 55,2310349 | 0,72905261 | 0,34325851 | 0,03367708 | 0,15515071 | PODNL1 |
| 91,560688 | 0,72782383 | 0,26838347 | 0,00669028 | 0,05307793 | CTSH |
| 287,159543 | 0,72779028 | 0,22188818 | 0,00103812 | 0,01388146 | PARVG |
| 84,279167 | 0,72660178 | 0,26217862 | 0,00558159 | 0,04694374 | LHB |
| 59,4209594 | 0,72497969 | 0,30414004 | 0,01713941 | 0,09945292 | RASGRP4 |
| 1427,76819 | 0,72487424 | 0,15951514 | 5,51E-06 | 0,00020823 | TXNIP |
| 42,2825197 | 0,72019996 | 0,30694211 | 0,01895725 | 0,10637308 | SNORD99 |
| 287,24303 | 0,71875568 | 0,15549963 | 3,80E-06 | 0,00015283 | IRF7 |
| 57,2261736 | 0,7184452 | 0,31290267 | 0,0216721 | 0,1161637 | SP6 |
| 1131,05206 | 0,71531425 | 0,12491506 | 1,03E-08 | 9,47E-07 | RELT |
| 53,6305225 | 0,71346177 | 0,29060617 | 0,01408527 | 0,08755589 | ITGB2-AS1 |
| 235,779842 | 0,71305837 | 0,22867215 | 0,00181924 | 0,02107895 | APC2 |
| 60,8570395 | 0,71104985 | 0,29577845 | 0,01621701 | 0,09617778 | RSPH6A |
| 70,9691764 | 0,71103983 | 0,25857543 | 0,00596252 | 0,04920057 | OR2W3 |
| 146,188298 | 0,71091418 | 0,27801364 | 0,01055429 | 0,07173048 | PIEZO1-AS1 |
| 64,2238021 | 0,71079094 | 0,28544659 | 0,01277067 | 0,0817991 | CAPS |
| 75,7057278 | 0,71077234 | 0,24023935 | 0,00309039 | 0,03071076 | CEMP1 |
| 107,012888 | 0,71059159 | 0,28644024 | 0,01310999 | 0,08315594 | TNK1 |
| 40,3505021 | 0,71033956 | 0,33151391 | 0,03213606 | 0,15073037 | NYAP1 |
| 118,410921 | 0,71004112 | 0,2498221 | 0,00448052 | 0,04013226 | MTSS1 |
| 55,1177113 | 0,70702599 | 0,27578152 | 0,01035577 | 0,07092602 | DDX11L2 |
| 106,793428 | 0,70689193 | 0,21166224 | 0,00083864 | 0,01182211 | LINC00863 |
| 1088,57123 | 0,70515007 | 0,11968368 | 3,82E-09 | 4,15E-07 | ATG2A |
| 40,0900264 | 0,70492523 | 0,31779125 | 0,02654105 | 0,13281758 | HAL |
| 266,840274 | 0,70480309 | 0,1922648 | 0,00024657 | 0,00462379 | BCL11B |
| 64,830694 | 0,70143648 | 0,25190844 | 0,0053612 | 0,04558586 | SNORA80B |

| | | | | | |
|---|---|---|---|---|---|
| 167,309775 | 0,70025226 | 0,18831634 | 0,00020042 | 0,00393781 | CBX7 |
| 178,429968 | 0,69952957 | 0,21927637 | 0,00142188 | 0,01754946 | YPEL2 |
| 183,307104 | 0,69893183 | 0,20120392 | 0,00051324 | 0,00815288 | ABTB1 |
| 231,292483 | 0,69830051 | 0,20066824 | 0,00050165 | 0,00800984 | RASSF7 |
| 43,821914 | 0,69607181 | 0,32726654 | 0,03342621 | 0,15440852 | TMEM150B |
| 70,909078 | 0,69138557 | 0,30076906 | 0,02152052 | 0,11564605 | LRIG1 |
| 58,728865 | 0,69036585 | 0,28527953 | 0,01552209 | 0,09362242 | ESRRB |
| 298,31344 | 0,68886626 | 0,17161598 | 5,97E-05 | 0,00148823 | MIR22HG |
| 198,70587 | 0,68864824 | 0,28533536 | 0,01580144 | 0,09465092 | SATB1 |
| 125,794233 | 0,68850155 | 0,20900275 | 0,00098694 | 0,01335762 | ZNF688 |
| 142,20951 | 0,68803012 | 0,24650981 | 0,00525315 | 0,04496241 | RHBDL1 |
| 44,8603771 | 0,6875775 | 0,3127758 | 0,02792717 | 0,13742784 | ANXA4 |
| 251,058166 | 0,68696793 | 0,20007979 | 0,00059591 | 0,00907094 | PBXIP1 |
| 172,735059 | 0,68462542 | 0,21852822 | 0,00173093 | 0,02030244 | FGL2 |
| 183,666488 | 0,68407724 | 0,18000934 | 0,00014456 | 0,00303567 | ADAMTSL4-AS1 |
| 62,5290614 | 0,68396649 | 0,26406731 | 0,0095942 | 0,06766002 | KIAA1614 |
| 278,021773 | 0,68233826 | 0,17811379 | 0,00012767 | 0,00275442 | TMEM198 |
| 67,3828624 | 0,6819135 | 0,25624587 | 0,00778699 | 0,05870028 | RANGRF |
| 385,769135 | 0,67864605 | 0,1854128 | 0,00025203 | 0,00471209 | SLC26A11 |
| 151,944363 | 0,6771976 | 0,22403606 | 0,00250516 | 0,02637741 | NKPD1 |
| 1410,02802 | 0,67614747 | 0,16672283 | 5,00E-05 | 0,00129656 | PTPRC |
| 108,592008 | 0,67510906 | 0,26246002 | 0,01010441 | 0,06985025 | PLAC8 |
| 41,9821004 | 0,67496802 | 0,31600795 | 0,03268585 | 0,1521774 | CCDC148 |
| 41,5930628 | 0,67283094 | 0,34107761 | 0,04853405 | 0,19881203 | C1R |
| 334,646573 | 0,67220088 | 0,18342024 | 0,00024752 | 0,00463694 | FBXW9 |
| 45,2055404 | 0,67197866 | 0,33455661 | 0,0445833 | 0,18783357 | GVINP1 |
| 209,765275 | 0,67104947 | 0,23135748 | 0,00372583 | 0,03512525 | ASPG |
| 418,541483 | 0,66937405 | 0,14690815 | 5,20E-06 | 0,0001993 | AMDHD2 |
| 92,5044407 | 0,6679221 | 0,33460926 | 0,04592025 | 0,19180069 | NOXA1 |
| 87,6583166 | 0,66510859 | 0,24769668 | 0,00724922 | 0,05609914 | IFIT3 |
| 186,099919 | 0,66497226 | 0,15656246 | 2,16E-05 | 0,00065914 | FAM110A |
| 87,9612645 | 0,66471501 | 0,32391517 | 0,04015734 | 0,17473983 | POU5F1P4 |
| 40,260476 | 0,66441033 | 0,33782756 | 0,04921615 | 0,20052083 | CARINH |
| 208,606488 | 0,66408494 | 0,22018467 | 0,00256103 | 0,02687586 | TEKTIP1 |
| 80,1103368 | 0,66296175 | 0,29246231 | 0,02340074 | 0,12234467 | PAX5 |
| 84,7676946 | 0,65813507 | 0,28006584 | 0,01877695 | 0,10567479 | ISG20 |
| 345,864582 | 0,65806011 | 0,1712957 | 0,0001222 | 0,00267638 | PLEKHA2 |
| 214,150628 | 0,65562803 | 0,27987985 | 0,0191533 | 0,10712371 | RYR2 |
| 56,0991895 | 0,65453181 | 0,27923636 | 0,01907784 | 0,10682802 | LOC100652768 |
| 335,184182 | 0,65439301 | 0,18097125 | 0,00029918 | 0,00536575 | HSALR1 |
| 1300,69452 | 0,6532013 | 0,17239067 | 0,00015121 | 0,0031509 | SORL1 |
| 118,54449 | 0,65217051 | 0,20025549 | 0,00112719 | 0,01465806 | DNM3 |
| 54,2730567 | 0,6520762 | 0,30292422 | 0,03134974 | 0,14821798 | RUNDC3A |
| 776,018597 | 0,65085541 | 0,14568171 | 7,91E-06 | 0,00028775 | MXD4 |
| 69,9636723 | 0,65059258 | 0,31901983 | 0,04141527 | 0,178445 | CBY2 |
| 813,630811 | 0,64984155 | 0,1675347 | 0,00010495 | 0,00237306 | DGKQ |
| 69,8276772 | 0,6494352 | 0,2876296 | 0,02395269 | 0,12397567 | MAFB |
| 179,980832 | 0,64926146 | 0,22285864 | 0,00357593 | 0,03417179 | CFAP157 |

| | | | | | |
|---|---|---|---|---|---|
| 56,8136782 | 0,64880749 | 0,32164671 | 0,04368057 | 0,18510191 | GGTLC2 |
| 56,0557599 | 0,64826204 | 0,3131022 | 0,03841035 | 0,16993845 | SPATA21 |
| 105,886551 | 0,64799913 | 0,24029351 | 0,00700307 | 0,05477375 | OSBPL5 |
| 115,574225 | 0,64766881 | 0,27252003 | 0,01747342 | 0,10067438 | RTEL1 |
| 92,601217 | 0,64632255 | 0,28587188 | 0,02376657 | 0,12357443 | ABCB9 |
| 407,508285 | 0,64600664 | 0,16736217 | 0,00011342 | 0,00253065 | AOAH |
| 558,174752 | 0,64553461 | 0,17566093 | 0,00023795 | 0,00450229 | DNAH1 |
| 95,3008269 | 0,64499735 | 0,26386329 | 0,01450782 | 0,08929811 | SNORD139 |
| 700,079 | 0,64498641 | 0,1523756 | 2,31E-05 | 0,00069405 | HAGHL |
| 290,790205 | 0,64467021 | 0,15345173 | 2,66E-05 | 0,00078031 | PHETA1 |
| 112,182812 | 0,64445545 | 0,308326 | 0,03660203 | 0,16444223 | SNORD26 |
| 217,336351 | 0,64315459 | 0,24795072 | 0,00948994 | 0,06708052 | SPOCD1 |
| 192,233469 | 0,64254301 | 0,16571942 | 0,00010562 | 0,00238487 | NKAIN3 |
| 153,915654 | 0,64240067 | 0,20551847 | 0,00177348 | 0,02072331 | SAC3D1 |
| 86,1160181 | 0,64180412 | 0,25624769 | 0,01225816 | 0,07974787 | LINC02613 |
| 1210,77503 | 0,63707545 | 0,15680215 | 4,85E-05 | 0,00126126 | AKNA |
| 833,852607 | 0,63676232 | 0,16425892 | 0,00010594 | 0,00238906 | FBXL6 |
| 90,4996972 | 0,63614346 | 0,27494562 | 0,02068383 | 0,112816 | ARRDC5 |
| 1158,65594 | 0,63429258 | 0,15243537 | 3,17E-05 | 0,00089636 | DUSP1 |
| 908,761733 | 0,63336295 | 0,150605 | 2,61E-05 | 0,0007703 | TMEM129 |
| 97,3665087 | 0,63270443 | 0,29363664 | 0,03118384 | 0,14776594 | MAPK8IP2 |
| 149,700655 | 0,63139255 | 0,2074911 | 0,00234243 | 0,02509717 | ASB16 |
| 122,801499 | 0,631044 | 0,27369583 | 0,02113077 | 0,11446185 | WDR97 |
| 82,430076 | 0,63068438 | 0,22219613 | 0,00453385 | 0,04041674 | EEPD1 |
| 57,8952121 | 0,62895332 | 0,30150687 | 0,03697562 | 0,16564627 | WHRN |
| 725,340842 | 0,62878476 | 0,1228063 | 3,05E-07 | 1,87E-05 | PER1 |
| 387,426688 | 0,62878169 | 0,18477145 | 0,00066644 | 0,00988228 | JMJD4 |
| 449,440866 | 0,62811034 | 0,16713609 | 0,00017122 | 0,00349485 | S1PR4 |
| 82,0591569 | 0,62782525 | 0,2490508 | 0,01170643 | 0,07717681 | CYP2D7 |
| 1743,21135 | 0,62634293 | 0,113743 | 3,66E-08 | 2,91E-06 | LRFN4 |
| 73,9114578 | 0,62554324 | 0,25425036 | 0,01388041 | 0,08674109 | SPTBN5 |
| 191,157509 | 0,62542763 | 0,16857442 | 0,00020718 | 0,00403711 | ZNF784 |
| 145,778811 | 0,62479899 | 0,18297155 | 0,00063845 | 0,00955698 | CIRBP-AS1 |
| 169,115444 | 0,62385846 | 0,2175083 | 0,00412807 | 0,03793689 | MBP |
| 731,538927 | 0,62360331 | 0,13284093 | 2,67E-06 | 0,00011245 | IQSEC1 |
| 145,704568 | 0,62356498 | 0,23512106 | 0,00799924 | 0,05977634 | TAS1R3 |
| 48,7499204 | 0,62327976 | 0,29518381 | 0,0347296 | 0,15838005 | TCTE1 |
| 644,467736 | 0,62040475 | 0,18985852 | 0,00108417 | 0,01431513 | TTN |
| 306,368088 | 0,62039326 | 0,16863426 | 0,00023422 | 0,00445388 | COL18A1 |
| 490,217247 | 0,62034534 | 0,17587591 | 0,00042002 | 0,00700806 | CCDC78 |
| 82,6962338 | 0,61950726 | 0,28979067 | 0,03253501 | 0,15177389 | SSH3 |
| 46,1150675 | 0,61889314 | 0,29320505 | 0,0347907 | 0,15858225 | AIRN |
| 50,7568637 | 0,61770071 | 0,29541952 | 0,03653459 | 0,16427543 | ZFP42 |
| 78,1371902 | 0,61750669 | 0,28006061 | 0,02746085 | 0,13612452 | SNORD55 |
| 100,916884 | 0,61677285 | 0,21952739 | 0,0049611 | 0,04317804 | DYNLT4 |
| 53,6253624 | 0,61642313 | 0,30210892 | 0,04131047 | 0,17821146 | MIR3140 |
| 55,126829 | 0,61566317 | 0,29964508 | 0,03991366 | 0,17421138 | FHL1 |
| 249,58939 | 0,61421586 | 0,18598168 | 0,00095806 | 0,01306015 | ADAT3 |

| | | | | | |
|---|---|---|---|---|---|
| 168,566337 | 0,61320835 | 0,23769975 | 0,00988689 | 0,06892775 | GABARAPL1 |
| 274,963936 | 0,61265927 | 0,22429104 | 0,00630397 | 0,05110426 | RASA4B |
| 796,572521 | 0,61179852 | 0,1135769 | 7,18E-08 | 5,16E-06 | ULK1 |
| 264,994144 | 0,61087922 | 0,22408115 | 0,00640775 | 0,05161391 | SPRN |
| 721,183091 | 0,61004854 | 0,12753746 | 1,72E-06 | 7,94E-05 | PLK3 |
| 1092,50053 | 0,60984093 | 0,12905097 | 2,29E-06 | 0,00010097 | TRMT61A |
| 75,574503 | 0,60903592 | 0,31034814 | 0,04971266 | 0,20176036 | TMEM38A |
| 120,926058 | 0,60850133 | 0,22091876 | 0,00587977 | 0,04864498 | RALGPS2 |
| 49,0781939 | 0,60774049 | 0,30167302 | 0,04395005 | 0,18582767 | INSRR |
| 168,05547 | 0,60602085 | 0,16963098 | 0,00035348 | 0,00613028 | MSX1 |
| 221,829134 | 0,60587954 | 0,16999242 | 0,00036503 | 0,00627312 | SMAD7 |
| 116,015962 | 0,60518171 | 0,2997423 | 0,04348652 | 0,18460633 | KIF26A |
| 2059,75435 | 0,6044765 | 0,1001049 | 1,56E-09 | 1,86E-07 | ABHD17A |
| 168,789813 | 0,60405923 | 0,29050494 | 0,03758589 | 0,16756897 | TERT |
| 704,507743 | 0,60335501 | 0,12295132 | 9,24E-07 | 4,64E-05 | C8orf82 |
| 108,474574 | 0,60264035 | 0,21505694 | 0,00507497 | 0,04388643 | KANK3 |
| 97,3488221 | 0,60237084 | 0,2652598 | 0,02315478 | 0,12155878 | OVGP1 |
| 1194,16044 | 0,60208346 | 0,13059867 | 4,02E-06 | 0,00015992 | E4F1 |
| 232,439984 | 0,60196327 | 0,27856757 | 0,03070123 | 0,1463219 | TEX29 |
| 5347,25967 | 0,5990493 | 0,11492645 | 1,86E-07 | 1,21E-05 | TMEM259 |
| 361,929725 | 0,59904559 | 0,1903687 | 0,00165087 | 0,01965777 | NME3 |
| 125,13539 | 0,59707728 | 0,21406592 | 0,00528349 | 0,04508825 | FRG2 |
| 867,072467 | 0,59599621 | 0,10373314 | 9,17E-09 | 8,76E-07 | RNF166 |
| 554,463209 | 0,59591723 | 0,17745225 | 0,00078457 | 0,01124603 | TIGD5 |
| 80,0316232 | 0,59574482 | 0,25930378 | 0,02159149 | 0,11586293 | CCDC107 |
| 262,113255 | 0,59574407 | 0,15316486 | 0,00010043 | 0,00230043 | SLC66A2 |
| 339,22332 | 0,59529916 | 0,14700226 | 5,13E-05 | 0,0013262 | VPS37B |
| 123,288475 | 0,59493879 | 0,1919032 | 0,00193388 | 0,02193706 | DACH1 |
| 545,174715 | 0,5944646 | 0,16201172 | 0,00024324 | 0,00457499 | PEX16 |
| 132,658475 | 0,59314461 | 0,27526725 | 0,03117785 | 0,14776594 | FAM157B |
| 1464,85166 | 0,59289119 | 0,10841504 | 4,53E-08 | 3,50E-06 | TENT5C |
| 155,136682 | 0,59267744 | 0,21511615 | 0,00586651 | 0,04862032 | UAP1L1 |
| 459,62931 | 0,59224561 | 0,16045323 | 0,0002233 | 0,00429384 | ZNF668 |
| 57,6965567 | 0,58914438 | 0,28746613 | 0,04041918 | 0,17556733 | KCNG2 |
| 401,09768 | 0,58887847 | 0,13971646 | 2,50E-05 | 0,00074371 | TEDC2 |
| 252,780263 | 0,5852462 | 0,25271112 | 0,02056535 | 0,11236411 | BTBD19 |
| 110,977982 | 0,58477485 | 0,2843059 | 0,03970057 | 0,17383052 | NAV1 |
| 104,802839 | 0,58418221 | 0,21271181 | 0,00602615 | 0,04948969 | MESP1 |
| 1044,4724 | 0,58362241 | 0,26987966 | 0,03057747 | 0,1459822 | OPLAH |
| 93,9660215 | 0,58303393 | 0,26993897 | 0,0307825 | 0,14656155 | SNED1 |
| 188,151739 | 0,58285364 | 0,21069681 | 0,00566938 | 0,04748578 | ICAM3 |
| 110,221092 | 0,58203733 | 0,22964267 | 0,01125967 | 0,0752275 | VASN |
| 337,223758 | 0,58186752 | 0,14589187 | 6,65E-05 | 0,0016265 | ZNF707 |
| 1593,31504 | 0,58081485 | 0,14004271 | 3,36E-05 | 0,00093977 | PGGHG |
| 150,579176 | 0,5803779 | 0,191392 | 0,00242614 | 0,02577486 | PSD |
| 95,1947785 | 0,57933891 | 0,23487396 | 0,01364046 | 0,08561948 | SNORD2 |
| 769,272488 | 0,57876036 | 0,12908514 | 7,34E-06 | 0,00027129 | SGSH |
| 51,126701 | 0,57860253 | 0,27905263 | 0,03813013 | 0,16919799 | LOC105369669 |

| | | | | | |
|---|---|---|---|---|---|
| 1534,74897 | 0,57851315 | 0,12494667 | 3,66E-06 | 0,00014746 | KIFC2 |
| 309,875856 | 0,57820637 | 0,14903677 | 0,00010462 | 0,00237065 | BRI3BP |
| 2252,26979 | 0,57763521 | 0,25878831 | 0,02560993 | 0,12994515 | WDR90 |
| 484,972875 | 0,57760244 | 0,16140073 | 0,00034533 | 0,00602636 | RASA4 |
| 872,787048 | 0,57633317 | 0,15517905 | 0,00020402 | 0,00397962 | GPCPD1 |
| 85,0793411 | 0,57623902 | 0,22747759 | 0,01130367 | 0,07541491 | SLC2A4 |
| 126,336579 | 0,57613477 | 0,22016255 | 0,00887425 | 0,06411213 | EGFL7 |
| 105,742005 | 0,57599847 | 0,2271717 | 0,01122804 | 0,07509571 | NPEPL1 |
| 149,469955 | 0,57485966 | 0,17126447 | 0,00078918 | 0,01130365 | CDK5R1 |
| 194,768965 | 0,57475764 | 0,15888424 | 0,00029751 | 0,00535088 | LINC03124 |
| 71,2962808 | 0,57452694 | 0,25228143 | 0,02276679 | 0,11992371 | DNM1P41 |
| 310,31689 | 0,57379155 | 0,13434622 | 1,95E-05 | 0,00060067 | INAFM1 |
| 436,840375 | 0,57349431 | 0,16534522 | 0,00052344 | 0,008274 | EME2 |
| 756,775412 | 0,57321476 | 0,13325025 | 1,69E-05 | 0,00054058 | TMC6 |
| 464,858263 | 0,57282251 | 0,16595498 | 0,00055712 | 0,00869025 | RXRA |
| 88,7338331 | 0,57203754 | 0,26331108 | 0,02981962 | 0,14327504 | BBS5 |
| 76,2800811 | 0,57188115 | 0,26229676 | 0,0292365 | 0,14163095 | EFCAB2-AS1 |
| 229,700262 | 0,57115323 | 0,16319532 | 0,00046558 | 0,00759423 | MAP3K8 |
| 2904,48627 | 0,56827688 | 0,14165317 | 6,03E-05 | 0,00149656 | BRAT1 |
| 573,528044 | 0,56760939 | 0,16155472 | 0,00044238 | 0,00729117 | TMEM160 |
| 243,215128 | 0,56680758 | 0,18732671 | 0,00248001 | 0,02617078 | GDPD5 |
| 907,953404 | 0,56673098 | 0,12975986 | 1,26E-05 | 0,00042567 | KLHL21 |
| 2790,89297 | 0,56664771 | 0,10457892 | 6,01E-08 | 4,41E-06 | SIPA1 |
| 188,010395 | 0,56560858 | 0,16492129 | 0,00060455 | 0,00915088 | NAGS |
| 1075,95806 | 0,56319608 | 0,13971116 | 5,55E-05 | 0,00140592 | PIGQ |
| 109,045506 | 0,56291766 | 0,2091711 | 0,00711992 | 0,05541587 | EFNA4 |
| 95,3141814 | 0,56244556 | 0,23588015 | 0,01710445 | 0,09945292 | TSACC |
| 139,993443 | 0,56206507 | 0,20278998 | 0,00557711 | 0,0469415 | NOL4L |
| 88,1021757 | 0,56151069 | 0,23671216 | 0,01768614 | 0,10149814 | TSPAN33 |
| 226,57609 | 0,56055738 | 0,20975636 | 0,0075306 | 0,05752383 | UBASH3B |
| 1199,8378 | 0,55925686 | 0,13854383 | 5,42E-05 | 0,00138547 | CTSD |
| 87,6661068 | 0,55918684 | 0,23037817 | 0,01521351 | 0,09235352 | ABCB5 |
| 232,32962 | 0,55902679 | 0,15650262 | 0,00035427 | 0,0061328 | MILIP |
| 491,392223 | 0,55833594 | 0,1568755 | 0,00037212 | 0,00635467 | FOXO3 |
| 814,721664 | 0,55714007 | 0,09928534 | 2,01E-08 | 1,73E-06 | PMF1 |
| 245,00211 | 0,55515556 | 0,18331743 | 0,00245866 | 0,0260235 | CCDC85C |
| 242,703532 | 0,55412642 | 0,20316565 | 0,00638238 | 0,05149048 | OBSL1 |
| 281,96008 | 0,55375707 | 0,20650812 | 0,0073287 | 0,05652531 | HES4 |
| 100,818216 | 0,55340757 | 0,21877953 | 0,01142181 | 0,07593537 | ICA1L |
| 754,859497 | 0,55276028 | 0,16453959 | 0,00078102 | 0,01121214 | SAMD11 |
| 396,646868 | 0,5523647 | 0,1692109 | 0,00109713 | 0,01440583 | SNORA71C |
| 99,5227126 | 0,55205904 | 0,26971981 | 0,04067899 | 0,17625078 | NICOL1 |
| 1990,87721 | 0,551927 | 0,16593054 | 0,00088022 | 0,01228217 | ADAM8 |
| 96,3293462 | 0,55187034 | 0,24631067 | 0,02505551 | 0,12785203 | MMP24OS |
| 2210,11007 | 0,55174889 | 0,09628348 | 1,00E-08 | 9,34E-07 | IFRD2 |
| 479,177688 | 0,55173518 | 0,14337289 | 0,00011896 | 0,00262956 | BBC3 |
| 671,037677 | 0,55092777 | 0,12399251 | 8,86E-06 | 0,00031877 | WDR13 |
| 1121,44497 | 0,55080184 | 0,14115617 | 9,54E-05 | 0,00219314 | FRMD8 |

| | | | | | |
|---|---|---|---|---|---|
| 136,94495 | 0,5505726 | 0,20372595 | 0,00688169 | 0,05414073 | CRACR2B |
| 170,00088 | 0,55049856 | 0,18932108 | 0,00364038 | 0,03459551 | ACTL10 |
| 194,337729 | 0,54836527 | 0,19575657 | 0,00509033 | 0,04397907 | TMEM86B |
| 1502,57701 | 0,54690506 | 0,13963016 | 8,97E-05 | 0,00208312 | KDM4B |
| 123,364983 | 0,54682784 | 0,25511258 | 0,03207485 | 0,15051792 | PSD2 |
| 166,253577 | 0,54677336 | 0,17543935 | 0,00182953 | 0,02117218 | LTBP3 |
| 205,33416 | 0,54652355 | 0,17310295 | 0,00159283 | 0,01914733 | PITX1 |
| 225,789463 | 0,54461677 | 0,18157056 | 0,00270443 | 0,02807577 | ZDHHC24 |
| 5596,759 | 0,54458357 | 0,27595714 | 0,04844599 | 0,19859422 | HSPA1B |
| 1326,83897 | 0,54455306 | 0,13566356 | 5,97E-05 | 0,00148823 | FAAP100 |
| 342,847041 | 0,54396443 | 0,15548687 | 0,00046795 | 0,00761977 | REC8 |
| 503,767796 | 0,54386893 | 0,14275338 | 0,00013905 | 0,00293637 | SLC9A3 |
| 360,302918 | 0,54382626 | 0,17538235 | 0,00192996 | 0,02191884 | SLC66A1 |
| 158,248286 | 0,54381272 | 0,17456389 | 0,0018378 | 0,02125489 | ID1 |
| 123,475358 | 0,54245471 | 0,2495192 | 0,02970514 | 0,14305163 | LINC00937 |
| 171,345869 | 0,54194247 | 0,23162486 | 0,01929705 | 0,10757795 | PRXL2B |
| 4600,45485 | 0,5406705 | 0,15420308 | 0,00045451 | 0,00744575 | H3C8 |
| 3075,10884 | 0,54026398 | 0,09542358 | 1,50E-08 | 1,32E-06 | RELB |
| 544,793647 | 0,54013047 | 0,21088778 | 0,01043046 | 0,07124721 | TNFRSF14 |
| 275,088834 | 0,53894024 | 0,17062997 | 0,00158566 | 0,01908826 | TP53I13 |
| 2000,10142 | 0,53892143 | 0,15501027 | 0,00050766 | 0,00808537 | TCIRG1 |
| 2028,33818 | 0,53877066 | 0,12389406 | 1,37E-05 | 0,00045632 | RPUSD1 |
| 581,496223 | 0,53766283 | 0,14192772 | 0,00015169 | 0,00315399 | VCL |
| 263,450035 | 0,5376077 | 0,195152 | 0,00587253 | 0,04862295 | PIM2 |
| 291,617104 | 0,53742829 | 0,25171237 | 0,03275375 | 0,15230615 | DENND6B |
| 123,784868 | 0,53723244 | 0,20846323 | 0,00996306 | 0,06920374 | TGFBR3L |
| 288,404373 | 0,53709999 | 0,15003936 | 0,00034395 | 0,00601473 | PKN3 |
| 2147,62734 | 0,53691145 | 0,12191695 | 1,06E-05 | 0,00036725 | ARHGAP45 |
| 104,172271 | 0,53626653 | 0,21148308 | 0,01122094 | 0,07507473 | MGC16275 |
| 2289,93032 | 0,53463461 | 0,11578007 | 3,88E-06 | 0,00015556 | LIMD2 |
| 84,8358702 | 0,53331072 | 0,22493985 | 0,01774466 | 0,10174141 | SNORA70G |
| 594,282953 | 0,53265708 | 0,1385613 | 0,00012095 | 0,00265789 | MTHFR |
| 684,543289 | 0,53249799 | 0,15843459 | 0,00077662 | 0,0111575 | ESRRA |
| 133,77764 | 0,53218577 | 0,1915477 | 0,00546364 | 0,04623425 | LOC729683 |
| 1089,20155 | 0,53210362 | 0,13319849 | 6,47E-05 | 0,00158888 | FASTK |
| 287,998884 | 0,53100223 | 0,2117031 | 0,01213341 | 0,07924623 | PDZD7 |
| 183,708184 | 0,53039054 | 0,20138916 | 0,00844701 | 0,06202406 | FOXO4 |
| 266,138378 | 0,52901551 | 0,21778854 | 0,01513916 | 0,09210549 | CBARP |
| 1541,69722 | 0,52876713 | 0,16231197 | 0,00112311 | 0,0146352 | TELO2 |
| 1251,5508 | 0,52865568 | 0,13715472 | 0,00011599 | 0,00257288 | ATP5F1D |
| 1357,05518 | 0,52856287 | 0,1312525 | 5,65E-05 | 0,00142289 | CSRNP1 |
| 749,55817 | 0,52613214 | 0,13321811 | 7,83E-05 | 0,00187406 | MFSD3 |
| 115,38 | 0,52593035 | 0,22817723 | 0,02117103 | 0,11453188 | SNHG20 |
| 1595,66571 | 0,52590569 | 0,11101824 | 2,17E-06 | 9,65E-05 | SUN2 |
| 240,884652 | 0,5255815 | 0,19638609 | 0,00744474 | 0,05707537 | OSBP2 |
| 500,655226 | 0,52514852 | 0,17020945 | 0,00203342 | 0,02271336 | PELI1 |
| 1910,64993 | 0,5248343 | 0,1435237 | 0,00025541 | 0,00475175 | SLC52A2 |
| 1525,14116 | 0,52372765 | 0,09381275 | 2,37E-08 | 2,00E-06 | NLRC5 |

| | | | | | |
|---|---|---|---|---|---|
| 784,805408 | 0,52319603 | 0,16499585 | 0,00151934 | 0,01848704 | MIB2 |
| 84,6342249 | 0,52302578 | 0,2305522 | 0,02329396 | 0,12195493 | PLTP |
| 1461,05079 | 0,5227015 | 0,12006425 | 1,34E-05 | 0,00044707 | GATD1 |
| 3361,18367 | 0,52187543 | 0,11134665 | 2,77E-06 | 0,00011507 | MC1R |
| 667,570335 | 0,52187461 | 0,14027588 | 0,00019895 | 0,0039212 | CIAO3 |
| 515,133934 | 0,52090608 | 0,15110051 | 0,00056598 | 0,00876737 | TSPOAP1 |
| 1021,25302 | 0,5196627 | 0,10963839 | 2,14E-06 | 9,57E-05 | KATNB1 |
| 173,647135 | 0,51915432 | 0,2012731 | 0,00989857 | 0,06893294 | DGCR6 |
| 272,123293 | 0,51795775 | 0,22289781 | 0,02013914 | 0,11093238 | MCRIP1 |
| 173,575838 | 0,51754926 | 0,17661524 | 0,00338551 | 0,0327826 | RGS3 |
| 613,958831 | 0,51627169 | 0,18409668 | 0,0050418 | 0,04371948 | TSPO |
| 106,22934 | 0,51626739 | 0,23165291 | 0,02583889 | 0,1305119 | MYH11 |
| 766,551167 | 0,51605855 | 0,16005296 | 0,00126282 | 0,01602531 | ALKBH7 |
| 167,720752 | 0,51600644 | 0,21354778 | 0,01567693 | 0,09425696 | ERMN |
| 404,505177 | 0,5151624 | 0,15840327 | 0,00114507 | 0,01483964 | HLA-H |
| 90,2907066 | 0,51371439 | 0,25193194 | 0,04144006 | 0,178445 | NTN3 |
| 528,47216 | 0,51358234 | 0,16250316 | 0,00157528 | 0,01899675 | KLHL24 |
| 122,603246 | 0,51311594 | 0,2301347 | 0,02577182 | 0,13031882 | JAZF1 |
| 161,331919 | 0,51240681 | 0,21283713 | 0,01606187 | 0,09568048 | PPM1M |
| 1795,55152 | 0,51199853 | 0,15041785 | 0,00066445 | 0,00987597 | AP5Z1 |
| 313,478871 | 0,51185089 | 0,14369543 | 0,00036796 | 0,00629499 | PCYT2 |
| 326,91448 | 0,51176347 | 0,16297601 | 0,00168881 | 0,01997127 | ZNF516 |
| 356,214822 | 0,50968101 | 0,12039139 | 2,30E-05 | 0,00069334 | IFFO1 |
| 5105,67676 | 0,50893007 | 0,1618569 | 0,00166473 | 0,01978536 | PKD1 |
| 380,68318 | 0,50851147 | 0,13967246 | 0,00027185 | 0,00500361 | PPP2R3B |
| 638,980077 | 0,50835194 | 0,09991923 | 3,63E-07 | 2,14E-05 | PLEKHO2 |
| 150,128513 | 0,50742695 | 0,18791219 | 0,00692685 | 0,05441508 | ADM2 |
| 357,461037 | 0,50733248 | 0,2206561 | 0,02149359 | 0,11553416 | AMH |
| 590,796421 | 0,50721502 | 0,13647297 | 0,00020192 | 0,00395099 | MSRB1 |
| 135,016968 | 0,50693047 | 0,18676147 | 0,00664119 | 0,05284186 | DUSP5 |
| 379,860679 | 0,50513102 | 0,17930563 | 0,00484517 | 0,0425606 | NECTIN2 |
| 114,282596 | 0,50477072 | 0,22930175 | 0,02771208 | 0,13665373 | UNC5CL |
| 75,5146644 | 0,50474836 | 0,24416743 | 0,03871321 | 0,17085056 | ID3 |
| 1005,16222 | 0,50457073 | 0,12991765 | 0,00010285 | 0,00233892 | MIIP |
| 665,342646 | 0,5038587 | 0,16147767 | 0,00180667 | 0,02098468 | ARSA |
| 161,975678 | 0,50375865 | 0,20707021 | 0,01498292 | 0,09140321 | ZDHHC11B |
| 219,762461 | 0,50365807 | 0,18797271 | 0,00737495 | 0,05670113 | TOR4A |
| 195,574667 | 0,50364681 | 0,21963935 | 0,02184439 | 0,11663667 | TP53INP1 |
| 554,383191 | 0,50151941 | 0,11094054 | 6,17E-06 | 0,00023148 | TSPAN14 |
| 3634,84356 | 0,50113477 | 0,12243699 | 4,26E-05 | 0,0011332 | CHTF18 |
| 1704,12866 | 0,50076613 | 0,12350145 | 5,02E-05 | 0,00129911 | TBL3 |
| 1292,50979 | 0,50031045 | 0,16042903 | 0,00181726 | 0,02106886 | RNPEPL1 |
| 151,121986 | 0,49984342 | 0,19510258 | 0,01040858 | 0,07112339 | RIPOR3 |
| 178,195248 | 0,4997728 | 0,22252683 | 0,02471048 | 0,12653605 | HCLS1 |
| 663,827829 | 0,49970068 | 0,1510309 | 0,00093764 | 0,01286519 | MARCHF2 |
| 95,5954813 | 0,49790195 | 0,22164297 | 0,02467748 | 0,12653605 | LOC100505502 |
| 124,14293 | 0,49781045 | 0,20136064 | 0,01342719 | 0,08465708 | STON2 |
| 192,793994 | 0,49752814 | 0,22208612 | 0,02507478 | 0,12787887 | KREMEN2 |

| | | | | | |
|---|---|---|---|---|---|
| 293,103821 | 0,49742104 | 0,18145341 | 0,00611937 | 0,05007985 | GLYCTK |
| 1151,95439 | 0,49713486 | 0,15906203 | 0,00177554 | 0,02072489 | LY6E |
| 366,450249 | 0,49646286 | 0,17473503 | 0,00449395 | 0,04020977 | DUSP2 |
| 249,588706 | 0,49585514 | 0,18787148 | 0,00830697 | 0,06132505 | UBTD1 |
| 1259,96645 | 0,49550338 | 0,14315553 | 0,00053759 | 0,0084342 | CLCN7 |
| 253,961902 | 0,49383299 | 0,23339705 | 0,03435763 | 0,15732833 | KCNJ11 |
| 90,4381277 | 0,49380531 | 0,23643801 | 0,03675103 | 0,16493508 | SRGAP2-AS1 |
| 113,648646 | 0,49366941 | 0,22013667 | 0,02492528 | 0,12731904 | EGR4 |
| 2702,59991 | 0,49335667 | 0,1189809 | 3,38E-05 | 0,00094205 | RHOT2 |
| 1041,175 | 0,49187379 | 0,11096117 | 9,30E-06 | 0,00033013 | TRIM11 |
| 138,936034 | 0,49117053 | 0,19019641 | 0,00981048 | 0,06859635 | PLPP1 |
| 126,092771 | 0,49109127 | 0,19880276 | 0,0135021 | 0,08495951 | PLA2G6 |
| 511,524463 | 0,4906889 | 0,1101914 | 8,47E-06 | 0,00030568 | PAQR4 |
| 456,257956 | 0,48995402 | 0,15379415 | 0,00144365 | 0,01778344 | ABCA7 |
| 1510,19641 | 0,48848073 | 0,0989105 | 7,87E-07 | 4,05E-05 | FPGS |
| 406,816241 | 0,48818584 | 0,17709849 | 0,00584096 | 0,04842983 | FBXL15 |
| 147,149237 | 0,48718 | 0,21870123 | 0,02590685 | 0,13065893 | BORCS6 |
| 181,979041 | 0,48717233 | 0,23006529 | 0,03421409 | 0,15690234 | ARVCF |
| 245,602724 | 0,48646845 | 0,18923442 | 0,01014877 | 0,06997994 | ZBTB20 |
| 1102,4204 | 0,48571083 | 0,12534689 | 0,00010665 | 0,00239952 | MPST |
| 849,074019 | 0,48507228 | 0,12035636 | 5,57E-05 | 0,00140722 | ZFTRAF1 |
| 760,045319 | 0,48491948 | 0,16220133 | 0,00279329 | 0,02869269 | SLC12A9 |
| 2492,90564 | 0,48448143 | 0,11249499 | 1,66E-05 | 0,00052964 | MBD3 |
| 855,669152 | 0,48402132 | 0,16174282 | 0,00276669 | 0,02854342 | UNC13D |
| 168,563848 | 0,48393974 | 0,1856431 | 0,00913851 | 0,06534642 | HIC1 |
| 424,346711 | 0,48388465 | 0,13524773 | 0,00034654 | 0,00603772 | NOS3 |
| 91,5373197 | 0,48286721 | 0,21502183 | 0,02472541 | 0,12657723 | DGCR11 |
| 2504,01521 | 0,48269822 | 0,11976361 | 5,57E-05 | 0,00140722 | NFKBIA |
| 298,483224 | 0,48204487 | 0,15012825 | 0,00132326 | 0,01659843 | CEBPD |
| 131,831331 | 0,48123301 | 0,20702256 | 0,02009637 | 0,1108232 | VIM-AS1 |
| 87,8316836 | 0,48102833 | 0,21951244 | 0,02842656 | 0,1392646 | EFNA1 |
| 109,679502 | 0,48049213 | 0,21570634 | 0,02591194 | 0,13065893 | TULP1 |
| 307,535486 | 0,48005662 | 0,13473989 | 0,00036686 | 0,00628754 | TBCC |
| 639,38502 | 0,48005231 | 0,15214206 | 0,00160339 | 0,01925789 | RFNG |
| 6453,26386 | 0,47967352 | 0,13083596 | 0,00024616 | 0,00462065 | ZBTB46 |
| 371,134282 | 0,47834945 | 0,16332252 | 0,00340203 | 0,03285787 | SLC6A8 |
| 3736,8769 | 0,47813585 | 0,12843467 | 0,00019703 | 0,00389144 | GNB2 |
| 658,697402 | 0,47795293 | 0,14000266 | 0,00064044 | 0,00957921 | IBA57 |
| 307,904254 | 0,47787665 | 0,14978465 | 0,00142064 | 0,01754563 | RASGRP1 |
| 176,206366 | 0,47723992 | 0,21578972 | 0,02699451 | 0,13458844 | MIR21 |
| 1618,02401 | 0,47700138 | 0,10812648 | 1,03E-05 | 0,00035769 | ZBED10P |
| 132,011433 | 0,47682776 | 0,23653232 | 0,04380943 | 0,18548183 | PLEKHF1 |
| 1120,77943 | 0,47668784 | 0,1390207 | 0,00060604 | 0,00916612 | ILVBL |
| 229,833629 | 0,47546975 | 0,18228846 | 0,00909833 | 0,0651588 | GNB1L |
| 163,40578 | 0,47534005 | 0,22105059 | 0,03152614 | 0,14875466 | ABCD1 |
| 564,578519 | 0,47506219 | 0,13586148 | 0,00047111 | 0,00764486 | TPRN |
| 128,091096 | 0,47481799 | 0,1920793 | 0,01343635 | 0,08466458 | TNS2 |
| 211,820041 | 0,47474193 | 0,16627162 | 0,00430071 | 0,03899079 | TREX1 |

| | | | | | |
|---|---|---|---|---|---|
| 417,609095 | 0,47428048 | 0,13981813 | 0,00069353 | 0,01019602 | CHMP6 |
| 1124,40603 | 0,47406309 | 0,11023296 | 1,70E-05 | 0,00054164 | TLNRD1 |
| 113,91206 | 0,47299263 | 0,20369499 | 0,02022953 | 0,11128894 | ITGB7 |
| 234,448799 | 0,47188812 | 0,15797301 | 0,00281597 | 0,02888019 | ATOSB |
| 1413,9713 | 0,47137753 | 0,09208016 | 3,07E-07 | 1,87E-05 | ZNF787 |
| 1206,3901 | 0,47063195 | 0,11365881 | 3,46E-05 | 0,00095902 | DGCR2 |
| 110,798776 | 0,47033834 | 0,21339612 | 0,02751996 | 0,13614715 | CTRL |
| 1677,56124 | 0,47009468 | 0,11095454 | 2,27E-05 | 0,00068742 | PGAP6 |
| 860,760727 | 0,46882624 | 0,11601297 | 5,32E-05 | 0,00136367 | H1-10 |
| 294,503758 | 0,46821065 | 0,19374566 | 0,01566514 | 0,09425428 | HSF4 |
| 769,026118 | 0,46701293 | 0,12634638 | 0,00021877 | 0,00421527 | MARCHF8 |
| 344,082375 | 0,46593447 | 0,15066614 | 0,00198481 | 0,02238055 | PSRC1 |
| 483,585033 | 0,46476482 | 0,13380109 | 0,00051362 | 0,00815288 | IRF5 |
| 1299,41043 | 0,46469597 | 0,08812869 | 1,34E-07 | 8,95E-06 | PTGES2 |
| 1610,79894 | 0,46439159 | 0,17263397 | 0,00714443 | 0,05556803 | MROH1 |
| 4175,50318 | 0,46403675 | 0,11944118 | 0,0001023 | 0,00233213 | HLA-A |
| 6160,91282 | 0,46344777 | 0,23514889 | 0,0487388 | 0,19934865 | HSPA1A |
| 1220,74465 | 0,46324386 | 0,17641387 | 0,00864221 | 0,06299274 | SLC49A3 |
| 439,361724 | 0,46312821 | 0,17053851 | 0,00661396 | 0,05264734 | SPIRE2 |
| 1036,6385 | 0,46177338 | 0,15485811 | 0,00286454 | 0,0292656 | ACAP3 |
| 1587,63452 | 0,46025739 | 0,13608964 | 0,00071956 | 0,01050523 | PPP1R16A |
| 836,289905 | 0,4601688 | 0,12251408 | 0,00017262 | 0,00351193 | CCDC85B |
| 125,878895 | 0,459669 | 0,21580301 | 0,03316833 | 0,15370444 | RUFY1-AS1 |
| 120,005445 | 0,45936467 | 0,19952671 | 0,02131986 | 0,11509077 | ZNF517 |
| 691,836059 | 0,45904841 | 0,1451301 | 0,00156145 | 0,01884199 | REEP2 |
| 1116,04681 | 0,45868603 | 0,14619255 | 0,00170368 | 0,02007313 | PPP1R15A |
| 128,768218 | 0,45819435 | 0,18973474 | 0,01573863 | 0,09448647 | CASZ1 |
| 2112,28223 | 0,45739386 | 0,10045303 | 5,28E-06 | 0,00020186 | TRABD |
| 176,498588 | 0,45701794 | 0,20644433 | 0,02684538 | 0,13402149 | BAD |
| 563,672112 | 0,45690464 | 0,18500736 | 0,01352428 | 0,08504252 | PGAM2 |
| 1364,82604 | 0,45672387 | 0,18579272 | 0,01396182 | 0,08704173 | RHPN1 |
| 2542,24836 | 0,45645468 | 0,11747664 | 0,00010212 | 0,00233083 | PTMS |
| 115,246812 | 0,45641357 | 0,202109 | 0,02392979 | 0,12397567 | ZNF611 |
| 668,047493 | 0,45574621 | 0,14490072 | 0,00165956 | 0,01974875 | PYCR3 |
| 323,848707 | 0,45552783 | 0,1623892 | 0,00502915 | 0,04364977 | F11R |
| 898,911322 | 0,45545566 | 0,12293503 | 0,00021152 | 0,00410895 | TSC22D4 |
| 2511,93263 | 0,45505435 | 0,12467552 | 0,00026233 | 0,00486151 | DVL1 |
| 242,185114 | 0,45484643 | 0,20248538 | 0,02468368 | 0,12653605 | ISG15 |
| 488,565651 | 0,45403877 | 0,17746936 | 0,01051531 | 0,07156625 | ECI1 |
| 308,760648 | 0,45317143 | 0,16807632 | 0,00701305 | 0,05478689 | MMD |
| 832,882218 | 0,45289113 | 0,11522289 | 8,47E-05 | 0,00199628 | SEMA6B |
| 345,905555 | 0,45280446 | 0,13008041 | 0,00049962 | 0,00799102 | TMEM140 |
| 363,983804 | 0,45273463 | 0,20561836 | 0,02767802 | 0,13655694 | SLC26A1 |
| 2866,73179 | 0,45256626 | 0,10985526 | 3,79E-05 | 0,00102855 | REPIN1 |
| 972,005121 | 0,45231607 | 0,11110429 | 4,68E-05 | 0,00122457 | DYNC2I2 |
| 791,071449 | 0,45213544 | 0,15855465 | 0,00434989 | 0,03928608 | SIVA1 |
| 421,52093 | 0,45212743 | 0,12084576 | 0,00018303 | 0,00368803 | STIM1 |
| 409,938868 | 0,45212442 | 0,1963424 | 0,02129379 | 0,1149828 | IDUA |

| | | | | | |
|---|---|---|---|---|---|
| 1196,90702 | 0,45200521 | 0,13830069 | 0,00108204 | 0,01429694 | PKMYT1 |
| 1145,13504 | 0,45173121 | 0,13714797 | 0,0009886 | 0,01337051 | KLHL17 |
| 508,576678 | 0,45168944 | 0,15450009 | 0,0034606 | 0,03323753 | NFKBIE |
| 1227,66911 | 0,45117104 | 0,14771776 | 0,00225603 | 0,02447614 | LLGL1 |
| 178,434826 | 0,4503765 | 0,1779231 | 0,01136411 | 0,07562245 | GARIN1A |
| 173,499642 | 0,45009058 | 0,21103306 | 0,03294142 | 0,15284076 | TPGS1 |
| 339,822179 | 0,44992434 | 0,16292261 | 0,00575219 | 0,04793127 | SNX18 |
| 5099,48786 | 0,44988924 | 0,1359019 | 0,00093164 | 0,01282001 | CAPN15 |
| 2435,99099 | 0,44978165 | 0,14200067 | 0,00153776 | 0,01863931 | SAPCD2 |
| 1856,34521 | 0,44960733 | 0,11549316 | 9,90E-05 | 0,00227141 | AMPD2 |
| 295,652583 | 0,44945274 | 0,16757906 | 0,0073176 | 0,05646671 | LAT |
| 346,680268 | 0,44924518 | 0,18724383 | 0,01642859 | 0,09701793 | MPP1 |
| 328,214592 | 0,44923326 | 0,15426193 | 0,00358955 | 0,03426732 | AVPR2 |
| 259,097359 | 0,44921066 | 0,13251956 | 0,00069952 | 0,01026807 | NKX2-5 |
| 604,006029 | 0,44895669 | 0,12860426 | 0,00048123 | 0,00777566 | LMBR1L |
| 226,71416 | 0,44864632 | 0,15882407 | 0,00473101 | 0,04188833 | CHDH |
| 1083,56704 | 0,44862122 | 0,13511931 | 0,00089958 | 0,01242399 | TNK2 |
| 2325,36894 | 0,44834944 | 0,09813204 | 4,90E-06 | 0,00019015 | CDCA4 |
| 381,493598 | 0,44785359 | 0,18968115 | 0,01822149 | 0,10386056 | MAMDC4 |
| 325,544645 | 0,44768936 | 0,21960454 | 0,04148835 | 0,17857168 | MXRA8 |
| 711,491777 | 0,44735079 | 0,11583473 | 0,00011247 | 0,00251849 | SELENOO |
| 418,224063 | 0,44630286 | 0,16878027 | 0,00818641 | 0,06076697 | NUDT14 |
| 476,899118 | 0,44541278 | 0,13921962 | 0,00137736 | 0,0171229 | TAF6L |
| 1123,9926 | 0,44471748 | 0,10844685 | 4,12E-05 | 0,00110349 | SPNS1 |
| 147,789518 | 0,4446996 | 0,20843876 | 0,0328855 | 0,1527686 | MX2 |
| 477,988163 | 0,44447053 | 0,19290363 | 0,02121666 | 0,11469726 | NAPRT |
| 338,926776 | 0,44416892 | 0,18180332 | 0,01456054 | 0,08956437 | DEXI |
| 299,629851 | 0,44412863 | 0,17620039 | 0,01171588 | 0,07721221 | BCL7C |
| 463,457398 | 0,44405194 | 0,16265631 | 0,00633341 | 0,05123318 | PLEKHM1P1 |
| 365,855541 | 0,44402357 | 0,17758843 | 0,01240897 | 0,08035256 | ZNF579 |
| 1668,39096 | 0,44378555 | 0,11035544 | 5,78E-05 | 0,00144973 | LMF2 |
| 1240,72267 | 0,44358442 | 0,10343288 | 1,80E-05 | 0,00056507 | DGAT1 |
| 1760,40372 | 0,44343282 | 0,12853646 | 0,00056088 | 0,00871378 | FBXW5 |
| 1714,95662 | 0,4434322 | 0,12919245 | 0,0005984 | 0,00909425 | VPS51 |
| 135,870512 | 0,44325935 | 0,22114456 | 0,04502859 | 0,18907809 | SNORA70 |
| 122,912936 | 0,44284815 | 0,21427422 | 0,03875908 | 0,17094651 | C1orf74 |
| 258,060128 | 0,44280599 | 0,19767989 | 0,02508993 | 0,12788968 | ELMO3 |
| 643,367142 | 0,44252439 | 0,17335766 | 0,01069016 | 0,07236799 | ZNF581 |
| 555,013047 | 0,44235459 | 0,13315944 | 0,00089377 | 0,01239242 | UBXN11 |
| 368,308146 | 0,44224052 | 0,15223723 | 0,0036732 | 0,03478507 | NATD1 |
| 587,731007 | 0,44107296 | 0,11858622 | 0,00019967 | 0,00392727 | NAGLU |
| 374,592488 | 0,44094493 | 0,13399641 | 0,00099931 | 0,01345757 | MMP25 |
| 519,146028 | 0,44071401 | 0,15673784 | 0,00492665 | 0,04295724 | ARMC5 |
| 554,339976 | 0,44057466 | 0,13533075 | 0,00113177 | 0,01470757 | TBC1D17 |
| 2735,36047 | 0,44045649 | 0,12599603 | 0,00047265 | 0,00766323 | ARFGAP1 |
| 3273,73007 | 0,43997487 | 0,14397196 | 0,00224329 | 0,02440855 | PSAP |
| 3548,37106 | 0,43995185 | 0,12972834 | 0,00069554 | 0,01021761 | DPP7 |
| 742,252474 | 0,43975087 | 0,1761008 | 0,01251945 | 0,08080714 | SERTAD1 |

| | | | | | |
|---|---|---|---|---|---|
| 302,240406 | 0,43904548 | 0,1345374 | 0,00110095 | 0,01443598 | CLUHP3 |
| 371,616494 | 0,43897282 | 0,19684685 | 0,02574598 | 0,13028608 | RNF215 |
| 178,631961 | 0,43888317 | 0,18635627 | 0,01851892 | 0,10481499 | LOC100128361 |
| 2042,93734 | 0,438443 | 0,16299867 | 0,00714828 | 0,05556803 | MFSD10 |
| 3215,66214 | 0,43780611 | 0,10550059 | 3,33E-05 | 0,00093279 | SH3BP2 |
| 1090,4434 | 0,43613024 | 0,1167561 | 0,00018742 | 0,00375654 | RGS14 |
| 1802,21582 | 0,43572221 | 0,12165209 | 0,00034136 | 0,00597483 | PPM1F |
| 159,941526 | 0,43534186 | 0,19746158 | 0,02747575 | 0,13612452 | LOC100288123 |
| 9625,64003 | 0,43525897 | 0,15007258 | 0,00372778 | 0,03512617 | PKD1P1 |
| 167,284197 | 0,43388259 | 0,21599023 | 0,04455765 | 0,18780917 | AQP11 |
| 932,756634 | 0,43365129 | 0,16496728 | 0,00857092 | 0,06266579 | MAFK |
| 1469,42525 | 0,43303122 | 0,13234286 | 0,00106774 | 0,01418554 | HDAC7 |
| 768,972849 | 0,43284489 | 0,14088683 | 0,00212425 | 0,02338155 | HDAC5 |
| 346,626358 | 0,4325434 | 0,16812153 | 0,01008789 | 0,06981454 | DGKA |
| 311,242647 | 0,43220136 | 0,19316729 | 0,02525721 | 0,12863841 | TNFRSF9 |
| 1020,76586 | 0,43210344 | 0,11573276 | 0,00018874 | 0,00377498 | CDC34 |
| 713,868776 | 0,43209972 | 0,13487598 | 0,00135682 | 0,01692317 | ASPSCR1 |
| 628,017534 | 0,43181699 | 0,14257341 | 0,00245581 | 0,02601696 | ATG4D |
| 1182,4644 | 0,43156142 | 0,12386584 | 0,00049379 | 0,00793795 | CNPPD1 |
| 2017,93172 | 0,4310511 | 0,11792423 | 0,00025686 | 0,00477409 | ULK3 |
| 1561,23222 | 0,43094162 | 0,12949926 | 0,00087549 | 0,0122433 | NADSYN1 |
| 1278,75127 | 0,43068273 | 0,12813864 | 0,00077642 | 0,0111575 | NUBP2 |
| 215,065061 | 0,43058301 | 0,1669431 | 0,00990237 | 0,06893402 | TMEM239 |
| 921,98979 | 0,43021348 | 0,12465384 | 0,00055796 | 0,00869343 | ZC3H12A |
| 330,964517 | 0,42990503 | 0,16213556 | 0,00801313 | 0,05983288 | ZBTB42 |
| 553,064217 | 0,42983072 | 0,18618439 | 0,02096416 | 0,1138249 | TEPSIN |
| 1689,85472 | 0,42962292 | 0,10624846 | 5,26E-05 | 0,00135336 | MCRS1 |
| 316,166334 | 0,42911183 | 0,17365727 | 0,01347256 | 0,08483006 | ZBTB10 |
| 250,566523 | 0,42906806 | 0,13535425 | 0,00152458 | 0,01851513 | MFSD14CP |
| 2121,85822 | 0,42846152 | 0,0783584 | 4,55E-08 | 3,50E-06 | GRK6 |
| 471,737953 | 0,42841984 | 0,15292975 | 0,00508789 | 0,04397806 | ABCA1 |
| 374,080498 | 0,42818799 | 0,16608756 | 0,00993493 | 0,06903373 | NUDT16L1 |
| 376,34258 | 0,42811181 | 0,17064602 | 0,01211519 | 0,07917823 | LINC01993 |
| 1883,39071 | 0,42758978 | 0,13924008 | 0,00213428 | 0,02346466 | NDOR1 |
| 6053,13646 | 0,42682894 | 0,08122127 | 1,48E-07 | 9,79E-06 | MAZ |
| 580,68715 | 0,42676555 | 0,12696264 | 0,00077563 | 0,0111575 | ZNF865 |
| 342,103355 | 0,42598749 | 0,16950997 | 0,01196915 | 0,07849851 | SFT2D3 |
| 201,951114 | 0,42591681 | 0,15943371 | 0,00755277 | 0,05758848 | ADGRB2 |
| 482,541642 | 0,42576883 | 0,15403544 | 0,00570806 | 0,04772526 | LINC00674 |
| 722,900543 | 0,4256906 | 0,16165144 | 0,00845383 | 0,06202406 | KRBA1 |
| 373,602038 | 0,42527025 | 0,14909523 | 0,00433987 | 0,03923299 | PKP3 |
| 231,304963 | 0,42521052 | 0,18315571 | 0,02025559 | 0,11131718 | CTBP1-AS |
| 1104,41478 | 0,4251631 | 0,13724368 | 0,00194917 | 0,02207074 | ENGASE |
| 2896,08399 | 0,42502896 | 0,11261382 | 0,00016051 | 0,00330869 | TRAF1 |
| 9067,0812 | 0,42498756 | 0,09802822 | 1,46E-05 | 0,0004731 | LENG8 |
| 145,713399 | 0,42496567 | 0,19634542 | 0,03043536 | 0,14549457 | SNX21 |
| 220,238562 | 0,42493178 | 0,17687909 | 0,01628853 | 0,09646173 | BRF2 |
| 1481,79653 | 0,42428367 | 0,1237574 | 0,00060726 | 0,00917176 | MAP3K10 |

| | | | | | |
|---|---|---|---|---|---|
| 127,789286 | 0,42415489 | 0,19415401 | 0,0289159 | 0,14061109 | HLA-DOB |
| 6713,31226 | 0,42392525 | 0,10520654 | 5,59E-05 | 0,00141039 | TUBB3 |
| 991,765538 | 0,4238678 | 0,16041145 | 0,0082326 | 0,06099019 | SPPL2B |
| 255,955374 | 0,42372236 | 0,15570544 | 0,00650243 | 0,05208825 | SLC25A25 |
| 588,377075 | 0,42275428 | 0,13778123 | 0,0021528 | 0,02362707 | ARRDC2 |
| 657,734341 | 0,42269546 | 0,16808927 | 0,01191309 | 0,07826655 | PKD1P6-NPIPP1 |
| 187,123117 | 0,42269464 | 0,19313193 | 0,02862363 | 0,139909 | GSN |
| 1364,28712 | 0,42261224 | 0,15151105 | 0,00528195 | 0,04508825 | PARP10 |
| 176,296048 | 0,42250523 | 0,18181944 | 0,02013822 | 0,11093238 | ABCB6 |
| 2282,90967 | 0,42231985 | 0,09631062 | 1,16E-05 | 0,00039761 | IRF2BP1 |
| 635,637053 | 0,42220983 | 0,13866561 | 0,00232831 | 0,02502175 | CUEDC1 |
| 689,019075 | 0,42203455 | 0,15811558 | 0,0076043 | 0,05785321 | TCEA2 |
| 237,19664 | 0,4215828 | 0,18547218 | 0,02302471 | 0,12101402 | AP4M1 |
| 2081,09617 | 0,42122852 | 0,10085243 | 2,96E-05 | 0,00084984 | GPAA1 |
| 371,421978 | 0,42070449 | 0,14516029 | 0,00375304 | 0,03532895 | ZNF628 |
| 582,27566 | 0,42041688 | 0,10911175 | 0,00011664 | 0,00258119 | KCTD17 |
| 434,829931 | 0,42009912 | 0,14817994 | 0,0045817 | 0,04073794 | ARHGAP27P1 |
| 782,669306 | 0,41998977 | 0,12891893 | 0,00112287 | 0,0146352 | HIP1R |
| 1131,89398 | 0,4197878 | 0,1409353 | 0,00289582 | 0,02944967 | SRGN |
| 900,8446 | 0,41952673 | 0,12208376 | 0,00058953 | 0,00901743 | SLC39A3 |
| 616,974214 | 0,41925081 | 0,15639409 | 0,00734611 | 0,05652531 | NEIL1 |
| 895,66466 | 0,41814152 | 0,10549348 | 7,38E-05 | 0,00177895 | LRCH4 |
| 530,031784 | 0,41769167 | 0,11930312 | 0,00046335 | 0,00756838 | HYAL2 |
| 699,968621 | 0,41760382 | 0,13478065 | 0,00194571 | 0,02205802 | NFKBID |
| 1004,40187 | 0,41741653 | 0,11119778 | 0,00017416 | 0,00353439 | FBXO31 |
| 467,390181 | 0,41729054 | 0,1255935 | 0,00089199 | 0,01239159 | CRY2 |
| 989,489969 | 0,4169094 | 0,11222026 | 0,00020312 | 0,00396626 | TNFRSF12A |
| 2181,23827 | 0,41599492 | 0,10585809 | 8,50E-05 | 0,00199874 | PIM3 |
| 640,139533 | 0,41590926 | 0,13513823 | 0,00208634 | 0,02312575 | DEDD2 |
| 2450,67055 | 0,41530868 | 0,11098 | 0,00018242 | 0,00368365 | MKNK2 |
| 151,175745 | 0,41516731 | 0,19989222 | 0,03780564 | 0,16823201 | RHBDF1 |
| 2581,86122 | 0,41500046 | 0,11260274 | 0,00022823 | 0,00436636 | SYNGR2 |
| 157,614321 | 0,41480314 | 0,20141677 | 0,03945335 | 0,17323187 | TRPV1 |
| 915,383735 | 0,41415173 | 0,12026615 | 0,00057397 | 0,00881496 | FLCN |
| 1404,22486 | 0,41392486 | 0,0859715 | 1,47E-06 | 7,01E-05 | TBC1D10B |
| 223,165425 | 0,41391747 | 0,18762189 | 0,0273752 | 0,13584218 | TM4SF19-AS1 |
| 14740,271 | 0,41386539 | 0,13125458 | 0,00161519 | 0,01935463 | HS6ST1 |
| 2070,72141 | 0,41377708 | 0,10420999 | 7,17E-05 | 0,00173679 | ADGRE5 |
| 651,140481 | 0,41332658 | 0,13695648 | 0,00254498 | 0,02673717 | SLC44A2 |
| 833,153642 | 0,41295391 | 0,14943748 | 0,00572045 | 0,04778662 | TPRA1 |
| 205,509564 | 0,41295128 | 0,20026864 | 0,039209 | 0,172609 | CTU1 |
| 372,730768 | 0,41293238 | 0,13827448 | 0,00282353 | 0,02894047 | FLYWCH2 |
| 1942,12739 | 0,41282927 | 0,11419153 | 0,00030007 | 0,00537662 | MVD |
| 1959,02705 | 0,41267605 | 0,12738659 | 0,00119716 | 0,01535689 | TUBGCP6 |
| 269,591036 | 0,41223782 | 0,18029 | 0,02222349 | 0,1177515 | DNASE1L1 |
| 1072,23209 | 0,41195074 | 0,10307914 | 6,43E-05 | 0,00158001 | ZBED1 |
| 259,544204 | 0,41148285 | 0,1794499 | 0,02184664 | 0,11663667 | PELI3 |
| 693,624692 | 0,41144405 | 0,11284191 | 0,00026615 | 0,00492257 | EMC10 |

| | | | | | |
|---|---|---|---|---|---|
| 845,188915 | 0,41124327 | 0,10032612 | 4,15E-05 | 0,00111016 | GMEB2 |
| 1038,70355 | 0,41101266 | 0,11188612 | 0,00023926 | 0,00451348 | EHD1 |
| 2676,36608 | 0,41082408 | 0,13294567 | 0,00200045 | 0,02248994 | SLC19A1 |
| 819,694082 | 0,41060849 | 0,14230693 | 0,00390951 | 0,03633193 | SPSB3 |
| 3559,5322 | 0,41022584 | 0,16723416 | 0,01416693 | 0,08794373 | AGRN |
| 192,951778 | 0,41011973 | 0,16742452 | 0,0143023 | 0,08841893 | SYT11 |
| 144,217476 | 0,41009043 | 0,19370916 | 0,03425629 | 0,15701806 | DNMT3A |
| 424,006361 | 0,40960973 | 0,18714845 | 0,02861946 | 0,139909 | CFAP410 |
| 1167,04162 | 0,40939149 | 0,1065825 | 0,0001225 | 0,00267638 | SH3GLB2 |
| 831,092338 | 0,40930848 | 0,11816129 | 0,00053224 | 0,00838883 | SNHG12 |
| 1821,74035 | 0,40920748 | 0,11878455 | 0,00057115 | 0,00879266 | NACC2 |
| 1458,02549 | 0,40908012 | 0,12595521 | 0,00116292 | 0,01505034 | AP5B1 |
| 843,142263 | 0,4088562 | 0,15510496 | 0,00838905 | 0,06171429 | SLC39A13 |
| 2940,82384 | 0,40782652 | 0,14173118 | 0,00400884 | 0,0370187 | CEP170B |
| 525,52425 | 0,40714879 | 0,1432564 | 0,00448176 | 0,04013226 | ZFTA |
| 211,252821 | 0,40646373 | 0,18993506 | 0,03235362 | 0,15126336 | MORN1 |
| 1098,60622 | 0,40618513 | 0,14768701 | 0,00595388 | 0,0491507 | TRIB3 |
| 329,971829 | 0,40601396 | 0,14946288 | 0,00659788 | 0,05260788 | THTPA |
| 3556,19396 | 0,40598641 | 0,1103354 | 0,00023363 | 0,00445169 | PKN1 |
| 3808,25602 | 0,40597622 | 0,15321602 | 0,00805637 | 0,06003723 | KCNAB2 |
| 332,054005 | 0,40596268 | 0,13862057 | 0,00340504 | 0,03285787 | LRRC8E |
| 189,438113 | 0,40569329 | 0,19864936 | 0,04112593 | 0,17780389 | KCP |
| 464,114888 | 0,40568058 | 0,13279259 | 0,00225065 | 0,02444578 | VPS37C |
| 2120,78311 | 0,40563102 | 0,09973212 | 4,76E-05 | 0,00124175 | ZNRF1 |
| 649,641792 | 0,40559125 | 0,10558563 | 0,00012236 | 0,00267638 | UNC93B1 |
| 2145,80508 | 0,40550138 | 0,10345444 | 8,87E-05 | 0,00206605 | DGKZ |
| 117,577293 | 0,40506102 | 0,19542192 | 0,03819544 | 0,16936863 | LINC01311 |
| 1518,26491 | 0,40463095 | 0,11778291 | 0,00059169 | 0,00904308 | GSDMD |
| 1329,28559 | 0,40421091 | 0,1277924 | 0,00156137 | 0,01884199 | MAP3K11 |
| 1498,34957 | 0,40412572 | 0,12228881 | 0,00095084 | 0,0129804 | SH2B1 |
| 2166,54054 | 0,40315145 | 0,12313689 | 0,00106033 | 0,01412609 | BCAR1 |
| 279,637917 | 0,40240014 | 0,1545975 | 0,00924419 | 0,06580486 | WDR83 |
| 460,597626 | 0,40183065 | 0,14277894 | 0,00488752 | 0,04278277 | PI4KAP1 |
| 558,102226 | 0,40130297 | 0,15203152 | 0,0083003 | 0,06132371 | TSPAN9 |
| 180,401467 | 0,40094283 | 0,17778781 | 0,02412227 | 0,12463614 | THNSL2 |
| 1739,42738 | 0,40089224 | 0,13446765 | 0,00287001 | 0,02928988 | TNFRSF1B |
| 723,31238 | 0,40086979 | 0,13347421 | 0,00267025 | 0,02779086 | SEMA4D |
| 177,621646 | 0,40065421 | 0,16373209 | 0,01440465 | 0,08889189 | ZFPM1 |
| 304,552216 | 0,40051559 | 0,16029873 | 0,01246999 | 0,0805822 | LENG9 |
| 207,25246 | 0,40003802 | 0,18556478 | 0,03110036 | 0,14770327 | KCTD11 |
| 1521,04395 | 0,39995814 | 0,13400373 | 0,00283876 | 0,02908076 | ELL |
| 816,741447 | 0,39987473 | 0,11126663 | 0,00032584 | 0,00575108 | MID1IP1 |
| 332,656571 | 0,39977352 | 0,18628885 | 0,03187398 | 0,14998449 | MROH6 |
| 909,804664 | 0,39956723 | 0,16960788 | 0,0184814 | 0,10469228 | ITGA5 |
| 162,667423 | 0,39920106 | 0,19615694 | 0,04183975 | 0,17955317 | MIRLET7BHG |
| 596,554489 | 0,39883917 | 0,13585644 | 0,00332753 | 0,03237031 | ABHD16B |
| 771,39657 | 0,39847028 | 0,10765399 | 0,00021441 | 0,00415524 | ITPRIP |
| 235,22267 | 0,39838328 | 0,15769758 | 0,01152879 | 0,07648536 | SMO |

| | | | | | |
|---|---|---|---|---|---|
| 851,262153 | 0,39819987 | 0,11872032 | 0,00079623 | 0,01139603 | FIS1 |
| 3371,45347 | 0,39748706 | 0,10086947 | 8,13E-05 | 0,00193177 | C19orf48P |
| 1001,79286 | 0,3970858 | 0,10346394 | 0,00012408 | 0,0026955 | MON1B |
| 243,769535 | 0,39697753 | 0,16535296 | 0,01635978 | 0,09667187 | NKD2 |
| 273,876449 | 0,39663926 | 0,17875089 | 0,02649014 | 0,13264488 | ADCK5 |
| 542,632987 | 0,39626772 | 0,12562756 | 0,0016088 | 0,0193026 | PNPLA2 |
| 163,236485 | 0,39623091 | 0,19449144 | 0,04162334 | 0,17898331 | MMP24 |
| 1134,67568 | 0,39537868 | 0,13468513 | 0,00332927 | 0,03237058 | CAPN10 |
| 10220,3945 | 0,39524905 | 0,0992932 | 6,87E-05 | 0,00166733 | SBNO2 |
| 643,331212 | 0,39523258 | 0,14230298 | 0,00547954 | 0,04634702 | RAD9A |
| 805,802926 | 0,39494413 | 0,14142367 | 0,00522813 | 0,04486848 | IER3 |
| 411,896298 | 0,39451028 | 0,17240486 | 0,02212133 | 0,11747275 | LOC100130872 |
| 1393,56772 | 0,39439013 | 0,08241154 | 1,70E-06 | 7,88E-05 | RPS6KA4 |
| 698,269049 | 0,39427461 | 0,11926231 | 0,00094657 | 0,01295011 | ZBTB47 |
| 1962,50705 | 0,39421076 | 0,14387471 | 0,00614469 | 0,05024361 | CHRD |
| 983,362119 | 0,39396649 | 0,13707494 | 0,00405186 | 0,03734306 | RGL2 |
| 2461,87262 | 0,3939053 | 0,11434457 | 0,00057128 | 0,00879266 | SPHK1 |
| 447,515666 | 0,3934513 | 0,15147899 | 0,00939329 | 0,06659065 | FHIP1B |
| 4510,03526 | 0,39326693 | 0,08566515 | 4,42E-06 | 0,00017338 | IRAK1 |
| 579,187093 | 0,39310085 | 0,13261606 | 0,00303469 | 0,03030041 | ANKRD9 |
| 1193,84984 | 0,392921 | 0,17956354 | 0,02865502 | 0,13995394 | HDAC10 |
| 363,170993 | 0,39276763 | 0,12803005 | 0,00215657 | 0,02364247 | CYB561D1 |
| 1320,86986 | 0,39201252 | 0,12561701 | 0,00180424 | 0,02097529 | HSPBP1 |
| 1287,69904 | 0,39185848 | 0,11801367 | 0,00089872 | 0,0124213 | ORAI2 |
| 383,040938 | 0,39148619 | 0,13665152 | 0,00417206 | 0,03825137 | PCGF3-AS1 |
| 386,217944 | 0,39124762 | 0,11738843 | 0,00085936 | 0,01205333 | GAPDHS |
| 698,612843 | 0,39120716 | 0,15608353 | 0,01219689 | 0,07949617 | MIGA2 |
| 1228,8014 | 0,39101342 | 0,12881577 | 0,0024018 | 0,02555926 | ZFYVE28 |
| 664,95682 | 0,39057414 | 0,14708165 | 0,00791928 | 0,05939419 | USP20 |
| 1013,29188 | 0,39048001 | 0,12706445 | 0,00211857 | 0,02335985 | ATG16L2 |
| 1474,08049 | 0,38982802 | 0,1360706 | 0,00417147 | 0,03825137 | PTOV1 |
| 493,759078 | 0,38971136 | 0,15182822 | 0,01026444 | 0,07049489 | GRAMD4 |
| 4631,82964 | 0,38904804 | 0,13192895 | 0,00318886 | 0,03137616 | ARHGAP4 |
| 4446,2471 | 0,38898246 | 0,12999647 | 0,00276926 | 0,02855428 | SBF1 |
| 999,594454 | 0,38869529 | 0,11133963 | 0,00048108 | 0,00777566 | MAP2K3 |
| 1305,91586 | 0,38868347 | 0,1042761 | 0,00019343 | 0,00384443 | ATP6V0E2 |
| 485,480801 | 0,38857757 | 0,156308 | 0,01291975 | 0,08236341 | FCGRT |
| 672,103116 | 0,3885484 | 0,1381304 | 0,00490954 | 0,04285407 | THAP7 |
| 1104,53737 | 0,3881913 | 0,10061634 | 0,00011426 | 0,00254393 | CC2D1B |
| 1912,71566 | 0,3880465 | 0,10951275 | 0,00039502 | 0,00667927 | ZBTB32 |
| 519,794227 | 0,38793912 | 0,16329352 | 0,01751491 | 0,10076002 | ZBTB7B |
| 449,653681 | 0,38781711 | 0,1401163 | 0,00564321 | 0,04730846 | RHBDD3 |
| 1429,55394 | 0,38771172 | 0,10962838 | 0,00040531 | 0,00684029 | SLC7A5P2 |
| 514,219746 | 0,38758061 | 0,15477691 | 0,01227549 | 0,07974787 | FDPSP2 |
| 1077,71757 | 0,38710756 | 0,1352438 | 0,00420586 | 0,03845582 | TBC1D24 |
| 194,395084 | 0,38708705 | 0,17998256 | 0,03150051 | 0,14867079 | ALOX12 |
| 1708,51727 | 0,38696493 | 0,13609521 | 0,00446434 | 0,04003396 | APRT |
| 528,301454 | 0,38653342 | 0,11842105 | 0,00109828 | 0,01441091 | UNKL |

| | | | | | |
|---|---|---|---|---|---|
| 1610,88939 | 0,38609582 | 0,14049518 | 0,00599404 | 0,0493101 | KIFC3 |
| 675,272264 | 0,38607335 | 0,15454661 | 0,01248599 | 0,08065805 | CLDND2 |
| 1880,15499 | 0,38593306 | 0,10788597 | 0,00034725 | 0,0060445 | FBXL19 |
| 928,215622 | 0,38568928 | 0,12125983 | 0,00146927 | 0,01799362 | SLC30A1 |
| 803,243584 | 0,38537498 | 0,1174411 | 0,00103281 | 0,01383006 | ISOC2 |
| 1169,65563 | 0,38485235 | 0,11585963 | 0,00089467 | 0,01239242 | BAK1 |
| 1290,50673 | 0,38464357 | 0,11324885 | 0,00068267 | 0,01007552 | RBM38 |
| 231,047787 | 0,38464325 | 0,18452555 | 0,03711484 | 0,16605579 | FKBP1AP1 |
| 3927,92692 | 0,38456159 | 0,10956131 | 0,00044809 | 0,00736596 | CYBC1 |
| 1737,95316 | 0,38439771 | 0,09294513 | 3,54E-05 | 0,00097403 | R3HDM4 |
| 2953,50493 | 0,38431113 | 0,12746897 | 0,00257031 | 0,02695836 | MAN2C1 |
| 1655,22886 | 0,38429732 | 0,10828367 | 0,00038672 | 0,00656828 | NEURL4 |
| 3911,37494 | 0,38422019 | 0,08977511 | 1,87E-05 | 0,00057926 | FLOT2 |
| 1654,51874 | 0,38396488 | 0,13533608 | 0,00455223 | 0,04051399 | SREBF1 |
| 580,899555 | 0,38335035 | 0,13083712 | 0,00338982 | 0,03280756 | SNAPC2 |
| 1646,8084 | 0,38319337 | 0,11314467 | 0,00070724 | 0,01034937 | CHMP1A |
| 2151,38721 | 0,38309203 | 0,12464121 | 0,00211522 | 0,02335341 | CD151 |
| 412,225473 | 0,38276929 | 0,14066824 | 0,00650693 | 0,05210229 | TMEM141 |
| 175,820037 | 0,38230245 | 0,16212409 | 0,01836946 | 0,10435766 | MYLK |
| 485,12574 | 0,38178868 | 0,16069802 | 0,01751027 | 0,10076002 | THEM6 |
| 1028,46225 | 0,3815988 | 0,12497423 | 0,00226249 | 0,02453214 | ABCB8 |
| 1422,40225 | 0,38146098 | 0,11771759 | 0,00119331 | 0,0153252 | PPP1R37 |
| 380,840512 | 0,38143144 | 0,17917664 | 0,03327072 | 0,15398267 | SARM1 |
| 8596,03053 | 0,38127097 | 0,08085257 | 2,41E-06 | 0,00010409 | EEF1D |
| 1142,15503 | 0,38126442 | 0,10472878 | 0,00027212 | 0,00500361 | TSPAN17 |
| 816,944033 | 0,38120061 | 0,11830967 | 0,00127273 | 0,01611874 | B3GAT3 |
| 813,695381 | 0,38069133 | 0,13919566 | 0,00623922 | 0,05078605 | E2F1 |
| 2932,62546 | 0,38053407 | 0,11390233 | 0,00083513 | 0,01180099 | RECQL4 |
| 176,188861 | 0,38045109 | 0,16544371 | 0,02147201 | 0,11550604 | GNAZ |
| 2092,44997 | 0,38019043 | 0,10296556 | 0,00022214 | 0,00427588 | PLEKHM2 |
| 224,862866 | 0,37966563 | 0,16189726 | 0,01902186 | 0,1066408 | GNMT |
| 621,134066 | 0,37955322 | 0,1631916 | 0,02002851 | 0,11054825 | SLC27A1 |
| 208,357552 | 0,37934234 | 0,16479305 | 0,02133903 | 0,11509571 | ZNF524 |
| 274,200359 | 0,37929131 | 0,14227312 | 0,00767739 | 0,05816328 | SLC35A2 |
| 1250,55268 | 0,37921367 | 0,14619715 | 0,00949075 | 0,06708052 | WDR81 |
| 1088,75567 | 0,37912664 | 0,15916961 | 0,01722341 | 0,0997503 | PNRC1 |
| 1724,95807 | 0,37895705 | 0,15434463 | 0,01407806 | 0,08755589 | ZNF316 |
| 490,578034 | 0,37862949 | 0,13118868 | 0,00389993 | 0,03626072 | ZER1 |
| 440,193135 | 0,37848718 | 0,14389399 | 0,00853046 | 0,06245052 | HPDL |
| 501,67018 | 0,37841082 | 0,14532864 | 0,00921889 | 0,06574845 | CLDN15 |
| 1606,57153 | 0,37818685 | 0,10717393 | 0,00041757 | 0,00697961 | KLF13 |
| 1583,69232 | 0,37805234 | 0,12755672 | 0,00303868 | 0,03032425 | CD320 |
| 1669,36128 | 0,37792549 | 0,11822246 | 0,00138994 | 0,01722271 | BTG1 |
| 228,843809 | 0,37769502 | 0,14949544 | 0,01152168 | 0,07646499 | LSR |
| 505,177519 | 0,37744691 | 0,1303747 | 0,00379046 | 0,03552218 | NECAB3 |
| 1144,63497 | 0,37717199 | 0,11002423 | 0,00060786 | 0,00917176 | ARHGEF18 |
| 495,686665 | 0,37715284 | 0,14097132 | 0,0074643 | 0,05717898 | RGS12 |
| 1039,7483 | 0,37686185 | 0,15108867 | 0,01262026 | 0,08128719 | THEMIS2 |

| | | | | | |
|---|---|---|---|---|---|
| 242,686055 | 0,37676183 | 0,14399845 | 0,00888547 | 0,06416866 | TMEM177 |
| 543,413948 | 0,37662505 | 0,18679457 | 0,04377356 | 0,18541294 | TMEM9 |
| 1321,33045 | 0,37633648 | 0,1036088 | 0,00028092 | 0,00512079 | NLRP1 |
| 336,085926 | 0,37616037 | 0,16199889 | 0,02023328 | 0,11128894 | C1orf122 |
| 269,843073 | 0,37609174 | 0,17832383 | 0,0349413 | 0,15900057 | ENDOV |
| 1596,34551 | 0,37607386 | 0,11262316 | 0,00084013 | 0,01182747 | ARF5 |
| 1374,57539 | 0,37559493 | 0,13040597 | 0,00397426 | 0,0368252 | GPR132 |
| 1395,0429 | 0,37526567 | 0,0893743 | 2,68E-05 | 0,00078712 | SEC14L1 |
| 530,366979 | 0,3752641 | 0,12329214 | 0,00233689 | 0,02506596 | CPTP |
| 803,016157 | 0,37458566 | 0,1273879 | 0,0032767 | 0,03199096 | CCDC142 |
| 1109,46619 | 0,37439228 | 0,12023395 | 0,00184654 | 0,02130389 | SHARPIN |
| 563,739391 | 0,37437415 | 0,12895113 | 0,00369341 | 0,03491294 | MICAL1 |
| 593,957547 | 0,37391751 | 0,14581454 | 0,01033733 | 0,07091341 | SDHAF1 |
| 279,805357 | 0,37365526 | 0,16221159 | 0,02125043 | 0,11478142 | ADM5 |
| 304,812455 | 0,3736017 | 0,18363643 | 0,04190484 | 0,17970205 | RRN3P3 |
| 676,197722 | 0,37302677 | 0,11329435 | 0,00099285 | 0,01340878 | ZNF444 |
| 371,609512 | 0,3727755 | 0,17400857 | 0,03217077 | 0,15081841 | MYL5 |
| 735,357989 | 0,3726121 | 0,10916914 | 0,00064213 | 0,00958945 | TMEM222 |
| 790,09876 | 0,37258781 | 0,10272697 | 0,00028676 | 0,00518726 | CORO1B |
| 1454,6522 | 0,37255861 | 0,11193924 | 0,00087402 | 0,01223179 | QPCTL |
| 519,147345 | 0,37242815 | 0,16158328 | 0,02117396 | 0,11453188 | PUSL1 |
| 550,382976 | 0,37174972 | 0,11491143 | 0,00121601 | 0,01558817 | WHAMM |
| 204,144711 | 0,37169499 | 0,15203009 | 0,01449012 | 0,08923944 | ADAM11 |
| 896,018282 | 0,37133914 | 0,15121953 | 0,0140638 | 0,08750452 | UVSSA |
| 2932,72863 | 0,3710524 | 0,08053228 | 4,08E-06 | 0,00016166 | APH1A |
| 154,794135 | 0,37072362 | 0,18075791 | 0,04027279 | 0,17512995 | APOBEC3F |
| 702,809932 | 0,37070418 | 0,12459234 | 0,00292667 | 0,02958095 | FBXL14 |
| 1781,16215 | 0,3706338 | 0,09243695 | 6,08E-05 | 0,00150639 | CDT1 |
| 1073,63401 | 0,37008952 | 0,13149498 | 0,00488568 | 0,04278277 | MAN2B1 |
| 216,5052 | 0,36959696 | 0,18848259 | 0,04988979 | 0,20201384 | SNORA61 |
| 939,463754 | 0,36951099 | 0,11123926 | 0,00089449 | 0,01239242 | CTU2 |
| 507,211862 | 0,36918474 | 0,129358 | 0,00431757 | 0,03906871 | PEX10 |
| 317,035595 | 0,36894994 | 0,17032149 | 0,03029612 | 0,14490212 | LY6G5B |
| 913,768802 | 0,36894684 | 0,14963062 | 0,01367391 | 0,08578386 | FAAP20 |
| 674,799813 | 0,36738788 | 0,13155827 | 0,00522885 | 0,04486848 | TEDC1 |
| 352,502684 | 0,36716592 | 0,15938264 | 0,02124086 | 0,11478142 | SLC41A3 |
| 205,671707 | 0,36696531 | 0,15911413 | 0,02109374 | 0,11429399 | RCAN3 |
| 2567,01983 | 0,3669555 | 0,12417058 | 0,00312413 | 0,03086766 | GPX1 |
| 326,691735 | 0,36661471 | 0,14172053 | 0,0096848 | 0,06809588 | GPATCH3 |
| 555,338808 | 0,36656072 | 0,12162334 | 0,00257912 | 0,02702076 | ARRDC1 |
| 293,535055 | 0,36653644 | 0,15062988 | 0,0149595 | 0,09133549 | ZNHIT2 |
| 1345,43254 | 0,36646821 | 0,09901956 | 0,00021478 | 0,00415524 | B4GALT2 |
| 2480,73485 | 0,36612901 | 0,10575027 | 0,00053577 | 0,0084197 | MFNG |
| 1858,62903 | 0,36609555 | 0,10673118 | 0,00060342 | 0,0091412 | TRIP6 |
| 4450,48036 | 0,36558029 | 0,1043733 | 0,0004607 | 0,00753824 | POLD1 |
| 806,713047 | 0,36549769 | 0,13391691 | 0,00634714 | 0,05131358 | SH2D5 |
| 3191,37635 | 0,36544218 | 0,11233122 | 0,00114091 | 0,01480598 | ZNF598 |
| 1853,4751 | 0,36482289 | 0,12850241 | 0,00452501 | 0,0403667 | TAF1C |

| | | | | | |
|---|---|---|---|---|---|
| 617,105607 | 0,36452623 | 0,14216912 | 0,01034639 | 0,07092602 | RARA |
| 3164,42304 | 0,36443357 | 0,15899319 | 0,02189796 | 0,11677881 | QSOX1 |
| 568,146641 | 0,36433387 | 0,12466883 | 0,0034733 | 0,03334264 | GNA11 |
| 455,455121 | 0,36431695 | 0,11450947 | 0,00146492 | 0,01795193 | LZTS3 |
| 1213,01407 | 0,36397169 | 0,13574598 | 0,00733432 | 0,05652531 | TMEM250 |
| 407,372217 | 0,3637576 | 0,17207107 | 0,03451558 | 0,15770813 | BSN |
| 415,807202 | 0,36313245 | 0,17734222 | 0,04059561 | 0,17605585 | CDKN1B |
| 232,639236 | 0,36311458 | 0,1726988 | 0,03550172 | 0,16085444 | TANGO2 |
| 650,387141 | 0,36244956 | 0,12769776 | 0,00453489 | 0,04041674 | SLC45A3 |
| 1446,92694 | 0,36228039 | 0,10355073 | 0,00046774 | 0,00761977 | MEF2D |
| 558,252144 | 0,36214389 | 0,14705821 | 0,0137938 | 0,08636418 | YJEFN3 |
| 633,445471 | 0,36202892 | 0,1166711 | 0,00191578 | 0,02182104 | TPST2 |
| 1131,88263 | 0,36202305 | 0,1165813 | 0,00190069 | 0,02171674 | GATD3 |
| 3760,78521 | 0,36143434 | 0,09361525 | 0,00011299 | 0,00252415 | MTCH1 |
| 1776,31655 | 0,36128848 | 0,13872557 | 0,00920515 | 0,06567523 | CDK10 |
| 557,903959 | 0,36096681 | 0,13958132 | 0,00970777 | 0,06818142 | HERPUD1 |
| 1083,08344 | 0,36065922 | 0,13200583 | 0,00629232 | 0,05107552 | GMIP |
| 1754,59164 | 0,36056384 | 0,10443045 | 0,00055507 | 0,00866538 | RETREG2 |
| 1451,50434 | 0,35969587 | 0,11100563 | 0,00119388 | 0,0153252 | NBPF25P |
| 322,960484 | 0,35949892 | 0,14501279 | 0,01317202 | 0,08343748 | KREMEN1 |
| 2396,69792 | 0,35931858 | 0,14221819 | 0,01151954 | 0,07646499 | MAPK11 |
| 3034,56339 | 0,35907636 | 0,13438226 | 0,0075389 | 0,05754077 | FAM83H |
| 344,082656 | 0,35897294 | 0,12996258 | 0,00574264 | 0,04790854 | NAGPA |
| 718,090857 | 0,35845495 | 0,13784302 | 0,00930995 | 0,06611673 | DYRK1B |
| 832,190454 | 0,35830317 | 0,14823588 | 0,01564407 | 0,09417801 | HPS6 |
| 880,278714 | 0,35830313 | 0,12904397 | 0,00549312 | 0,04643319 | SNHG4 |
| 223,49347 | 0,35795157 | 0,17029365 | 0,0355562 | 0,16102415 | HDHD3 |
| 1377,39169 | 0,35791929 | 0,10178909 | 0,00043763 | 0,00722553 | SLC2A4RG |
| 1787,66191 | 0,35747444 | 0,14643967 | 0,01464245 | 0,0899469 | SLC17A9 |
| 590,179028 | 0,3571944 | 0,12288078 | 0,00365103 | 0,03464454 | CACTIN |
| 3224,73114 | 0,35670754 | 0,16096803 | 0,02669049 | 0,13345952 | HELZ2 |
| 3299,02092 | 0,35648494 | 0,11567075 | 0,00205688 | 0,0228662 | PCNX3 |
| 2814,23318 | 0,35560907 | 0,11582248 | 0,00213848 | 0,02348357 | FERMT3 |
| 2335,53844 | 0,35544384 | 0,12426037 | 0,00423005 | 0,03859963 | PFKL |
| 374,73688 | 0,35543656 | 0,14308732 | 0,01298963 | 0,08258627 | FHL3 |
| 546,641331 | 0,35536048 | 0,15468529 | 0,02160094 | 0,11588076 | TOM1 |
| 560,485812 | 0,35529109 | 0,15669736 | 0,02336755 | 0,12227028 | GPSM1 |
| 3767,54067 | 0,35524573 | 0,14091083 | 0,01169994 | 0,07716087 | SLC16A3 |
| 1071,18959 | 0,35520405 | 0,12366113 | 0,00407369 | 0,03750767 | ST3GAL2 |
| 416,218034 | 0,35495055 | 0,14277193 | 0,01291383 | 0,08235344 | ACBD4 |
| 2738,16662 | 0,35479162 | 0,09806625 | 0,00029703 | 0,00534738 | NSMF |
| 267,77024 | 0,35476955 | 0,15926739 | 0,02591327 | 0,13065893 | RPP25L |
| 1456,92683 | 0,3546863 | 0,0922211 | 0,00012004 | 0,00264411 | MFSD12 |
| 1105,31023 | 0,35455491 | 0,10192355 | 0,00050397 | 0,0080402 | SPC24 |
| 610,498533 | 0,35444242 | 0,13555507 | 0,00892932 | 0,06436251 | CLBA1 |
| 969,851067 | 0,35442583 | 0,13913041 | 0,01085177 | 0,07314486 | SNX11 |
| 7224,11302 | 0,35439378 | 0,11313379 | 0,00173313 | 0,02030498 | OAZ1 |
| 333,29254 | 0,3542815 | 0,15945183 | 0,02629198 | 0,13206016 | TNFSF9 |

| | | | | | |
|---|---|---|---|---|---|
| 602,774629 | 0,35417148 | 0,10912541 | 0,00117226 | 0,01514014 | MEX3D |
| 1669,78577 | 0,35395586 | 0,09470699 | 0,00018595 | 0,00373497 | AXIN1 |
| 1019,65853 | 0,35384224 | 0,14833833 | 0,01706181 | 0,09931121 | BNIP3L |
| 2166,31041 | 0,35358029 | 0,09745647 | 0,00028553 | 0,00517486 | NADK |
| 691,020571 | 0,35357902 | 0,14683705 | 0,01604156 | 0,09564473 | SH2B3 |
| 231,950456 | 0,35321442 | 0,17854477 | 0,04789531 | 0,19709161 | CHRNG |
| 234,619452 | 0,35313209 | 0,17172516 | 0,03974596 | 0,17388053 | SGK1 |
| 1542,54291 | 0,35301583 | 0,09660065 | 0,0002578 | 0,0047822 | MAFG |
| 714,877365 | 0,35236606 | 0,13446816 | 0,00878157 | 0,06368577 | C19orf25 |
| 840,351468 | 0,35228415 | 0,09869397 | 0,00035772 | 0,0061699 | CDR2L |
| 792,714974 | 0,3522566 | 0,13910045 | 0,01132879 | 0,07547605 | RNASEH2C |
| 664,508554 | 0,35190516 | 0,1235958 | 0,00441021 | 0,03969816 | ZNF48 |
| 544,018095 | 0,35189313 | 0,14222851 | 0,01335576 | 0,08429099 | ZNF324 |
| 1900,66245 | 0,35181021 | 0,11490363 | 0,00220021 | 0,02403594 | FAM193B |
| 1538,9702 | 0,35175568 | 0,11701743 | 0,002647 | 0,0275972 | FZR1 |
| 990,069452 | 0,35147195 | 0,11219322 | 0,00173184 | 0,02030244 | ECSIT |
| 565,762367 | 0,35130255 | 0,13880936 | 0,01137938 | 0,07567985 | MAPK13 |
| 818,792305 | 0,35082013 | 0,14520447 | 0,01569024 | 0,094276 | STK11IP |
| 818,769582 | 0,35052112 | 0,11677136 | 0,00268412 | 0,02791988 | SH3BP5L |
| 1118,56407 | 0,35037318 | 0,13940135 | 0,01195692 | 0,07847273 | ZNF276 |
| 378,633329 | 0,35025519 | 0,15977351 | 0,02836521 | 0,13907698 | RAB26 |
| 499,09182 | 0,35000011 | 0,13779316 | 0,01108399 | 0,0742635 | SLC25A10 |
| 1624,23282 | 0,34993406 | 0,13678426 | 0,01051878 | 0,07156625 | CAPN1 |
| 1155,776 | 0,34987355 | 0,10499772 | 0,00086162 | 0,01207607 | ANKRD13D |
| 821,801501 | 0,34942133 | 0,14139762 | 0,01346617 | 0,08481806 | ST6GALNAC6 |
| 1202,72385 | 0,3494076 | 0,08934129 | 9,19E-05 | 0,00212667 | SNHG7 |
| 1384,66354 | 0,34939468 | 0,1015508 | 0,00058045 | 0,00889987 | RIC8A |
| 3175,27667 | 0,34927952 | 0,11926693 | 0,00340544 | 0,03285787 | NCLN |
| 654,233012 | 0,34925107 | 0,15872124 | 0,02777815 | 0,13687248 | APBA3 |
| 2256,56095 | 0,3491419 | 0,11921949 | 0,00340532 | 0,03285787 | SCAF1 |
| 1281,4601 | 0,34884643 | 0,1196091 | 0,00353921 | 0,03389453 | MAP3K6 |
| 484,867593 | 0,34871721 | 0,17239695 | 0,04309816 | 0,18359279 | COL7A1 |
| 2957,93705 | 0,3484639 | 0,15820948 | 0,02762666 | 0,13645109 | H6PD |
| 523,433568 | 0,34845213 | 0,16581517 | 0,03560157 | 0,16119103 | RCE1 |
| 906,520661 | 0,34842559 | 0,14548509 | 0,01662385 | 0,09786557 | CYLD |
| 799,788954 | 0,34826124 | 0,11099862 | 0,00170379 | 0,02007313 | TAB1 |
| 925,922728 | 0,34794506 | 0,15439302 | 0,02421933 | 0,12499313 | GHDC |
| 3766,16177 | 0,34748305 | 0,13764761 | 0,0115883 | 0,0766919 | ACAP1 |
| 718,023041 | 0,34744005 | 0,12303349 | 0,00474362 | 0,04192738 | SAMD10 |
| 831,050061 | 0,34665766 | 0,12540476 | 0,00570433 | 0,0477151 | LRWD1 |
| 1084,70319 | 0,34473011 | 0,12458086 | 0,00565541 | 0,0473897 | EPHB4 |
| 177,446654 | 0,34471404 | 0,1579471 | 0,02907522 | 0,14109575 | LOC728743 |
| 1269,26655 | 0,34452462 | 0,11148406 | 0,00199922 | 0,02248941 | BCL2L1 |
| 480,569403 | 0,34427107 | 0,12695623 | 0,0066933 | 0,05307793 | INPP5K |
| 598,703795 | 0,34421242 | 0,15436819 | 0,02575976 | 0,13031882 | UBA7 |
| 385,971191 | 0,34410278 | 0,13184493 | 0,00905673 | 0,06498384 | HEXIM2 |
| 602,758864 | 0,34395215 | 0,14732784 | 0,01956412 | 0,10871446 | TOLLIP |
| 1549,29732 | 0,34375612 | 0,12969725 | 0,00803845 | 0,05992733 | NCAPH2 |

| | | | | | |
|---|---|---|---|---|---|
| 508,265168 | 0,34365564 | 0,17052506 | 0,04387538 | 0,18567796 | HMOX1 |
| 2222,21331 | 0,34326247 | 0,13307528 | 0,00989547 | 0,06893294 | MGAT4B |
| 338,130752 | 0,34305949 | 0,15015283 | 0,02232835 | 0,11816188 | TMEM175 |
| 5816,99711 | 0,34303808 | 0,13505078 | 0,01108311 | 0,0742635 | INTS1 |
| 442,140244 | 0,342905 | 0,14620631 | 0,01900922 | 0,10660149 | GAMT |
| 1081,38621 | 0,34288435 | 0,12207502 | 0,00497265 | 0,04325862 | HIC2 |
| 758,644278 | 0,34279447 | 0,13688362 | 0,0122702 | 0,07974787 | SSNA1 |
| 262,901619 | 0,34211668 | 0,16310576 | 0,03594806 | 0,16223259 | HS1BP3 |
| 2125,1764 | 0,34199224 | 0,08959224 | 0,00013497 | 0,00286618 | THOP1 |
| 292,394013 | 0,3415458 | 0,17147223 | 0,04638827 | 0,19311605 | EVI5L |
| 3024,63001 | 0,34151206 | 0,11160602 | 0,00221353 | 0,02412564 | MGAT1 |
| 5288,05886 | 0,3414489 | 0,12180706 | 0,00505991 | 0,04382298 | HLA-E |
| 2476,79201 | 0,34132796 | 0,12105987 | 0,0048099 | 0,04233651 | SQSTM1 |
| 1353,43686 | 0,34101781 | 0,11584084 | 0,00324159 | 0,03174647 | CENPB |
| 795,374315 | 0,34093912 | 0,11156501 | 0,00224335 | 0,02440855 | GPR137 |
| 414,369234 | 0,34068074 | 0,1278006 | 0,00768234 | 0,05816328 | IFNGR2 |
| 2222,92862 | 0,33994464 | 0,12221341 | 0,00540974 | 0,04585916 | POLM |
| 325,566879 | 0,33988421 | 0,14230074 | 0,01691766 | 0,09876841 | C11orf68 |
| 675,546406 | 0,33971864 | 0,13634923 | 0,01271931 | 0,0816911 | SMAD3 |
| 1056,69213 | 0,33931126 | 0,13554897 | 0,01230629 | 0,07982425 | PNKP |
| 1789,60098 | 0,3392101 | 0,10256804 | 0,00094243 | 0,01291524 | ATG4B |
| 519,778822 | 0,33858482 | 0,14082732 | 0,01620546 | 0,09615035 | ANKS3 |
| 1105,13647 | 0,33847485 | 0,10814871 | 0,00174975 | 0,020487 | NCKAP5L |
| 892,235728 | 0,33831773 | 0,10120524 | 0,00082912 | 0,01174228 | DCAF11 |
| 2177,39856 | 0,33826478 | 0,12427879 | 0,00649231 | 0,0520292 | FAM234A |
| 664,522004 | 0,33775453 | 0,12070884 | 0,00514054 | 0,04427131 | PLEKHJ1 |
| 448,366409 | 0,33766056 | 0,12846315 | 0,00857718 | 0,06268399 | SUV39H1 |
| 269,621907 | 0,33765134 | 0,15779053 | 0,03236521 | 0,15128017 | FAM171A2 |
| 2063,32754 | 0,33736759 | 0,10837321 | 0,00185186 | 0,0213392 | ATAD3B |
| 806,154155 | 0,33653184 | 0,11544861 | 0,00355697 | 0,03402499 | TK1 |
| 801,206318 | 0,33612562 | 0,14133952 | 0,01740007 | 0,10037398 | STX11 |
| 915,62048 | 0,3359146 | 0,1045989 | 0,00132065 | 0,01658129 | ARPC4 |
| 288,846873 | 0,33548367 | 0,16265567 | 0,03915645 | 0,17249805 | DDIT3 |
| 2008,296 | 0,33521396 | 0,10926995 | 0,0021567 | 0,02364247 | POR |
| 439,722525 | 0,33511378 | 0,1604429 | 0,03673659 | 0,16490937 | MST1 |
| 1931,64333 | 0,33505627 | 0,098006 | 0,00062914 | 0,00945503 | NSUN5 |
| 1788,75532 | 0,33501158 | 0,13404716 | 0,01244716 | 0,08051718 | TRPV2 |
| 482,036586 | 0,33492087 | 0,15689166 | 0,03278331 | 0,15238185 | NTHL1 |
| 4441,98296 | 0,33471093 | 0,09378502 | 0,00035846 | 0,00617707 | DDX39A |
| 705,451486 | 0,33460394 | 0,14223322 | 0,01864759 | 0,10535414 | CYB561A3 |
| 3377,08115 | 0,33440365 | 0,10880164 | 0,00211552 | 0,02335341 | PITPNM1 |
| 374,926657 | 0,3341389 | 0,13951869 | 0,01662309 | 0,09786557 | GPR153 |
| 407,303605 | 0,33410629 | 0,14966701 | 0,02559311 | 0,12990526 | OGFOD2 |
| 8116,85302 | 0,33387206 | 0,14814207 | 0,02421323 | 0,12499313 | NIBAN2 |
| 1998,67779 | 0,33376499 | 0,10987792 | 0,00238472 | 0,02542691 | PPP6R2 |
| 3073,35319 | 0,333703 | 0,11394557 | 0,00340473 | 0,03285787 | CTDSP1 |
| 717,362783 | 0,33349964 | 0,14013427 | 0,01731932 | 0,10012184 | PTDSS2 |
| 3121,91233 | 0,33339431 | 0,08263201 | 5,47E-05 | 0,00139234 | CSNK1G2 |

| | | | | | |
|---|---|---|---|---|---|
| 3769,66291 | 0,33323928 | 0,14412738 | 0,02077127 | 0,11319505 | SCRIB |
| 392,643496 | 0,33315687 | 0,16602002 | 0,04477873 | 0,18836336 | DECR2 |
| 1663,83507 | 0,33308834 | 0,08686762 | 0,00012585 | 0,00272455 | RPS6KB2 |
| 663,49399 | 0,33260918 | 0,11364244 | 0,00342464 | 0,0329927 | RASSF5 |
| 1276,26837 | 0,33240678 | 0,11856953 | 0,00505551 | 0,04381826 | MPDU1 |
| 1239,71607 | 0,33232861 | 0,11989988 | 0,00557614 | 0,0469415 | FLYWCH1 |
| 305,18821 | 0,33225257 | 0,13690953 | 0,01523249 | 0,09241712 | ZNF212 |
| 623,377074 | 0,33223121 | 0,12915669 | 0,0101022 | 0,06985025 | ANKRD54 |
| 569,080646 | 0,3322061 | 0,14545073 | 0,02237289 | 0,11824508 | SLC4A3 |
| 1079,53861 | 0,33214548 | 0,0946369 | 0,00044864 | 0,00736871 | MUS81 |
| 928,976119 | 0,33214132 | 0,12389667 | 0,0073448 | 0,05652531 | WASH3P |
| 1422,15148 | 0,33206274 | 0,10300829 | 0,00126568 | 0,01605086 | RNF31 |
| 691,073586 | 0,33174856 | 0,12528061 | 0,0080959 | 0,06028435 | SIRT6 |
| 409,259398 | 0,33173773 | 0,1529707 | 0,03011035 | 0,14429626 | NR4A1 |
| 3400,9906 | 0,33139416 | 0,11362708 | 0,00353975 | 0,03389453 | PLXNB2 |
| 2099,92145 | 0,33100921 | 0,12989987 | 0,01082843 | 0,07306708 | GAA |
| 584,33127 | 0,3306409 | 0,11233495 | 0,00324683 | 0,03178129 | COQ8A |
| 764,52893 | 0,33035826 | 0,15118737 | 0,02888213 | 0,14055523 | LRRC75A |
| 487,352156 | 0,3302269 | 0,1258902 | 0,00871249 | 0,06334882 | ZNF785 |
| 6612,69391 | 0,33001917 | 0,09434759 | 0,00046892 | 0,00762897 | FURIN |
| 647,9789 | 0,32940307 | 0,14333559 | 0,02155469 | 0,11579679 | PSKH1 |
| 494,917886 | 0,32902895 | 0,14811096 | 0,0263169 | 0,13208183 | DGCR6L |
| 707,700402 | 0,32886748 | 0,14808032 | 0,02635961 | 0,13222435 | CCDC69 |
| 928,838481 | 0,3284281 | 0,14209717 | 0,02081671 | 0,11324699 | LZTS2 |
| 723,880142 | 0,3280833 | 0,12277997 | 0,00753728 | 0,05754077 | NR2F6 |
| 1341,93711 | 0,32800867 | 0,10573363 | 0,00192077 | 0,02184069 | XRCC3 |
| 651,685699 | 0,32741972 | 0,14896509 | 0,02795176 | 0,13751306 | TMEM120A |
| 252,392313 | 0,32710517 | 0,14886425 | 0,02799627 | 0,13766043 | RUSC2 |
| 1142,93335 | 0,3270343 | 0,12899898 | 0,01123929 | 0,07514438 | C1orf159 |
| 740,183616 | 0,32696481 | 0,09392118 | 0,00049905 | 0,00799102 | ZNF526 |
| 4388,62374 | 0,32676461 | 0,10393828 | 0,00166751 | 0,01980594 | GIGYF1 |
| 999,646663 | 0,32623963 | 0,14634337 | 0,0257956 | 0,13039771 | INPP5E |
| 3014,48266 | 0,32610821 | 0,11009594 | 0,00305611 | 0,03041077 | JUND |
| 874,609244 | 0,32599488 | 0,11870135 | 0,00602633 | 0,04948969 | USF1 |
| 246,222382 | 0,32579404 | 0,16007937 | 0,04183044 | 0,17955317 | UBE2Q2P1 |
| 1832,55485 | 0,3252064 | 0,13052168 | 0,0127173 | 0,0816911 | CLSTN3 |
| 416,794463 | 0,32466782 | 0,13602767 | 0,01699676 | 0,09916415 | TFEB |
| 360,710732 | 0,32453088 | 0,13767098 | 0,0184087 | 0,10447831 | MAP3K7CL |
| 1212,60815 | 0,32420468 | 0,12758746 | 0,01105236 | 0,07413037 | GTPBP6 |
| 1505,17973 | 0,32412132 | 0,13469987 | 0,01611731 | 0,09589177 | ACTR1B |
| 897,536567 | 0,32411935 | 0,11136475 | 0,00360931 | 0,0343866 | THAP11 |
| 4095,11292 | 0,32358739 | 0,12910037 | 0,01219403 | 0,07949617 | GALNS |
| 1123,75449 | 0,3235506 | 0,1429234 | 0,02358616 | 0,122905 | DDX51 |
| 573,235697 | 0,3233962 | 0,12320151 | 0,0086665 | 0,0630688 | RGL4 |
| 461,127443 | 0,3232825 | 0,14883614 | 0,02985039 | 0,14338644 | AGO4 |
| 475,930936 | 0,32297004 | 0,10330711 | 0,00177015 | 0,02070025 | PLCG2 |
| 6249,66924 | 0,32257507 | 0,09393388 | 0,0005946 | 0,00906563 | SLC25A39 |
| 589,241017 | 0,3224106 | 0,12173267 | 0,00808466 | 0,06022439 | LINC03009 |

| | | | | | |
|---|---|---|---|---|---|
| 8091,03044 | 0,32228273 | 0,12631482 | 0,01072836 | 0,07257462 | PIEZO1 |
| 1600,89947 | 0,32228222 | 0,12435848 | 0,00955424 | 0,0674285 | RNH1 |
| 2500,37448 | 0,32215685 | 0,13140496 | 0,01422093 | 0,08816323 | POLR2J3 |
| 545,643507 | 0,32203 | 0,16394319 | 0,04949787 | 0,20119208 | FAM174C |
| 645,337892 | 0,32091374 | 0,14907199 | 0,03133886 | 0,14821798 | ARHGEF19 |
| 1571,87027 | 0,32077492 | 0,14004341 | 0,02199008 | 0,11717101 | ADISSP |
| 1583,04548 | 0,32055415 | 0,11115254 | 0,00392768 | 0,03642927 | GTPBP3 |
| 1073,37082 | 0,32024778 | 0,11955702 | 0,00739263 | 0,05677248 | SLC8B1 |
| 702,889386 | 0,31992701 | 0,1437871 | 0,02608091 | 0,13138597 | NUTM2A |
| 2624,81596 | 0,3198799 | 0,15347784 | 0,03714116 | 0,16609499 | SLC2A1 |
| 1364,47122 | 0,31984764 | 0,14964377 | 0,03256599 | 0,15188095 | ALDH16A1 |
| 758,088053 | 0,31973273 | 0,15450408 | 0,03850736 | 0,17019338 | LRFN1 |
| 1325,794 | 0,31967637 | 0,12555496 | 0,01089318 | 0,07337481 | LOC102724023 |
| 886,004783 | 0,31963347 | 0,10073296 | 0,00150832 | 0,01838852 | CENPBD2P |
| 1865,02944 | 0,31953178 | 0,1177869 | 0,00667183 | 0,05299648 | PAK4 |
| 2238,73012 | 0,3192334 | 0,07983791 | 6,37E-05 | 0,0015683 | ACADVL |
| 274,952099 | 0,31906364 | 0,16113956 | 0,04769843 | 0,19665846 | SPSB2 |
| 1292,43881 | 0,31902173 | 0,09385972 | 0,00067652 | 0,01001799 | MGRN1 |
| 1806,33534 | 0,31882428 | 0,13992248 | 0,02269216 | 0,11963176 | SEMA4C |
| 182,195853 | 0,31856041 | 0,15948394 | 0,04577686 | 0,19145533 | TCTA |
| 1020,88046 | 0,31828468 | 0,12994322 | 0,0143089 | 0,08841893 | HPS1 |
| 238,325548 | 0,31811405 | 0,1438013 | 0,02695437 | 0,13445919 | EPHX1 |
| 586,79641 | 0,31809223 | 0,13538022 | 0,01879253 | 0,10573103 | SERTAD3 |
| 507,318487 | 0,31759908 | 0,12768502 | 0,01286938 | 0,082153 | DOLPP1 |
| 1015,52522 | 0,31730237 | 0,10844508 | 0,00343432 | 0,03301864 | PEPD |
| 559,001246 | 0,31705578 | 0,13616956 | 0,01989142 | 0,11004844 | STK16 |
| 1451,89614 | 0,31687557 | 0,10635817 | 0,00288884 | 0,02940277 | AGAP3 |
| 816,322958 | 0,31670274 | 0,1320656 | 0,01648166 | 0,09727056 | GUSBP11 |
| 880,752734 | 0,31648125 | 0,10456629 | 0,00247314 | 0,02612779 | WDR24 |
| 1787,74829 | 0,31612804 | 0,09523502 | 0,00090194 | 0,01244762 | PREB |
| 414,821586 | 0,31610408 | 0,12756308 | 0,01321132 | 0,08360244 | SGTB |
| 1411,09548 | 0,31603649 | 0,12879642 | 0,01413683 | 0,08781449 | NHERF1 |
| 2405,23481 | 0,31596201 | 0,13348478 | 0,01793165 | 0,10249258 | DNAJB1 |
| 921,602974 | 0,3155697 | 0,13256773 | 0,01729191 | 0,10005512 | GIPC1 |
| 1377,74332 | 0,3150094 | 0,10591689 | 0,00293831 | 0,02968277 | KIAA2013 |
| 1905,26377 | 0,31482711 | 0,1469456 | 0,03215537 | 0,15078359 | NPDC1 |
| 1936,18777 | 0,31434385 | 0,11816924 | 0,00781138 | 0,05881381 | KHNYN |
| 2393,17193 | 0,31370859 | 0,08785403 | 0,0003559 | 0,0061554 | PGAM5 |
| 2318,87986 | 0,31309812 | 0,09227319 | 0,0006909 | 0,01017318 | GUCD1 |
| 1994,92949 | 0,31308391 | 0,108975 | 0,00406609 | 0,03745592 | DHX37 |
| 520,81582 | 0,31259582 | 0,10251253 | 0,00229343 | 0,02474583 | ALG1 |
| 447,00559 | 0,31244246 | 0,14064357 | 0,02631578 | 0,13208183 | RAB40C |
| 703,007778 | 0,3124 | 0,13967863 | 0,02531495 | 0,12889777 | SERHL2 |
| 1311,10718 | 0,31203589 | 0,11282119 | 0,0056791 | 0,04754612 | PDXK |
| 3013,67194 | 0,31163325 | 0,13544301 | 0,02140045 | 0,11526271 | TONSL |
| 1085,78759 | 0,31150595 | 0,10800398 | 0,00392399 | 0,0364129 | ZNF672 |
| 2858,33042 | 0,31143594 | 0,10449057 | 0,00287762 | 0,02932755 | EML3 |
| 335,445285 | 0,31143358 | 0,14110169 | 0,0273031 | 0,13576922 | CSAD |

| | | | | | |
|---|---|---|---|---|---|
| 4082,31389 | 0,31132742 | 0,08912352 | 0,00047725 | 0,00772452 | GRAMD1A |
| 656,138015 | 0,31105927 | 0,15600711 | 0,04616531 | 0,19244191 | VPS9D1 |
| 364,562297 | 0,3106541 | 0,15398603 | 0,04365198 | 0,18506364 | FDXR |
| 646,004309 | 0,31063238 | 0,13448888 | 0,02090337 | 0,11365305 | HEXD |
| 647,924963 | 0,31056358 | 0,14731907 | 0,03502219 | 0,15925376 | FAM98C |
| 4844,41823 | 0,31045484 | 0,1038847 | 0,00280391 | 0,02878609 | TBRG4 |
| 2372,27369 | 0,31029531 | 0,10086195 | 0,00209491 | 0,02315297 | ARAP1 |
| 3352,78624 | 0,31025568 | 0,10006683 | 0,00193204 | 0,02192932 | PDLIM7 |
| 969,759948 | 0,31009787 | 0,09569289 | 0,00119298 | 0,0153252 | IKBKG |
| 1248,2645 | 0,30980416 | 0,10515587 | 0,00321764 | 0,03156152 | ZSWIM8 |
| 4616,24545 | 0,30925721 | 0,12034538 | 0,01017728 | 0,07007432 | MLLT6 |
| 1852,93735 | 0,30922437 | 0,12601135 | 0,01413003 | 0,0878011 | CHPF2 |
| 2311,2659 | 0,30826873 | 0,10829447 | 0,00441918 | 0,03974111 | POLRMT |
| 651,771695 | 0,30812362 | 0,14555145 | 0,03426497 | 0,15701806 | FCSK |
| 1275,28122 | 0,30790751 | 0,10817702 | 0,00442263 | 0,03975328 | ABHD2 |
| 406,499975 | 0,30789153 | 0,13210127 | 0,01976795 | 0,10958961 | USP43 |
| 1250,63922 | 0,30776312 | 0,13514841 | 0,02277289 | 0,11992371 | CERCAM |
| 4559,2609 | 0,30753422 | 0,11907161 | 0,00980115 | 0,06858965 | DENND4B |
| 5259,41709 | 0,30749635 | 0,11991741 | 0,01034037 | 0,07091341 | VASP |
| 1893,2727 | 0,30641253 | 0,11178752 | 0,00612476 | 0,0501023 | CCNF |
| 2202,33185 | 0,30607899 | 0,12356282 | 0,01324502 | 0,0837317 | SGSM2 |
| 1050,36669 | 0,3060604 | 0,11678307 | 0,0087734 | 0,06365452 | SLC12A4 |
| 532,466026 | 0,30563051 | 0,14245539 | 0,03191711 | 0,15002517 | COMMD5 |
| 3900,48203 | 0,30545284 | 0,09592242 | 0,00145068 | 0,0178351 | TYK2 |
| 1049,81484 | 0,3054076 | 0,14035596 | 0,02955897 | 0,1425657 | JRK |
| 2033,52458 | 0,30527355 | 0,10577151 | 0,00389973 | 0,03626072 | TRIM65 |
| 468,00835 | 0,30494768 | 0,14616675 | 0,03695146 | 0,16559891 | KAZALD1 |
| 5599,67585 | 0,3045187 | 0,13955479 | 0,02910417 | 0,1411638 | RALGDS |
| 1362,65838 | 0,30440667 | 0,11431166 | 0,0077458 | 0,05849564 | STARD3 |
| 688,217397 | 0,30374907 | 0,1155211 | 0,00855403 | 0,06258732 | SHROOM1 |
| 4365,50475 | 0,30370615 | 0,09823887 | 0,00199143 | 0,02244187 | BOP1 |
| 1495,76924 | 0,30332466 | 0,14997176 | 0,04312006 | 0,18363146 | TYMP |
| 1061,42917 | 0,30332146 | 0,1118724 | 0,00670167 | 0,05309977 | TRMT2A |
| 1322,77117 | 0,30267152 | 0,15176095 | 0,04610874 | 0,19241805 | STON1 |
| 379,733342 | 0,30225917 | 0,14008291 | 0,0309499 | 0,14709951 | TXNRD2 |
| 417,345663 | 0,30222831 | 0,15381592 | 0,04942927 | 0,20103246 | DMAC1 |
| 1043,50602 | 0,30185636 | 0,11577659 | 0,00912776 | 0,06529544 | GET4 |
| 622,520356 | 0,30185453 | 0,12670873 | 0,01720623 | 0,09970308 | KLHDC2 |
| 1484,48909 | 0,30154231 | 0,1158213 | 0,00922736 | 0,06578408 | MED16 |
| 1544,09309 | 0,30151593 | 0,12074715 | 0,01252188 | 0,08080714 | BCAT2 |
| 1975,26265 | 0,30082253 | 0,11330862 | 0,0079333 | 0,05945598 | VAT1 |
| 747,328912 | 0,30075349 | 0,11572616 | 0,009354 | 0,06634852 | CLN8 |
| 1892,39494 | 0,30057895 | 0,13361749 | 0,02447766 | 0,12592219 | BASP1P1 |
| 2117,79189 | 0,30054246 | 0,07731911 | 0,00010147 | 0,00231883 | MAP3K3 |
| 1196,18899 | 0,3005182 | 0,11198942 | 0,00728665 | 0,05629675 | ZNF768 |
| 1409,70327 | 0,30047086 | 0,10804936 | 0,0054214 | 0,04593744 | LIMD1 |
| 1339,41323 | 0,300467 | 0,09001921 | 0,00084442 | 0,01187895 | PRR7 |
| 1096,18697 | 0,30019755 | 0,1311354 | 0,02206668 | 0,11738083 | SLC35E2A |

| | | | | | |
|---|---|---|---|---|---|
| 1349,03008 | 0,30013319 | 0,13052676 | 0,02148225 | 0,11550604 | EPB41 |
| 520,92817 | 0,299916 | 0,13736545 | 0,02901051 | 0,14085396 | SLC25A29 |
| 954,776283 | 0,29990844 | 0,12488456 | 0,01632867 | 0,09663894 | STK11 |
| 4742,4217 | 0,29963667 | 0,10368123 | 0,00385266 | 0,03596264 | ARHGEF1 |
| 699,267297 | 0,29924715 | 0,13941728 | 0,03183999 | 0,14986182 | BCR |
| 380,831359 | 0,29921065 | 0,13250324 | 0,02393704 | 0,12397567 | DDX28 |
| 1858,83751 | 0,29909586 | 0,10944874 | 0,00628083 | 0,0510041 | HSD3B7 |
| 1684,36726 | 0,29903343 | 0,10218979 | 0,00343069 | 0,03301736 | TRAF4 |
| 2426,85688 | 0,29902164 | 0,10454158 | 0,00423223 | 0,03859963 | RNF187 |
| 561,281495 | 0,29827077 | 0,12968151 | 0,02144678 | 0,11547939 | TKFC |
| 1908,75579 | 0,29821578 | 0,11546525 | 0,00980215 | 0,06858965 | GGA1 |
| 1439,54103 | 0,2979784 | 0,10733053 | 0,00549868 | 0,04644662 | LITAF |
| 640,759569 | 0,29747301 | 0,14424866 | 0,03918644 | 0,1725841 | DDAH2 |
| 960,403829 | 0,2964312 | 0,0905779 | 0,00106538 | 0,01416592 | FAM53B |
| 772,549586 | 0,29638806 | 0,12409058 | 0,01691815 | 0,09876841 | TSEN54 |
| 2547,38582 | 0,29623837 | 0,11049857 | 0,0073419 | 0,05652531 | GRK2 |
| 1695,46884 | 0,29611 | 0,1263387 | 0,01908947 | 0,10686151 | PGPEP1 |
| 528,942755 | 0,29606945 | 0,14462021 | 0,04063645 | 0,17618748 | FSD1 |
| 1521,2397 | 0,29584237 | 0,13230199 | 0,02534429 | 0,12897774 | SPG7 |
| 1065,69004 | 0,29564358 | 0,10206583 | 0,00377234 | 0,03544015 | TMUB2 |
| 3124,30894 | 0,29558996 | 0,10848511 | 0,00643589 | 0,05171486 | HLA-C |
| 380,599098 | 0,29506395 | 0,14373839 | 0,04009348 | 0,17470962 | TTC5 |
| 483,349371 | 0,29459705 | 0,1224838 | 0,01616397 | 0,0960548 | ZCCHC2 |
| 289,388036 | 0,29445464 | 0,141085 | 0,03688157 | 0,16534453 | BCL2 |
| 1085,62675 | 0,2942542 | 0,12602537 | 0,01954965 | 0,10869791 | NEU1 |
| 962,018637 | 0,29420799 | 0,10544626 | 0,00526881 | 0,04504336 | PLCB2 |
| 1250,60512 | 0,2941982 | 0,1047041 | 0,00495714 | 0,04316339 | LRRC47 |
| 2523,57637 | 0,29419272 | 0,09954373 | 0,00312252 | 0,03086766 | CPNE1 |
| 1195,31856 | 0,29418744 | 0,11454271 | 0,01021796 | 0,07024456 | KLF16 |
| 2272,7458 | 0,29318311 | 0,10583185 | 0,00560093 | 0,04703741 | NOL6 |
| 1329,20915 | 0,29303326 | 0,09777517 | 0,00272641 | 0,02825102 | WSB2 |
| 3413,70249 | 0,29302396 | 0,10627081 | 0,0058275 | 0,04836062 | SSBP4 |
| 460,737832 | 0,29297896 | 0,13128695 | 0,02564192 | 0,13003289 | FZD1 |
| 422,046903 | 0,29267469 | 0,12125583 | 0,01579164 | 0,0946448 | FIZ1 |
| 1293,59794 | 0,29262399 | 0,11323488 | 0,00975992 | 0,06844603 | WDR18 |
| 1994,6342 | 0,29215712 | 0,11541445 | 0,01136167 | 0,07562245 | INPPL1 |
| 1208,76099 | 0,29194143 | 0,11869025 | 0,01390564 | 0,08683454 | THOC6 |
| 323,479132 | 0,29166686 | 0,12618013 | 0,02080461 | 0,11324699 | C21orf58 |
| 4548,92838 | 0,29141272 | 0,07873829 | 0,00021473 | 0,00415524 | CLASRP |
| 800,314327 | 0,29120366 | 0,12909098 | 0,02408305 | 0,12446747 | IGSF8 |
| 569,559652 | 0,29119564 | 0,14199429 | 0,04029072 | 0,17512995 | PHF13 |
| 1941,09875 | 0,29113656 | 0,08978002 | 0,00118371 | 0,01522567 | ARMC6 |
| 759,130732 | 0,29113519 | 0,12502111 | 0,01987559 | 0,11002518 | HOMER3 |
| 937,552161 | 0,29106356 | 0,11108114 | 0,00878579 | 0,06369192 | C7orf50 |
| 721,170523 | 0,29084773 | 0,14625934 | 0,04674808 | 0,19410149 | IER5L |
| 2398,04382 | 0,29078646 | 0,1335549 | 0,02945965 | 0,14241342 | SLC4A2 |
| 1556,04492 | 0,29030809 | 0,10561764 | 0,00598375 | 0,04929354 | GPRIN1 |
| 1632,38064 | 0,28963123 | 0,12016593 | 0,01594112 | 0,09514889 | DDX49 |

| | | | | | |
|---|---|---|---|---|---|
| 1983,58393 | 0,28897718 | 0,09033767 | 0,00137974 | 0,01714123 | RASAL3 |
| 4045,16109 | 0,28896295 | 0,12016909 | 0,01618857 | 0,09611044 | ARID5A |
| 620,475473 | 0,28894178 | 0,13942986 | 0,03823661 | 0,16949094 | ASL |
| 552,242993 | 0,28893234 | 0,12556642 | 0,02138983 | 0,11526271 | IRF9 |
| 5213,63755 | 0,28869563 | 0,09865258 | 0,00342924 | 0,03301736 | SMG5 |
| 748,22456 | 0,28816379 | 0,13301338 | 0,03027856 | 0,14485478 | MFSD13A |
| 1683,28553 | 0,28785132 | 0,09454048 | 0,0023288 | 0,02502175 | CENPT |
| 876,810684 | 0,28733936 | 0,13216568 | 0,02969878 | 0,14305163 | MNT |
| 1122,07638 | 0,2872858 | 0,11879747 | 0,01559398 | 0,09399611 | PSMG3 |
| 4076,6827 | 0,28728404 | 0,0947214 | 0,00242186 | 0,02574385 | HGS |
| 777,494696 | 0,2871516 | 0,09588205 | 0,00274587 | 0,02839048 | FAM78A |
| 854,313499 | 0,28688654 | 0,13126156 | 0,02884417 | 0,14051508 | MTG1 |
| 1058,73238 | 0,28678176 | 0,1023377 | 0,00507384 | 0,04388643 | UNC119B |
| 279,192935 | 0,28669702 | 0,12984397 | 0,02724345 | 0,13557951 | USP35 |
| 8303,65622 | 0,28661231 | 0,11856879 | 0,01563759 | 0,09416893 | TNRC18 |
| 565,7433 | 0,28621075 | 0,1215167 | 0,01850675 | 0,10477742 | DRG2 |
| 1652,61178 | 0,28572243 | 0,11213284 | 0,01083205 | 0,07306708 | PRKD2 |
| 1071,51033 | 0,28569171 | 0,10569036 | 0,00686959 | 0,05406801 | EPOP |
| 1959,94414 | 0,28476473 | 0,09619944 | 0,0030749 | 0,03057284 | FHOD1 |
| 674,398413 | 0,28470603 | 0,09358888 | 0,0023494 | 0,02514306 | ILKAP |
| 3173,82933 | 0,28430159 | 0,10896648 | 0,00907876 | 0,06504328 | FARSA |
| 1689,86862 | 0,28426253 | 0,1271127 | 0,02533192 | 0,12894951 | NELFB |
| 698,169393 | 0,28402648 | 0,13215576 | 0,03162016 | 0,14897537 | TMED1 |
| 416,76994 | 0,28363994 | 0,14303805 | 0,04737094 | 0,19578541 | JMJD8 |
| 1566,1793 | 0,28363728 | 0,11215714 | 0,01144116 | 0,07601061 | LRRC61 |
| 915,032257 | 0,2829728 | 0,10409367 | 0,00655898 | 0,05240811 | TBC1D22A |
| 739,762099 | 0,2829149 | 0,09595939 | 0,0031955 | 0,03142521 | FAM83G |
| 4197,97829 | 0,28276318 | 0,10135378 | 0,00527303 | 0,04504336 | MIDN |
| 429,819706 | 0,28248887 | 0,12300608 | 0,0216448 | 0,11605021 | AP4S1 |
| 1430,60514 | 0,28232279 | 0,13549457 | 0,03719233 | 0,16619784 | MAD2L2 |
| 1082,46361 | 0,28198926 | 0,0923945 | 0,00227312 | 0,02461916 | HECA |
| 829,673181 | 0,28198586 | 0,11997428 | 0,01875396 | 0,10563965 | VAV2 |
| 383,684081 | 0,28188344 | 0,14287706 | 0,04850592 | 0,19873984 | SH2B2 |
| 980,354732 | 0,28160071 | 0,13957445 | 0,04363644 | 0,18503925 | FADS3 |
| 1773,24628 | 0,28132783 | 0,1121167 | 0,01209909 | 0,07915819 | NR1H2 |
| 767,427653 | 0,28073148 | 0,12056629 | 0,01988871 | 0,11004844 | HES7 |
| 1069,37868 | 0,28042246 | 0,13126591 | 0,03265553 | 0,15211108 | PLCB3 |
| 1264,96988 | 0,28037957 | 0,10800527 | 0,00943214 | 0,06681603 | GALE |
| 12536,8151 | 0,28017933 | 0,08610362 | 0,00113801 | 0,0147785 | TOMM40 |
| 19316,9314 | 0,2789258 | 0,11305257 | 0,01361658 | 0,0855359 | RPSA |
| 1079,01303 | 0,27876354 | 0,1046348 | 0,00771815 | 0,05836738 | HGH1 |
| 1009,77611 | 0,27874967 | 0,11810203 | 0,01826289 | 0,10395673 | DAPK3 |
| 268,119466 | 0,27851646 | 0,13896744 | 0,04505024 | 0,18908507 | ZNF513 |
| 786,808418 | 0,27848582 | 0,12642934 | 0,02761595 | 0,13645109 | PHPT1 |
| 1415,33601 | 0,27756841 | 0,08212256 | 0,00072505 | 0,01055277 | MED22 |
| 4373,16618 | 0,27732796 | 0,10949623 | 0,01131675 | 0,07542243 | PSTPIP1 |
| 1035,72841 | 0,27653585 | 0,09660335 | 0,00420193 | 0,03844523 | FANCG |
| 10089,3872 | 0,27649095 | 0,11678326 | 0,01790599 | 0,10241815 | TNFRSF8 |

| | | | | | |
|---|---|---|---|---|---|
| 1374,51087 | 0,27620685 | 0,104601 | 0,00827653 | 0,06123603 | SAMD1 |
| 1859,89298 | 0,27618463 | 0,10356189 | 0,00765645 | 0,05815646 | PHRF1 |
| 1850,54065 | 0,27606234 | 0,10605661 | 0,00924196 | 0,06580486 | USP19 |
| 1388,27061 | 0,27576265 | 0,09277176 | 0,002954 | 0,02971657 | GRWD1 |
| 1009,11061 | 0,27572192 | 0,10630682 | 0,00949651 | 0,06709613 | CYTH1 |
| 506,561692 | 0,27556248 | 0,12235459 | 0,02431198 | 0,1253425 | ATOSA |
| 1053,88649 | 0,27551748 | 0,1313151 | 0,03589276 | 0,16216022 | SGSM3 |
| 1180,20825 | 0,27536942 | 0,09735089 | 0,00467481 | 0,04148551 | RIOK3 |
| 1512,46419 | 0,27514283 | 0,10291682 | 0,00750758 | 0,05744073 | PPP1R12C |
| 478,540375 | 0,27489654 | 0,11586855 | 0,01766882 | 0,10142956 | TRAPPC14 |
| 859,677526 | 0,27467845 | 0,13467679 | 0,04139635 | 0,17841921 | INTS5 |
| 974,435144 | 0,27439673 | 0,11725709 | 0,01927707 | 0,10756163 | IRF3 |
| 699,620669 | 0,27435587 | 0,13178292 | 0,03735367 | 0,16680879 | TWF2 |
| 1846,17595 | 0,27429551 | 0,1101213 | 0,01274383 | 0,08173473 | TAP1 |
| 608,391382 | 0,27418421 | 0,10053604 | 0,00638699 | 0,05149048 | CTDP1 |
| 606,399692 | 0,2740849 | 0,13941571 | 0,04930391 | 0,20070551 | FAM3A |
| 1218,4391 | 0,27394894 | 0,12005903 | 0,02250202 | 0,11879482 | ECE1 |
| 550,92793 | 0,27387878 | 0,11648898 | 0,01871734 | 0,10554062 | HASPIN |
| 442,160181 | 0,27358411 | 0,13769043 | 0,04692785 | 0,19446384 | NUDT8 |
| 1143,34737 | 0,27355917 | 0,10610615 | 0,00993266 | 0,06903373 | CDIPT |
| 852,630706 | 0,27340652 | 0,13943003 | 0,04989222 | 0,20201384 | LRSAM1 |
| 883,543794 | 0,27319068 | 0,13377255 | 0,04113142 | 0,17780389 | EBP |
| 636,825506 | 0,27292529 | 0,12361832 | 0,0272578 | 0,13561524 | FBXO44 |
| 658,68322 | 0,27286328 | 0,10042165 | 0,00658416 | 0,05258715 | WRAP53 |
| 855,823411 | 0,27282489 | 0,13874782 | 0,04925977 | 0,20064983 | CGAS |
| 837,757546 | 0,27280781 | 0,10210852 | 0,00754582 | 0,05757035 | NOC4L |
| 641,335291 | 0,27274456 | 0,10889525 | 0,01225724 | 0,07974787 | FANCE |
| 723,577614 | 0,27257075 | 0,09543648 | 0,00428956 | 0,038927 | TMCO6 |
| 987,76746 | 0,27240271 | 0,09890344 | 0,00588306 | 0,04865096 | CXXC1 |
| 2709,59842 | 0,272365 | 0,0928053 | 0,00333763 | 0,03241739 | EDC4 |
| 1380,95987 | 0,27229349 | 0,09457121 | 0,00398631 | 0,03690069 | CERK |
| 1899,28117 | 0,2717993 | 0,12012695 | 0,02366032 | 0,12308962 | GAS2L1 |
| 988,22813 | 0,27173818 | 0,10899209 | 0,01266004 | 0,08136571 | AKAP17A |
| 531,800205 | 0,27134212 | 0,11149833 | 0,01494958 | 0,09133142 | SNHG10 |
| 603,13979 | 0,27105849 | 0,1094874 | 0,01329734 | 0,08397831 | ASF1B |
| 1257,7425 | 0,27094989 | 0,1214566 | 0,02569192 | 0,13015177 | FKRP |
| 2011,00823 | 0,27054561 | 0,09444681 | 0,0041763 | 0,03825137 | SDF4 |
| 891,700384 | 0,2702989 | 0,12264479 | 0,0275302 | 0,13614715 | HTRA2 |
| 2497,48395 | 0,27026597 | 0,0980437 | 0,00584081 | 0,04842983 | GAK |
| 775,059772 | 0,26991517 | 0,11762929 | 0,0217546 | 0,11644095 | CDAN1 |
| 962,250695 | 0,26977489 | 0,10489445 | 0,01011509 | 0,06987498 | DAGLB |
| 1492,65888 | 0,26967195 | 0,13462507 | 0,045163 | 0,18943229 | TMEM201 |
| 3243,89244 | 0,26945488 | 0,09430948 | 0,00427484 | 0,03882106 | CTBP1 |
| 535,08678 | 0,26929133 | 0,11333076 | 0,01749416 | 0,10073252 | KCTD13 |
| 2126,13477 | 0,26905299 | 0,11260542 | 0,01687853 | 0,09868931 | CYB561 |
| 1374,01483 | 0,2687195 | 0,08764075 | 0,00216836 | 0,02372907 | IP6K1 |
| 955,527913 | 0,26832674 | 0,11574422 | 0,02043447 | 0,11192241 | SCAMP5 |
| 1613,99987 | 0,26824884 | 0,13053921 | 0,03988584 | 0,17421138 | SLC35E2B |

| | | | | | |
|---|---|---|---|---|---|
| 710,31875 | 0,26816756 | 0,10031572 | 0,00751235 | 0,05745397 | ZBTB34 |
| 2291,24746 | 0,26814831 | 0,13487471 | 0,04679744 | 0,19413398 | SNX17 |
| 870,789885 | 0,26808469 | 0,12702203 | 0,03481213 | 0,15864171 | PIK3R1 |
| 479,179253 | 0,26739625 | 0,13103684 | 0,04128872 | 0,17819492 | MIF4GD |
| 2523,95739 | 0,26702432 | 0,07343351 | 0,00027661 | 0,00506756 | MAP7D1 |
| 594,237733 | 0,26694811 | 0,1071701 | 0,01274263 | 0,08173473 | TARBP2 |
| 1068,24162 | 0,266836 | 0,11810888 | 0,02386861 | 0,12386672 | RTL10 |
| 1387,82781 | 0,26657693 | 0,12445513 | 0,03219734 | 0,15082232 | DCAF12 |
| 873,045367 | 0,26642085 | 0,12418338 | 0,03192227 | 0,15002517 | SFXN5 |
| 799,449078 | 0,26622606 | 0,13038485 | 0,04116659 | 0,17783402 | NAT9 |
| 3746,33031 | 0,2660963 | 0,09338039 | 0,00437749 | 0,03947881 | CSNK1D |
| 2169,60939 | 0,2657997 | 0,11017671 | 0,01584417 | 0,09477949 | PLIN3 |
| 2737,02802 | 0,2651393 | 0,09962742 | 0,00778376 | 0,05870028 | IMPDH1 |
| 1767,31275 | 0,26507683 | 0,13236561 | 0,04521906 | 0,18958135 | VARS2 |
| 2509,82961 | 0,26502718 | 0,08482311 | 0,00178126 | 0,02077884 | PYGB |
| 1350,72801 | 0,26407709 | 0,09585721 | 0,005871 | 0,04862295 | DVL2 |
| 625,185927 | 0,26315338 | 0,09609844 | 0,0061744 | 0,05040878 | SLC25A26 |
| 1314,33176 | 0,26292981 | 0,10917261 | 0,01602322 | 0,09560876 | RPS6KL1 |
| 1457,90583 | 0,26223405 | 0,11078976 | 0,01793525 | 0,10249258 | RNF123 |
| 1165,87526 | 0,26219954 | 0,11895797 | 0,02751478 | 0,13614715 | UBXN6 |
| 3162,35284 | 0,26204687 | 0,11450323 | 0,02210526 | 0,11747275 | MICALL1 |
| 2667,42662 | 0,26193804 | 0,1123977 | 0,019782 | 0,10963536 | DNAJC5 |
| 5356,9775 | 0,261645 | 0,09867718 | 0,00801292 | 0,05983288 | NFKB2 |
| 1531,78169 | 0,26143585 | 0,12750908 | 0,04033212 | 0,17522939 | STMN3 |
| 4543,58417 | 0,26083496 | 0,11203416 | 0,0199029 | 0,11007639 | UBE2S |
| 1129,29214 | 0,2603322 | 0,08697172 | 0,00275981 | 0,02850348 | ZNF574 |
| 993,491455 | 0,26012279 | 0,12090322 | 0,03143708 | 0,14844545 | SLC38A10 |
| 1116,91043 | 0,25987906 | 0,12352871 | 0,03539635 | 0,16064627 | TMUB1 |
| 518,924728 | 0,25944376 | 0,11451535 | 0,02347708 | 0,12257443 | PLPP5 |
| 1425,77268 | 0,25923195 | 0,11184163 | 0,02045757 | 0,1119689 | POLR3H |
| 1830,24856 | 0,25922213 | 0,09532611 | 0,00654164 | 0,05230397 | DCAF15 |
| 824,15186 | 0,25917562 | 0,13101677 | 0,04790766 | 0,19709955 | RASSF1 |
| 1070,83491 | 0,25901792 | 0,12514076 | 0,03846989 | 0,17006749 | ATP6V0B |
| 1734,47524 | 0,25890822 | 0,12383983 | 0,03655765 | 0,16434007 | FBXL18 |
| 1007,82094 | 0,25877982 | 0,0976032 | 0,00801718 | 0,05983945 | RNF11 |
| 1910,96014 | 0,25856288 | 0,11092057 | 0,01975002 | 0,10955444 | SNHG17 |
| 1492,61404 | 0,2582642 | 0,10780649 | 0,01659194 | 0,09776893 | SETD1B |
| 2403,67005 | 0,25817799 | 0,08694126 | 0,00298219 | 0,02988663 | BAP1 |
| 1748,3579 | 0,25780444 | 0,11880768 | 0,03001209 | 0,14390743 | SZT2 |
| 1264,55828 | 0,2574077 | 0,09331999 | 0,00580968 | 0,04825505 | TRIM8 |
| 3778,32242 | 0,25720302 | 0,09796041 | 0,00865012 | 0,06299274 | EFHD2 |
| 2103,77211 | 0,25688696 | 0,09095251 | 0,00473681 | 0,04191405 | TRAPPC12 |
| 8149,97953 | 0,25649991 | 0,11419691 | 0,02469636 | 0,12653605 | NBEAL2 |
| 1578,2922 | 0,25642584 | 0,09754881 | 0,00857138 | 0,06266579 | DNPEP |
| 935,839762 | 0,25636309 | 0,10623037 | 0,01580989 | 0,09466423 | MAP1LC3B |
| 2469,15427 | 0,25618601 | 0,08640074 | 0,00302594 | 0,03024502 | SCAP |
| 1888,81323 | 0,25618073 | 0,10486791 | 0,01457019 | 0,08959462 | VAC14 |
| 1767,65979 | 0,25577574 | 0,07237834 | 0,00040952 | 0,00688759 | FGD5-AS1 |

| | | | | | |
|---|---|---|---|---|---|
| 1015,23907 | 0,25574778 | 0,12214185 | 0,03627256 | 0,16325242 | SNX8 |
| 1204,26474 | 0,25549345 | 0,10434094 | 0,01433965 | 0,08858001 | AGTRAP |
| 2933,50295 | 0,25445879 | 0,09269319 | 0,00604791 | 0,04962383 | NCS1 |
| 663,097704 | 0,25377828 | 0,1070616 | 0,0177691 | 0,10185067 | MARCHF9 |
| 755,012885 | 0,25359723 | 0,11483447 | 0,02721836 | 0,13549129 | HROB |
| 608,210788 | 0,25357355 | 0,12207828 | 0,03778874 | 0,16823201 | GPN2 |
| 611,556628 | 0,25328907 | 0,10589416 | 0,01676106 | 0,09824751 | TNFAIP1 |
| 7626,53009 | 0,25324391 | 0,11826095 | 0,03224185 | 0,15092728 | FKBP8 |
| 2369,06722 | 0,25315967 | 0,09574662 | 0,00819185 | 0,06077898 | H2AX |
| 3319,05797 | 0,25305554 | 0,08864631 | 0,00430822 | 0,03902634 | TEAD3 |
| 2981,5609 | 0,25299903 | 0,12457665 | 0,04226815 | 0,18098116 | IRF2BPL |
| 2336,9164 | 0,2525818 | 0,085881 | 0,00327084 | 0,03195017 | PACS2 |
| 1184,03828 | 0,25241049 | 0,09607455 | 0,00860817 | 0,06288615 | PUS1 |
| 1645,02302 | 0,25159809 | 0,11119779 | 0,02365926 | 0,12308962 | NSUN5P1 |
| 1557,64544 | 0,25153462 | 0,11930403 | 0,03500075 | 0,15919452 | SNAPC4 |
| 1181,77037 | 0,251425 | 0,10907053 | 0,02115763 | 0,11451083 | MRPL38 |
| 2641,13561 | 0,25126755 | 0,10786185 | 0,01983094 | 0,10987446 | DUS1L |
| 3478,94473 | 0,2512246 | 0,08449992 | 0,00294832 | 0,02971657 | RANGAP1 |
| 1275,19849 | 0,25081762 | 0,11223329 | 0,02543126 | 0,12931601 | TSC22D2 |
| 2531,37959 | 0,25062841 | 0,12486933 | 0,04473632 | 0,18822679 | ADAM15 |
| 1851,54152 | 0,25055559 | 0,08632253 | 0,00370136 | 0,03496431 | KLHDC3 |
| 551,768537 | 0,25037801 | 0,11165189 | 0,02492986 | 0,12731904 | SHPK |
| 622,753684 | 0,25035737 | 0,10300648 | 0,01507796 | 0,09182171 | ACAA1 |
| 1659,48328 | 0,25035665 | 0,09348842 | 0,00740758 | 0,05685959 | LYST |
| 2471,57475 | 0,24997441 | 0,10052117 | 0,01289037 | 0,08225928 | USF2 |
| 3013,80716 | 0,24976134 | 0,120003 | 0,03740715 | 0,16700818 | PTPA |
| 988,134113 | 0,24969911 | 0,12650185 | 0,04839571 | 0,19850321 | MIER2 |
| 2209,19952 | 0,24961531 | 0,08044738 | 0,00191674 | 0,02182104 | NT5DC2 |
| 3375,9534 | 0,24953178 | 0,09211087 | 0,00674787 | 0,05335416 | FBRSL1 |
| 1457,67484 | 0,24927911 | 0,11687606 | 0,03293703 | 0,15284076 | ZBTB17 |
| 561,562189 | 0,24909162 | 0,1175034 | 0,03401724 | 0,15637515 | TUFT1 |
| 900,197208 | 0,24865342 | 0,10385727 | 0,01665738 | 0,09801249 | YIPF2 |
| 734,130486 | 0,24822293 | 0,09504022 | 0,00900756 | 0,06475381 | OGFOD3 |
| 782,56673 | 0,24798885 | 0,10437528 | 0,01750456 | 0,10076002 | ARHGAP26 |
| 349,097773 | 0,24797333 | 0,12258432 | 0,04308555 | 0,18359279 | NCBP2AS2 |
| 1851,79683 | 0,24778711 | 0,08551691 | 0,00376124 | 0,03538854 | SLC35C2 |
| 634,712297 | 0,24764953 | 0,1057844 | 0,01922815 | 0,10735195 | RUSC1 |
| 4667,96782 | 0,24733879 | 0,09485352 | 0,00911829 | 0,06525235 | PPP6R1 |
| 656,620506 | 0,24723338 | 0,12507736 | 0,04808191 | 0,19753132 | TUT1 |
| 905,20933 | 0,24685151 | 0,12538723 | 0,04898602 | 0,19997776 | NPHP4 |
| 4194,54866 | 0,24683446 | 0,08682006 | 0,00446829 | 0,04004955 | DHX30 |
| 1358,05763 | 0,24676264 | 0,11133352 | 0,02666226 | 0,13335358 | BRD1 |
| 1553,37664 | 0,2467546 | 0,10693627 | 0,0210274 | 0,11403253 | MAP3K14 |
| 1549,20428 | 0,24671045 | 0,09530449 | 0,00963516 | 0,06783714 | NFKBIB |
| 46065,8243 | 0,24663395 | 0,1087156 | 0,02329169 | 0,12195493 | RPLP0 |
| 672,472159 | 0,24626984 | 0,11213847 | 0,02808317 | 0,13798017 | BIN3 |
| 1895,5112 | 0,24580723 | 0,10144359 | 0,015389 | 0,09302723 | HDHD5 |
| 3071,17922 | 0,24571715 | 0,10246047 | 0,01647743 | 0,09727056 | TCF25 |

| | | | | | |
|---|---|---|---|---|---|
| 1912,78573 | 0,24534123 | 0,10414643 | 0,01848617 | 0,10469228 | PTPRJ |
| 1372,61754 | 0,24529921 | 0,08091423 | 0,00243265 | 0,02582105 | GATC |
| 2213,62536 | 0,24507112 | 0,07963479 | 0,00208788 | 0,02312931 | DBNL |
| 2065,15312 | 0,24496794 | 0,12014383 | 0,04145445 | 0,17846635 | ZDHHC8 |
| 1305,22101 | 0,24490236 | 0,10988183 | 0,02582859 | 0,13049472 | RNF126 |
| 751,200643 | 0,24420854 | 0,11860329 | 0,03949071 | 0,17323187 | KPTN |
| 898,946824 | 0,24409845 | 0,11459224 | 0,03315938 | 0,15370129 | NAA40 |
| 1337,74104 | 0,24395516 | 0,09673006 | 0,0116683 | 0,07705967 | CDCA5 |
| 501,14645 | 0,24374612 | 0,11123497 | 0,02843208 | 0,1392646 | SNX29 |
| 841,35043 | 0,24364284 | 0,09656217 | 0,01163019 | 0,07690552 | OTUD5 |
| 929,550761 | 0,24362784 | 0,08390139 | 0,00368734 | 0,03489169 | KCTD5 |
| 2019,7678 | 0,24337968 | 0,08809284 | 0,00573143 | 0,04785719 | INTS11 |
| 943,538416 | 0,24329329 | 0,09559175 | 0,01092375 | 0,0734904 | ZNF646 |
| 610,528208 | 0,24328853 | 0,10829594 | 0,024671 | 0,12653605 | RPUSD3 |
| 46879,8122 | 0,24314879 | 0,10992135 | 0,02696493 | 0,1344764 | FLNA |
| 741,302486 | 0,2429814 | 0,1056521 | 0,0214581 | 0,11550604 | MTMR14 |
| 1075,57344 | 0,24236383 | 0,11699006 | 0,038297 | 0,16965994 | PCBP4 |
| 1477,42272 | 0,2422389 | 0,12101728 | 0,04531824 | 0,18987297 | ENTPD6 |
| 1099,09042 | 0,24189361 | 0,10511478 | 0,02137846 | 0,11524271 | MAST3 |
| 2076,04517 | 0,24142603 | 0,1151322 | 0,03599834 | 0,16232712 | DEF8 |
| 3842,52681 | 0,2405001 | 0,09713556 | 0,01328925 | 0,08395521 | MAN2A2 |
| 1130,2087 | 0,24025885 | 0,1114715 | 0,03113533 | 0,1477581 | INTS15 |
| 870,190073 | 0,23908461 | 0,11794248 | 0,04264924 | 0,18212339 | HSD17B1 |
| 3355,37191 | 0,23889264 | 0,098441 | 0,01523434 | 0,09241712 | ATP13A1 |
| 757,384142 | 0,23884056 | 0,10734417 | 0,02608137 | 0,13138597 | BTBD6 |
| 547,733241 | 0,23867324 | 0,10423689 | 0,02203758 | 0,117292 | METTL22 |
| 1121,80097 | 0,23824986 | 0,09588101 | 0,01296063 | 0,08245733 | TBC1D2B |
| 2704,03207 | 0,23779556 | 0,10010723 | 0,01752953 | 0,10081347 | ITPK1 |
| 2161,83694 | 0,23743475 | 0,10798038 | 0,02788721 | 0,1373026 | WDR6 |
| 745,532545 | 0,23724735 | 0,1198233 | 0,04770614 | 0,19665846 | BRSK1 |
| 381,176741 | 0,23645068 | 0,11689696 | 0,04310127 | 0,18359279 | SLC45A4 |
| 1047,20772 | 0,23644767 | 0,10863762 | 0,02951934 | 0,1425215 | RXRB |
| 4368,24105 | 0,23579111 | 0,09407184 | 0,01219329 | 0,07949617 | POLD2 |
| 1126,90102 | 0,23565068 | 0,1000061 | 0,01845487 | 0,10462449 | SMIM3 |
| 1225,61529 | 0,23564074 | 0,10165644 | 0,02044873 | 0,11195326 | ZNF282 |
| 1223,9614 | 0,23563118 | 0,11034611 | 0,03272977 | 0,15223204 | TOB2 |
| 1504,34904 | 0,23510027 | 0,11349069 | 0,03830856 | 0,1696679 | GFUS |
| 6465,64865 | 0,23505833 | 0,08815879 | 0,00766898 | 0,05816328 | PCED1A |
| 1036,73295 | 0,23472211 | 0,09095677 | 0,00986318 | 0,0688131 | MOCS3 |
| 4861,20512 | 0,23460642 | 0,10032258 | 0,01936026 | 0,10786674 | PRDX2 |
| 6059,5418 | 0,23452317 | 0,1014576 | 0,02080312 | 0,11324699 | SHKBP1 |
| 1637,70194 | 0,23452069 | 0,09895532 | 0,01778975 | 0,10190732 | RBM15B |
| 1804,96802 | 0,2336835 | 0,10203579 | 0,02200908 | 0,11720627 | DUS3L |
| 12173,9268 | 0,23344883 | 0,10208926 | 0,02221258 | 0,11773606 | JAK3 |
| 9643,99165 | 0,23325337 | 0,09078426 | 0,01018997 | 0,07013622 | MYBL2 |
| 1282,13749 | 0,23296033 | 0,10811358 | 0,03117953 | 0,14776594 | DPH2 |
| 8222,72481 | 0,23183717 | 0,09491544 | 0,01458329 | 0,08964599 | COLGALT1 |
| 5322,80475 | 0,23177341 | 0,07527095 | 0,00207566 | 0,02304782 | RNPS1 |

| | | | | | |
|---|---|---|---|---|---|
| 2669,38003 | 0,2317381 | 0,10105261 | 0,02183407 | 0,11663537 | STAT6 |
| 1508,63784 | 0,23160862 | 0,09286119 | 0,01262634 | 0,08128719 | STK10 |
| 1293,84334 | 0,23118009 | 0,09873253 | 0,01920753 | 0,10730018 | ZNF687 |
| 2248,36563 | 0,23103247 | 0,09650776 | 0,0166691 | 0,09801249 | RAI1 |
| 851,872155 | 0,23067464 | 0,11241657 | 0,04017326 | 0,17473983 | WRAP73 |
| 1232,58187 | 0,23061414 | 0,0915528 | 0,01177155 | 0,07749811 | KDM2B |
| 7282,98387 | 0,23054633 | 0,09421962 | 0,01440893 | 0,08889189 | TPD52L2 |
| 713,396276 | 0,23049745 | 0,11449146 | 0,04409062 | 0,18625555 | SLC25A11 |
| 3997,68462 | 0,23045661 | 0,10847536 | 0,03362781 | 0,15503698 | CPSF1 |
| 1021,84953 | 0,23020462 | 0,10223002 | 0,02433302 | 0,1254168 | NCDN |
| 2051,64556 | 0,22994552 | 0,09237535 | 0,01280122 | 0,08189429 | ERF |
| 3944,92541 | 0,22908458 | 0,10698631 | 0,03225377 | 0,15094572 | SLC39A1 |
| 2490,99689 | 0,22861867 | 0,101683 | 0,02455407 | 0,12614376 | SMPD4BP |
| 614,826197 | 0,22842443 | 0,1136555 | 0,04445273 | 0,18745044 | MAD2L1BP |
| 624,246063 | 0,22808919 | 0,11338932 | 0,04426655 | 0,18679023 | DXO |
| 1402,16253 | 0,22786675 | 0,10318328 | 0,02721856 | 0,13549129 | ARHGEF2 |
| 2610,02737 | 0,22750932 | 0,09115124 | 0,01256188 | 0,08103762 | ALKBH5 |
| 6474,21186 | 0,22749999 | 0,0859374 | 0,00811432 | 0,06037398 | AKT2 |
| 1092,05402 | 0,22744638 | 0,09037326 | 0,01184448 | 0,077924 | LEMD2 |
| 1123,58595 | 0,22743711 | 0,1031736 | 0,0274954 | 0,13612452 | PI4K2A |
| 1951,783 | 0,22620254 | 0,10627861 | 0,03330459 | 0,15403473 | REXO1 |
| 1152,22012 | 0,22602105 | 0,10579966 | 0,03265402 | 0,15211108 | WDR77 |
| 8136,41475 | 0,22598949 | 0,11305877 | 0,0456227 | 0,19093722 | CLIC1 |
| 10296,7091 | 0,2254431 | 0,09346015 | 0,01585727 | 0,09479174 | CALM3 |
| 3601,75959 | 0,22531572 | 0,07542891 | 0,00281613 | 0,02888019 | CSK |
| 789,206357 | 0,22529403 | 0,10294583 | 0,02863527 | 0,13992972 | KDM6A |
| 661,874534 | 0,2247343 | 0,1063958 | 0,0346652 | 0,15820072 | VPS33A |
| 1032,63257 | 0,22346392 | 0,09337555 | 0,01670333 | 0,09805887 | CBFA2T2 |
| 2358,58999 | 0,22337551 | 0,08785313 | 0,01100306 | 0,0739045 | ANKRD10 |
| 812,563682 | 0,2231385 | 0,09662143 | 0,02092085 | 0,11368275 | BLOC1S3 |
| 2335,7548 | 0,22291925 | 0,09433022 | 0,01811888 | 0,10344821 | CLPTM1L |
| 1429,19533 | 0,22231803 | 0,10241038 | 0,02994218 | 0,14360862 | TBL2 |
| 1211,30671 | 0,22190811 | 0,08639005 | 0,01020877 | 0,07022914 | UBE2J2 |
| 1153,32533 | 0,22183598 | 0,09777863 | 0,02328309 | 0,12195493 | CRCP |
| 1569,90127 | 0,22149764 | 0,09333328 | 0,01763517 | 0,10132859 | CARS2 |
| 1577,31801 | 0,221426 | 0,09813906 | 0,02405515 | 0,12435722 | BSDC1 |
| 1132,90295 | 0,22120847 | 0,0886983 | 0,0126333 | 0,08130428 | LINC00540 |
| 1593,82761 | 0,22114364 | 0,08341621 | 0,00802332 | 0,05986165 | LPCAT1 |
| 2001,46434 | 0,22092583 | 0,10702605 | 0,0389964 | 0,17195313 | CD81 |
| 4552,67387 | 0,22085182 | 0,09901796 | 0,02571944 | 0,13022146 | ERCC2 |
| 1702,76421 | 0,22076285 | 0,09019553 | 0,014381 | 0,08877745 | FEM1A |
| 1127,96056 | 0,22038047 | 0,10108124 | 0,02924034 | 0,14163095 | PEX26 |
| 1144,14763 | 0,2200154 | 0,10064379 | 0,02880974 | 0,14044843 | PISD |
| 6493,17524 | 0,2197189 | 0,09957455 | 0,02734363 | 0,13584218 | TAGLN2 |
| 1181,66666 | 0,2196368 | 0,11130357 | 0,04845989 | 0,19859422 | ACSF3 |
| 4335,36612 | 0,21933351 | 0,08268158 | 0,00798391 | 0,05970894 | TTLL12 |
| 6323,2292 | 0,21919174 | 0,08844674 | 0,01320344 | 0,08358054 | TIMM50 |
| 714,923519 | 0,21883588 | 0,11006027 | 0,04677547 | 0,19412999 | STRADB |

| | | | | | |
|---|---|---|---|---|---|
| 1071,64603 | 0,21880042 | 0,08927163 | 0,01424792 | 0,08827274 | TRIM35 |
| 2175,58025 | 0,21864955 | 0,07882587 | 0,00554006 | 0,04672949 | ST3GAL1 |
| 9142,33146 | 0,21855 | 0,08134025 | 0,00721268 | 0,05593078 | SURF4 |
| 6323,29475 | 0,21807776 | 0,07627128 | 0,0042466 | 0,03868556 | ARHGDIA |
| 1510,05313 | 0,21794501 | 0,08184557 | 0,00774748 | 0,05849564 | HAUS5 |
| 1055,6563 | 0,21776201 | 0,10565948 | 0,03930502 | 0,17281218 | SLC35B2 |
| 2176,12905 | 0,2177573 | 0,09401991 | 0,0205541 | 0,11233508 | STX1A |
| 3280,9156 | 0,21702035 | 0,09257361 | 0,01906281 | 0,10680706 | PRPF19 |
| 1053,18642 | 0,21695369 | 0,10568611 | 0,04009084 | 0,17470962 | DPH7 |
| 4633,43611 | 0,21685 | 0,08675157 | 0,01243103 | 0,0804403 | MAPKAPK2 |
| 504,37444 | 0,21678654 | 0,10994135 | 0,0486281 | 0,19911107 | SFT2D2 |
| 556,562671 | 0,21671939 | 0,10982313 | 0,04845579 | 0,19859422 | ZBTB45 |
| 1339,55704 | 0,21621149 | 0,10695553 | 0,04322717 | 0,18383936 | MRPL49 |
| 3138,68773 | 0,21616928 | 0,09231199 | 0,01919486 | 0,10726107 | RABL6 |
| 2344,46262 | 0,21533094 | 0,08663417 | 0,01293624 | 0,082413 | NDST2 |
| 2896,7081 | 0,21465996 | 0,0955697 | 0,02469703 | 0,12653605 | PUF60 |
| 7418,29213 | 0,21429261 | 0,08327849 | 0,01007616 | 0,06975888 | USP22 |
| 1669,94934 | 0,21413954 | 0,10464082 | 0,04071461 | 0,17632437 | EPN1 |
| 1353,59373 | 0,21413762 | 0,10400223 | 0,03949698 | 0,17323187 | GIT1 |
| 664,498272 | 0,21398042 | 0,10916759 | 0,049983 | 0,20227277 | ABHD16A |
| 1914,96983 | 0,21357337 | 0,09791509 | 0,0291679 | 0,14140042 | DIP2A |
| 1354,46958 | 0,21344663 | 0,10477517 | 0,04163128 | 0,17898331 | GGT1 |
| 1382,88179 | 0,21273065 | 0,10574058 | 0,04423927 | 0,18671676 | ARFGAP2 |
| 3483,00243 | 0,21245266 | 0,10195634 | 0,0371819 | 0,16619784 | ZNF496 |
| 2467,46908 | 0,2123974 | 0,08075157 | 0,00853203 | 0,06245052 | CIRBP |
| 2575,31675 | 0,2123066 | 0,09123097 | 0,0199582 | 0,11028885 | BCAP31 |
| 5612,74067 | 0,21186342 | 0,07613039 | 0,00538751 | 0,04573218 | RCC2 |
| 6818,7223 | 0,21169869 | 0,08800863 | 0,0161534 | 0,09602217 | DNM2 |
| 2562,18449 | 0,21169404 | 0,10205356 | 0,0380475 | 0,1689502 | SKIC2 |
| 767,182481 | 0,21145787 | 0,10614967 | 0,04636307 | 0,19308891 | EOLA2 |
| 1761,62933 | 0,21079932 | 0,07994562 | 0,00836961 | 0,06161923 | ENTR1 |
| 2586,02276 | 0,21062802 | 0,10696222 | 0,04893226 | 0,19992386 | TSC2 |
| 2255,18984 | 0,21054554 | 0,08854711 | 0,01741721 | 0,10041166 | MFHAS1 |
| 812,986826 | 0,21051532 | 0,10689392 | 0,04890886 | 0,19987141 | LMNA |
| 679,195271 | 0,21048012 | 0,09827361 | 0,03221146 | 0,15082232 | OSER1 |
| 3140,53827 | 0,21045928 | 0,08436413 | 0,01260801 | 0,08125213 | SKI |
| 3211,80917 | 0,2103738 | 0,10162071 | 0,03843491 | 0,16995254 | MYO9B |
| 5304,03844 | 0,20990332 | 0,08299721 | 0,01143748 | 0,07601061 | PPIF |
| 3168,16493 | 0,20989959 | 0,08727541 | 0,01617134 | 0,09606841 | NPLOC4 |
| 899,084881 | 0,20947366 | 0,08782149 | 0,01706887 | 0,09931121 | BPNT1 |
| 1294,0395 | 0,20903999 | 0,08630691 | 0,01543306 | 0,09320426 | LSM14B |
| 1155,85518 | 0,20863672 | 0,08752717 | 0,01714053 | 0,09945292 | BIRC5 |
| 2440,14044 | 0,20861152 | 0,09493052 | 0,02798348 | 0,13763334 | PDPK1 |
| 35519,6264 | 0,2085533 | 0,09374777 | 0,02610666 | 0,13144334 | ACTG1 |
| 3004,09664 | 0,20849808 | 0,07000115 | 0,00289668 | 0,02944967 | RAPGEF1 |
| 1889,81214 | 0,20796324 | 0,10367687 | 0,04486918 | 0,18865994 | RBM42 |
| 1192,81193 | 0,20796303 | 0,08871991 | 0,01907611 | 0,10682802 | ARAF |
| 967,72282 | 0,20750683 | 0,10088896 | 0,03970697 | 0,17383052 | CLK3 |

| | | | | | |
|---|---|---|---|---|---|
| 2378,25587 | 0,20716041 | 0,07784448 | 0,00778607 | 0,05870028 | POLDIP2 |
| 3044,78586 | 0,20685643 | 0,08037433 | 0,01006283 | 0,06969209 | E2F4 |
| 1003,60043 | 0,20657114 | 0,09977295 | 0,03841376 | 0,16993845 | SHC1 |
| 3151,19261 | 0,20568033 | 0,08091492 | 0,01102412 | 0,07399341 | PES1 |
| 1656,16485 | 0,20458559 | 0,09383607 | 0,02923933 | 0,14163095 | TRMT1 |
| 3435,99375 | 0,20443258 | 0,09759011 | 0,03618803 | 0,16302705 | NOC2L |
| 1655,75286 | 0,20325979 | 0,08804959 | 0,02097304 | 0,11383561 | RAB35 |
| 978,34737 | 0,20314631 | 0,09370193 | 0,03015826 | 0,14442532 | VCPIP1 |
| 3450,77382 | 0,20301161 | 0,10298522 | 0,04869307 | 0,19924774 | SNHG3 |
| 8359,17453 | 0,20218129 | 0,06988708 | 0,00381617 | 0,03569239 | PREX1 |
| 1227,47807 | 0,20204396 | 0,10269382 | 0,04913247 | 0,20028572 | SFI1 |
| 4080,08661 | 0,20156616 | 0,08559442 | 0,01852777 | 0,10483373 | RTN4 |
| 2792,80644 | 0,20156571 | 0,1024529 | 0,04913727 | 0,20028572 | LIMK1 |
| 1742,97234 | 0,20119852 | 0,08565374 | 0,01882515 | 0,10578875 | TMEM30A |
| 1188,41726 | 0,2011446 | 0,08359579 | 0,01612133 | 0,09589177 | PARP12 |
| 1662,62214 | 0,20032131 | 0,10029986 | 0,04580083 | 0,19151326 | ATAD3A |
| 1477,52178 | 0,20031937 | 0,08592203 | 0,01973183 | 0,10948559 | ZDHHC18 |
| 1740,67243 | 0,19985891 | 0,0970035 | 0,03936732 | 0,17302418 | FIBP |
| 801,934071 | 0,1993159 | 0,10023877 | 0,04676623 | 0,19412999 | ARFIP2 |
| 2165,02639 | 0,19841784 | 0,10027823 | 0,04785281 | 0,19704526 | TATDN2 |
| 2065,84646 | 0,19835223 | 0,09450704 | 0,03583373 | 0,16204829 | TUBA1A |
| 7977,01351 | 0,19758785 | 0,07648929 | 0,00978859 | 0,06858965 | SEPTIN9 |
| 1568,42908 | 0,19742731 | 0,08914759 | 0,02678671 | 0,13376389 | FAM53C |
| 1113,82697 | 0,19597703 | 0,0957077 | 0,04059312 | 0,17605585 | INO80E |
| 1060,62175 | 0,19546308 | 0,09317496 | 0,0359222 | 0,16223259 | PLAG1 |
| 1053,82127 | 0,19475839 | 0,09442442 | 0,03915208 | 0,17249805 | RNF26 |
| 2490,99688 | 0,19388308 | 0,09677575 | 0,04513156 | 0,18934239 | HSF1 |
| 1012,94347 | 0,19370825 | 0,09858645 | 0,04943083 | 0,20103246 | ZNF623 |
| 1245,57095 | 0,19320765 | 0,08862828 | 0,02925914 | 0,14163095 | C15orf39 |
| 1005,52765 | 0,19256539 | 0,09643388 | 0,0458399 | 0,19159194 | BAX |
| 2466,55361 | 0,19218936 | 0,0897195 | 0,03218434 | 0,15082232 | AP1B1 |
| 2972,21628 | 0,19195432 | 0,08252933 | 0,0200243 | 0,11054825 | MFN2 |
| 1225,04058 | 0,19184286 | 0,09502242 | 0,04349532 | 0,18460633 | WDR73 |
| 3456,48564 | 0,19134396 | 0,08740626 | 0,02858715 | 0,13983906 | TCF3 |
| 1117,24008 | 0,1902052 | 0,09086569 | 0,03632626 | 0,16345521 | USP4 |
| 2372,72018 | 0,18921975 | 0,08722868 | 0,03006463 | 0,14412285 | AEN |
| 5005,36525 | 0,18907988 | 0,06600802 | 0,0041767 | 0,03825137 | CERS2 |
| 7375,41103 | 0,18884954 | 0,0619922 | 0,00231643 | 0,02494547 | STRN4 |
| 1920,38786 | 0,18815036 | 0,09548138 | 0,04877593 | 0,1994574 | MAU2 |
| 931,487694 | 0,18733608 | 0,09378318 | 0,04576606 | 0,19145248 | PIAS4 |
| 3985,23205 | 0,18663674 | 0,06301147 | 0,00305699 | 0,03041077 | RNF10 |
| 2137,39121 | 0,18622988 | 0,0880818 | 0,03449109 | 0,15770813 | GEMIN4 |
| 3428,60001 | 0,18560533 | 0,09133809 | 0,04214664 | 0,18054259 | RBM3 |
| 2266,36557 | 0,18532944 | 0,08915542 | 0,03764281 | 0,16774366 | RAF1 |
| 5048,22821 | 0,18526246 | 0,07880271 | 0,01872476 | 0,10554062 | XPO6 |
| 2199,81948 | 0,18469158 | 0,09197263 | 0,04463113 | 0,1879514 | MAF1 |
| 3196,66589 | 0,18457615 | 0,0780512 | 0,01803939 | 0,10305657 | IPO4 |
| 2394,43351 | 0,18396716 | 0,08579774 | 0,03201719 | 0,15043387 | JADE2 |

| | | | | | |
|---|---|---|---|---|---|
| 1353,30742 | 0,18378749 | 0,08943656 | 0,03988401 | 0,17421138 | ACOT7 |
| 1371,46323 | 0,18372896 | 0,08091768 | 0,02317324 | 0,12159204 | AMBRA1 |
| 5865,04882 | 0,18321036 | 0,07510442 | 0,01471133 | 0,09028643 | VARS1 |
| 2030,25625 | 0,18287671 | 0,08339586 | 0,02831637 | 0,13890953 | MRPS16 |
| 3079,8021 | 0,18264474 | 0,09291979 | 0,04934285 | 0,20082085 | AGPAT3 |
| 1836,21229 | 0,18238459 | 0,08863942 | 0,03962842 | 0,17360717 | PEX19 |
| 937,141852 | 0,18180903 | 0,09103711 | 0,04581573 | 0,19153322 | ATG16L1 |
| 1654,41209 | 0,18157696 | 0,08822989 | 0,03959029 | 0,17348031 | SPRING1 |
| 3019,41428 | 0,18016016 | 0,0739945 | 0,01490094 | 0,09115444 | HAPSTR1 |
| 2153,76823 | 0,17991436 | 0,08090816 | 0,02616956 | 0,13158499 | RNF4 |
| 4294,65868 | 0,17978044 | 0,08277762 | 0,02986709 | 0,14339387 | KDM5C |
| 9659,21941 | 0,17955298 | 0,0902815 | 0,0467219 | 0,19403537 | TRIM28 |
| 1101,13039 | 0,17806393 | 0,0894991 | 0,04663929 | 0,19381991 | TJAP1 |
| 1239,98477 | 0,17651584 | 0,07744215 | 0,02264774 | 0,11943083 | SMCR8 |
| 1373,83537 | 0,17462592 | 0,08242173 | 0,03411685 | 0,15668659 | CEP68 |
| 2016,85355 | 0,17258894 | 0,07909944 | 0,02911516 | 0,14118091 | WDR5 |
| 1396,09152 | 0,17192662 | 0,08736879 | 0,049088 | 0,20025745 | APMAP |
| 3291,5676 | 0,17059752 | 0,08533673 | 0,04559646 | 0,19091184 | ABR |
| 2544,12438 | 0,17050228 | 0,07456316 | 0,02221435 | 0,11773606 | ABL1 |
| 4863,25228 | 0,16764364 | 0,08475376 | 0,04792777 | 0,19713654 | MTA1 |
| 3712,79433 | 0,16610492 | 0,07801576 | 0,03324434 | 0,15390655 | CCAR2 |
| 1926,7293 | 0,165682 | 0,08308443 | 0,04613673 | 0,19244191 | CDC42EP4 |
| 2781,63985 | 0,16354954 | 0,08181536 | 0,04560752 | 0,1909159 | GTPBP2 |
| 11636,8563 | 0,16340005 | 0,07935074 | 0,03947386 | 0,17323187 | ITGA3 |
| 2283,91053 | 0,15848531 | 0,08045509 | 0,04885435 | 0,19969174 | DMAC2 |
| 6287,72702 | 0,1560475 | 0,07312726 | 0,03284958 | 0,15263924 | TLN1 |
| 3605,21987 | 0,14830906 | 0,07304398 | 0,04231511 | 0,18114121 | ELAVL1 |
| 4522,9183 | 0,14569807 | 0,06704565 | 0,02977149 | 0,14322295 | DAZAP1 |
| 6375,80537 | 0,14269768 | 0,06759487 | 0,03476608 | 0,15850821 | PPP1R9B |
| 4493,63312 | -0,1264292 | 0,06435669 | 0,04947122 | 0,20112696 | SRSF11 |
| 33973,9324 | -0,1333474 | 0,06715586 | 0,04707332 | 0,19485328 | PRKDC |
| 5260,60602 | -0,1349879 | 0,06578395 | 0,04017082 | 0,17473983 | NSD1 |
| 3885,43315 | -0,1372873 | 0,06222842 | 0,02737119 | 0,13584218 | DCTN1 |
| 1959,38607 | -0,1438658 | 0,07331367 | 0,04972371 | 0,20176036 | DCAF5 |
| 5718,391 | -0,1447685 | 0,06507252 | 0,0260997 | 0,1314433 | FAM120A |
| 6157,8515 | -0,1496298 | 0,06787806 | 0,02749685 | 0,13612452 | MAP4 |
| 5123,05168 | -0,1500992 | 0,0646915 | 0,0203284 | 0,11162499 | SSRP1 |
| 1716,79757 | -0,1513049 | 0,07474445 | 0,0429397 | 0,18327591 | TTBK2 |
| 6917,83395 | -0,1527726 | 0,07676608 | 0,04657928 | 0,19364818 | RPS6KA3 |
| 3389,97117 | -0,1531191 | 0,06948151 | 0,02754273 | 0,13617353 | DZIP1 |
| 2954,38638 | -0,153913 | 0,07798058 | 0,0484119 | 0,19852657 | MPZL1 |
| 1968,14142 | -0,1549352 | 0,07309649 | 0,03403997 | 0,15644166 | RAB3GAP1 |
| 32045,9528 | -0,1554056 | 0,07879327 | 0,04857301 | 0,19892854 | PABPC1 |
| 25073,317 | -0,1557913 | 0,06933509 | 0,02464433 | 0,12653605 | SCD |
| 7371,33489 | -0,1564768 | 0,07065241 | 0,02677783 | 0,13375485 | SMARCA4 |
| 3646,29826 | -0,1566811 | 0,07989165 | 0,04985927 | 0,20201384 | RALGAPB |
| 9411,46642 | -0,1572825 | 0,07735358 | 0,04202298 | 0,18013522 | RAN |
| 6227,90218 | -0,1588695 | 0,06485551 | 0,01430184 | 0,08841893 | ACIN1 |

| | | | | | |
|---|---|---|---|---|---|
| 2724,22947 | -0,1596235 | 0,07515206 | 0,03366954 | 0,15515071 | ABCF1 |
| 1338,95044 | -0,1601961 | 0,07978964 | 0,04467193 | 0,18808136 | XRCC1 |
| 7449,3346 | -0,1621167 | 0,07579387 | 0,03244245 | 0,15152889 | PPP6R3 |
| 15687,8494 | -0,1633287 | 0,07046895 | 0,02046342 | 0,1119689 | EIF2S3 |
| 1255,27917 | -0,1638771 | 0,07897752 | 0,03798804 | 0,16880498 | RIPK1 |
| 1754,27981 | -0,1659113 | 0,08354172 | 0,04703667 | 0,19474416 | MAPK9 |
| 2095,49782 | -0,1659752 | 0,07096916 | 0,01935123 | 0,10784817 | RTN3 |
| 3572,2243 | -0,1666726 | 0,08128004 | 0,0403062 | 0,17515701 | PDCD6IP |
| 5913,93817 | -0,1688235 | 0,08507543 | 0,0472115 | 0,19520849 | TCP1 |
| 13931,2036 | -0,1688546 | 0,06635977 | 0,01094241 | 0,07354934 | EIF4G1 |
| 23971,3061 | -0,169245 | 0,08485968 | 0,04610724 | 0,19241805 | IL1RAP |
| 2440,05478 | -0,1695841 | 0,07054813 | 0,01622533 | 0,09617778 | PEAK1 |
| 2032,42734 | -0,1696561 | 0,08250034 | 0,03974122 | 0,17388053 | CNDP2 |
| 3756,19233 | -0,1700315 | 0,07829716 | 0,02988453 | 0,14344121 | FKBP4 |
| 1807,13762 | -0,1704587 | 0,08634406 | 0,04836129 | 0,19840503 | VIRMA |
| 1659,94919 | -0,1709291 | 0,08126142 | 0,03542672 | 0,1606685 | GDAP2 |
| 4624,04698 | -0,1712725 | 0,08160998 | 0,03584587 | 0,16205885 | HMBOX1 |
| 2573,85903 | -0,1716103 | 0,08388046 | 0,04076709 | 0,17647087 | ASPH |
| 1653,60972 | -0,1721468 | 0,08513134 | 0,0431627 | 0,18364774 | VPS8 |
| 3367,63294 | -0,1727094 | 0,08798782 | 0,04966032 | 0,20163582 | IMMT |
| 2136,32098 | -0,1727119 | 0,07727303 | 0,02541168 | 0,12925118 | KLC1 |
| 4030,34662 | -0,1728951 | 0,08247084 | 0,03604327 | 0,16247423 | YME1L1 |
| 4600,67991 | -0,1731962 | 0,07990312 | 0,03019089 | 0,14447805 | VPS13A |
| 6442,413 | -0,1741364 | 0,06775794 | 0,0101705 | 0,07005314 | IARS1 |
| 10465,6361 | -0,1743407 | 0,07554573 | 0,02101304 | 0,1139873 | GTF2I |
| 2893,62056 | -0,1750516 | 0,08677781 | 0,04367062 | 0,18510121 | SLC25A40 |
| 9777,89325 | -0,1753023 | 0,08015044 | 0,02873041 | 0,14017744 | USP34 |
| 7103,36139 | -0,1755341 | 0,06740019 | 0,00920473 | 0,06567523 | UHRF2 |
| 2036,25861 | -0,1756451 | 0,08425061 | 0,03708806 | 0,16597519 | SLC35B1 |
| 2348,53842 | -0,1758312 | 0,0866441 | 0,04242264 | 0,18143733 | PRR11 |
| 1455,89864 | -0,176075 | 0,08685723 | 0,04264435 | 0,18212339 | ERC1 |
| 1416,27622 | -0,1775694 | 0,08405939 | 0,03464973 | 0,15816828 | OSBP |
| 4116,90118 | -0,177625 | 0,08076453 | 0,02785694 | 0,13718928 | ZNF800 |
| 1284,71705 | -0,17764 | 0,09044883 | 0,04953198 | 0,2012875 | FBXO38 |
| 5825,77556 | -0,1776527 | 0,08429861 | 0,03508104 | 0,15942156 | CLPTM1 |
| 1553,25493 | -0,1776925 | 0,0784923 | 0,02358516 | 0,122905 | SART3 |
| 1201,79408 | -0,1777055 | 0,08433467 | 0,03510485 | 0,15947631 | SRPK2 |
| 4675,2602 | -0,1795655 | 0,08889387 | 0,04338351 | 0,18429714 | TUG1 |
| 2953,22318 | -0,1796456 | 0,07535596 | 0,01712748 | 0,09945292 | HADHA |
| 2726,0798 | -0,1805892 | 0,0716301 | 0,01169766 | 0,07716087 | GUSBP14 |
| 1767,54748 | -0,1807442 | 0,09018008 | 0,04504241 | 0,18908507 | GNAL |
| 2143,77882 | -0,1808525 | 0,08510493 | 0,0335822 | 0,15490218 | NOL10 |
| 17912,6281 | -0,1812673 | 0,09038357 | 0,04490604 | 0,18873104 | HMGA1 |
| 2994,91534 | -0,1815195 | 0,08543001 | 0,0336055 | 0,15497187 | PKP4 |
| 1656,42989 | -0,1821632 | 0,086894 | 0,03604814 | 0,16247423 | SPAG5 |
| 911,988224 | -0,1834395 | 0,0921793 | 0,04658773 | 0,19364818 | RHOT1 |
| 27925,4061 | -0,1834445 | 0,07665111 | 0,01670033 | 0,09805887 | HUWE1 |
| 4392,82983 | -0,1837056 | 0,07606273 | 0,01572722 | 0,09446815 | PRKAR1A |

| | | | | | |
|---|---|---|---|---|---|
| 2576,47506 | -0,1837745 | 0,08089157 | 0,02309481 | 0,12128139 | NCAPH |
| 4357,44871 | -0,1842338 | 0,09109805 | 0,04313834 | 0,18364774 | EIF3D |
| 1075,20621 | -0,1845648 | 0,09090972 | 0,04233636 | 0,1811912 | ZBTB21 |
| 12625,9046 | -0,1849003 | 0,08643145 | 0,0324137 | 0,151432 | PEX1 |
| 2833,56773 | -0,185014 | 0,08721991 | 0,03390194 | 0,15603469 | UTP18 |
| 1359,30152 | -0,1851509 | 0,08953955 | 0,03865776 | 0,17071739 | ORC2 |
| 786,528279 | -0,1852268 | 0,09124577 | 0,04235887 | 0,18124653 | CDK7 |
| 1542,26328 | -0,1860246 | 0,0826687 | 0,0244336 | 0,1257297 | OCIAD1 |
| 5010,94211 | -0,1860838 | 0,08005522 | 0,02010175 | 0,1108232 | EIF3L |
| 6363,13077 | -0,1874923 | 0,06593248 | 0,00445927 | 0,04003396 | CHST11 |
| 2617,72762 | -0,1875702 | 0,08629172 | 0,02972949 | 0,14305992 | AHI1 |
| 3857,13015 | -0,1877057 | 0,07469023 | 0,01196678 | 0,07849851 | CHD8 |
| 1097,05085 | -0,1891678 | 0,09070789 | 0,03702745 | 0,16582163 | UIMC1 |
| 806,163146 | -0,1895831 | 0,09586869 | 0,04798186 | 0,19723326 | ZNG1B |
| 1683,0488 | -0,1896174 | 0,09572393 | 0,04760498 | 0,19653806 | TSN |
| 3850,43065 | -0,1897126 | 0,09559872 | 0,04720374 | 0,19520849 | PGD |
| 2269,51145 | -0,1897735 | 0,07821753 | 0,01525669 | 0,09246376 | DDX23 |
| 1824,90923 | -0,1898608 | 0,09323609 | 0,04171572 | 0,17922414 | RDH11 |
| 12511,4023 | -0,1902804 | 0,07155003 | 0,00782791 | 0,05891485 | EIF4B |
| 1839,79694 | -0,1907392 | 0,08386181 | 0,02293917 | 0,12066499 | NFX1 |
| 1128,50426 | -0,1911782 | 0,08856009 | 0,03087036 | 0,146869 | RBFOX2 |
| 3242,31733 | -0,1912396 | 0,09339581 | 0,04059678 | 0,17605585 | SINHCAF |
| 799,763936 | -0,1912528 | 0,09300242 | 0,03974131 | 0,17388053 | SRSF9 |
| 875,252362 | -0,1913511 | 0,08614813 | 0,02633829 | 0,13215243 | SIX1 |
| 1690,67204 | -0,191774 | 0,08112731 | 0,01808542 | 0,10328833 | DOCK7 |
| 6135,01691 | -0,1919848 | 0,08476565 | 0,02351946 | 0,12276183 | NF1 |
| 4834,61307 | -0,1919885 | 0,070765 | 0,00666686 | 0,05297925 | TTC3 |
| 3215,3595 | -0,1920781 | 0,09391887 | 0,04084013 | 0,17674658 | ATP5MC3 |
| 933,107055 | -0,1923715 | 0,09624811 | 0,04564036 | 0,19096886 | UBE2C |
| 2610,71027 | -0,1926794 | 0,07601504 | 0,01125264 | 0,07520709 | MSMO1 |
| 847,904831 | -0,1927131 | 0,09278638 | 0,03780575 | 0,16823201 | KLHL7 |
| 2787,25652 | -0,192752 | 0,09257405 | 0,03732985 | 0,16674177 | BATF3 |
| 1533,75709 | -0,1928922 | 0,09297215 | 0,0380115 | 0,16886959 | ZMYND11 |
| 3078,95795 | -0,1929147 | 0,08648748 | 0,02571092 | 0,13021316 | KARS1 |
| 6592,17669 | -0,1936102 | 0,09585989 | 0,04341239 | 0,18437842 | IL17RA |
| 2480,02681 | -0,1936504 | 0,0817087 | 0,01778763 | 0,10190732 | CD164 |
| 4413,65151 | -0,193685 | 0,08785934 | 0,02748991 | 0,13612452 | ACACA |
| 2067,06728 | -0,1940759 | 0,08078445 | 0,01628823 | 0,09646173 | TRIM14 |
| 6552,83984 | -0,194306 | 0,08649286 | 0,02467214 | 0,12653605 | USP24 |
| 2158,85569 | -0,1943154 | 0,08986618 | 0,03059694 | 0,1459822 | TGS1 |
| 1836,56113 | -0,1945958 | 0,09127128 | 0,03300195 | 0,15308406 | HDAC1 |
| 1443,33202 | -0,1953669 | 0,08528118 | 0,02197177 | 0,11710643 | LUZP1 |
| 3989,95781 | -0,1954826 | 0,08858027 | 0,02732516 | 0,13584218 | RBMS1 |
| 3282,49469 | -0,1957272 | 0,09674996 | 0,04307094 | 0,18358758 | ANKRD36B |
| 1058,64526 | -0,1958896 | 0,09176477 | 0,03278614 | 0,15238185 | BLMH |
| 4662,13925 | -0,1968203 | 0,0913524 | 0,03119967 | 0,14777599 | IKZF1 |
| 12133,8601 | -0,1968748 | 0,0875223 | 0,02448554 | 0,12592846 | MYBPC2 |
| 2447,75754 | -0,1970133 | 0,09367785 | 0,03545755 | 0,16073126 | TTC17 |

| | | | | | |
|---|---|---|---|---|---|
| 6125,43517 | -0,1971173 | 0,09961661 | 0,04784311 | 0,19704526 | CKAP5 |
| 1911,05669 | -0,1973031 | 0,09155376 | 0,03115778 | 0,14776594 | SEC31A |
| 861,727325 | -0,1974319 | 0,09271594 | 0,0332189 | 0,15386407 | GAN |
| 5213,06112 | -0,1978462 | 0,07534344 | 0,00864138 | 0,06299274 | ZNF207 |
| 3080,32128 | -0,1978898 | 0,10088931 | 0,04982602 | 0,20196902 | CLIC4 |
| 4626,80649 | -0,1981831 | 0,09832522 | 0,04384311 | 0,1855829 | FMNL2 |
| 11001,0036 | -0,1983964 | 0,09752053 | 0,04191065 | 0,17970205 | YWHAZ |
| 902,460388 | -0,1984884 | 0,09882048 | 0,04458212 | 0,18783357 | HUS1 |
| 1786,34156 | -0,1985916 | 0,08698194 | 0,02242233 | 0,11844024 | WDR12 |
| 2489,55311 | -0,199398 | 0,08359475 | 0,01706551 | 0,09931121 | TSPAN3 |
| 4084,38673 | -0,199943 | 0,08733202 | 0,02205278 | 0,11733988 | PARP4 |
| 911,343492 | -0,2000965 | 0,09127463 | 0,02836174 | 0,13907698 | NR2C1 |
| 3380,73169 | -0,2001038 | 0,08725181 | 0,02182464 | 0,11663537 | PDIA3 |
| 5476,14807 | -0,2003421 | 0,07488288 | 0,00746388 | 0,05717898 | OSMR |
| 6654,34313 | -0,2005596 | 0,09104884 | 0,02761099 | 0,13645109 | MSN |
| 852,132491 | -0,2006117 | 0,09914639 | 0,04303309 | 0,18356243 | TMEM237 |
| 1201,62591 | -0,200665 | 0,09095116 | 0,02736343 | 0,13584218 | CSNK2A2 |
| 1514,77926 | -0,2007064 | 0,09307452 | 0,03105206 | 0,14751092 | GNL2 |
| 4294,56601 | -0,2015339 | 0,10204043 | 0,04826366 | 0,19804744 | ANLN |
| 1230,6338 | -0,2020097 | 0,0894682 | 0,0239523 | 0,12397567 | NAB1 |
| 1392,52994 | -0,202079 | 0,09534279 | 0,03404825 | 0,15644169 | SHOC2 |
| 3221,45945 | -0,2021342 | 0,08231007 | 0,01405851 | 0,08750034 | POLR1B |
| 1424,32287 | -0,2021896 | 0,08227033 | 0,01398591 | 0,0871058 | TSHZ3 |
| 1795,86423 | -0,2023263 | 0,10277296 | 0,04899067 | 0,19997776 | SMU1 |
| 1492,46753 | -0,2024957 | 0,10248257 | 0,04816573 | 0,19773134 | CUL4B |
| 1182,73155 | -0,2028113 | 0,09340523 | 0,02990809 | 0,1434815 | CHAMP1 |
| 654,348688 | -0,2029108 | 0,10292389 | 0,04867076 | 0,19919954 | VPS4B |
| 3916,33706 | -0,2029323 | 0,0994506 | 0,04129722 | 0,17819492 | LMNB1 |
| 983,621595 | -0,203218 | 0,09660376 | 0,0354112 | 0,1606685 | INVS |
| 9891,01004 | -0,2032191 | 0,0830923 | 0,01445698 | 0,08913016 | FNDC3B |
| 644,542736 | -0,2033455 | 0,0983698 | 0,03871963 | 0,17085056 | SNRNP48 |
| 1323,25556 | -0,2038939 | 0,10298207 | 0,04771507 | 0,19665846 | UBFD1 |
| 2037,02785 | -0,2041434 | 0,09112833 | 0,0250796 | 0,12787887 | RBBP6 |
| 2868,25946 | -0,2044734 | 0,08148838 | 0,01209935 | 0,07915819 | MSH6 |
| 1045,58473 | -0,2045739 | 0,0965621 | 0,03412653 | 0,15668722 | BTRC |
| 1728,1815 | -0,2048176 | 0,08085138 | 0,01130071 | 0,07541491 | TSR1 |
| 7977,71727 | -0,204819 | 0,06349055 | 0,00125544 | 0,01596465 | CHD4 |
| 2542,42237 | -0,2049468 | 0,08476047 | 0,01560812 | 0,09405133 | AFF1 |
| 2337,75135 | -0,2050497 | 0,09219982 | 0,02615038 | 0,13158499 | UBE2K |
| 7018,26538 | -0,2050925 | 0,08573055 | 0,01674354 | 0,09820352 | KMT2A |
| 855,970847 | -0,2053345 | 0,09646338 | 0,03328519 | 0,15398267 | MYH10 |
| 1048,33191 | -0,2054651 | 0,10038208 | 0,04067472 | 0,17625078 | HMG20A |
| 1495,56164 | -0,2056221 | 0,09916617 | 0,03812476 | 0,16919799 | SHPRH |
| 889,654493 | -0,205638 | 0,09388475 | 0,02850079 | 0,13952481 | MRRF |
| 1871,19451 | -0,2057487 | 0,08651128 | 0,01739323 | 0,10037398 | OXSR1 |
| 1915,417 | -0,2058901 | 0,08580041 | 0,01641119 | 0,09694541 | LTA4H |
| 863,536535 | -0,2060235 | 0,09569316 | 0,03132181 | 0,14819706 | GLOD4 |
| 3928,26494 | -0,2060827 | 0,07404958 | 0,00538527 | 0,04573218 | PDCD11 |

| | | | | | |
|---|---|---|---|---|---|
| 2105,45412 | -0,2061476 | 0,09188426 | 0,02486088 | 0,12716756 | KMT2E |
| 33698,058 | -0,2061793 | 0,08668016 | 0,01737751 | 0,10033566 | ACTN4 |
| 1108,27248 | -0,2063737 | 0,09202954 | 0,02493087 | 0,12731904 | BABAM2 |
| 1739,29988 | -0,2064454 | 0,08138027 | 0,01118712 | 0,07490143 | COX15 |
| 1352,65435 | -0,2065444 | 0,08823439 | 0,01923936 | 0,10738289 | RAB5B |
| 19690,526 | -0,206615 | 0,1039637 | 0,04688067 | 0,19431087 | EEF1G |
| 1564,31416 | -0,20666 | 0,10227175 | 0,04331132 | 0,18403179 | MXRA7 |
| 17597,4087 | -0,206747 | 0,10385229 | 0,0465051 | 0,19347473 | SLC2A3 |
| 2246,363 | -0,2069032 | 0,09512491 | 0,02962498 | 0,14281125 | SMARCAD1 |
| 10895,2009 | -0,2075159 | 0,07523157 | 0,00580918 | 0,04825505 | SLC26A4 |
| 1357,58381 | -0,207659 | 0,09090545 | 0,02235154 | 0,11819829 | PLEKHA7 |
| 6988,07201 | -0,2077309 | 0,06914705 | 0,00266289 | 0,02772956 | HDLBP |
| 691,608051 | -0,2077876 | 0,10319446 | 0,04405637 | 0,18616664 | MICB |
| 5762,52504 | -0,2079594 | 0,09910922 | 0,03588001 | 0,16214131 | UPK1B |
| 738,607425 | -0,2079851 | 0,10605659 | 0,04987005 | 0,20201384 | COA7 |
| 1863,24514 | -0,2079912 | 0,09034916 | 0,02133051 | 0,11509571 | NSF |
| 1217,69708 | -0,208589 | 0,09287564 | 0,02471068 | 0,12653605 | FAM120B |
| 1563,58122 | -0,2085939 | 0,09156202 | 0,02271643 | 0,11972633 | HEATR6 |
| 3465,80366 | -0,2086542 | 0,07167601 | 0,00360189 | 0,03433717 | PAK2 |
| 1813,01661 | -0,2087154 | 0,07752723 | 0,00709917 | 0,05532266 | UQCRC2 |
| 1249,14903 | -0,2087644 | 0,10135054 | 0,03941527 | 0,1731532 | ANAPC7 |
| 1358,18626 | -0,2087746 | 0,10646827 | 0,04988965 | 0,20201384 | DOCK4 |
| 1074,07427 | -0,208796 | 0,08845654 | 0,01825345 | 0,10393424 | DHX57 |
| 1027,61591 | -0,2093066 | 0,10438144 | 0,04494069 | 0,18883475 | IFT172 |
| 1172,25679 | -0,2093479 | 0,09796498 | 0,03260074 | 0,15200556 | DNAJC11 |
| 1755,03925 | -0,2095307 | 0,08871479 | 0,01818421 | 0,10375855 | RAB6A |
| 694,025806 | -0,209776 | 0,09861157 | 0,0333955 | 0,15435758 | SH3YL1 |
| 3653,36904 | -0,2097871 | 0,09777949 | 0,03191191 | 0,15002517 | ANKRD40 |
| 3704,30887 | -0,2097916 | 0,08930814 | 0,01882009 | 0,10578875 | CSNK2A1 |
| 1247,02388 | -0,2098471 | 0,08427661 | 0,01277503 | 0,0817991 | ROCK2 |
| 962,338846 | -0,2098479 | 0,0965353 | 0,02972055 | 0,14305332 | AGK |
| 3793,91099 | -0,2102322 | 0,08665575 | 0,01526367 | 0,09247641 | PTCD3 |
| 3101,39781 | -0,2102854 | 0,1024229 | 0,04006196 | 0,17469749 | NUP50 |
| 1572,5657 | -0,2106161 | 0,10578605 | 0,04648495 | 0,19343341 | DLG1 |
| 10010,6458 | -0,2106491 | 0,06255483 | 0,00075872 | 0,0109586 | SIPA1L1 |
| 4625,65717 | -0,2111885 | 0,07580437 | 0,00533683 | 0,04542427 | QKI |
| 801,023184 | -0,2115234 | 0,09890452 | 0,03246297 | 0,15153214 | RALGAPA2 |
| 1275,16797 | -0,2116081 | 0,10135034 | 0,03680795 | 0,16515133 | REPS1 |
| 1568,6361 | -0,2116493 | 0,07917842 | 0,00751611 | 0,05745948 | RERE |
| 1822,40913 | -0,2116867 | 0,08060004 | 0,00862974 | 0,06299274 | ADIPOR2 |
| 3374,73405 | -0,2117533 | 0,09881716 | 0,03212277 | 0,1507054 | TRPM7 |
| 3060,25341 | -0,2119067 | 0,10353628 | 0,04068847 | 0,17625148 | RBBP7 |
| 1764,96031 | -0,2119963 | 0,10631815 | 0,04615423 | 0,19244191 | ESD |
| 4351,92391 | -0,2125122 | 0,09390522 | 0,02363259 | 0,12307324 | PRDX1 |
| 17497,0434 | -0,212546 | 0,08438678 | 0,0117785 | 0,07751685 | DICER1 |
| 1281,81366 | -0,2127223 | 0,08877188 | 0,01656244 | 0,09768638 | SCAMP1 |
| 4848,2241 | -0,2128396 | 0,09228158 | 0,02108748 | 0,11429284 | KIF5B |
| 2111,80862 | -0,2128903 | 0,09058691 | 0,01876728 | 0,10567479 | CAMSAP2 |

| | | | | | |
|---|---|---|---|---|---|
| 1962,30103 | -0,2130718 | 0,10141967 | 0,03565045 | 0,16130616 | BCAP29 |
| 2038,29499 | -0,2135994 | 0,08505033 | 0,01202373 | 0,07874805 | FYCO1 |
| 883,473403 | -0,2138926 | 0,09347279 | 0,02212077 | 0,11747275 | SHQ1 |
| 2533,51976 | -0,2140015 | 0,08146098 | 0,00861303 | 0,06289743 | AURKA |
| 4439,93064 | -0,2140497 | 0,09265547 | 0,02087887 | 0,11355249 | MTRR |
| 832,158448 | -0,2145703 | 0,10547162 | 0,04191243 | 0,17970205 | PPP1R8 |
| 1076,37991 | -0,2145717 | 0,09832503 | 0,02908969 | 0,14112974 | COX11 |
| 3317,95313 | -0,2146616 | 0,09620046 | 0,02565471 | 0,13003289 | KAT6A |
| 4249,95378 | -0,2147479 | 0,09629765 | 0,02574456 | 0,13028608 | CAPNS1 |
| 642,297461 | -0,2149381 | 0,09493691 | 0,02357348 | 0,122905 | TAF11 |
| 2018,35522 | -0,2151256 | 0,08482851 | 0,01121249 | 0,07504472 | ZMYM4 |
| 1031,50514 | -0,2151894 | 0,10843229 | 0,0471945 | 0,19520849 | MED14 |
| 1467,06999 | -0,2153799 | 0,09805642 | 0,02805689 | 0,13788687 | RIMS2 |
| 955,294526 | -0,2154481 | 0,10892647 | 0,04793749 | 0,19713654 | XPNPEP1 |
| 1278,52742 | -0,2154804 | 0,08140603 | 0,00812133 | 0,06040236 | CRYBG3 |
| 928,061039 | -0,2157131 | 0,0868109 | 0,01296016 | 0,08245733 | SEC24D |
| 4146,74914 | -0,2159696 | 0,07683143 | 0,00493949 | 0,0430295 | NFAT5 |
| 3284,3697 | -0,2160991 | 0,07331748 | 0,00320413 | 0,03149373 | DDX18 |
| 2186,44793 | -0,2162036 | 0,10690984 | 0,04314552 | 0,18364774 | RNF138 |
| 1575,29986 | -0,2163293 | 0,09400084 | 0,02137158 | 0,11523842 | VDAC3 |
| 18744,68 | -0,2163417 | 0,0934344 | 0,0205889 | 0,11246034 | RPS4X |
| 8088,31307 | -0,2165959 | 0,07983369 | 0,00666593 | 0,05297925 | WNK1 |
| 12254,9692 | -0,2167554 | 0,09731493 | 0,02592324 | 0,13065893 | HMGCS1 |
| 2276,43492 | -0,2168802 | 0,09051557 | 0,01657265 | 0,09771612 | HAUS6 |
| 797,779582 | -0,2169323 | 0,10461381 | 0,03811195 | 0,16919665 | TMEM267 |
| 1783,05312 | -0,2170094 | 0,09758954 | 0,02616895 | 0,13158499 | SEC23IP |
| 1841,93282 | -0,2173467 | 0,09585517 | 0,02336284 | 0,12227028 | REEP5 |
| 1487,17485 | -0,2173832 | 0,0935658 | 0,02016202 | 0,11102606 | PRC1 |
| 11233,599 | -0,2179212 | 0,08077304 | 0,00697689 | 0,05466244 | NPEPPS |
| 6465,14063 | -0,2180773 | 0,0969814 | 0,02453475 | 0,126113 | IL1R2 |
| 970,857066 | -0,2182645 | 0,09951539 | 0,02828768 | 0,13884078 | MYO10 |
| 2185,62801 | -0,2184546 | 0,09666465 | 0,02382608 | 0,12371385 | POGK |
| 2622,44458 | -0,2188303 | 0,10071299 | 0,02979452 | 0,14322718 | RPS6KB1 |
| 6098,72051 | -0,2190739 | 0,07911113 | 0,00561956 | 0,04717289 | HMGCR |
| 763,95634 | -0,2191311 | 0,10036014 | 0,02900291 | 0,14085318 | ARL1 |
| 5047,87639 | -0,2193949 | 0,07790711 | 0,00486092 | 0,04261977 | AFDN |
| 757,694489 | -0,2194038 | 0,10641425 | 0,03922773 | 0,1726513 | PALB2 |
| 2983,90223 | -0,2194391 | 0,07471753 | 0,00331493 | 0,03228093 | SUPT6H |
| 12753,0922 | -0,2195742 | 0,06635225 | 0,00093556 | 0,01285532 | MACF1 |
| 2263,27508 | -0,2195989 | 0,10667807 | 0,03954027 | 0,17338157 | NSD3 |
| 5399,80286 | -0,219727 | 0,10288653 | 0,03271005 | 0,15221518 | BCL3 |
| 3129,69809 | -0,219749 | 0,08322713 | 0,00828196 | 0,06123603 | PPT1 |
| 1113,2343 | -0,2209549 | 0,09935469 | 0,02615518 | 0,13158499 | MTF2 |
| 3462,5851 | -0,2210438 | 0,08075914 | 0,00619882 | 0,0505479 | UGGT1 |
| 3639,50375 | -0,2210459 | 0,10768544 | 0,04010168 | 0,17470962 | ENTPD7 |
| 3620,39 | -0,2212699 | 0,09189427 | 0,0160457 | 0,09564473 | ASXL2 |
| 812,8223 | -0,221272 | 0,10486451 | 0,03485191 | 0,15874649 | ZCCHC7 |
| 5007,3478 | -0,2213026 | 0,10294311 | 0,03157449 | 0,1489085 | ATP2B4 |

| | | | | | |
|---|---|---|---|---|---|
| 1045,3916 | -0,221325 | 0,09479813 | 0,01955918 | 0,10871446 | CNTRL |
| 2073,78904 | -0,22145 | 0,09106835 | 0,01502858 | 0,09157995 | PCNT |
| 1784,06818 | -0,2214891 | 0,10621705 | 0,03704659 | 0,1658681 | DPY19L3 |
| 983,812602 | -0,2216407 | 0,09216441 | 0,01617952 | 0,09608685 | OSTC |
| 9793,02117 | -0,2219169 | 0,08074697 | 0,00599052 | 0,04930259 | MCTP2 |
| 643,052573 | -0,2219845 | 0,0940624 | 0,01827637 | 0,10400219 | RMDN1 |
| 510,887974 | -0,2220873 | 0,10811351 | 0,03995585 | 0,17435532 | EXD2 |
| 2361,47095 | -0,2221645 | 0,10415675 | 0,03292621 | 0,15284076 | LIN7C |
| 6528,18849 | -0,2222413 | 0,09121896 | 0,01483633 | 0,09090615 | CCT6A |
| 36091,9857 | -0,2225822 | 0,10478366 | 0,03365284 | 0,15511461 | RACK1 |
| 3658,55303 | -0,2230577 | 0,0999876 | 0,02569083 | 0,13015177 | BMP2K |
| 683,480325 | -0,2234489 | 0,11095615 | 0,04402546 | 0,18610493 | TJP2 |
| 933,360463 | -0,2234533 | 0,10319611 | 0,03036266 | 0,14518368 | MAPKAPK5 |
| 1677,77691 | -0,2235333 | 0,09553357 | 0,019292 | 0,10757795 | RFC1 |
| 1052,4397 | -0,2235741 | 0,09087216 | 0,01388151 | 0,08674109 | SDHD |
| 2224,80755 | -0,2237179 | 0,09067009 | 0,01361046 | 0,08552758 | SCCPDH |
| 5692,20562 | -0,2241656 | 0,08143323 | 0,00590965 | 0,04882814 | MAP4K4 |
| 1465,55692 | -0,2243296 | 0,09370817 | 0,01666953 | 0,09801249 | ASNS |
| 2053,74148 | -0,2244583 | 0,11278246 | 0,04657025 | 0,19364818 | UGGT2 |
| 3525,95208 | -0,2244817 | 0,10311239 | 0,02947618 | 0,14245689 | RAD23B |
| 3567,04588 | -0,2244853 | 0,09731459 | 0,02106595 | 0,11420886 | SRP72 |
| 892,987928 | -0,2247218 | 0,09557991 | 0,01871596 | 0,10554062 | IMP3 |
| 4709,23076 | -0,2248 | 0,09926475 | 0,0235341 | 0,12277577 | DPYD |
| 3611,63375 | -0,2248209 | 0,0936263 | 0,01633881 | 0,09665673 | EIF4A2 |
| 10257,8204 | -0,2249067 | 0,10945572 | 0,03990092 | 0,17421138 | RANBP2 |
| 1957,28505 | -0,2255011 | 0,09285706 | 0,01516245 | 0,09219342 | ALG13 |
| 932,932962 | -0,2256279 | 0,10409303 | 0,03019218 | 0,14447805 | TPD52 |
| 3450,58532 | -0,2256817 | 0,09356263 | 0,01586125 | 0,09479174 | HELZ |
| 1169,22571 | -0,2257182 | 0,10172459 | 0,02649252 | 0,13264488 | SNX2 |
| 713,360005 | -0,2258729 | 0,10786326 | 0,03625398 | 0,16320761 | UBTD2 |
| 683,920531 | -0,2259288 | 0,11357987 | 0,04668332 | 0,19391771 | ATF7 |
| 2874,51941 | -0,2259336 | 0,08031702 | 0,00490781 | 0,04285407 | SF3B2 |
| 5017,62149 | -0,2260463 | 0,09959084 | 0,02322274 | 0,12181797 | THRAP3 |
| 4275,50552 | -0,2261589 | 0,0832759 | 0,006612 | 0,05264734 | PPP5C |
| 3433,68323 | -0,2261677 | 0,08228334 | 0,00598421 | 0,04929354 | ADD2 |
| 7299,96234 | -0,2261744 | 0,08152449 | 0,00553185 | 0,04668519 | SLC43A3 |
| 4782,78711 | -0,2263808 | 0,08519787 | 0,00788118 | 0,05919801 | SMARCC1 |
| 3502,31118 | -0,2264664 | 0,09590294 | 0,01820546 | 0,10381722 | GART |
| 812,027887 | -0,2264738 | 0,10422655 | 0,02978786 | 0,14322718 | PCMT1 |
| 767,572067 | -0,2270054 | 0,10490538 | 0,03047152 | 0,14563064 | CTPS2 |
| 1024,12542 | -0,2271219 | 0,09985664 | 0,02293715 | 0,12066499 | RRP36 |
| 1052,26862 | -0,2273804 | 0,11472582 | 0,04748523 | 0,19608644 | ZGRF1 |
| 1749,81225 | -0,2274884 | 0,10329703 | 0,02764596 | 0,13646992 | PARP9 |
| 1612,48815 | -0,2277493 | 0,07851057 | 0,00372123 | 0,03509938 | MAP3K5 |
| 430,404553 | -0,2277633 | 0,11592185 | 0,04943735 | 0,20103246 | ZSCAN23 |
| 1186,65044 | -0,2278803 | 0,08138494 | 0,00510977 | 0,04408665 | RGMB |
| 660,25853 | -0,2279666 | 0,11231466 | 0,04238562 | 0,18132001 | WDR7 |
| 1032,88585 | -0,228328 | 0,11029297 | 0,03843401 | 0,16995254 | CEP250 |

| | | | | | |
|---|---|---|---|---|---|
| 1305,12635 | -0,2283639 | 0,1088812 | 0,03596102 | 0,16223259 | TMEM165 |
| 2649,66558 | -0,2284574 | 0,08321804 | 0,00604581 | 0,04962383 | EXOC2 |
| 5163,20212 | -0,2285645 | 0,09578282 | 0,01701986 | 0,09920913 | AQP9 |
| 874,477754 | -0,2287114 | 0,10389005 | 0,02770236 | 0,13664139 | FAM3C |
| 595,973626 | -0,2287721 | 0,1123611 | 0,04174593 | 0,17931255 | GLMN |
| 3207,58684 | -0,2291351 | 0,09208703 | 0,01283752 | 0,08197731 | THOC2 |
| 4337,75066 | -0,2293668 | 0,10371999 | 0,02700789 | 0,13461198 | NCOR1 |
| 6319,75552 | -0,2294904 | 0,08928185 | 0,01015801 | 0,06999261 | ZKSCAN1 |
| 1294,1017 | -0,2296098 | 0,11597172 | 0,0477173 | 0,19665846 | PSMD4 |
| 6349,94626 | -0,2297426 | 0,0904095 | 0,01104937 | 0,07413037 | POM121C |
| 1519,61615 | -0,2298909 | 0,11581536 | 0,04714696 | 0,19511539 | ARHGAP21 |
| 1957,73568 | -0,229941 | 0,08625983 | 0,00768331 | 0,05816328 | SPOP |
| 1269,09927 | -0,2301154 | 0,10611434 | 0,03011606 | 0,14429626 | OSBPL9 |
| 2542,25381 | -0,2303211 | 0,08640714 | 0,00768664 | 0,05816328 | PJA2 |
| 911,06593 | -0,2303961 | 0,10165482 | 0,0234235 | 0,12239605 | GGNBP2 |
| 3504,71663 | -0,230503 | 0,08267377 | 0,00530177 | 0,04517843 | TARDBP |
| 1940,65621 | -0,230649 | 0,1023756 | 0,02426114 | 0,12514858 | DCAF1 |
| 795,135447 | -0,2308839 | 0,09898609 | 0,01967504 | 0,10920251 | MALSU1 |
| 5999,37874 | -0,2310207 | 0,0862858 | 0,00741982 | 0,05693043 | PYGL |
| 586,290639 | -0,2311574 | 0,10954923 | 0,03485165 | 0,15874649 | WASHC2A |
| 1959,8098 | -0,23126 | 0,09736588 | 0,01754114 | 0,10084958 | NCOA1 |
| 427,034215 | -0,2313718 | 0,11672176 | 0,04745116 | 0,19603131 | CZIB |
| 978,738599 | -0,2313862 | 0,0919564 | 0,01186078 | 0,07800408 | PHF10 |
| 1889,58213 | -0,231483 | 0,11390697 | 0,04213226 | 0,18052187 | HBS1L |
| 3598,0872 | -0,2317464 | 0,09600368 | 0,01578138 | 0,09461327 | MORF4L2 |
| 12532,6668 | -0,2320545 | 0,08779223 | 0,00821207 | 0,0608858 | USP32 |
| 828,470902 | -0,2323887 | 0,1165446 | 0,04615324 | 0,19244191 | SNX4 |
| 640,078534 | -0,232423 | 0,10134567 | 0,02182676 | 0,11663537 | ERMP1 |
| 703,029246 | -0,2324496 | 0,11278079 | 0,03929592 | 0,17281218 | THAP6 |
| 1905,94336 | -0,2326812 | 0,09376831 | 0,01308499 | 0,0830809 | APPL1 |
| 888,828484 | -0,2327462 | 0,10446318 | 0,02587911 | 0,13064531 | SP2 |
| 1627,23862 | -0,2328252 | 0,10979891 | 0,0339665 | 0,15625581 | PCNP |
| 1130,095 | -0,2328323 | 0,09538382 | 0,0146465 | 0,0899469 | KDM1B |
| 852,106167 | -0,2329362 | 0,10341737 | 0,02429766 | 0,12530281 | PPP1R11 |
| 1951,58003 | -0,2331981 | 0,08361047 | 0,00528544 | 0,04508825 | ACP1 |
| 8474,5606 | -0,2333439 | 0,07755848 | 0,00262438 | 0,02742286 | CHSY1 |
| 548,993813 | -0,2334965 | 0,11695459 | 0,04588259 | 0,1916857 | GORAB |
| 613,81472 | -0,23406 | 0,10380365 | 0,02414375 | 0,12471305 | RCHY1 |
| 12086,279 | -0,2341158 | 0,11892463 | 0,0489983 | 0,19997776 | RPS8 |
| 818,18876 | -0,2346237 | 0,10660115 | 0,02773964 | 0,13675397 | MRM2 |
| 901,940234 | -0,2347046 | 0,10424194 | 0,02435154 | 0,12547805 | GALNT1 |
| 35773,832 | -0,2349924 | 0,0721549 | 0,00112685 | 0,01465806 | HSP90AB1 |
| 688,279336 | -0,235035 | 0,11526266 | 0,04143756 | 0,178445 | STXBP4 |
| 2874,49065 | -0,2352517 | 0,08068261 | 0,00354815 | 0,03395776 | HIVEP1 |
| 1672,58625 | -0,2357001 | 0,08171624 | 0,00392195 | 0,03641175 | NPEPPSP1 |
| 1690,9351 | -0,2357273 | 0,11130733 | 0,03419157 | 0,15687163 | CSTF1 |
| 3592,81124 | -0,2357409 | 0,08024922 | 0,00330753 | 0,03225309 | NRDC |
| 1701,03207 | -0,2359493 | 0,09855894 | 0,01666613 | 0,09801249 | RPE |

| | | | | | |
|---|---|---|---|---|---|
| 2380,22551 | -0,2359686 | 0,1135361 | 0,03767642 | 0,1678539 | ATP5PB |
| 1087,41275 | -0,23599 | 0,09946949 | 0,01766882 | 0,10142956 | N4BP1 |
| 1430,35054 | -0,236069 | 0,09514837 | 0,01309915 | 0,08311504 | SP100 |
| 3282,61646 | -0,2360991 | 0,07893062 | 0,00277862 | 0,02862357 | DKC1 |
| 1927,88164 | -0,2363259 | 0,09927051 | 0,0172833 | 0,10003588 | CFI |
| 1025,9732 | -0,2364087 | 0,10813705 | 0,02880131 | 0,14044843 | STRN3 |
| 791,997974 | -0,2366372 | 0,10946995 | 0,03064413 | 0,14612339 | DRG1 |
| 5131,19977 | -0,2366864 | 0,08713868 | 0,00660353 | 0,05263077 | MTMR7 |
| 1407,43479 | -0,2369034 | 0,10973811 | 0,03086517 | 0,146869 | TBCB |
| 1302,84471 | -0,2370446 | 0,11444026 | 0,03832709 | 0,1696679 | PSMD6 |
| 871,873157 | -0,2372224 | 0,10486015 | 0,0236805 | 0,12316072 | DTWD1 |
| 2136,32342 | -0,2372567 | 0,08105396 | 0,0034209 | 0,0329734 | CD274 |
| 2047,40468 | -0,2372859 | 0,09018238 | 0,00850896 | 0,06232986 | CPSF2 |
| 3009,43302 | -0,2373263 | 0,09397378 | 0,01155493 | 0,07660507 | SLK |
| 678,51917 | -0,2376398 | 0,10256825 | 0,02050949 | 0,11215609 | ATXN3 |
| 3026,39673 | -0,2377848 | 0,10210477 | 0,01986798 | 0,11001524 | MED1 |
| 1469,53521 | -0,2378776 | 0,11930236 | 0,04616245 | 0,19244191 | TRNT1 |
| 923,31219 | -0,2380365 | 0,10148656 | 0,01900171 | 0,10659099 | NCEH1 |
| 2518,31791 | -0,238081 | 0,112602 | 0,03448469 | 0,15770813 | RAPGEF4 |
| 9660,69612 | -0,2381533 | 0,08794443 | 0,00676905 | 0,05345461 | POM121 |
| 1390,1885 | -0,2384653 | 0,11699526 | 0,0415252 | 0,17868961 | SDAD1 |
| 3111,44435 | -0,2386077 | 0,07883676 | 0,00247317 | 0,02612779 | TMEM131 |
| 2731,25074 | -0,238794 | 0,09737601 | 0,01419511 | 0,08808974 | LAIR1 |
| 1141,08782 | -0,2387964 | 0,09284033 | 0,01010782 | 0,06985025 | TIMM17A |
| 2263,27844 | -0,2389319 | 0,08545342 | 0,00517318 | 0,04447145 | NCR3LG1 |
| 1146,39053 | -0,2389638 | 0,09953504 | 0,01635925 | 0,09667187 | STC2 |
| 1307,2363 | -0,2389866 | 0,08827422 | 0,00678288 | 0,05354144 | LIX1L |
| 7857,47257 | -0,2390167 | 0,07686614 | 0,00187392 | 0,02152791 | CDK5RAP2 |
| 630,310083 | -0,2390601 | 0,12083877 | 0,04789043 | 0,19709161 | ARMC1 |
| 7326,60834 | -0,2391562 | 0,10417722 | 0,02169496 | 0,11622038 | PTPN13 |
| 5432,67476 | -0,2393421 | 0,08879433 | 0,00702901 | 0,05486633 | TOP2A |
| 698,789918 | -0,2398828 | 0,10958848 | 0,02860106 | 0,13987092 | REXO4 |
| 581,04238 | -0,2402458 | 0,11598838 | 0,03833148 | 0,1696679 | WDR89 |
| 2018,37879 | -0,2403661 | 0,08400037 | 0,00421657 | 0,03850467 | YEATS2 |
| 834,272806 | -0,240413 | 0,10604116 | 0,02338005 | 0,12227028 | LYRM4 |
| 1765,53449 | -0,2404179 | 0,08642521 | 0,00540579 | 0,04584623 | FUBP3 |
| 473,128944 | -0,2408637 | 0,12277445 | 0,04978127 | 0,20191028 | PBX3 |
| 1701,99339 | -0,2409616 | 0,11052722 | 0,02924922 | 0,14163095 | TIFA |
| 3066,68801 | -0,2409662 | 0,0951374 | 0,01131481 | 0,07542243 | LRRFIP1 |
| 591,203838 | -0,2411369 | 0,10335156 | 0,01963914 | 0,10903523 | ARMCX6 |
| 1856,3385 | -0,2411397 | 0,12121229 | 0,0466571 | 0,19385133 | GABARAP |
| 465,133713 | -0,2411951 | 0,12117452 | 0,0465384 | 0,19357069 | LINC01355 |
| 937,805743 | -0,241316 | 0,09813236 | 0,01392908 | 0,08693106 | PPP1R7 |
| 594,436514 | -0,241435 | 0,11203156 | 0,03115698 | 0,14776594 | TIMMDC1 |
| 1564,53919 | -0,2414486 | 0,09267721 | 0,00918035 | 0,06559722 | CNOT2 |
| 2374,02539 | -0,2414799 | 0,09238299 | 0,00895152 | 0,06444891 | TAB3 |
| 1801,69095 | -0,2415279 | 0,11208857 | 0,03117784 | 0,14776594 | STRN |
| 2564,22734 | -0,2417823 | 0,08359606 | 0,0038247 | 0,0357369 | PDIA6 |

| | | | | | |
|---|---|---|---|---|---|
| 816,35104 | -0,2418565 | 0,11125147 | 0,02970798 | 0,14305163 | RAB23 |
| 5663,11165 | -0,2419325 | 0,11362289 | 0,03323291 | 0,1538913 | ACTR3 |
| 2426,94889 | -0,2419753 | 0,08115551 | 0,00286723 | 0,02927735 | TAF1 |
| 646,029741 | -0,2420049 | 0,10482695 | 0,02096505 | 0,1138249 | MTA3 |
| 6152,33547 | -0,2420061 | 0,0807873 | 0,0027391 | 0,02835143 | SRI |
| 1783,2835 | -0,2424202 | 0,07561356 | 0,00134575 | 0,01682212 | PTPN3 |
| 1744,127 | -0,24248 | 0,10054619 | 0,01588147 | 0,09488266 | SF3A3 |
| 1089,24896 | -0,2425347 | 0,11987064 | 0,04304184 | 0,18356243 | GTF2A1 |
| 3635,33172 | -0,2426135 | 0,11098518 | 0,02881564 | 0,14044843 | WAC |
| 424,131123 | -0,2432457 | 0,11389819 | 0,03270865 | 0,15221518 | SLC35B3 |
| 5823,3851 | -0,2432761 | 0,07103148 | 0,00061501 | 0,0092575 | PARP1 |
| 5378,41758 | -0,2433623 | 0,08208157 | 0,00302798 | 0,0302494 | PTPN11 |
| 8686,21879 | -0,2435841 | 0,07573614 | 0,00129891 | 0,01634092 | LUC7L3 |
| 860,851416 | -0,2437122 | 0,11323445 | 0,03137535 | 0,14822795 | HELB |
| 491,758077 | -0,2439603 | 0,11198819 | 0,02937279 | 0,14206607 | NCK1 |
| 1239,73533 | -0,2439925 | 0,10722994 | 0,02288109 | 0,12042646 | EXOC4 |
| 784,537552 | -0,2440194 | 0,10539667 | 0,02059937 | 0,11248507 | GLDC |
| 1953,53275 | -0,2441496 | 0,106944 | 0,02243233 | 0,11845999 | GOLGB1 |
| 4149,47195 | -0,2441581 | 0,1036101 | 0,01844765 | 0,10462449 | KPNA2 |
| 13697,649 | -0,244235 | 0,07755392 | 0,00163702 | 0,01954201 | HNRNPUL1 |
| 6979,99104 | -0,2442779 | 0,080378 | 0,00237275 | 0,02533569 | SAMD4B |
| 1334,84332 | -0,2443749 | 0,11400449 | 0,03206853 | 0,15051792 | AP3M1 |
| 2136,23784 | -0,2443822 | 0,09165234 | 0,00766673 | 0,05816328 | UBAP2 |
| 3829,47827 | -0,2446883 | 0,10586236 | 0,02081178 | 0,11324699 | HSPA4 |
| 2631,24951 | -0,2448851 | 0,08556703 | 0,00421098 | 0,03847217 | DNAJC13 |
| 2658,58159 | -0,245042 | 0,1113513 | 0,02776289 | 0,13683291 | DROSHA |
| 638,441526 | -0,2451557 | 0,119112 | 0,03957141 | 0,17343771 | ZNF197 |
| 3335,5366 | -0,2453389 | 0,12484242 | 0,04939242 | 0,20097937 | ITGB1 |
| 662,979624 | -0,2455341 | 0,10522048 | 0,01962085 | 0,10899769 | CFAP36 |
| 539,821188 | -0,2457459 | 0,11608085 | 0,03425785 | 0,15701806 | PIGK |
| 1578,0826 | -0,2460987 | 0,10356643 | 0,01749008 | 0,10073252 | USP53 |
| 645,822034 | -0,2461325 | 0,09770477 | 0,011764 | 0,07747536 | COMMD2 |
| 830,030723 | -0,2461927 | 0,08561905 | 0,00403461 | 0,03721095 | TMEM135 |
| 740,161401 | -0,2462019 | 0,11808359 | 0,03707111 | 0,16593859 | PSAT1 |
| 2717,07548 | -0,2463238 | 0,09361503 | 0,00850734 | 0,06232986 | MRPL19 |
| 560,709661 | -0,2465678 | 0,11282969 | 0,02886618 | 0,14055523 | NTPCR |
| 967,421636 | -0,2472116 | 0,1065591 | 0,02034367 | 0,11162499 | ENOPH1 |
| 2970,48306 | -0,2473635 | 0,10757532 | 0,0214797 | 0,11550604 | LARP4 |
| 1332,45448 | -0,2474586 | 0,09681429 | 0,01058793 | 0,07188166 | PARD3 |
| 1481,06084 | -0,2475487 | 0,09484821 | 0,00905565 | 0,06498384 | METAP1 |
| 1128,87513 | -0,2475724 | 0,10148508 | 0,01470779 | 0,09028643 | RAD17 |
| 2265,01001 | -0,2475727 | 0,10288069 | 0,01611035 | 0,09588672 | PPFIBP1 |
| 5987,8647 | -0,2476849 | 0,10295372 | 0,01613761 | 0,0959584 | FAM133B |
| 1703,72474 | -0,2478978 | 0,12043977 | 0,03956402 | 0,17343771 | NOL11 |
| 4571,81667 | -0,2480854 | 0,11874291 | 0,03668391 | 0,16471198 | TARS1 |
| 20240,8681 | -0,2481714 | 0,09100586 | 0,00639164 | 0,05150608 | CANX |
| 1019,61334 | -0,2482723 | 0,11915717 | 0,0371993 | 0,16619784 | ZNF431 |
| 822,161824 | -0,2484522 | 0,11353885 | 0,02865087 | 0,13995394 | ACTL6A |

| | | | | | |
|---|---|---|---|---|---|
| 2848,37089 | -0,2485149 | 0,10438448 | 0,01727674 | 0,1000285 | MAML2 |
| 5742,46658 | -0,2485686 | 0,08405531 | 0,00310444 | 0,03080182 | ANKRD36 |
| 1872,68854 | -0,2486819 | 0,12459646 | 0,04594497 | 0,19186155 | EIF4E |
| 499,310364 | -0,2487868 | 0,11767698 | 0,03450245 | 0,15770813 | KLHL20 |
| 1004,65161 | -0,2489361 | 0,11631903 | 0,0323455 | 0,15126278 | SKAP2 |
| 2485,38539 | -0,249586 | 0,09886538 | 0,01158617 | 0,0766919 | ZKSCAN8 |
| 556,505164 | -0,2496539 | 0,11384604 | 0,02831398 | 0,13890953 | MTMR9 |
| 964,125388 | -0,2499395 | 0,10295402 | 0,01519572 | 0,09233085 | EML5 |
| 1286,66615 | -0,2500041 | 0,09556233 | 0,00889312 | 0,06417492 | SPTLC1 |
| 2037,77323 | -0,2500856 | 0,11354456 | 0,02762772 | 0,13645109 | NUP107 |
| 2229,11566 | -0,2504223 | 0,12342815 | 0,04246935 | 0,18155502 | CHD9 |
| 953,327729 | -0,2504995 | 0,10382961 | 0,01583914 | 0,09477936 | INTS2 |
| 1511,56574 | -0,25061 | 0,10541455 | 0,01743632 | 0,10049118 | ZMYM3 |
| 1941,71312 | -0,2506513 | 0,11068662 | 0,02354257 | 0,12278084 | DHX36 |
| 702,345388 | -0,2506627 | 0,09817569 | 0,01067365 | 0,07229099 | YIPF5 |
| 1166,86021 | -0,2508934 | 0,11893076 | 0,03489494 | 0,15886602 | PSMC2 |
| 702,011562 | -0,2513357 | 0,11061267 | 0,02307355 | 0,12123703 | CHM |
| 2493,12541 | -0,2518898 | 0,10708614 | 0,0186619 | 0,10540352 | UCHL5 |
| 25731,8392 | -0,252155 | 0,08929308 | 0,00474428 | 0,04192738 | RPL4 |
| 3104,32404 | -0,2522448 | 0,1014996 | 0,01294859 | 0,08245733 | POLA1 |
| 675,957726 | -0,2523939 | 0,11709905 | 0,03113146 | 0,1477581 | IQCB1 |
| 7198,49342 | -0,2524149 | 0,09691216 | 0,00919885 | 0,06567523 | GTPBP10 |
| 335,569429 | -0,2526539 | 0,12763187 | 0,04775391 | 0,19676649 | RTL8A |
| 561,661037 | -0,2526919 | 0,11964741 | 0,03468889 | 0,15823251 | GGPS1 |
| 21209,0366 | -0,2530096 | 0,07412535 | 0,00064189 | 0,00958945 | ANKIB1 |
| 1660,66022 | -0,2530761 | 0,1028213 | 0,01384268 | 0,08661289 | RBAK |
| 1610,60115 | -0,2537507 | 0,09662126 | 0,00863337 | 0,06299274 | CUL1 |
| 2038,47429 | -0,2538231 | 0,08584584 | 0,00310919 | 0,03083277 | CTPS1 |
| 1312,30633 | -0,2539327 | 0,08691474 | 0,00348204 | 0,03340961 | LRP11 |
| 617,585212 | -0,2539452 | 0,11844508 | 0,0320336 | 0,15043931 | SHCBP1 |
| 787,273593 | -0,2540912 | 0,12610411 | 0,04391177 | 0,18574885 | ZNF322 |
| 420,549634 | -0,2541971 | 0,11661478 | 0,02927214 | 0,14165168 | SNAPIN |
| 1419,25961 | -0,2543465 | 0,11090361 | 0,02182493 | 0,11663537 | GABBR1 |
| 714,402177 | -0,2544277 | 0,12398449 | 0,04016059 | 0,17473983 | ALKAL2 |
| 3469,26672 | -0,2545272 | 0,07210531 | 0,00041566 | 0,00696608 | HNRNPM |
| 2302,83538 | -0,2546715 | 0,07809574 | 0,00111014 | 0,01452619 | FRAS1 |
| 2431,6204 | -0,2548721 | 0,09785001 | 0,00919487 | 0,06567523 | CSNK2B |
| 448,326381 | -0,2548839 | 0,11234048 | 0,02327736 | 0,12195493 | HARS2 |
| 608,564397 | -0,2553308 | 0,1284334 | 0,04680721 | 0,19413398 | PSMG1 |
| 1037,66604 | -0,255419 | 0,12149042 | 0,03552002 | 0,16089879 | MRPS10 |
| 993,37731 | -0,2555048 | 0,11013217 | 0,02034179 | 0,11162499 | NHP2 |
| 2738,31113 | -0,2558139 | 0,08694154 | 0,00325714 | 0,0318493 | TP53BP1 |
| 1501,47528 | -0,2559922 | 0,08806001 | 0,0036489 | 0,03464163 | SPTLC2 |
| 5129,12181 | -0,2560647 | 0,08243779 | 0,00189536 | 0,02169298 | CTNNA1 |
| 1177,46724 | -0,2561308 | 0,09478318 | 0,00688655 | 0,05415645 | KIF5A |
| 772,841316 | -0,2561606 | 0,11078744 | 0,02076773 | 0,11319505 | MRPS11 |
| 583,478966 | -0,2562612 | 0,11964724 | 0,0322091 | 0,15082232 | ZW10 |
| 1906,4662 | -0,2562802 | 0,10009955 | 0,0104596 | 0,07139472 | SCAF4 |

| | | | | | |
|---|---|---|---|---|---|
| 746,368922 | -0,2566245 | 0,11749323 | 0,02895017 | 0,14070544 | GADD45B |
| 1179,29908 | -0,2567405 | 0,09591132 | 0,00743173 | 0,05699865 | PCYOX1 |
| 1474,81847 | -0,2568109 | 0,08360948 | 0,00212948 | 0,02342557 | LAS1L |
| 2031,08068 | -0,2571757 | 0,08531665 | 0,00257516 | 0,02699426 | REST |
| 4945,57694 | -0,2572193 | 0,12439475 | 0,03866202 | 0,17071739 | HNRNPDL |
| 1974,82337 | -0,2572568 | 0,09470129 | 0,00659746 | 0,05260788 | THOC3 |
| 22716,5724 | -0,2572701 | 0,09960979 | 0,00980079 | 0,06858965 | CEMIP |
| 9392,408 | -0,257285 | 0,10329595 | 0,01274716 | 0,08173473 | ODC1 |
| 516,547384 | -0,2573489 | 0,1184289 | 0,0297785 | 0,14322295 | DHRS7 |
| 2007,37559 | -0,2574043 | 0,09226364 | 0,00527279 | 0,04504336 | USP48 |
| 5215,62531 | -0,2574068 | 0,10460257 | 0,01386248 | 0,0866964 | ZNF638 |
| 2000,09916 | -0,2575327 | 0,10379335 | 0,01309387 | 0,08310938 | CPNE8 |
| 1571,03498 | -0,2578695 | 0,10804567 | 0,01700166 | 0,09916415 | PSMA3 |
| 2293,21857 | -0,2579318 | 0,0934576 | 0,00578226 | 0,04811177 | QRICH1 |
| 1111,7423 | -0,2579475 | 0,11859009 | 0,02962122 | 0,14281125 | SRP54 |
| 4554,07128 | -0,2581487 | 0,11281867 | 0,02212742 | 0,11747275 | XRN2 |
| 643,299648 | -0,2584127 | 0,12006223 | 0,03137193 | 0,14822795 | SLC25A4 |
| 2275,73224 | -0,2584798 | 0,10129025 | 0,01071458 | 0,07250737 | AQR |
| 1145,30332 | -0,2586552 | 0,11412012 | 0,02341975 | 0,12239605 | LEF1 |
| 941,390355 | -0,2588452 | 0,13184197 | 0,04961144 | 0,20152385 | RFC4 |
| 613,322515 | -0,2588691 | 0,11334045 | 0,02237181 | 0,11824508 | PHTF1 |
| 930,315819 | -0,2588817 | 0,10751845 | 0,01604947 | 0,09564473 | CAPRIN2 |
| 3220,75713 | -0,2588879 | 0,10006915 | 0,00967903 | 0,06808066 | STT3A |
| 5364,05461 | -0,2592566 | 0,10857476 | 0,01694881 | 0,09891684 | HNRNPAB |
| 642,574153 | -0,2593016 | 0,11916474 | 0,02955582 | 0,1425657 | ELMOD2 |
| 924,65566 | -0,2595232 | 0,1237422 | 0,03596795 | 0,16223259 | GPR135 |
| 2181,44482 | -0,2595464 | 0,09128239 | 0,00446442 | 0,04003396 | HACD3 |
| 2187,14103 | -0,2595692 | 0,08930229 | 0,00365339 | 0,03464953 | ATP11C |
| 391,3422 | -0,2596794 | 0,12440005 | 0,03684707 | 0,16525138 | MED20 |
| 434,009081 | -0,259805 | 0,12026156 | 0,03074688 | 0,14646568 | ZNF277 |
| 935,106215 | -0,2598905 | 0,10166101 | 0,01057487 | 0,07181875 | BCLAF3 |
| 3505,93625 | -0,2600024 | 0,09480308 | 0,00609637 | 0,04997807 | ITPR2 |
| 1104,53931 | -0,2600305 | 0,0951907 | 0,00630124 | 0,05110408 | STK26 |
| 1193,15559 | -0,260035 | 0,10285275 | 0,01146405 | 0,07613592 | ZNF512 |
| 648,516753 | -0,2601334 | 0,12413336 | 0,03611805 | 0,16275055 | NUP35 |
| 341,116233 | -0,2604427 | 0,12054891 | 0,03073606 | 0,146451 | BORA |
| 522,102897 | -0,2607721 | 0,11549582 | 0,02395506 | 0,12397567 | ANAPC10 |
| 1389,37808 | -0,2610689 | 0,08466587 | 0,00204568 | 0,02280864 | TFG |
| 13031,8841 | -0,26112 | 0,11287229 | 0,02070006 | 0,11287201 | RPL32 |
| 913,966281 | -0,2611616 | 0,09587064 | 0,00644762 | 0,05171486 | MKRN2 |
| 1243,28925 | -0,2611891 | 0,09560993 | 0,00629855 | 0,05110408 | PTCH1 |
| 1889,47149 | -0,2612471 | 0,10555917 | 0,0133278 | 0,08414255 | DNAJC7 |
| 2032,40852 | -0,2614319 | 0,08564809 | 0,00227022 | 0,02460191 | BBX |
| 545,689065 | -0,261641 | 0,12046798 | 0,02986532 | 0,14339387 | MBD5 |
| 863,209765 | -0,2616941 | 0,11409328 | 0,02180855 | 0,11663537 | NAA20 |
| 857,559284 | -0,2617553 | 0,09151486 | 0,00423309 | 0,03859963 | LAP3 |
| 718,524387 | -0,2618781 | 0,12998933 | 0,04394528 | 0,18582767 | SRFBP1 |
| 7374,1689 | -0,261957 | 0,10903091 | 0,01627927 | 0,09646173 | CCT7 |

| | | | | | |
|---|---|---|---|---|---|
| 2114,42532 | -0,2620202 | 0,11992967 | 0,02890484 | 0,14059345 | GORASP2 |
| 2942,1469 | -0,2621745 | 0,08567378 | 0,00221227 | 0,02412564 | ARCN1 |
| 688,819315 | -0,2623774 | 0,11160179 | 0,0187223 | 0,10554062 | SNRNP27 |
| 1019,61812 | -0,262625 | 0,11283137 | 0,01993397 | 0,11018718 | DSTN |
| 591,16918 | -0,2630647 | 0,11460532 | 0,02171041 | 0,11627022 | BICD1 |
| 2015,62901 | -0,2631343 | 0,08047617 | 0,00107655 | 0,01425433 | ELP1 |
| 984,718523 | -0,2632834 | 0,11932464 | 0,02735277 | 0,13584218 | AMER1 |
| 1081,52394 | -0,2633117 | 0,09885871 | 0,00773289 | 0,05845542 | CLCN5 |
| 5489,66943 | -0,2633525 | 0,10294741 | 0,01052377 | 0,0715745 | HIF1A |
| 454,143003 | -0,2634221 | 0,12375173 | 0,03328454 | 0,15398267 | WDR44 |
| 733,080358 | -0,2636575 | 0,11524164 | 0,0221453 | 0,11747716 | CTSZ |
| 904,428325 | -0,2636726 | 0,13302474 | 0,04746446 | 0,19604346 | TBK1 |
| 1632,43482 | -0,2640589 | 0,12160697 | 0,02989998 | 0,14347896 | IL13RA1 |
| 3058,41279 | -0,2641994 | 0,08946214 | 0,003145 | 0,03102533 | GALNT3 |
| 795,408984 | -0,2642391 | 0,11355402 | 0,01996579 | 0,11029861 | PTTG1 |
| 1025,0265 | -0,2644007 | 0,12623577 | 0,03621557 | 0,1630735 | MCM8 |
| 3073,44083 | -0,2644783 | 0,08986321 | 0,00324921 | 0,03178818 | SPECC1 |
| 774,877162 | -0,264946 | 0,11601808 | 0,02239144 | 0,11831007 | MRPL39 |
| 2288,61747 | -0,2649878 | 0,11340849 | 0,01946112 | 0,1082693 | FARSB |
| 1734,18116 | -0,2652801 | 0,08595298 | 0,00202637 | 0,02267358 | RBM17 |
| 606,713202 | -0,2653361 | 0,1119087 | 0,01773978 | 0,10174141 | MRPL57 |
| 808,262488 | -0,2655796 | 0,10863076 | 0,01449357 | 0,08923944 | TMEM230 |
| 370,466832 | -0,2655987 | 0,11761654 | 0,02393478 | 0,12397567 | TFB1M |
| 1701,9167 | -0,2656963 | 0,10152029 | 0,0088661 | 0,06411213 | TMEM147 |
| 1304,81382 | -0,2657269 | 0,10130513 | 0,00871502 | 0,06334882 | C11orf24 |
| 1186,45602 | -0,2657445 | 0,09782624 | 0,00659771 | 0,05260788 | TMEM68 |
| 3315,22624 | -0,2660075 | 0,11017061 | 0,01575661 | 0,09452465 | PSMD1 |
| 1505,8421 | -0,2660658 | 0,08473651 | 0,00168994 | 0,01997215 | PSMD7 |
| 951,173383 | -0,266198 | 0,12939854 | 0,03966752 | 0,17373825 | PPWD1 |
| 637,450212 | -0,2662669 | 0,12388322 | 0,03160761 | 0,14895333 | MRPL52 |
| 11130,4459 | -0,2664515 | 0,10365997 | 0,010157 | 0,06999261 | DDX21 |
| 2041,12707 | -0,2664842 | 0,12577882 | 0,03411811 | 0,15668659 | DIS3 |
| 697,659542 | -0,2664927 | 0,11861962 | 0,0246646 | 0,12653605 | ISOC1 |
| 763,026417 | -0,2664952 | 0,12567256 | 0,03395955 | 0,15625581 | SLC4A1AP |
| 765,742418 | -0,2666667 | 0,12196933 | 0,02879056 | 0,14043471 | LAMTOR5 |
| 2294,80565 | -0,2667322 | 0,08361947 | 0,00142355 | 0,01755862 | NDUFS1 |
| 910,035893 | -0,2668623 | 0,10930164 | 0,01462552 | 0,08987642 | PPP2CB |
| 11409,8805 | -0,2669116 | 0,09097332 | 0,00334673 | 0,0324736 | HNRNPH1 |
| 1271,14103 | -0,2671962 | 0,1245438 | 0,03192093 | 0,15002517 | AFG2A |
| 1185,0232 | -0,2672676 | 0,09071798 | 0,0032177 | 0,03156152 | PITRM1 |
| 455,680274 | -0,2673125 | 0,10930694 | 0,01446428 | 0,08913428 | KIF16B |
| 841,526561 | -0,2674435 | 0,10770405 | 0,01302325 | 0,08277224 | ODR4 |
| 2244,54056 | -0,267574 | 0,09500585 | 0,00485659 | 0,04260154 | DARS1 |
| 1852,82219 | -0,2677891 | 0,0950725 | 0,00485224 | 0,04258316 | SEC24B |
| 743,307946 | -0,2677948 | 0,09654737 | 0,00554204 | 0,04672949 | FBXW7 |
| 518,926692 | -0,2680361 | 0,13062202 | 0,04016988 | 0,17473983 | GPATCH2 |
| 470,289193 | -0,2681675 | 0,12449471 | 0,03123647 | 0,14790423 | ALG5 |
| 4998,60061 | -0,2681795 | 0,10153981 | 0,00826308 | 0,06116812 | U2SURP |

| | | | | | |
|---|---|---|---|---|---|
| 802,211005 | -0,2682492 | 0,11244789 | 0,01705395 | 0,09931121 | BRK1 |
| 2094,06457 | -0,2683659 | 0,13028471 | 0,03941365 | 0,1731532 | ID2 |
| 5869,3533 | -0,2685005 | 0,10576681 | 0,01112944 | 0,07454161 | TXN |
| 1053,0733 | -0,2685012 | 0,13352885 | 0,04434548 | 0,18708159 | UTP11 |
| 4248,82753 | -0,2685359 | 0,09854473 | 0,00642985 | 0,05170547 | CBX5 |
| 1067,22081 | -0,2686418 | 0,08741671 | 0,00211835 | 0,02335985 | GPHN |
| 561,72571 | -0,2686649 | 0,1307111 | 0,03983945 | 0,17421138 | SCP2 |
| 4674,02776 | -0,268717 | 0,11743134 | 0,02212059 | 0,11747275 | UBXN2B |
| 4062,86619 | -0,268814 | 0,07624021 | 0,00042207 | 0,00703619 | TRPS1 |
| 956,397344 | -0,2691301 | 0,09626058 | 0,00517635 | 0,04447854 | CLPX |
| 1404,54627 | -0,2692461 | 0,12919782 | 0,03716168 | 0,16614748 | NLRP7 |
| 3542,10597 | -0,2700489 | 0,09754877 | 0,00563413 | 0,04725327 | LSM14A |
| 1881,75449 | -0,2701202 | 0,11323218 | 0,01705393 | 0,09931121 | GFM1 |
| 1030,74579 | -0,2706661 | 0,1101523 | 0,0140025 | 0,08718042 | FAM111A |
| 904,386532 | -0,2707252 | 0,11798043 | 0,02175249 | 0,11644095 | EXOSC9 |
| 930,168781 | -0,2710797 | 0,11117461 | 0,01475554 | 0,09049907 | L3HYPDH |
| 465,631396 | -0,2712116 | 0,13167321 | 0,03942413 | 0,1731532 | SGO1 |
| 1213,38262 | -0,2714564 | 0,08624484 | 0,00164668 | 0,01962019 | CTNNBL1 |
| 1050,89714 | -0,2715107 | 0,11501646 | 0,01824417 | 0,10391266 | PRKACA |
| 802,723158 | -0,2717549 | 0,10803595 | 0,01188935 | 0,07816483 | CEP43 |
| 639,305215 | -0,2717914 | 0,12408207 | 0,02849406 | 0,13952481 | AIG1 |
| 1247,8235 | -0,2718347 | 0,11685776 | 0,02000781 | 0,11049845 | ATP5PO |
| 832,497367 | -0,2719808 | 0,11481171 | 0,01783966 | 0,10216228 | PRPF38A |
| 332,033179 | -0,2721048 | 0,13418805 | 0,04258202 | 0,18195445 | C6orf163 |
| 1108,86689 | -0,2721434 | 0,11156706 | 0,01471652 | 0,09028898 | RNF6 |
| 3023,65228 | -0,2721728 | 0,0953952 | 0,00432939 | 0,03915695 | DNAJC10 |
| 735,703703 | -0,2722033 | 0,13246872 | 0,03989351 | 0,17421138 | SUGT1 |
| 2735,66292 | -0,2723809 | 0,09350881 | 0,003581 | 0,03420299 | SLF2 |
| 432,997356 | -0,2725835 | 0,12525406 | 0,02953697 | 0,1425657 | UTP23 |
| 845,022347 | -0,273011 | 0,12797717 | 0,03290184 | 0,152807 | BOLA2B |
| 647,982657 | -0,2732408 | 0,12694524 | 0,03136348 | 0,14822795 | PLEKHA5 |
| 579,653692 | -0,2735522 | 0,10938972 | 0,0123944 | 0,08028565 | AZI2 |
| 4740,33658 | -0,2735941 | 0,11465146 | 0,01701834 | 0,09920913 | DEK |
| 1428,60587 | -0,27375 | 0,13424481 | 0,04143155 | 0,178445 | HIGD2A |
| 614,225458 | -0,2739924 | 0,13454797 | 0,04171103 | 0,17922414 | MED10 |
| 1973,40556 | -0,2740841 | 0,08407599 | 0,00111429 | 0,01455174 | ARGLU1 |
| 4342,31614 | -0,2741228 | 0,10474783 | 0,00887119 | 0,06411213 | ZFR |
| 509,35541 | -0,2741336 | 0,13313616 | 0,03948972 | 0,17323187 | TRAPPC6B |
| 1636,07434 | -0,2741666 | 0,11122939 | 0,0137061 | 0,08592879 | PUS7 |
| 1641,18922 | -0,2742503 | 0,12551388 | 0,0288874 | 0,14055523 | RSL24D1 |
| 2406,60404 | -0,2742683 | 0,08822229 | 0,00187828 | 0,0215649 | THUMPD2 |
| 2107,10441 | -0,2744023 | 0,10404339 | 0,00835484 | 0,06158234 | ADAM9 |
| 576,821887 | -0,2746641 | 0,11278873 | 0,0148832 | 0,09110483 | RPA3 |
| 2232,30664 | -0,2748714 | 0,11555827 | 0,01737669 | 0,10033566 | LOC653513 |
| 873,523551 | -0,2759115 | 0,10252275 | 0,00711906 | 0,05541587 | ITSN1 |
| 625,612053 | -0,2760789 | 0,10726211 | 0,01005674 | 0,06967546 | ERCC8 |
| 644,135444 | -0,2760884 | 0,12645888 | 0,02901909 | 0,14085947 | GOPC |
| 2708,13246 | -0,2768133 | 0,09087465 | 0,0023183 | 0,02495141 | NUP214 |

| | | | | | |
|---|---|---|---|---|---|
| 436,323936 | -0,2769661 | 0,11967301 | 0,02064812 | 0,1126862 | ALKBH1 |
| 5421,84097 | -0,2769871 | 0,13480821 | 0,0399104 | 0,17421138 | CCT2 |
| 2907,58502 | -0,2771237 | 0,09460433 | 0,00339735 | 0,03285787 | DDX1 |
| 628,621774 | -0,2773663 | 0,13775182 | 0,04405973 | 0,18616664 | RTCA |
| 10355,6426 | -0,2775984 | 0,09868901 | 0,00491029 | 0,04285407 | RPL10A |
| 969,406376 | -0,2776824 | 0,09925239 | 0,00514615 | 0,04429947 | MYO6 |
| 988,20265 | -0,2777884 | 0,13407191 | 0,03827139 | 0,16958619 | FAM171B |
| 820,002884 | -0,2778061 | 0,10826084 | 0,01028547 | 0,07058812 | CEP112 |
| 85848,1338 | -0,2779134 | 0,1062009 | 0,00887415 | 0,06411213 | GAPDH |
| 1850,95106 | -0,2779809 | 0,11307598 | 0,0139575 | 0,08704173 | LCLAT1 |
| 607,590103 | -0,2780888 | 0,11015221 | 0,01158345 | 0,0766919 | ZNG1A |
| 2995,21435 | -0,2781703 | 0,10239918 | 0,00659705 | 0,05260788 | SMS |
| 2557,92638 | -0,2783745 | 0,0807568 | 0,0005667 | 0,00876737 | CSGALNACT2 |
| 519,433634 | -0,2785543 | 0,13572851 | 0,04014148 | 0,17473983 | GTF2E2 |
| 1836,90736 | -0,2787868 | 0,11842559 | 0,01856713 | 0,10499361 | MTUS1 |
| 1279,17012 | -0,2792172 | 0,08289529 | 0,0007563 | 0,01093622 | BECN1 |
| 419,229424 | -0,279224 | 0,12429553 | 0,02467503 | 0,12653605 | NAPEPLD |
| 1743,15806 | -0,2793115 | 0,11143811 | 0,01219582 | 0,07949617 | CEP295 |
| 2230,84652 | -0,279479 | 0,08546902 | 0,00107568 | 0,01425279 | PPP1R12A |
| 10430,1807 | -0,2795218 | 0,10512024 | 0,00783573 | 0,05895025 | RPL29 |
| 1451,03059 | -0,2795234 | 0,0853804 | 0,00106089 | 0,01412609 | COPS8 |
| 1428,95332 | -0,279754 | 0,09134139 | 0,00219328 | 0,02397403 | SYAP1 |
| 5732,30995 | -0,2800909 | 0,11568863 | 0,01547465 | 0,09339582 | CALM2 |
| 1502,50761 | -0,2801383 | 0,11758132 | 0,01719522 | 0,09967855 | PRELID3B |
| 4355,86053 | -0,2802469 | 0,10931473 | 0,0103572 | 0,07092602 | CD2AP |
| 355,915533 | -0,2802828 | 0,12295109 | 0,02262973 | 0,11936917 | ZNF397 |
| 2157,27982 | -0,2803035 | 0,10836549 | 0,00969144 | 0,06811682 | API5 |
| 3947,4554 | -0,2805319 | 0,12998821 | 0,03091747 | 0,1470192 | CHD1 |
| 7415,49246 | -0,2811339 | 0,08668677 | 0,00118237 | 0,01521873 | RUNX2 |
| 1386,94049 | -0,2812059 | 0,10991459 | 0,0105154 | 0,07156625 | TMX1 |
| 1127,71052 | -0,2812584 | 0,10530208 | 0,0075633 | 0,05763398 | PTRH2 |
| 27527,5511 | -0,2812673 | 0,0854431 | 0,00099525 | 0,01341246 | PPIA |
| 386,907404 | -0,2814264 | 0,13158176 | 0,0324523 | 0,15153214 | BCKDHB |
| 5620,13222 | -0,2814493 | 0,08677875 | 0,00118155 | 0,01521873 | ZEB2 |
| 3238,14111 | -0,2815063 | 0,0986293 | 0,00431473 | 0,03906177 | NOP58 |
| 571,414401 | -0,2815567 | 0,10561412 | 0,00767824 | 0,05816328 | TULP3 |
| 2066,89714 | -0,2820512 | 0,09264717 | 0,00233177 | 0,02503937 | APC |
| 1268,10529 | -0,2824229 | 0,09915654 | 0,004396 | 0,03960794 | PPHLN1 |
| 2635,81566 | -0,2824953 | 0,12106726 | 0,0196285 | 0,10900818 | RAB5C |
| 304,873456 | -0,2828788 | 0,14388301 | 0,04929479 | 0,20070551 | FBXO4 |
| 673,46293 | -0,2832803 | 0,1350538 | 0,03594612 | 0,16223259 | CHMP2B |
| 3070,38883 | -0,2835008 | 0,0917674 | 0,00200601 | 0,0225123 | PLOD2 |
| 1175,52569 | -0,2835714 | 0,13137358 | 0,03088806 | 0,14691628 | ZNF22 |
| 938,1682 | -0,2835984 | 0,11320611 | 0,01223991 | 0,07969421 | RBM41 |
| 1349,72663 | -0,2839803 | 0,11255247 | 0,01163277 | 0,07690552 | SEC62 |
| 889,030995 | -0,2842229 | 0,10533173 | 0,0069682 | 0,05463009 | CRACD |
| 1991,40336 | -0,284449 | 0,09049619 | 0,00167103 | 0,0198306 | ETV1 |
| 5287,28816 | -0,2844987 | 0,13799874 | 0,03924551 | 0,17268939 | LOC100132287 |

| | | | | | |
|---|---|---|---|---|---|
| 1727,15446 | -0,2847449 | 0,10386598 | 0,0061166 | 0,05007985 | ZNF664 |
| 2068,36573 | -0,2848687 | 0,08889457 | 0,00135265 | 0,0168823 | KIDINS220 |
| 3848,059 | -0,2849032 | 0,12656262 | 0,02438019 | 0,12552313 | H3C2 |
| 1064,36142 | -0,2855365 | 0,14449764 | 0,04814762 | 0,19769989 | METTL5 |
| 512,542569 | -0,2856347 | 0,1172125 | 0,01481391 | 0,0907982 | NDUFB11 |
| 402,404287 | -0,2857675 | 0,13783605 | 0,03814977 | 0,16924544 | NDUFAF4 |
| 1089,03326 | -0,2858951 | 0,11104832 | 0,01003818 | 0,06959782 | EMSY |
| 821,805623 | -0,2859881 | 0,11215499 | 0,01077425 | 0,07283299 | NANS |
| 1312,25749 | -0,2863526 | 0,10923629 | 0,00875681 | 0,06362813 | TMEM154 |
| 625,918086 | -0,2864656 | 0,12338423 | 0,02024717 | 0,11131718 | CCDC90B |
| 619,220088 | -0,2866746 | 0,09851428 | 0,00361452 | 0,03440161 | SCFD2 |
| 1329,57326 | -0,2868212 | 0,09843338 | 0,00356989 | 0,03413133 | MRPS2 |
| 1584,33263 | -0,2870632 | 0,11332957 | 0,01130927 | 0,07542243 | NIPA2 |
| 7298,10509 | -0,2871575 | 0,12014502 | 0,01684421 | 0,09863662 | PPA1 |
| 18139,2 | -0,2872288 | 0,10110795 | 0,00449986 | 0,04023721 | LDHA |
| 1120,30272 | -0,2872944 | 0,105105 | 0,00626834 | 0,05094641 | IRF2 |
| 7568,22092 | -0,28764 | 0,08360714 | 0,00058091 | 0,00889987 | LTBP1 |
| 401,937766 | -0,2879928 | 0,14325526 | 0,04439449 | 0,18724659 | ZNF430 |
| 2462,72412 | -0,2880544 | 0,10718031 | 0,00719744 | 0,05585839 | MTREX |
| 648,284044 | -0,2881853 | 0,11881908 | 0,015291 | 0,09258275 | RIOK2 |
| 1308,36666 | -0,2883369 | 0,1302918 | 0,02689707 | 0,13420872 | TMPRSS11F |
| 383,384421 | -0,2883853 | 0,1287347 | 0,02508105 | 0,12787887 | E2F8 |
| 446,779122 | -0,2885195 | 0,13446872 | 0,03190289 | 0,15002517 | PPP2R3C |
| 614,526617 | -0,2890238 | 0,12568337 | 0,02146984 | 0,11550604 | ANKRD29 |
| 1040,48869 | -0,2890248 | 0,09846932 | 0,00333358 | 0,03239584 | NDUFA9 |
| 2021,88257 | -0,2892559 | 0,1059402 | 0,00632634 | 0,0512152 | PSMD14 |
| 1872,95316 | -0,289264 | 0,12864812 | 0,02454497 | 0,12613127 | KIF11 |
| 2164,41957 | -0,2894271 | 0,09531546 | 0,00239328 | 0,02548295 | GOLGA4 |
| 2344,85779 | -0,2895966 | 0,08486051 | 0,00064342 | 0,00960102 | WAPL |
| 1799,00713 | -0,2896346 | 0,12228016 | 0,01785485 | 0,10221835 | ARL5A |
| 1082,77234 | -0,2896479 | 0,10009869 | 0,00380825 | 0,03563598 | POLR1D |
| 939,663589 | -0,2897139 | 0,09293685 | 0,00182504 | 0,0211331 | STN1 |
| 755,063261 | -0,2899124 | 0,10383267 | 0,00523653 | 0,04491401 | PARN |
| 1021,05099 | -0,2900363 | 0,09743893 | 0,00291468 | 0,02949127 | KIF21A |
| 1601,98537 | -0,2902807 | 0,13007708 | 0,02564108 | 0,13003289 | CREB1 |
| 608,528757 | -0,2903176 | 0,10703946 | 0,00668286 | 0,05306181 | PCGF1 |
| 4172,33632 | -0,2912525 | 0,08515853 | 0,00062593 | 0,00941435 | GAS5 |
| 643,589392 | -0,2912917 | 0,13640317 | 0,03271854 | 0,15221723 | ISCU |
| 623,081037 | -0,2914461 | 0,13267207 | 0,02803901 | 0,1378348 | EIF2B3 |
| 633,909634 | -0,2917126 | 0,12256165 | 0,01730657 | 0,10009407 | ANAPC4 |
| 1336,70357 | -0,2918229 | 0,11447373 | 0,01079529 | 0,0729231 | COG5 |
| 2637,759 | -0,291989 | 0,1073772 | 0,00654235 | 0,05230397 | SMARCA5 |
| 471,821084 | -0,2920991 | 0,11344011 | 0,01002636 | 0,06954131 | HEBP2 |
| 664,400874 | -0,2922514 | 0,09578416 | 0,0022797 | 0,02466223 | TMEM184C |
| 318,878295 | -0,292268 | 0,1443922 | 0,04295717 | 0,18330912 | ATP10D |
| 371,701559 | -0,2924768 | 0,13389029 | 0,02892909 | 0,14063908 | SDF2 |
| 364,618798 | -0,2926929 | 0,14916203 | 0,04973365 | 0,20176036 | NQO2 |
| 1491,12648 | -0,2929082 | 0,11154882 | 0,0086438 | 0,06299274 | CDCA7 |

| | | | | | |
|---|---|---|---|---|---|
| 1402,72424 | -0,2936585 | 0,10059144 | 0,00350797 | 0,03364135 | EHBP1 |
| 1033,02519 | -0,2939676 | 0,09612047 | 0,00222579 | 0,0242453 | MCTS1 |
| 880,091114 | -0,2940703 | 0,11369328 | 0,00969497 | 0,06811682 | MRPL35 |
| 2107,3587 | -0,2941649 | 0,10798148 | 0,00644544 | 0,05171486 | APEX1 |
| 2704,39812 | -0,2942721 | 0,10960415 | 0,00725597 | 0,05612844 | RAD21 |
| 725,355151 | -0,2942953 | 0,1040889 | 0,00469357 | 0,04161526 | DPYSL2 |
| 2053,13788 | -0,2943296 | 0,1091529 | 0,00700747 | 0,05477375 | ADAM10 |
| 659,549906 | -0,2944619 | 0,13632242 | 0,03076958 | 0,14653694 | SUCLA2 |
| 9962,12831 | -0,294529 | 0,07672577 | 0,00012368 | 0,00268973 | MAP3K20 |
| 542,721763 | -0,2947646 | 0,10910772 | 0,00690083 | 0,05424621 | TMEM243 |
| 3177,9618 | -0,2948525 | 0,10328939 | 0,00430876 | 0,03902634 | PPM1G |
| 744,196558 | -0,2950701 | 0,1032711 | 0,00427345 | 0,03882106 | BRCC3 |
| 885,152175 | -0,2953945 | 0,12168975 | 0,01520583 | 0,09235352 | TAOK3 |
| 659,814222 | -0,295397 | 0,10705831 | 0,00579402 | 0,0481884 | DTNBP1 |
| 611,737624 | -0,2954167 | 0,14046563 | 0,03545487 | 0,16073126 | GOLGA5 |
| 630,20944 | -0,2957444 | 0,117888 | 0,01211798 | 0,07917823 | LIN28B |
| 374,930282 | -0,2958887 | 0,12732835 | 0,02013493 | 0,11093238 | ZNF280B |
| 1075,70986 | -0,2959668 | 0,10364039 | 0,00429409 | 0,03894939 | TMEM209 |
| 557,292864 | -0,2959722 | 0,11446975 | 0,00972116 | 0,06819948 | ENTPD5 |
| 962,324608 | -0,2959753 | 0,11404109 | 0,00944976 | 0,06689071 | RSBN1L |
| 2155,20533 | -0,2963589 | 0,10928751 | 0,00669319 | 0,05307793 | CD63 |
| 1025,24751 | -0,2967025 | 0,11814895 | 0,01203016 | 0,07876218 | PLPBP |
| 570,720625 | -0,2967205 | 0,11038391 | 0,00718647 | 0,05579613 | AIMP2 |
| 1665,33483 | -0,2968678 | 0,12686965 | 0,01928668 | 0,10757795 | HDAC2 |
| 4201,08336 | -0,2968843 | 0,09811146 | 0,0024782 | 0,02616635 | FDFT1 |
| 1490,66122 | -0,2970226 | 0,11526669 | 0,0099711 | 0,06923421 | PSIP1 |
| 1059,26754 | -0,297027 | 0,12337967 | 0,01606558 | 0,09568048 | PLRG1 |
| 477,915406 | -0,2971904 | 0,1177046 | 0,01157367 | 0,0766919 | RPAIN |
| 361,842139 | -0,2972119 | 0,13953984 | 0,03317631 | 0,15370444 | PTS |
| 1915,82498 | -0,2972728 | 0,10543653 | 0,00481064 | 0,04233651 | ELOVL6 |
| 1647,06547 | -0,2973083 | 0,11099423 | 0,00739323 | 0,05677248 | ECT2 |
| 7495,28874 | -0,2973649 | 0,1189462 | 0,01241951 | 0,08039328 | SAMD9L |
| 7075,07838 | -0,2973732 | 0,0912184 | 0,00111407 | 0,01455174 | ANKRD17 |
| 2948,70408 | -0,2975552 | 0,12482279 | 0,01713395 | 0,09945292 | COX6B1 |
| 17785,4274 | -0,2978783 | 0,1116935 | 0,00765485 | 0,05815646 | IL2RA |
| 707,666671 | -0,2979502 | 0,10313008 | 0,00386381 | 0,0360489 | PON2 |
| 1361,07843 | -0,2980476 | 0,13207218 | 0,02402669 | 0,12424405 | DNAJA2 |
| 8997,43378 | -0,2980483 | 0,10238558 | 0,00360231 | 0,03433717 | NAP1L1 |
| 1255,02735 | -0,2980669 | 0,1012267 | 0,00323427 | 0,03169113 | DYNC2H1 |
| 410,515687 | -0,2980878 | 0,12877367 | 0,02062282 | 0,11258061 | ADGRA3 |
| 634,633658 | -0,2981083 | 0,14280995 | 0,03684771 | 0,16525138 | ZHX1 |
| 396,480686 | -0,2981482 | 0,11582067 | 0,01004656 | 0,06963041 | ERC2 |
| 1273,86539 | -0,2981948 | 0,084803 | 0,00043757 | 0,00722553 | SMPDL3A |
| 3113,14306 | -0,2982035 | 0,07565308 | 8,09E-05 | 0,0019278 | PHF14 |
| 5100,10421 | -0,2982876 | 0,09690652 | 0,00208328 | 0,0231189 | EPRS1 |
| 574,86786 | -0,2986546 | 0,13027821 | 0,02188044 | 0,11675122 | RABL3 |
| 951,03933 | -0,2987013 | 0,08906246 | 0,00079698 | 0,0113981 | DNAJC21 |
| 1323,92157 | -0,298719 | 0,115141 | 0,00947628 | 0,06705337 | RIMOC1 |

| | | | | | |
|---|---|---|---|---|---|
| 11499,6768 | -0,2988036 | 0,10388559 | 0,00402398 | 0,03714035 | VCAN |
| 1934,84644 | -0,2989108 | 0,0870196 | 0,00059259 | 0,00904959 | PSMC1 |
| 1099,67192 | -0,2991478 | 0,11206822 | 0,00760012 | 0,05785321 | STK39 |
| 281,895595 | -0,2994898 | 0,13585596 | 0,02749163 | 0,13612452 | SMCO4 |
| 838,987649 | -0,2995164 | 0,08801422 | 0,00066639 | 0,00988228 | ZNF83 |
| 749,191112 | -0,2995509 | 0,10944577 | 0,00620061 | 0,0505479 | CLIC5 |
| 651,031719 | -0,2995554 | 0,13578276 | 0,0273744 | 0,13584218 | KRR1 |
| 1785,66746 | -0,2998711 | 0,1170382 | 0,01040222 | 0,07110562 | GCC2 |
| 983,382159 | -0,29995 | 0,14140041 | 0,03389817 | 0,15603469 | CENPJ |
| 2429,7759 | -0,3000059 | 0,11893835 | 0,01165689 | 0,07701116 | PAF1 |
| 1434,71855 | -0,3001668 | 0,10825824 | 0,00555947 | 0,04683472 | RSU1 |
| 215,039878 | -0,3003535 | 0,1497198 | 0,04484514 | 0,1886008 | PDCL |
| 2978,50997 | -0,3009907 | 0,10460232 | 0,0040087 | 0,0370187 | DDX46 |
| 2662,63965 | -0,3009964 | 0,0833016 | 0,00030229 | 0,0053958 | GIGYF2 |
| 1517,4242 | -0,3010324 | 0,13543588 | 0,02623652 | 0,13185155 | PNISR |
| 12246,9219 | -0,301039 | 0,08026624 | 0,00017648 | 0,00357128 | KRIT1 |
| 4483,45682 | -0,3010901 | 0,12075043 | 0,01264939 | 0,08136571 | HNRNPR |
| 9882,51769 | -0,3012749 | 0,10684381 | 0,00480583 | 0,04233651 | HSPA9 |
| 1426,44413 | -0,301593 | 0,13170134 | 0,02202267 | 0,11724562 | BRIX1 |
| 430,174123 | -0,3020365 | 0,14106747 | 0,03226771 | 0,15097361 | CLHC1 |
| 1161,09498 | -0,3021715 | 0,10638146 | 0,00450496 | 0,04024473 | SEC63 |
| 672,274921 | -0,302298 | 0,1047778 | 0,00391247 | 0,03634158 | TMEM69 |
| 4819,20729 | -0,3026085 | 0,11408413 | 0,00798975 | 0,05972895 | SELPLG |
| 666,816326 | -0,3027163 | 0,11529797 | 0,00865185 | 0,06299274 | FBXW8 |
| 724,384259 | -0,3027745 | 0,10959707 | 0,00573404 | 0,04785785 | FER |
| 6086,47761 | -0,3032826 | 0,09283566 | 0,00108741 | 0,01432794 | CBX3 |
| 1034,18467 | -0,3039497 | 0,12323001 | 0,01364319 | 0,08561948 | PSMD5 |
| 557,416395 | -0,3041584 | 0,12048164 | 0,01158562 | 0,0766919 | NDUFC2 |
| 677,634368 | -0,3041625 | 0,09915522 | 0,00215829 | 0,02364623 | METTL15 |
| 1309,83313 | -0,3043726 | 0,09055087 | 0,00077564 | 0,0111575 | CPSF3 |
| 374,581607 | -0,3044915 | 0,12973553 | 0,0189244 | 0,10622029 | LIMA1 |
| 1011,56853 | -0,3046293 | 0,08643781 | 0,00042467 | 0,00707318 | SP110 |
| 2097,73941 | -0,3046946 | 0,13583994 | 0,02489415 | 0,12730328 | H4C6 |
| 826,064041 | -0,3047177 | 0,12961873 | 0,0187292 | 0,10554062 | DNAJC2 |
| 1408,16124 | -0,3047786 | 0,09587521 | 0,0014783 | 0,01806909 | EFL1 |
| 1559,15709 | -0,3052389 | 0,11553091 | 0,0082405 | 0,06102485 | ZNF37BP |
| 6301,58978 | -0,3055697 | 0,08393616 | 0,00027211 | 0,00500361 | CHD7 |
| 3110,61774 | -0,3056134 | 0,09349097 | 0,00107963 | 0,01427513 | CUL3 |
| 930,374871 | -0,3056486 | 0,12609523 | 0,0153527 | 0,09288624 | VTA1 |
| 1293,8467 | -0,3056682 | 0,09462041 | 0,00123588 | 0,01581069 | POMP |
| 494,605597 | -0,3057007 | 0,11577919 | 0,00828138 | 0,06123603 | SDCCAG8 |
| 582,998039 | -0,3058101 | 0,13375797 | 0,02223704 | 0,11779034 | SGCB |
| 9163,51685 | -0,3058539 | 0,12839187 | 0,01720999 | 0,09970308 | FTH1 |
| 1321,90587 | -0,3061431 | 0,13555405 | 0,02391713 | 0,12397567 | CKAP2 |
| 2185,51877 | -0,3065134 | 0,13983832 | 0,02838578 | 0,1391418 | MYOF |
| 1082,02175 | -0,3066958 | 0,11539431 | 0,0078651 | 0,05914766 | TYW1 |
| 692,605659 | -0,307493 | 0,10710716 | 0,00409316 | 0,03766865 | CSTF2 |
| 389,05726 | -0,3076683 | 0,14149872 | 0,02967859 | 0,14303324 | TRMT13 |

| | | | | | |
|---|---|---|---|---|---|
| 413,551655 | -0,3077123 | 0,11874532 | 0,00955965 | 0,06744149 | ALG6 |
| 1072,79217 | -0,3077381 | 0,10442594 | 0,00320924 | 0,03152757 | KLHL8 |
| 430,626289 | -0,3077929 | 0,1233737 | 0,01260276 | 0,08124594 | HDAC8 |
| 539,2867 | -0,30797 | 0,15451297 | 0,04624329 | 0,19272451 | BCAS2 |
| 684,423951 | -0,3080405 | 0,13971488 | 0,02746959 | 0,13612452 | HBEGF |
| 5544,60128 | -0,3082224 | 0,08273573 | 0,00019502 | 0,00386786 | TANC2 |
| 537,01152 | -0,3084111 | 0,13242355 | 0,01986042 | 0,11000557 | SAV1 |
| 1066,7243 | -0,3084404 | 0,13422462 | 0,02156511 | 0,11581988 | MAK16 |
| 1188,98152 | -0,3084684 | 0,12911768 | 0,01689207 | 0,09873551 | DLGAP5 |
| 1656,89247 | -0,308598 | 0,09402594 | 0,00103055 | 0,01380949 | PLD1 |
| 470,593054 | -0,3086423 | 0,1327766 | 0,02009748 | 0,1108232 | TIGAR |
| 6170,28581 | -0,309067 | 0,11644541 | 0,00795016 | 0,05950358 | PRPF40A |
| 563,657918 | -0,30922 | 0,13937269 | 0,02651017 | 0,13269814 | EMC3 |
| 1538,54674 | -0,3093106 | 0,08202684 | 0,00016269 | 0,0033424 | PRPF38B |
| 1425,36419 | -0,3094235 | 0,11726475 | 0,00832309 | 0,06139616 | SMC2 |
| 411,516425 | -0,3094281 | 0,15645673 | 0,04795992 | 0,19718592 | ITGB3BP |
| 1808,20246 | -0,3094515 | 0,11050138 | 0,00510344 | 0,0440521 | NFE2L2 |
| 563,073238 | -0,3097585 | 0,11980065 | 0,0097205 | 0,06819948 | FERRY3 |
| 681,074923 | -0,3097666 | 0,14565914 | 0,03344882 | 0,15447525 | SPDYE3 |
| 696,076648 | -0,3097976 | 0,11011619 | 0,00490253 | 0,04285407 | SCO1 |
| 583,735358 | -0,3098228 | 0,13665273 | 0,02337636 | 0,12227028 | BLOC1S2 |
| 311,256319 | -0,3098528 | 0,13722595 | 0,02394721 | 0,12397567 | CCDC51 |
| 524,448343 | -0,3099473 | 0,12968539 | 0,01684866 | 0,09863662 | DNAJC1 |
| 387,711316 | -0,3099518 | 0,13422979 | 0,02093728 | 0,11373936 | TDRD3 |
| 2327,81664 | -0,3102807 | 0,11143624 | 0,00536304 | 0,04558586 | SOS1 |
| 572,776126 | -0,3106181 | 0,1343844 | 0,02080995 | 0,11324699 | THAP5 |
| 1273,64934 | -0,3109427 | 0,08670885 | 0,00033572 | 0,0058816 | NBEAL1 |
| 3529,53116 | -0,311321 | 0,10073236 | 0,00199769 | 0,02248555 | MYL12A |
| 701,641027 | -0,3114264 | 0,10458739 | 0,00290452 | 0,02946723 | NEK1 |
| 2552,12019 | -0,3114339 | 0,11154858 | 0,00523974 | 0,04492124 | ACSL3 |
| 212,049047 | -0,3118785 | 0,151763 | 0,03987551 | 0,17421138 | ZFYVE21 |
| 1152,7281 | -0,3120244 | 0,12088758 | 0,00984825 | 0,06874025 | PIAS1 |
| 430,129712 | -0,3121253 | 0,13450155 | 0,02030802 | 0,11157027 | MRPL53 |
| 3296,68154 | -0,3123497 | 0,1030685 | 0,00244144 | 0,02589378 | ATRX |
| 2438,33303 | -0,3123986 | 0,12475607 | 0,01227718 | 0,07974787 | UBE2L3 |
| 1854,1058 | -0,3124589 | 0,08982959 | 0,00050451 | 0,00804193 | VPS41 |
| 1821,13154 | -0,3125205 | 0,14359015 | 0,02951967 | 0,1425215 | KIF20B |
| 1926,33575 | -0,3125516 | 0,09629845 | 0,0011718 | 0,01514014 | ANK2 |
| 1044,97386 | -0,3126707 | 0,09874052 | 0,00154238 | 0,01865945 | GOLIM4 |
| 3967,33143 | -0,3129298 | 0,11991314 | 0,00906382 | 0,06498546 | ARPC2 |
| 3191,93796 | -0,3129463 | 0,09339339 | 0,00080567 | 0,01149629 | SRSF5 |
| 823,262895 | -0,3130874 | 0,10875078 | 0,0039901 | 0,03691766 | MLH1 |
| 2917,84874 | -0,313528 | 0,12010591 | 0,00904286 | 0,06495825 | GNL3 |
| 950,780674 | -0,3135547 | 0,09994272 | 0,00170486 | 0,02007323 | SPATS2L |
| 1069,23132 | -0,3139997 | 0,08352185 | 0,00017026 | 0,00347891 | TAF1B |
| 622,504361 | -0,3142724 | 0,1516117 | 0,03818371 | 0,1693563 | POLE2 |
| 556,162377 | -0,314273 | 0,10392198 | 0,00249354 | 0,02628427 | SPATS2 |
| 683,914388 | -0,3143209 | 0,11831997 | 0,00789487 | 0,05925377 | ZNF195 |

| | | | | | |
|---|---|---|---|---|---|
| 2113,83175 | -0,3144834 | 0,1035387 | 0,00238667 | 0,02542691 | AGPS |
| 1998,61177 | -0,3146449 | 0,10645394 | 0,0031197 | 0,03086766 | MAP4K5 |
| 5359,37766 | -0,3149636 | 0,06689562 | 2,50E-06 | 0,00010694 | DLGAP4 |
| 757,140477 | -0,3149883 | 0,12656051 | 0,01281622 | 0,08192421 | GCLC |
| 4746,99626 | -0,3150542 | 0,09589347 | 0,00101816 | 0,0136628 | TM4SF1 |
| 536,645254 | -0,3154031 | 0,12909716 | 0,01455986 | 0,08956437 | ZNF248 |
| 1683,6437 | -0,3156658 | 0,09588828 | 0,00099472 | 0,01341246 | WDR3 |
| 3626,40064 | -0,3157649 | 0,08012122 | 8,11E-05 | 0,00193047 | EZR |
| 4120,65835 | -0,315799 | 0,11540076 | 0,00620887 | 0,05059349 | NIPBL |
| 4981,16219 | -0,3159555 | 0,10198273 | 0,00194748 | 0,0220648 | SPEN |
| 2939,91039 | -0,3160634 | 0,12566325 | 0,01189776 | 0,07819299 | RARS1 |
| 3354,03028 | -0,3161911 | 0,10039485 | 0,00163564 | 0,01953786 | QSER1 |
| 10265,4491 | -0,3162187 | 0,09354464 | 0,00072379 | 0,01055077 | SERBP1 |
| 731,088429 | -0,3163577 | 0,12256558 | 0,00984774 | 0,06874025 | CYSTM1 |
| 229,431872 | -0,3166558 | 0,15620992 | 0,04265043 | 0,18212339 | MREG |
| 9270,89674 | -0,3168947 | 0,09375074 | 0,00072437 | 0,01055111 | LDHB |
| 4610,04232 | -0,3168985 | 0,08666007 | 0,00025538 | 0,00475175 | ERO1A |
| 2313,17676 | -0,3173391 | 0,10092025 | 0,00166397 | 0,01978536 | ATN1 |
| 2256,70286 | -0,3173622 | 0,1216063 | 0,00906081 | 0,06498546 | LOC102723728 |
| 1176,04596 | -0,3174116 | 0,10675409 | 0,00294619 | 0,02971657 | TMX3 |
| 2442,25872 | -0,3176241 | 0,08821087 | 0,00031732 | 0,00560592 | ELOVL5 |
| 2937,2936 | -0,3177202 | 0,10341381 | 0,00212403 | 0,02338155 | MTHFD1 |
| 615,398474 | -0,3177734 | 0,12704727 | 0,01237658 | 0,08022515 | TMEM14B |
| 730,91073 | -0,3182577 | 0,11584871 | 0,00601095 | 0,04942769 | MRPS7 |
| 944,436337 | -0,3184685 | 0,10722763 | 0,00297778 | 0,02985823 | ELOF1 |
| 483,325699 | -0,318547 | 0,14247636 | 0,02536563 | 0,12905167 | FIGNL1 |
| 273,302149 | -0,3185761 | 0,14636793 | 0,02951466 | 0,1425215 | PRPF18 |
| 1366,17626 | -0,3190183 | 0,14224765 | 0,0249164 | 0,12731904 | ERG28 |
| 373,532542 | -0,3191474 | 0,13655077 | 0,01942844 | 0,10811927 | ARMT1 |
| 884,196711 | -0,3194174 | 0,11329096 | 0,00481071 | 0,04233651 | C5orf22 |
| 834,929543 | -0,3194764 | 0,12138027 | 0,00848766 | 0,0622461 | RAB13 |
| 2029,88001 | -0,3203246 | 0,08988714 | 0,00036576 | 0,00627874 | SP140 |
| 1643,09243 | -0,3204047 | 0,09319687 | 0,00058617 | 0,00897323 | PPIG |
| 1326,37707 | -0,3204362 | 0,11702266 | 0,00617689 | 0,05040878 | INTS13 |
| 656,850961 | -0,3206219 | 0,12482337 | 0,0102109 | 0,07022914 | GXYLT1 |
| 3666,54096 | -0,3207028 | 0,09087996 | 0,00041735 | 0,00697961 | CEP170 |
| 391,043127 | -0,3208796 | 0,11910283 | 0,00705707 | 0,05506261 | PAAF1 |
| 962,520322 | -0,3210111 | 0,12612202 | 0,01092019 | 0,0734904 | MAP3K7 |
| 1212,11826 | -0,3210991 | 0,09507237 | 0,0007317 | 0,01062739 | MRPS5 |
| 2396,79964 | -0,3212277 | 0,11913023 | 0,00700847 | 0,05477375 | CMAHP |
| 673,572115 | -0,3213138 | 0,12425543 | 0,00971213 | 0,06818673 | NDUFAF7 |
| 1233,5574 | -0,3214672 | 0,11786981 | 0,00638535 | 0,05149048 | RSRC2 |
| 1831,93268 | -0,3215025 | 0,10393539 | 0,00197949 | 0,0223339 | RCN1 |
| 1334,6469 | -0,3215097 | 0,11065283 | 0,00366583 | 0,03473267 | SPCS2 |
| 1288,39077 | -0,3217933 | 0,11554502 | 0,00535272 | 0,04553902 | SEPHS1 |
| 991,703746 | -0,3218623 | 0,10421232 | 0,00201153 | 0,02254749 | BASP1 |
| 2924,30841 | -0,3219242 | 0,09979171 | 0,0012555 | 0,01596465 | RAC2 |
| 3327,63609 | -0,3219868 | 0,08858759 | 0,00027833 | 0,00508828 | MAPK1IP1L |

| | | | | | |
|---|---|---|---|---|---|
| 3589,06249 | -0,32211 | 0,09008332 | 0,00034931 | 0,00606908 | RBM25 |
| 281,88278 | -0,3221813 | 0,14540695 | 0,02671061 | 0,13352481 | PLAAT3 |
| 301,552388 | -0,3224598 | 0,14764474 | 0,02896043 | 0,14071379 | TEC |
| 733,700427 | -0,3226019 | 0,10810701 | 0,00284416 | 0,02911883 | PMP22 |
| 628,76449 | -0,3226345 | 0,13269991 | 0,01504437 | 0,09164665 | TRERF1 |
| 468,948579 | -0,3226487 | 0,12232368 | 0,00834795 | 0,06155554 | PDIA5 |
| 191456,307 | -0,3228027 | 0,14538124 | 0,02639249 | 0,13231908 | RN7SK |
| 1644,23009 | -0,3233773 | 0,10691829 | 0,00249022 | 0,02626393 | SLC25A23 |
| 499,564148 | -0,3238562 | 0,1417099 | 0,02229254 | 0,11801825 | PPIL3 |
| 240,325814 | -0,3238599 | 0,1652065 | 0,04995676 | 0,2022319 | TMEM35B |
| 347,390878 | -0,3241059 | 0,13905429 | 0,01976461 | 0,10958961 | PBDC1 |
| 721,654543 | -0,3245561 | 0,1213452 | 0,0074807 | 0,05728142 | CASP8AP2 |
| 275,828173 | -0,3245696 | 0,15265986 | 0,0334952 | 0,15461398 | DYRK4 |
| 1321,67871 | -0,3246763 | 0,12652012 | 0,01028187 | 0,07058812 | DENR |
| 1034,61859 | -0,3247108 | 0,10243062 | 0,00152409 | 0,01851513 | PRR3 |
| 363,865994 | -0,3250583 | 0,14016381 | 0,02038792 | 0,11179276 | ATOX1 |
| 4269,86227 | -0,3253275 | 0,09020564 | 0,00031034 | 0,00550844 | DST |
| 882,769499 | -0,3257804 | 0,10366831 | 0,00167497 | 0,01984669 | FTSJ3 |
| 660,548062 | -0,326583 | 0,10867823 | 0,00265541 | 0,02766686 | PTPRB |
| 3850,66248 | -0,3266678 | 0,13863546 | 0,01845762 | 0,10462449 | EIF2S2 |
| 1727,37788 | -0,3267785 | 0,11057493 | 0,00312398 | 0,03086766 | ZNF148 |
| 351,150643 | -0,3268385 | 0,12183182 | 0,007303 | 0,05637702 | PTPRK |
| 623,229527 | -0,3270865 | 0,11622973 | 0,00489082 | 0,04278277 | PSMG2 |
| 1337,1712 | -0,3273188 | 0,11416795 | 0,00414391 | 0,03806163 | CSTF3 |
| 1565,64212 | -0,3273486 | 0,145052 | 0,02402264 | 0,12424405 | BCL2A1 |
| 920,977851 | -0,3274203 | 0,12748639 | 0,01022056 | 0,07024456 | MCTP1 |
| 1036,13101 | -0,3275077 | 0,08884247 | 0,00022746 | 0,00435907 | RBM28 |
| 626,290625 | -0,3275229 | 0,12215966 | 0,00733794 | 0,05652531 | SMAP1 |
| 186,566959 | -0,3277152 | 0,16515303 | 0,04722104 | 0,19520849 | DHRS4-AS1 |
| 537,61553 | -0,3277377 | 0,13318678 | 0,01386519 | 0,0866964 | ABITRAM |
| 552,224831 | -0,3280674 | 0,1132738 | 0,0037767 | 0,03544594 | FANCL |
| 2109,95883 | -0,328151 | 0,09636055 | 0,00066055 | 0,00982572 | FXR1 |
| 217,718946 | -0,3281852 | 0,15105351 | 0,02980728 | 0,1432521 | LOC100294145 |
| 472,406848 | -0,32824 | 0,13269657 | 0,01337543 | 0,08438698 | CETN2 |
| 1853,71145 | -0,3284788 | 0,12037095 | 0,00635483 | 0,05132202 | EIF2AK2 |
| 8743,4236 | -0,3286081 | 0,09439296 | 0,00049903 | 0,00799102 | CSDE1 |
| 389,080962 | -0,3286456 | 0,13248863 | 0,01311772 | 0,08317708 | PTGR2 |
| 7521,97685 | -0,3287891 | 0,10807491 | 0,00234829 | 0,02514306 | HSP90B1 |
| 1811,8444 | -0,3288092 | 0,10644144 | 0,00200758 | 0,02251655 | MCCC2 |
| 3512,56733 | -0,3289069 | 0,12389532 | 0,00793752 | 0,05945598 | USP7 |
| 1374,21337 | -0,3294211 | 0,11344984 | 0,00368814 | 0,03489169 | NOL8 |
| 545,521557 | -0,3294869 | 0,13214519 | 0,01265368 | 0,08136571 | GANC |
| 6552,90685 | -0,3296075 | 0,07946391 | 3,36E-05 | 0,00093921 | CALM1 |
| 6160,21489 | -0,3298789 | 0,0985565 | 0,0008166 | 0,01161726 | BAZ1B |
| 4282,33125 | -0,330124 | 0,08535286 | 0,00010984 | 0,0024683 | PTPN12 |
| 361,916262 | -0,330231 | 0,12626636 | 0,00891355 | 0,06427333 | POU3F2 |
| 768,067586 | -0,3304593 | 0,09439126 | 0,0004636 | 0,00756838 | SLU7 |
| 379,212448 | -0,3305747 | 0,13483465 | 0,01421814 | 0,08816323 | ZNHIT3 |

| | | | | | |
|---|---|---|---|---|---|
| 4903,59058 | -0,3307857 | 0,1265302 | 0,0089415 | 0,06442578 | NPIPB4 |
| 429,059361 | -0,3314509 | 0,12362176 | 0,00733654 | 0,05652531 | MED8 |
| 1622,33835 | -0,3320098 | 0,10626834 | 0,00178253 | 0,02078086 | PARK7 |
| 1386,52816 | -0,3320647 | 0,09011878 | 0,00022893 | 0,00437538 | PRR13 |
| 601,499372 | -0,3320936 | 0,120245 | 0,00574823 | 0,04793127 | PCBD1 |
| 4600,33909 | -0,3321791 | 0,11539961 | 0,00399557 | 0,03695027 | TCERG1 |
| 1748,86399 | -0,3321964 | 0,10645231 | 0,00180475 | 0,02097529 | LOC101929128 |
| 24600,9096 | -0,3327002 | 0,12258524 | 0,00664698 | 0,05286571 | HSP90AA1 |
| 428,264908 | -0,3330214 | 0,15721418 | 0,03415259 | 0,15676882 | ZNF33B |
| 1080,65743 | -0,3332214 | 0,10186139 | 0,00107046 | 0,0142035 | SREK1IP1 |
| 2026,65965 | -0,3334015 | 0,13558835 | 0,0139354 | 0,08693432 | ANXA3 |
| 3535,4997 | -0,3337031 | 0,11864398 | 0,00491362 | 0,04286338 | NAA15 |
| 704,60262 | -0,3342439 | 0,12679887 | 0,00838857 | 0,06171429 | DNAJA4 |
| 3063,14171 | -0,3346287 | 0,14804342 | 0,02380005 | 0,12368052 | RAP1B |
| 311,507589 | -0,334933 | 0,14231985 | 0,01860349 | 0,10516779 | APOO |
| 862,668919 | -0,3351316 | 0,1247667 | 0,00722988 | 0,05601827 | ABCC9 |
| 2421,67146 | -0,3352525 | 0,1053534 | 0,00146176 | 0,0179364 | TOMM6 |
| 408,586324 | -0,3352741 | 0,13534246 | 0,01324075 | 0,0837317 | UQCC2 |
| 443,729435 | -0,3353507 | 0,13644957 | 0,0139834 | 0,0871058 | ITFG1 |
| 1664,96737 | -0,3357678 | 0,13484694 | 0,01277459 | 0,0817991 | NUSAP1 |
| 1530,32468 | -0,3360323 | 0,14549597 | 0,0209123 | 0,11366896 | HAX1 |
| 1829,37105 | -0,3360349 | 0,10896156 | 0,00204254 | 0,02280049 | ZNF121 |
| 1581,29363 | -0,3362427 | 0,09151492 | 0,00023861 | 0,00450793 | CLCN3 |
| 830,069148 | -0,3363007 | 0,12667452 | 0,00793458 | 0,05945598 | CNOT4 |
| 5346,13357 | -0,3364599 | 0,11086145 | 0,00240569 | 0,02558629 | UBA2 |
| 3364,5257 | -0,3364828 | 0,08024576 | 2,75E-05 | 0,00080204 | PRP4K |
| 267,755738 | -0,3367575 | 0,16499663 | 0,04125117 | 0,17811809 | CAAP1 |
| 547,191971 | -0,3368613 | 0,12555225 | 0,0072957 | 0,05634367 | TMEM18 |
| 322,787869 | -0,3369329 | 0,13765824 | 0,01438097 | 0,08877745 | ATPSCKMT |
| 4892,5187 | -0,3371038 | 0,10836686 | 0,00186604 | 0,02145037 | STIP1 |
| 353,945618 | -0,3371376 | 0,12960053 | 0,00928549 | 0,06599957 | RRS1 |
| 309,30637 | -0,3373201 | 0,15953245 | 0,03447862 | 0,15770813 | ALKBH8 |
| 2552,0049 | -0,3373782 | 0,08471773 | 6,82E-05 | 0,00165805 | NCOA6 |
| 564,719675 | -0,3375541 | 0,13100402 | 0,00997572 | 0,06924085 | RAD51B |
| 3633,45743 | -0,337622 | 0,10026775 | 0,00075934 | 0,01095922 | PHF3 |
| 575,87492 | -0,3378131 | 0,1134582 | 0,00290682 | 0,02947482 | ERCC4 |
| 7963,13041 | -0,3379324 | 0,06564223 | 2,63E-07 | 1,65E-05 | ZFP36L1 |
| 872,191972 | -0,3379634 | 0,11181899 | 0,00250766 | 0,02638904 | IDI1 |
| 638,85042 | -0,3388263 | 0,11433417 | 0,00304188 | 0,03032425 | VPS9D1-AS1 |
| 2065,29798 | -0,3388627 | 0,08599864 | 8,14E-05 | 0,00193177 | ZNF609 |
| 371,384303 | -0,3390792 | 0,14616251 | 0,02034748 | 0,11162499 | MBIP |
| 4074,11604 | -0,3390917 | 0,11588852 | 0,00343333 | 0,03301864 | ABCE1 |
| 208,266646 | -0,3391557 | 0,14600023 | 0,02018019 | 0,11106148 | CASP6 |
| 452,041984 | -0,3394403 | 0,14198134 | 0,0168145 | 0,09850395 | GPATCH11 |
| 207,584131 | -0,3396966 | 0,16298013 | 0,03713451 | 0,16609499 | CTXN1 |
| 1111,47574 | -0,3397316 | 0,11531223 | 0,00321716 | 0,03156152 | PMS1 |
| 419,601206 | -0,339954 | 0,13589964 | 0,01236659 | 0,08018787 | ZNF438 |
| 4576,45609 | -0,3402246 | 0,11533634 | 0,00317931 | 0,03133112 | EWSR1 |

| | | | | | |
|---|---|---|---|---|---|
| 1296,1859 | -0,3403713 | 0,105723 | 0,00128431 | 0,01620027 | CSPP1 |
| 258,17444 | -0,340396 | 0,14871867 | 0,02208756 | 0,11742589 | ARSK |
| 553,410498 | -0,3414314 | 0,15894615 | 0,0317062 | 0,14930637 | RPF1 |
| 3696,99599 | -0,3414323 | 0,07279178 | 2,72E-06 | 0,00011407 | SMC1A |
| 3655,15937 | -0,3424291 | 0,08274342 | 3,50E-05 | 0,00096726 | GPATCH8 |
| 185,677392 | -0,3424949 | 0,17412605 | 0,0491905 | 0,20045948 | EFCAB7 |
| 1214,0665 | -0,3426982 | 0,09870915 | 0,00051699 | 0,00818568 | CTCF |
| 1305,76946 | -0,3428895 | 0,09713614 | 0,00041558 | 0,00696608 | SAP130 |
| 544,785794 | -0,3429272 | 0,11879718 | 0,00389353 | 0,0362547 | NDUFV3 |
| 594,814076 | -0,3429354 | 0,17116288 | 0,04511702 | 0,1893234 | ASGR2 |
| 471,60469 | -0,3432195 | 0,1301473 | 0,00836028 | 0,06159849 | AIMP1 |
| 607,764333 | -0,3432885 | 0,11789138 | 0,00359232 | 0,0342765 | ECHDC1 |
| 3875,68525 | -0,3433784 | 0,1156725 | 0,00299218 | 0,02997093 | SRRM1 |
| 199,551105 | -0,3436463 | 0,14812843 | 0,02034513 | 0,11162499 | CLN5 |
| 1265,7243 | -0,3439039 | 0,12748724 | 0,00698506 | 0,0547038 | C14orf119 |
| 1570,23583 | -0,3440732 | 0,11161459 | 0,00205139 | 0,02284536 | RFX7 |
| 1091,57233 | -0,3441002 | 0,14215524 | 0,01549513 | 0,09348961 | SAP18 |
| 1418,89515 | -0,3442056 | 0,08411447 | 4,27E-05 | 0,00113597 | NDRG3 |
| 2522,05728 | -0,3446584 | 0,09674939 | 0,0003675 | 0,00629279 | ST13 |
| 1235,39834 | -0,3448439 | 0,10307823 | 0,00082147 | 0,01166892 | IGF2BP2 |
| 11722,4425 | -0,3450451 | 0,09247868 | 0,00019066 | 0,0038014 | HNRNPK |
| 257,143727 | -0,3452849 | 0,140491 | 0,01398295 | 0,0871058 | ST7 |
| 6821,61537 | -0,3453193 | 0,08845958 | 9,47E-05 | 0,00218834 | CCT5 |
| 1465,43348 | -0,3453586 | 0,08959551 | 0,0001159 | 0,00257288 | RSF1 |
| 719,000613 | -0,3454908 | 0,11337612 | 0,00230909 | 0,02489476 | HSBP1 |
| 14303,6793 | -0,3456088 | 0,12472484 | 0,00558895 | 0,04695763 | RPS6 |
| 510,486657 | -0,3457749 | 0,12283055 | 0,00487687 | 0,04273979 | ZNF117 |
| 1091,41392 | -0,3461328 | 0,11028104 | 0,00169732 | 0,02003431 | CEP152 |
| 422,398763 | -0,3461579 | 0,14486848 | 0,01687303 | 0,09868931 | TBP |
| 615,358295 | -0,3465061 | 0,12718456 | 0,00644116 | 0,05171486 | WDR70 |
| 2100,10559 | -0,3466258 | 0,08650437 | 6,15E-05 | 0,00151878 | PALS2 |
| 4989,64074 | -0,3466283 | 0,08096477 | 1,86E-05 | 0,00057842 | UBAP2L |
| 260,18151 | -0,3466295 | 0,16408376 | 0,03464224 | 0,15816828 | EBAG9 |
| 1652,40892 | -0,3467237 | 0,16128318 | 0,03157246 | 0,1489085 | PPP1R14B |
| 2286,66758 | -0,3480185 | 0,09013311 | 0,00011285 | 0,0025239 | COPB2 |
| 387,335389 | -0,3483619 | 0,12765808 | 0,00635524 | 0,05132202 | COA5 |
| 299,495955 | -0,3485144 | 0,14612686 | 0,01707852 | 0,09933684 | PAPSS1 |
| 323,207854 | -0,3485371 | 0,13235377 | 0,0084541 | 0,06202406 | CRYZ |
| 1187,65077 | -0,3487685 | 0,09110083 | 0,00012899 | 0,00277343 | MLKL |
| 1756,10703 | -0,3491975 | 0,09925074 | 0,00043426 | 0,00718869 | TRIP11 |
| 171,392442 | -0,3492393 | 0,17120987 | 0,04136709 | 0,17834194 | ZNF449 |
| 2702,31518 | -0,3495292 | 0,11340069 | 0,00205443 | 0,02286578 | HMGN1 |
| 260,045296 | -0,3498501 | 0,15227523 | 0,0215911 | 0,11586293 | WASF1 |
| 843,467729 | -0,3503616 | 0,10614823 | 0,0009645 | 0,0130913 | IL1RL2 |
| 852,699673 | -0,3510819 | 0,12130272 | 0,0038005 | 0,03559866 | PDHB |
| 432,705554 | -0,3513525 | 0,12473091 | 0,0048492 | 0,04257619 | CBR4 |
| 1028,56309 | -0,3513554 | 0,1091341 | 0,00128422 | 0,01620027 | MBD6 |
| 1029,71428 | -0,3514784 | 0,10911933 | 0,00127724 | 0,01614335 | NCBP2 |

| | | | | | |
|---|---|---|---|---|---|
| 514,360321 | -0,3518215 | 0,17843218 | 0,04863951 | 0,19911472 | PMS2P7 |
| 24640,9422 | -0,3519317 | 0,08183673 | 1,70E-05 | 0,00054164 | SRRM2 |
| 457,389163 | -0,3521843 | 0,12163897 | 0,00378766 | 0,03551355 | HABP4 |
| 359,615443 | -0,3523125 | 0,14535088 | 0,01535586 | 0,09288624 | NDUFA2 |
| 2069,1649 | -0,3524644 | 0,11183952 | 0,00162423 | 0,01944137 | RABEP1 |
| 1606,28548 | -0,3525586 | 0,10679112 | 0,00096209 | 0,01306796 | ATP5MC1 |
| 1040,60689 | -0,3525691 | 0,10227659 | 0,0005664 | 0,00876737 | PUS7L |
| 1207,21775 | -0,3526802 | 0,11256694 | 0,00172988 | 0,02030244 | CEP290 |
| 255,641276 | -0,3527339 | 0,14187867 | 0,01291277 | 0,08235344 | PLCE1 |
| 800,23045 | -0,3530201 | 0,16056654 | 0,02790704 | 0,13736449 | TLN2 |
| 989,393025 | -0,3530696 | 0,14175862 | 0,01275135 | 0,08173473 | SAFB2 |
| 3870,54833 | -0,3531741 | 0,10711902 | 0,00097715 | 0,01324398 | FDPS |
| 1778,65092 | -0,353363 | 0,12069799 | 0,00341521 | 0,03293534 | SLC25A46 |
| 1539,63472 | -0,353384 | 0,11608488 | 0,00233311 | 0,02503956 | SNRPC |
| 5519,95831 | -0,3535806 | 0,08532814 | 3,42E-05 | 0,0009506 | CLIP1 |
| 762,343405 | -0,3535838 | 0,10261327 | 0,00056939 | 0,00878254 | TLR6 |
| 4055,9473 | -0,3536413 | 0,11937093 | 0,00305111 | 0,03038423 | WDR43 |
| 2179,05945 | -0,3536836 | 0,11053466 | 0,00137545 | 0,01712169 | CFAP97 |
| 261,370929 | -0,3540854 | 0,15764815 | 0,02470091 | 0,12653605 | SLC7A11 |
| 1071,8101 | -0,3541434 | 0,09633341 | 0,00023671 | 0,00448776 | LINC00342 |
| 447,856924 | -0,3542233 | 0,1189358 | 0,00289877 | 0,02944967 | POLR2J |
| 11268,3775 | -0,3542512 | 0,13919166 | 0,01092588 | 0,0734904 | RPS14 |
| 2800,11354 | -0,354344 | 0,10371761 | 0,00063448 | 0,00951925 | TOP1 |
| 460,5331 | -0,3544368 | 0,16261579 | 0,02928737 | 0,14168914 | NFYB |
| 457,175995 | -0,3544678 | 0,12375731 | 0,00418042 | 0,03826691 | UBE2E2 |
| 1078,22304 | -0,3546277 | 0,14981481 | 0,01792774 | 0,10249258 | COX7A2 |
| 504,240283 | -0,3546668 | 0,10832838 | 0,00106038 | 0,01412609 | HACD2 |
| 1105,53807 | -0,3549966 | 0,12614576 | 0,00489021 | 0,04278277 | H2BC11 |
| 594,352962 | -0,3551173 | 0,10450044 | 0,00067822 | 0,01003324 | DDRGK1 |
| 1332,32946 | -0,3551357 | 0,13752605 | 0,00981398 | 0,06859635 | COX7A2L |
| 5795,04019 | -0,3551509 | 0,09454768 | 0,00017243 | 0,00351193 | LARS1 |
| 6108,67539 | -0,3552985 | 0,08725053 | 4,66E-05 | 0,00122404 | FUS |
| 1045,94304 | -0,3555394 | 0,1166924 | 0,00231285 | 0,02492108 | RRBP1 |
| 903,489186 | -0,3559604 | 0,10007936 | 0,00037543 | 0,00640528 | COIL |
| 690,421391 | -0,3560553 | 0,11105989 | 0,00134605 | 0,01682212 | IL18R1 |
| 354,237579 | -0,3561742 | 0,14223858 | 0,01227766 | 0,07974787 | SMIM26 |
| 562,590239 | -0,3562345 | 0,13141326 | 0,0067122 | 0,05313873 | FAM222B |
| 744,205646 | -0,3565739 | 0,10730861 | 0,00089092 | 0,01238587 | ELP3 |
| 1667,46642 | -0,357938 | 0,10535962 | 0,00068055 | 0,01005204 | RBM6 |
| 761,403195 | -0,3580534 | 0,10374959 | 0,00055824 | 0,00869343 | CCDC144BP |
| 601,222966 | -0,3583036 | 0,14385076 | 0,01274555 | 0,08173473 | LSM7 |
| 260,570412 | -0,3583485 | 0,172275 | 0,03751676 | 0,16730016 | TSPAN31 |
| 295,271642 | -0,3588207 | 0,16314481 | 0,02784948 | 0,13718824 | RNF113A |
| 541,35558 | -0,3592577 | 0,15639987 | 0,02161614 | 0,11592941 | TERF1 |
| 599,202735 | -0,3593568 | 0,15868201 | 0,0235351 | 0,12277577 | THOC7 |
| 221,919301 | -0,359683 | 0,18160547 | 0,04763914 | 0,19663615 | TSGA10 |
| 736,095247 | -0,3601662 | 0,13191654 | 0,00632848 | 0,0512152 | MANF |
| 3349,51996 | -0,3601745 | 0,09899739 | 0,00027453 | 0,00504302 | YLPM1 |

| | | | | | |
|---|---|---|---|---|---|
| 523,534083 | -0,3603262 | 0,1428495 | 0,01165511 | 0,07701116 | POLR3B |
| 273,314689 | -0,360337 | 0,15274707 | 0,01832209 | 0,10416833 | NSUN3 |
| 571,867487 | -0,3612537 | 0,12731869 | 0,00454829 | 0,040498 | RLIG1 |
| 502,670688 | -0,3612939 | 0,12472692 | 0,00377135 | 0,03544015 | COBLL1 |
| 2443,18615 | -0,3614969 | 0,11401355 | 0,00152099 | 0,01849528 | RAD50 |
| 900,529498 | -0,3618868 | 0,13652117 | 0,00803074 | 0,05989346 | SUCLG1 |
| 2093,00423 | -0,3619555 | 0,12178966 | 0,00295894 | 0,02971657 | PDHA1 |
| 690,062527 | -0,3620901 | 0,117296 | 0,00202203 | 0,02263835 | EXOSC8 |
| 340,167182 | -0,3621666 | 0,15132466 | 0,01669718 | 0,09805887 | SRRD |
| 426,200256 | -0,3622134 | 0,14240064 | 0,01097098 | 0,07371516 | NDUFC1 |
| 1008,82042 | -0,3630282 | 0,11111301 | 0,00108618 | 0,01432794 | AIFM1 |
| 1054,43798 | -0,3631468 | 0,10336676 | 0,00044277 | 0,00729117 | KAT6B |
| 815,135436 | -0,3631812 | 0,10723309 | 0,00070703 | 0,01034937 | RDX |
| 1518,16074 | -0,3633133 | 0,14919939 | 0,01488827 | 0,0911064 | PRKXP1 |
| 1033,68699 | -0,3634411 | 0,12752791 | 0,00437338 | 0,03946049 | BST2 |
| 311,558162 | -0,3639071 | 0,14524582 | 0,01222934 | 0,07966248 | ETFDH |
| 1362,539 | -0,3640404 | 0,0834747 | 1,29E-05 | 0,00043417 | UBE2Q1 |
| 935,43923 | -0,3640461 | 0,10055885 | 0,00029434 | 0,00530404 | RIOX2 |
| 704,660413 | -0,3640812 | 0,0998997 | 0,00026795 | 0,00494615 | TPMT |
| 1457,78229 | -0,3640924 | 0,12255597 | 0,00297001 | 0,02979612 | ATXN2 |
| 1815,23493 | -0,3642862 | 0,0951932 | 0,00012981 | 0,00278366 | HARS1 |
| 3270,16325 | -0,3644934 | 0,09670161 | 0,00016373 | 0,00336008 | MORF4L1 |
| 274,959366 | -0,3654163 | 0,15201224 | 0,01622295 | 0,09617778 | SPATA7 |
| 683,649953 | -0,3654504 | 0,13038675 | 0,00506581 | 0,04384733 | BEX3 |
| 607,598559 | -0,3654713 | 0,09721231 | 0,00017024 | 0,00347891 | CRNKL1 |
| 5646,7393 | -0,3655396 | 0,14912819 | 0,014239 | 0,08824634 | RPL34 |
| 330,906805 | -0,3655453 | 0,1522476 | 0,01635069 | 0,09667187 | SIMC1P1 |
| 1188,86198 | -0,3656375 | 0,11124192 | 0,00101308 | 0,01360598 | ZMYND8 |
| 7345,49538 | -0,3658131 | 0,10123815 | 0,00030222 | 0,0053958 | UBB |
| 262,931023 | -0,3658701 | 0,13498505 | 0,00671922 | 0,05317203 | UPF3B |
| 284,615318 | -0,366239 | 0,14770715 | 0,0131569 | 0,08336965 | CDIN1 |
| 652,134593 | -0,3666327 | 0,12739298 | 0,00400248 | 0,03699606 | NUCB2 |
| 1404,84774 | -0,3667681 | 0,10528734 | 0,00049491 | 0,00794923 | USP8 |
| 219,602881 | -0,3671968 | 0,15569033 | 0,01834859 | 0,1042877 | ANKMY2 |
| 1133,3611 | -0,3672541 | 0,11909886 | 0,00204507 | 0,02280864 | PROSER3 |
| 1249,59576 | -0,3674806 | 0,1252212 | 0,00333923 | 0,03241739 | PSMC6 |
| 9287,60372 | -0,367497 | 0,13847753 | 0,00795819 | 0,05954014 | RPL7 |
| 3251,87256 | -0,3677102 | 0,08786122 | 2,85E-05 | 0,00082583 | ADAM22 |
| 2617,04925 | -0,3678125 | 0,12055748 | 0,00228133 | 0,02466578 | CALU |
| 1610,31323 | -0,3680366 | 0,10679105 | 0,00056827 | 0,00877731 | RND3 |
| 439,334824 | -0,3681241 | 0,13803067 | 0,00765378 | 0,05815646 | TRAF3IP1 |
| 180,942273 | -0,3681773 | 0,16843931 | 0,02882908 | 0,1404777 | NIFK-AS1 |
| 1480,98422 | -0,3682236 | 0,12535316 | 0,00330882 | 0,03225309 | PREPL |
| 1126,00373 | -0,3682733 | 0,10731937 | 0,00060011 | 0,00911292 | CWC22 |
| 737,197762 | -0,3684435 | 0,12256257 | 0,00264564 | 0,0275972 | PAK1IP1 |
| 6928,19999 | -0,368636 | 0,0968739 | 0,00014162 | 0,00298401 | SYNCRIP |
| 1220,78885 | -0,3687723 | 0,10643191 | 0,0005305 | 0,00837156 | PSMC5 |
| 1001,68415 | -0,3689532 | 0,1120654 | 0,00099371 | 0,0134108 | AATF |

| | | | | | |
|---|---|---|---|---|---|
| 1070,28108 | -0,369108 | 0,13459384 | 0,0060995 | 0,04998208 | SKA2 |
| 288,021005 | -0,3691371 | 0,15654433 | 0,01837193 | 0,10435766 | CIR1 |
| 1459,20252 | -0,3693431 | 0,10099683 | 0,00025521 | 0,00475175 | EFCAB13 |
| 522,483828 | -0,3693894 | 0,16645784 | 0,02647876 | 0,13264488 | SCARNA12 |
| 174,468817 | -0,3695436 | 0,18407191 | 0,04468534 | 0,18809595 | FTLP10 |
| 1826,9385 | -0,369603 | 0,11244955 | 0,0010132 | 0,01360598 | USO1 |
| 982,833286 | -0,3696632 | 0,12983969 | 0,00441231 | 0,03969817 | WDHD1 |
| 2417,58789 | -0,3700541 | 0,11144018 | 0,00089807 | 0,0124213 | TAX1BP1 |
| 3280,15388 | -0,3706619 | 0,08642323 | 1,80E-05 | 0,00056507 | CAST |
| 1595,61904 | -0,3708854 | 0,11651646 | 0,00145699 | 0,01790112 | TTLL5 |
| 282,373012 | -0,3710835 | 0,17444901 | 0,03340558 | 0,15435758 | EEF1DP3 |
| 721,87745 | -0,3712579 | 0,13197865 | 0,00490794 | 0,04285407 | DMD |
| 4283,51102 | -0,3712877 | 0,11131505 | 0,00085156 | 0,01196171 | SUPT16H |
| 266,135183 | -0,3717546 | 0,15746156 | 0,01822953 | 0,10386056 | SUGCT |
| 75465,6606 | -0,3721997 | 0,11412968 | 0,00110942 | 0,01452619 | SERPINA1 |
| 3456,13475 | -0,372277 | 0,09804457 | 0,00014645 | 0,00306858 | PLSCR1 |
| 913,068886 | -0,3723878 | 0,12099652 | 0,00208623 | 0,02312575 | ACBD6 |
| 436,957882 | -0,3724013 | 0,16682093 | 0,02559239 | 0,12990526 | PSMB9 |
| 446,942475 | -0,3725116 | 0,17797424 | 0,03634337 | 0,1634933 | HMMR |
| 298,510762 | -0,3732024 | 0,17544653 | 0,03340702 | 0,15435758 | LRRTM2 |
| 733,989216 | -0,3734098 | 0,13191031 | 0,00464338 | 0,04126701 | LINC00997 |
| 653,939096 | -0,3739195 | 0,1508713 | 0,0131973 | 0,08356968 | MRPL32 |
| 2894,70779 | -0,3739993 | 0,09511304 | 8,42E-05 | 0,00198617 | PTPN2 |
| 419,505139 | -0,3741387 | 0,11919469 | 0,00169591 | 0,02003014 | SNAPC3 |
| 3004,02439 | -0,3743726 | 0,08471281 | 9,90E-06 | 0,00034693 | SIPA1L2 |
| 343,115617 | -0,3745259 | 0,1739809 | 0,03134353 | 0,14821798 | CEP162 |
| 302,770109 | -0,3746164 | 0,14432722 | 0,00944245 | 0,06686403 | TIMM8A |
| 1406,59722 | -0,3749594 | 0,10396387 | 0,0003102 | 0,00550844 | PROSER1 |
| 933,240042 | -0,3765679 | 0,10260122 | 0,00024235 | 0,00456729 | PWAR5 |
| 202,222976 | -0,3770451 | 0,15702432 | 0,01634189 | 0,09665673 | DIAPH2 |
| 6369,18382 | -0,3775318 | 0,09273456 | 4,68E-05 | 0,00122457 | ANKRD36C |
| 1694,98502 | -0,3777843 | 0,10771234 | 0,0004526 | 0,00742088 | SEC11A |
| 2182,95376 | -0,3778327 | 0,10456597 | 0,00030227 | 0,0053958 | MYL6 |
| 964,416134 | -0,3778833 | 0,14218817 | 0,00786926 | 0,05915506 | ATP6V1G1 |
| 1489,24419 | -0,3779647 | 0,14432987 | 0,00882511 | 0,06392798 | RTRAF |
| 548,37105 | -0,3779651 | 0,10874188 | 0,00050933 | 0,00810517 | OTUD6B |
| 435,572059 | -0,3780172 | 0,14344893 | 0,00840868 | 0,06181064 | SPC25 |
| 216,617759 | -0,3782479 | 0,18058879 | 0,03621314 | 0,1630735 | SULT1B1 |
| 203,659105 | -0,3792361 | 0,17444965 | 0,02971264 | 0,14305163 | LOC105369140 |
| 1144,84825 | -0,3797374 | 0,1155027 | 0,00101012 | 0,01358384 | ADH5 |
| 430,047128 | -0,3798241 | 0,11341987 | 0,00081153 | 0,01156248 | DAAM1 |
| 2142,81098 | -0,3800437 | 0,09972901 | 0,00013854 | 0,00292881 | PLP2 |
| 1978,53832 | -0,3800664 | 0,13539397 | 0,00499874 | 0,04342569 | ZC3H15 |
| 3426,88955 | -0,3804227 | 0,10669695 | 0,00036323 | 0,0062479 | RABGGTB |
| 509,177454 | -0,3804897 | 0,16202497 | 0,01885732 | 0,10593805 | CHMP5 |
| 1924,36074 | -0,3806594 | 0,11271001 | 0,00073192 | 0,01062739 | METAP2 |
| 329,85774 | -0,3817218 | 0,11861353 | 0,00128999 | 0,01624799 | TTF1 |
| 427,339966 | -0,3823281 | 0,14030364 | 0,00643005 | 0,05170547 | IER3IP1 |

| | | | | | |
|---|---|---|---|---|---|
| 797,707465 | -0,3823805 | 0,10585662 | 0,00030355 | 0,0054133 | RPAP3 |
| 1330,73348 | -0,3828998 | 0,12881491 | 0,00295404 | 0,02971657 | CLNS1A |
| 509,679881 | -0,3836527 | 0,1186334 | 0,00122097 | 0,01563294 | OSBPL1A |
| 1019,56497 | -0,383834 | 0,1558122 | 0,0137611 | 0,08621649 | MRPL9 |
| 601,518162 | -0,3840391 | 0,11104464 | 0,00054335 | 0,00851762 | RIOK1 |
| 3785,52201 | -0,3840952 | 0,10704426 | 0,00033298 | 0,00585478 | RPS13 |
| 205,827629 | -0,3842266 | 0,15556183 | 0,01351402 | 0,08500623 | SMIM10 |
| 246,956424 | -0,3846534 | 0,15065762 | 0,01067497 | 0,07229099 | TTC9C |
| 235,498748 | -0,3847697 | 0,19601343 | 0,04964892 | 0,2016328 | IL26 |
| 493,792559 | -0,3854164 | 0,15092724 | 0,01065982 | 0,072266 | LINC00174 |
| 347,71911 | -0,3855931 | 0,14461746 | 0,00766919 | 0,05816328 | NBDY |
| 283,839823 | -0,38564 | 0,18547458 | 0,03759836 | 0,16758508 | ZNF41 |
| 765,793413 | -0,3858856 | 0,12065366 | 0,00138244 | 0,01716353 | SCAPER |
| 1101,67779 | -0,3860591 | 0,14303897 | 0,00695526 | 0,05458337 | TNFSF10 |
| 8684,43017 | -0,3864563 | 0,0979515 | 7,97E-05 | 0,00190098 | MFF |
| 284,693975 | -0,386659 | 0,14929245 | 0,00959918 | 0,06766989 | CFDP1 |
| 1526,31217 | -0,3866598 | 0,13892385 | 0,00538173 | 0,04572416 | ZFP36 |
| 1645,47743 | -0,3870797 | 0,11969468 | 0,00122116 | 0,01563294 | SMC5 |
| 505,309876 | -0,3874481 | 0,13884523 | 0,00526261 | 0,04501529 | UTP14C |
| 1548,0701 | -0,3876152 | 0,10108361 | 0,00012577 | 0,00272455 | OAT |
| 394,070366 | -0,3877638 | 0,13410352 | 0,00383376 | 0,03580387 | C1orf52 |
| 268,920084 | -0,3878753 | 0,14505939 | 0,00749734 | 0,05738555 | POLR2L |
| 430,975294 | -0,3880865 | 0,15142442 | 0,01038004 | 0,07103098 | UFSP2 |
| 2248,00912 | -0,3889681 | 0,09877003 | 8,21E-05 | 0,00194474 | RBM8A |
| 497,245045 | -0,3892396 | 0,12539676 | 0,00190882 | 0,02179647 | MICOS10 |
| 764,317561 | -0,3893122 | 0,09592927 | 4,94E-05 | 0,00128283 | SART1 |
| 905,190593 | -0,3893526 | 0,12888924 | 0,00252082 | 0,02651274 | BRD10 |
| 1090,59822 | -0,3893666 | 0,11933909 | 0,0011036 | 0,01446062 | MIR1244-2 |
| 502,079946 | -0,3894001 | 0,13012309 | 0,00276655 | 0,02854342 | ZCRB1 |
| 713,046317 | -0,3898884 | 0,13646486 | 0,00427584 | 0,03882106 | SPDL1 |
| 511,969182 | -0,3905079 | 0,14900669 | 0,0087739 | 0,06365452 | RALB |
| 358,794038 | -0,3906091 | 0,14714981 | 0,00794262 | 0,05947065 | ANKRD18B |
| 286,912644 | -0,3907191 | 0,19380209 | 0,04379236 | 0,18545105 | FRG1 |
| 415,067605 | -0,3907883 | 0,15555715 | 0,01199863 | 0,07863735 | POLE4 |
| 244,349668 | -0,3908794 | 0,16108604 | 0,01524417 | 0,09244711 | MAP9 |
| 152,677221 | -0,3908854 | 0,18201513 | 0,03175005 | 0,14947567 | NUDT11 |
| 4365,32052 | -0,3913343 | 0,10783226 | 0,00028441 | 0,00516438 | BDP1 |
| 555,2892 | -0,3914356 | 0,13998344 | 0,00516915 | 0,04447145 | NDUFA7 |
| 620,954771 | -0,3916228 | 0,14942738 | 0,00877176 | 0,06365452 | PROS1 |
| 1589,45118 | -0,3919038 | 0,10487113 | 0,00018623 | 0,00373657 | COX4I1 |
| 1629,26508 | -0,392587 | 0,11491781 | 0,00063492 | 0,00951925 | HAT1 |
| 243,878871 | -0,3929919 | 0,14599458 | 0,00710621 | 0,05535474 | XAF1 |
| 230,864899 | -0,3934912 | 0,153992 | 0,01061054 | 0,07200931 | CISD1 |
| 8152,49678 | -0,3939914 | 0,13029371 | 0,00249563 | 0,02629165 | RPS15A |
| 751,869961 | -0,3941567 | 0,11860484 | 0,00088967 | 0,01237748 | HMGB3 |
| 13590,7411 | -0,394391 | 0,10266512 | 0,00012227 | 0,00267638 | HSPD1 |
| 1417,44894 | -0,3944255 | 0,09433665 | 2,90E-05 | 0,00083568 | CDC5L |
| 241,87427 | -0,3948135 | 0,15513989 | 0,01093131 | 0,07350084 | COQ7 |

| | | | | | |
|---|---|---|---|---|---|
| 827,776054 | -0,3949139 | 0,11604392 | 0,00066616 | 0,00988228 | MRPS23 |
| 322,202781 | -0,3950539 | 0,16570035 | 0,01711878 | 0,09945292 | LMLN |
| 419,457805 | -0,3955381 | 0,13384397 | 0,00312448 | 0,03086766 | HCCS |
| 369,039487 | -0,3957445 | 0,14698496 | 0,00709366 | 0,05530253 | POPDC3 |
| 7015,69147 | -0,3958783 | 0,10361788 | 0,00013315 | 0,00284026 | CTSL |
| 1024,6232 | -0,3958837 | 0,11145762 | 0,00038249 | 0,00650826 | MTIF2 |
| 284,555391 | -0,396484 | 0,17101336 | 0,0204255 | 0,11192241 | GPN3 |
| 214,513885 | -0,3965025 | 0,17206626 | 0,02120246 | 0,11465327 | ZNF182 |
| 743,345792 | -0,3965484 | 0,13626672 | 0,00361325 | 0,03440161 | ACADM |
| 926,950163 | -0,3965962 | 0,10475084 | 0,00015304 | 0,00317499 | H4C3 |
| 466,253919 | -0,3966493 | 0,14539549 | 0,00637057 | 0,0514239 | RPGRIP1L |
| 221,268115 | -0,3967364 | 0,17481851 | 0,02324289 | 0,12182235 | APPAT |
| 887,021281 | -0,3968368 | 0,10481259 | 0,000153 | 0,00317499 | CBLL1 |
| 1260,25683 | -0,3968386 | 0,14944208 | 0,00791985 | 0,05939419 | NOP10 |
| 1645,82719 | -0,3971046 | 0,12352016 | 0,00130491 | 0,01639457 | NEMF |
| 186,831214 | -0,3971223 | 0,17991071 | 0,02729102 | 0,13574483 | CHMP4A |
| 1776,32673 | -0,3972745 | 0,13735297 | 0,00382358 | 0,0357369 | STEAP1 |
| 363,527949 | -0,3980827 | 0,11492447 | 0,00053248 | 0,00838883 | DCP1B |
| 798,622339 | -0,3980931 | 0,12566275 | 0,00153519 | 0,01862006 | ATG3 |
| 393,522276 | -0,3983996 | 0,11421266 | 0,00048624 | 0,0078499 | ZSCAN26 |
| 1123,91919 | -0,3984992 | 0,10927916 | 0,00026572 | 0,00491939 | FAM98A |
| 213,122825 | -0,3985536 | 0,18746914 | 0,03350582 | 0,15462527 | IQCH |
| 1383,08852 | -0,3988236 | 0,13595238 | 0,00335106 | 0,03248228 | ST18 |
| 452,082112 | -0,3988413 | 0,15619399 | 0,01066464 | 0,0722728 | HDDC2 |
| 404,77329 | -0,4001055 | 0,13051481 | 0,00217237 | 0,02375925 | ELP4 |
| 668,485244 | -0,4003017 | 0,14007047 | 0,00426509 | 0,03877934 | DUSP11 |
| 287,565306 | -0,4006761 | 0,17235979 | 0,0200907 | 0,1108232 | H2BC6 |
| 558,749467 | -0,4007147 | 0,12050561 | 0,0008833 | 0,01230706 | VCAN-AS1 |
| 5761,82205 | -0,4009534 | 0,10477963 | 0,00012991 | 0,00278366 | CCT8 |
| 251,065577 | -0,4010294 | 0,14483143 | 0,00562391 | 0,0471885 | PABIR1 |
| 1332,26626 | -0,4011034 | 0,12588907 | 0,00144172 | 0,01777123 | SERF2 |
| 569,800966 | -0,402154 | 0,13594755 | 0,00309486 | 0,03073903 | ZNF326 |
| 894,22481 | -0,4023218 | 0,0938906 | 1,83E-05 | 0,00057066 | UTP6 |
| 1052,58195 | -0,4023479 | 0,11487528 | 0,00046096 | 0,00753824 | SNRPA1 |
| 201,026262 | -0,4027146 | 0,17386447 | 0,02054429 | 0,11231391 | SLC26A10P |
| 622,983564 | -0,4031958 | 0,12171795 | 0,00092452 | 0,01273136 | POP4 |
| 420,212025 | -0,4032688 | 0,12690149 | 0,00148392 | 0,01812607 | STX4 |
| 3442,66305 | -0,4036876 | 0,09533523 | 2,29E-05 | 0,00069265 | NIN |
| 7992,36752 | -0,4038236 | 0,12976723 | 0,00185883 | 0,0213935 | NPIPB5 |
| 757,056783 | -0,4040291 | 0,13157807 | 0,002136 | 0,0234699 | ZNHIT6 |
| 1108,5445 | -0,4046015 | 0,10647505 | 0,00014472 | 0,00303567 | NADK2 |
| 6820,26954 | -0,404996 | 0,10581603 | 0,00012953 | 0,00278178 | SNRPD2 |
| 818,28054 | -0,4053042 | 0,13326862 | 0,00235592 | 0,02519866 | NRP1 |
| 528,872748 | -0,4055924 | 0,11886087 | 0,0006441 | 0,00960368 | UXT |
| 455,438837 | -0,4059317 | 0,15255437 | 0,00779324 | 0,05870939 | SAAL1 |
| 8734,83126 | -0,4063002 | 0,12956273 | 0,00171308 | 0,02014491 | CDK6-AS1 |
| 234,158246 | -0,4063663 | 0,15985265 | 0,01101791 | 0,07397796 | PABPN1 |
| 5081,2112 | -0,4064623 | 0,10716237 | 0,00014886 | 0,00311232 | ZFAS1 |

| | | | | | |
|---|---|---|---|---|---|
| 1862,63231 | -0,4077361 | 0,07838671 | 1,98E-07 | 1,28E-05 | WBP11 |
| 233,175922 | -0,408305 | 0,1589907 | 0,01022558 | 0,07025352 | RPP40 |
| 17369,647 | -0,4084481 | 0,10432962 | 9,04E-05 | 0,00209388 | AKAP9 |
| 1702,84047 | -0,408887 | 0,11578553 | 0,00041334 | 0,00694566 | MIA3 |
| 189,793059 | -0,4089026 | 0,16812591 | 0,01501069 | 0,09150039 | SAP30 |
| 1609,70391 | -0,4091364 | 0,17960111 | 0,0227249 | 0,11973765 | IFITM2 |
| 306,402474 | -0,4093242 | 0,1523823 | 0,00722771 | 0,05601827 | SLC10A7 |
| 1321,55318 | -0,4093911 | 0,15998348 | 0,01049862 | 0,07156625 | BUB1B |
| 486,546874 | -0,4094472 | 0,14685965 | 0,00530317 | 0,04517843 | EPDR1 |
| 306,031856 | -0,4097429 | 0,1298641 | 0,00160406 | 0,01925789 | COPRS |
| 8885,65891 | -0,4097841 | 0,0979796 | 2,89E-05 | 0,00083346 | IQGAP1 |
| 1835,48866 | -0,4099778 | 0,14494554 | 0,00467674 | 0,04148551 | CST7 |
| 2937,99338 | -0,4101013 | 0,10736278 | 0,00013358 | 0,00284614 | SNHG5 |
| 847,791975 | -0,4101734 | 0,09480338 | 1,51E-05 | 0,00049068 | STXBP1 |
| 481,475827 | -0,4108929 | 0,17959756 | 0,02214605 | 0,11747716 | PRIM1 |
| 225,182075 | -0,4113222 | 0,18894942 | 0,02948874 | 0,14248124 | MORN4 |
| 368,75018 | -0,4118305 | 0,13039614 | 0,00158691 | 0,01908826 | KIF27 |
| 5540,55052 | -0,412166 | 0,13084364 | 0,00163234 | 0,01951073 | HMGB1 |
| 319,112263 | -0,4122044 | 0,16439628 | 0,01216288 | 0,07938388 | FRA10AC1 |
| 596,791047 | -0,4124277 | 0,16799058 | 0,01408594 | 0,08755589 | ZNF721 |
| 595,001009 | -0,413528 | 0,11942818 | 0,00053505 | 0,0084197 | MYB |
| 3272,01725 | -0,4136255 | 0,09163448 | 6,37E-06 | 0,00023851 | RSL1D1 |
| 523,936652 | -0,4143512 | 0,12867484 | 0,00128127 | 0,01618356 | SRP19 |
| 533,816454 | -0,4146112 | 0,1492299 | 0,00546376 | 0,04623425 | COX6C |
| 222,458987 | -0,4149709 | 0,18516808 | 0,02502284 | 0,12771975 | TMEM116 |
| 1578,89965 | -0,4153429 | 0,0999656 | 3,25E-05 | 0,00091645 | SH3KBP1 |
| 318,81394 | -0,4156046 | 0,19889388 | 0,03665559 | 0,16462387 | NDUFA8 |
| 1632,50476 | -0,4157097 | 0,09278528 | 7,45E-06 | 0,00027334 | SF3B4 |
| 121,54009 | -0,4157618 | 0,2099768 | 0,04769937 | 0,19665846 | DHFR2 |
| 123,414984 | -0,4162913 | 0,18058771 | 0,02115521 | 0,11451083 | EEF1AKMT1 |
| 3021,39868 | -0,4164589 | 0,11823472 | 0,00042781 | 0,00711925 | KTN1 |
| 979,527081 | -0,4168629 | 0,12583071 | 0,00092338 | 0,01272492 | MBD2 |
| 191,555722 | -0,4168744 | 0,16041169 | 0,00935563 | 0,06634852 | NOPCHAP1 |
| 2118,45214 | -0,4168821 | 0,08838564 | 2,40E-06 | 0,00010406 | DDX42 |
| 2220,15628 | -0,4174109 | 0,08885565 | 2,63E-06 | 0,00011116 | ZC3H13 |
| 1335,88482 | -0,4175214 | 0,12486307 | 0,00082629 | 0,01171105 | H4C2 |
| 1088,32148 | -0,4177906 | 0,09992292 | 2,90E-05 | 0,00083568 | ANKRD26 |
| 17696,0655 | -0,4181223 | 0,13201954 | 0,00153954 | 0,01864898 | RPL8 |
| 633,424031 | -0,4185463 | 0,12963666 | 0,00124393 | 0,01587081 | ATF1 |
| 572,687435 | -0,4188959 | 0,14851748 | 0,00479466 | 0,04231328 | CEP128 |
| 1235,30574 | -0,4189503 | 0,08906068 | 2,55E-06 | 0,00010818 | GOLM1 |
| 1526,14174 | -0,4190372 | 0,09969087 | 2,63E-05 | 0,00077383 | TRIM44 |
| 356,783211 | -0,4190467 | 0,14120998 | 0,0030019 | 0,0300206 | SH3BP5 |
| 301,817342 | -0,4193283 | 0,15480692 | 0,00675438 | 0,05338326 | CHMP2A |
| 991,097532 | -0,4195472 | 0,12781479 | 0,00102905 | 0,01379916 | STAU2 |
| 4900,78719 | -0,4199543 | 0,11896253 | 0,00041534 | 0,00696608 | LSM12 |
| 562,786616 | -0,4202118 | 0,11074804 | 0,00014806 | 0,00309889 | DPY30 |
| 618,154037 | -0,4205507 | 0,13552284 | 0,00191458 | 0,02182104 | PIBF1 |

| | | | | | |
|---|---|---|---|---|---|
| 3276,58411 | -0,4212894 | 0,09571825 | 1,08E-05 | 0,00037077 | SAT1 |
| 18991,9946 | -0,4213113 | 0,11051893 | 0,00013778 | 0,00291605 | RPS20 |
| 1655,03312 | -0,4217207 | 0,08516548 | 7,35E-07 | 3,81E-05 | CBX1 |
| 1846,35262 | -0,4217497 | 0,15978614 | 0,00830373 | 0,06132505 | RPL38 |
| 4238,23444 | -0,4219011 | 0,14194737 | 0,00295639 | 0,02971657 | NPIPB3 |
| 4520,85107 | -0,4224481 | 0,12131649 | 0,00049731 | 0,00798108 | CENPF |
| 177,025364 | -0,4225566 | 0,17485665 | 0,0156667 | 0,09425428 | TMEM60 |
| 139,876553 | -0,422717 | 0,1963238 | 0,03130593 | 0,14815896 | MAP4K3-DT |
| 536,28817 | -0,423405 | 0,10489459 | 5,43E-05 | 0,00138547 | HMGN3 |
| 527,748738 | -0,4234434 | 0,10719655 | 7,81E-05 | 0,00187053 | THG1L |
| 152,350879 | -0,424187 | 0,19182494 | 0,02701346 | 0,13461198 | RBKS |
| 366,31075 | -0,426161 | 0,16681387 | 0,01062763 | 0,07209947 | RBIS |
| 273,327884 | -0,4262683 | 0,17251713 | 0,01347828 | 0,08483783 | SLC12A2-DT |
| 573,922833 | -0,4262906 | 0,12274324 | 0,00051462 | 0,00816195 | SEC61B |
| 603,445732 | -0,4266564 | 0,13679238 | 0,00181462 | 0,0210511 | ZWINT |
| 1282,99718 | -0,4269893 | 0,09638004 | 9,41E-06 | 0,00033286 | MDC1 |
| 146,245704 | -0,4275339 | 0,20296921 | 0,03516983 | 0,15969483 | KRT18 |
| 394,32731 | -0,4276581 | 0,14218931 | 0,00263261 | 0,02748988 | GTF3C6 |
| 505,172323 | -0,4278795 | 0,14032186 | 0,00229397 | 0,02474583 | LARP7 |
| 226,638561 | -0,4281746 | 0,18946309 | 0,02382516 | 0,12371385 | H4C16 |
| 491,275673 | -0,4283519 | 0,13379551 | 0,00136694 | 0,01703347 | DYNLL2 |
| 172,15173 | -0,4287469 | 0,16080925 | 0,0076718 | 0,05816328 | MAPDA |
| 5990,11215 | -0,4289473 | 0,13684864 | 0,00172168 | 0,02023346 | RPS15 |
| 506,113889 | -0,4296234 | 0,17231716 | 0,01265929 | 0,08136571 | DRAP1 |
| 669,493227 | -0,4296302 | 0,11083785 | 0,0001061 | 0,00238987 | ALYREF |
| 677,966985 | -0,4296533 | 0,10487257 | 4,19E-05 | 0,00111785 | HIBCH |
| 2833,18622 | -0,429706 | 0,14121208 | 0,00234246 | 0,02509717 | SUMO2 |
| 388,393049 | -0,4301086 | 0,16472582 | 0,00902626 | 0,06486363 | TPRKB |
| 556,636146 | -0,4320725 | 0,19594265 | 0,02744733 | 0,13612452 | LOC101929950 |
| 103,91715 | -0,4320968 | 0,21860494 | 0,04808568 | 0,19753132 | STOX1 |
| 1756,07521 | -0,4323083 | 0,13709054 | 0,00161352 | 0,01934694 | SUMO1 |
| 644,761754 | -0,43281 | 0,15299339 | 0,00467017 | 0,04146608 | PDCD10 |
| 2772,84242 | -0,4335245 | 0,09801609 | 9,73E-06 | 0,00034235 | CCAR1 |
| 1011,87921 | -0,4337625 | 0,11410309 | 0,00014382 | 0,00302368 | H4C9 |
| 1889,94592 | -0,434106 | 0,14225358 | 0,00227599 | 0,02463611 | EIF3J |
| 3218,64753 | -0,4342364 | 0,14490397 | 0,00272902 | 0,02826265 | SEPTIN7 |
| 737,712759 | -0,4351543 | 0,1081693 | 5,75E-05 | 0,00144447 | TTC1 |
| 448,344191 | -0,4354503 | 0,1446476 | 0,00260885 | 0,02730202 | SNTB1 |
| 7978,52247 | -0,4354861 | 0,12638456 | 0,00056954 | 0,00878254 | IFI16 |
| 205,218659 | -0,4356302 | 0,14974517 | 0,00362421 | 0,03447642 | C18orf21 |
| 457,187487 | -0,4363191 | 0,14256006 | 0,00220895 | 0,02410357 | ETAA1 |
| 794,122686 | -0,4379022 | 0,13403992 | 0,00108711 | 0,01432794 | GLRX5 |
| 1152,66364 | -0,4380519 | 0,1206354 | 0,0002821 | 0,00513727 | SMIM15 |
| 2046,4602 | -0,4387318 | 0,0859583 | 3,33E-07 | 2,00E-05 | BRWD3 |
| 135,706797 | -0,4392215 | 0,1894647 | 0,02043718 | 0,11192241 | ZNF649 |
| 615,713276 | -0,4392622 | 0,14887687 | 0,00317253 | 0,03128059 | CCDC43 |
| 2159,2801 | -0,4392757 | 0,12145788 | 0,00029839 | 0,00535663 | ANP32A |
| 1056,12211 | -0,4393523 | 0,12082489 | 0,00027661 | 0,00506756 | LOC729218 |

| | | | | | |
|---|---|---|---|---|---|
| 1254,28191 | -0,4394861 | 0,09513982 | 3,85E-06 | 0,00015462 | EID1 |
| 565,608005 | -0,4400542 | 0,1112964 | 7,69E-05 | 0,00184388 | UQCR10 |
| 865,627493 | -0,440174 | 0,1082567 | 4,78E-05 | 0,00124635 | RBM34 |
| 628,476686 | -0,4404207 | 0,13146366 | 0,00080773 | 0,01151698 | SIMC1 |
| 1669,22544 | -0,4405742 | 0,10649702 | 3,52E-05 | 0,00097211 | FKBP5 |
| 556,943974 | -0,4416915 | 0,1421049 | 0,00188226 | 0,02159745 | RPF2 |
| 592,869774 | -0,4419594 | 0,11821968 | 0,00018515 | 0,00372687 | LINC00857 |
| 2810,75916 | -0,442118 | 0,13991942 | 0,00157875 | 0,01902648 | SMN1 |
| 816,757537 | -0,4422445 | 0,16752887 | 0,00829524 | 0,06131024 | PRDX5 |
| 369,76288 | -0,4426689 | 0,14047225 | 0,00162551 | 0,01944137 | NDUFB6 |
| 998,390477 | -0,4427704 | 0,14895309 | 0,00295338 | 0,02971657 | UBA3 |
| 353,26776 | -0,4427792 | 0,14625361 | 0,00246606 | 0,02608174 | MTIF3 |
| 195,705172 | -0,4436084 | 0,15561292 | 0,00436208 | 0,03937731 | YAE1 |
| 3562,50612 | -0,4437536 | 0,10969423 | 5,22E-05 | 0,00134483 | NREP |
| 581,886929 | -0,4438661 | 0,13469942 | 0,0009834 | 0,01331915 | HSDL2 |
| 399,407478 | -0,444293 | 0,12887986 | 0,00056613 | 0,00876737 | BTN3A2 |
| 569,923967 | -0,4446552 | 0,15098027 | 0,00322828 | 0,03164885 | GTF2A2 |
| 1315,98873 | -0,4462514 | 0,10961351 | 4,68E-05 | 0,00122457 | CCNK |
| 1180,93356 | -0,4469403 | 0,10632699 | 2,63E-05 | 0,00077383 | NDUFV2 |
| 115,413851 | -0,446976 | 0,22707819 | 0,04902441 | 0,20004114 | LUZP2 |
| 216,339629 | -0,4474763 | 0,14776388 | 0,00245917 | 0,0260235 | NUDT9 |
| 2010,21933 | -0,4478935 | 0,12483591 | 0,00033341 | 0,00585478 | ERH |
| 1043,20662 | -0,4479114 | 0,11662115 | 0,00012266 | 0,00267698 | VRK1 |
| 240,445294 | -0,4483987 | 0,1505631 | 0,00289999 | 0,02944967 | BBIP1 |
| 2935,56449 | -0,4494101 | 0,13927029 | 0,00125143 | 0,01595573 | ZNF146 |
| 3722,19822 | -0,4505066 | 0,12424518 | 0,0002879 | 0,00520288 | SUB1 |
| 18320,0824 | -0,4511485 | 0,09784938 | 4,01E-06 | 0,0001599 | PRRC2C |
| 276,064982 | -0,4512447 | 0,18158479 | 0,01295384 | 0,08245733 | KIAA1549L |
| 254,188508 | -0,4518099 | 0,15911238 | 0,00451751 | 0,04031882 | BEND6 |
| 85,0063677 | -0,4521937 | 0,22982659 | 0,04912065 | 0,20028572 | HOXB7 |
| 1072,07244 | -0,4525513 | 0,12858346 | 0,00043234 | 0,00716939 | ACAT1 |
| 2520,55872 | -0,4533812 | 0,15115686 | 0,00270504 | 0,02807577 | PSMA4 |
| 406,049195 | -0,4540842 | 0,13519716 | 0,00078315 | 0,0112343 | KNOP1 |
| 1369,98608 | -0,4540886 | 0,09082474 | 5,74E-07 | 3,12E-05 | KCNC3 |
| 2143,19695 | -0,4546553 | 0,1027367 | 9,62E-06 | 0,00033912 | EPB41L2 |
| 137,998441 | -0,4548545 | 0,18320977 | 0,01303916 | 0,08281771 | ZNF419 |
| 391,207032 | -0,4555475 | 0,13952719 | 0,00109488 | 0,01440118 | RSRC1 |
| 837,612456 | -0,4556347 | 0,12937208 | 0,00042848 | 0,00712407 | UQCRQ |
| 4040,16011 | -0,4557696 | 0,13465698 | 0,00071264 | 0,01041219 | RPS28 |
| 270,594372 | -0,4561423 | 0,14657867 | 0,0018587 | 0,0213935 | THYN1 |
| 1185,18901 | -0,4566249 | 0,11326732 | 5,54E-05 | 0,00140592 | UPF2 |
| 108,279434 | -0,4573413 | 0,21610499 | 0,03431997 | 0,15723202 | POGLUT2 |
| 679,768828 | -0,4580992 | 0,14327905 | 0,00138744 | 0,01720296 | KNSTRN |
| 439,330551 | -0,4581352 | 0,11361815 | 5,52E-05 | 0,00140317 | ACOT13 |
| 364,261659 | -0,4583006 | 0,14445123 | 0,00151028 | 0,0184005 | ATP5PF |
| 827,608375 | -0,4586293 | 0,09602041 | 1,78E-06 | 8,13E-05 | MAP7D3 |
| 7101,17786 | -0,45872 | 0,22488343 | 0,041369 | 0,17834194 | PA2G4 |
| 992,632312 | -0,4587548 | 0,10954604 | 2,82E-05 | 0,00081874 | DCAF6 |

| | | | | | |
|---|---|---|---|---|---|
| 724,320051 | -0,4591769 | 0,13046485 | 0,00043229 | 0,00716939 | MRPL18 |
| 166,120053 | -0,4594279 | 0,19030595 | 0,01577179 | 0,09458577 | KRCC1 |
| 4028,68502 | -0,4601452 | 0,12309478 | 0,00018539 | 0,00372767 | S100A10 |
| 121,934007 | -0,4608437 | 0,22381275 | 0,03948858 | 0,17323187 | ALDH1L2 |
| 668,127344 | -0,4613042 | 0,14786716 | 0,00181023 | 0,02101307 | F5 |
| 342,756674 | -0,4615071 | 0,13964527 | 0,00095026 | 0,0129804 | SLX4IP |
| 575,82111 | -0,461662 | 0,14094824 | 0,00105512 | 0,01407894 | FAM162A |
| 198,632933 | -0,4616854 | 0,17670483 | 0,00898172 | 0,0645926 | PSG8-AS1 |
| 216,656142 | -0,4631344 | 0,1474214 | 0,00168046 | 0,01989732 | TIMM21 |
| 848,291094 | -0,4632712 | 0,1952371 | 0,01765074 | 0,10138728 | NMI |
| 351,626297 | -0,463312 | 0,14930539 | 0,00191494 | 0,02182104 | DNAJC19 |
| 1058,41362 | -0,4635972 | 0,10668829 | 1,39E-05 | 0,00045999 | SMC6 |
| 344,073931 | -0,4645937 | 0,17641601 | 0,00845067 | 0,06202406 | EMC2 |
| 597,541092 | -0,4650236 | 0,13738362 | 0,00071216 | 0,01041219 | CASP1 |
| 131,463518 | -0,465227 | 0,18878311 | 0,01372633 | 0,0860271 | RUNX1T1 |
| 993,844931 | -0,4660105 | 0,10928595 | 2,01E-05 | 0,0006184 | BRD7 |
| 316,818264 | -0,4661783 | 0,14103627 | 0,00094848 | 0,01296691 | MPHOSPH6 |
| 794,703462 | -0,4685111 | 0,10729079 | 1,26E-05 | 0,00042611 | TIMM13 |
| 2469,59154 | -0,4686303 | 0,15667065 | 0,002779 | 0,02862357 | H2BC21 |
| 713,007797 | -0,4686863 | 0,12666805 | 0,0002155 | 0,00416498 | CNPY2 |
| 160,779753 | -0,4710335 | 0,18189077 | 0,00960735 | 0,06770228 | CCPG1 |
| 7569,15256 | -0,4723454 | 0,09072467 | 1,93E-07 | 1,25E-05 | NOLC1 |
| 714,598112 | -0,4724561 | 0,14162761 | 0,00085022 | 0,01195178 | UTP14A |
| 738,902038 | -0,4727177 | 0,11321394 | 2,97E-05 | 0,00085263 | FGF9 |
| 599,067725 | -0,4727184 | 0,13000263 | 0,00027667 | 0,00506756 | CMAS |
| 203,599035 | -0,472719 | 0,15808373 | 0,00278697 | 0,02865881 | CCDC77 |
| 1189,73569 | -0,4727734 | 0,15546912 | 0,00235835 | 0,02521038 | COX7B |
| 647,012099 | -0,4730982 | 0,11448121 | 3,59E-05 | 0,0009824 | NDUFAB1 |
| 958,818707 | -0,4735006 | 0,12594662 | 0,00017023 | 0,00347891 | RAB27A |
| 177,653673 | -0,4745331 | 0,19830321 | 0,01671274 | 0,09808367 | CMC1 |
| 655,96272 | -0,4745483 | 0,16664525 | 0,00440425 | 0,03966345 | RAD51C |
| 1128,19695 | -0,4747556 | 0,1093137 | 1,41E-05 | 0,00046157 | TARS3 |
| 6993,40808 | -0,4751794 | 0,10603603 | 7,42E-06 | 0,0002731 | TPR |
| 1825,76882 | -0,4754404 | 0,100157 | 2,07E-06 | 9,26E-05 | NDUFB8 |
| 371,460987 | -0,4761659 | 0,19655342 | 0,01541082 | 0,09312942 | ZNF480 |
| 247,081482 | -0,4772235 | 0,20056151 | 0,01733909 | 0,10020556 | CDKN3 |
| 1102,58108 | -0,4773058 | 0,1281897 | 0,00019653 | 0,00388872 | CHCHD3 |
| 1996,78757 | -0,4777807 | 0,09451838 | 4,31E-07 | 2,43E-05 | BOD1L1 |
| 1721,20508 | -0,4780315 | 0,1078251 | 9,28E-06 | 0,00032991 | HLTF |
| 44822,346 | -0,4786125 | 0,07680117 | 4,61E-10 | 6,71E-08 | NAMPT |
| 29929,1268 | -0,4798986 | 0,10988312 | 1,26E-05 | 0,00042567 | RPS16 |
| 396,160021 | -0,4800329 | 0,17704414 | 0,0067005 | 0,05309977 | SAP30L-AS1 |
| 545,462543 | -0,4815796 | 0,17223423 | 0,00517276 | 0,04447145 | MPC2 |
| 4373,37442 | -0,4819586 | 0,12964254 | 0,00020113 | 0,00394356 | CASP4 |
| 514,906207 | -0,4855418 | 0,16563937 | 0,00337526 | 0,0327001 | CRIP1 |
| 211,78535 | -0,4882704 | 0,20442164 | 0,01691521 | 0,09876841 | PLAC4 |
| 107,355168 | -0,4884745 | 0,20762281 | 0,0186376 | 0,10532914 | DPP4 |
| 1037,19604 | -0,4889172 | 0,09834318 | 6,64E-07 | 3,53E-05 | BCCIP |

| | | | | | |
|---|---|---|---|---|---|
| 446,425438 | -0,4892509 | 0,13716279 | 0,00036118 | 0,00621827 | ECI2 |
| 198,253058 | -0,4896364 | 0,17367744 | 0,00481392 | 0,04234505 | CEP63 |
| 1135,91057 | -0,4896787 | 0,13643543 | 0,00033184 | 0,00584064 | THOC1 |
| 109,84179 | -0,4898414 | 0,20229853 | 0,01546176 | 0,09334778 | TMCO5A |
| 325,742854 | -0,4916244 | 0,15356393 | 0,00136747 | 0,01703347 | MFAP1 |
| 498,418055 | -0,4918455 | 0,12167758 | 5,30E-05 | 0,00135942 | PARP2 |
| 522,823133 | -0,4921216 | 0,11947104 | 3,80E-05 | 0,00102926 | ZFP62 |
| 856,565689 | -0,4926709 | 0,12175144 | 5,20E-05 | 0,00134 | MTX2 |
| 124,889876 | -0,4928538 | 0,23498086 | 0,03595631 | 0,16223259 | MACROH2A2 |
| 1799,58446 | -0,4928864 | 0,13639105 | 0,00030177 | 0,0053958 | GSTP1 |
| 112,871399 | -0,4934792 | 0,24664903 | 0,04542104 | 0,19026157 | SPDYE16 |
| 6981,57209 | -0,4947995 | 0,09764426 | 4,03E-07 | 2,33E-05 | HNRNPA3 |
| 447,462074 | -0,4950142 | 0,13257888 | 0,00018866 | 0,00377498 | OSTF1 |
| 13045,9403 | -0,4953781 | 0,12637311 | 8,86E-05 | 0,00206605 | NPM1 |
| 479,272444 | -0,4954296 | 0,10493668 | 2,34E-06 | 0,00010244 | TRMT6 |
| 931,076269 | -0,4963664 | 0,15146434 | 0,00104873 | 0,0140134 | CFH |
| 106,413306 | -0,4970321 | 0,21219472 | 0,01916328 | 0,10714787 | UACA |
| 1146,1791 | -0,4976708 | 0,10862445 | 4,61E-06 | 0,0001793 | CCDC47 |
| 405,751249 | -0,4981376 | 0,13047606 | 0,00013462 | 0,00286199 | TMEM258 |
| 1101,20217 | -0,4982132 | 0,1015433 | 9,28E-07 | 4,64E-05 | C5orf15 |
| 1921,83098 | -0,4982937 | 0,08134196 | 9,02E-10 | 1,17E-07 | MRFAP1 |
| 596,521716 | -0,4987974 | 0,11891401 | 2,73E-05 | 0,00079938 | ZC3H8 |
| 337,65722 | -0,4991785 | 0,15471608 | 0,00125351 | 0,01596071 | CWF19L2 |
| 72,2730831 | -0,4992671 | 0,25129665 | 0,04694858 | 0,19450715 | ZNF22-AS1 |
| 487,06957 | -0,5000836 | 0,13500452 | 0,00021206 | 0,00411526 | CHRAC1 |
| 16328,1794 | -0,5000904 | 0,08611941 | 6,36E-09 | 6,44E-07 | HNRNPA2B1 |
| 418,929985 | -0,5002838 | 0,1683283 | 0,00295797 | 0,02971657 | MITD1 |
| 355,981102 | -0,50067 | 0,21998053 | 0,02284757 | 0,12028348 | NSFP1 |
| 406,365162 | -0,5008596 | 0,15300825 | 0,00106256 | 0,01413832 | MRPL50 |
| 1812,268 | -0,5018216 | 0,11024581 | 5,32E-06 | 0,00020286 | AHSA1 |
| 14237,2942 | -0,5022764 | 0,08895824 | 1,64E-08 | 1,44E-06 | HNRNPU |
| 328,76668 | -0,502763 | 0,17184165 | 0,00343644 | 0,03302229 | AK6 |
| 750,211108 | -0,5032775 | 0,11646139 | 1,55E-05 | 0,00050058 | NDUFB4 |
| 561,343275 | -0,503333 | 0,15243294 | 0,00095999 | 0,01306421 | EIF1AY |
| 655,825351 | -0,5049058 | 0,13791079 | 0,00025113 | 0,00469994 | UTP3 |
| 272,731369 | -0,5049649 | 0,15564783 | 0,00117749 | 0,01519727 | FAM81A |
| 198,014564 | -0,5065764 | 0,16395788 | 0,00200376 | 0,02250042 | NDUFAF1 |
| 497,465754 | -0,5066134 | 0,14947834 | 0,00070095 | 0,01028107 | TAF3 |
| 82,0194841 | -0,506911 | 0,25067761 | 0,0431595 | 0,18364774 | MMAA |
| 519,502262 | -0,5071928 | 0,11274699 | 6,84E-06 | 0,00025439 | QNG1 |
| 80,543762 | -0,507536 | 0,25779908 | 0,04898444 | 0,19997776 | FAM228B |
| 268,792554 | -0,5081944 | 0,1770291 | 0,00409586 | 0,03767516 | TXNDC17 |
| 459,924012 | -0,508858 | 0,12090745 | 2,57E-05 | 0,00076067 | ARL13B |
| 677,322708 | -0,5090012 | 0,180189 | 0,00473074 | 0,04188833 | RNF181 |
| 5857,59369 | -0,5091618 | 0,142281 | 0,00034549 | 0,00602636 | RPL36 |
| 732,892902 | -0,5100162 | 0,11490726 | 9,06E-06 | 0,00032279 | NOP16 |
| 214,579359 | -0,5103227 | 0,19506644 | 0,0088928 | 0,06417492 | MRPL48 |
| 457,967657 | -0,5109532 | 0,13786293 | 0,00021036 | 0,00409484 | EXOSC1 |

| | | | | | |
|---|---|---|---|---|---|
| 740,125963 | -0,5109608 | 0,1266037 | 5,44E-05 | 0,00138699 | GTSF1 |
| 1900,66755 | -0,5113568 | 0,12656212 | 5,34E-05 | 0,00136643 | ZNF107 |
| 79,9412384 | -0,5124481 | 0,24143977 | 0,0337985 | 0,15563432 | RAB3A |
| 121,946646 | -0,5126679 | 0,19648142 | 0,00907426 | 0,06503562 | BLOC1S1 |
| 1893,63231 | -0,5146627 | 0,10162135 | 4,09E-07 | 2,35E-05 | R3HDM1 |
| 336,536313 | -0,5149568 | 0,12436066 | 3,46E-05 | 0,00095902 | CASC15 |
| 526,64566 | -0,5150848 | 0,13227176 | 9,85E-05 | 0,00226289 | NFE4 |
| 1141,99588 | -0,5162453 | 0,11879786 | 1,39E-05 | 0,00045999 | SNW1 |
| 101,219785 | -0,5165984 | 0,20699352 | 0,01257009 | 0,08106295 | OTX1 |
| 628,966522 | -0,5166398 | 0,14508737 | 0,0003696 | 0,00631732 | EGR1 |
| 4450,15626 | -0,5184371 | 0,11017138 | 2,53E-06 | 0,0001078 | CYTIP |
| 2149,2199 | -0,5186908 | 0,08882733 | 5,24E-09 | 5,42E-07 | ANP32B |
| 781,869995 | -0,5190263 | 0,14111981 | 0,00023515 | 0,00446419 | FKBP3 |
| 440,166233 | -0,5199739 | 0,11217566 | 3,56E-06 | 0,00014406 | ENY2 |
| 1576,00165 | -0,5213102 | 0,14247008 | 0,00025312 | 0,00472776 | CHORDC1 |
| 126,318766 | -0,5217281 | 0,24215823 | 0,03120158 | 0,14777599 | LOC642131 |
| 15963,2843 | -0,5226269 | 0,1089151 | 1,60E-06 | 7,54E-05 | PTMA |
| 235,892793 | -0,5228795 | 0,18048994 | 0,00376749 | 0,03542976 | SNX22 |
| 72,4866962 | -0,5233844 | 0,2530622 | 0,03862081 | 0,17061506 | DPH5-DT |
| 162,426154 | -0,525812 | 0,20995403 | 0,01226542 | 0,07974787 | CCDC112 |
| 6938,64144 | -0,527134 | 0,11088692 | 2,00E-06 | 9,01E-05 | UBA52 |
| 480,887148 | -0,5275461 | 0,16756448 | 0,00164213 | 0,01957834 | CCDC59 |
| 99,4537265 | -0,5278836 | 0,22388144 | 0,01838007 | 0,10437256 | AASS |
| 213,398428 | -0,5279695 | 0,16992218 | 0,00188916 | 0,0216373 | USE1 |
| 207,929067 | -0,5279876 | 0,16672329 | 0,001541 | 0,01865469 | ALKBH3 |
| 2679,16919 | -0,5281486 | 0,09141387 | 7,58E-09 | 7,47E-07 | EIF5B |
| 325,778474 | -0,5296783 | 0,1348704 | 8,59E-05 | 0,00201652 | RBMX2 |
| 78,6167981 | -0,5297713 | 0,23111478 | 0,0218916 | 0,11677785 | TBX18 |
| 1447,80334 | -0,52996 | 0,12585068 | 2,54E-05 | 0,00075511 | ARPC3 |
| 231,521912 | -0,5311566 | 0,14638736 | 0,00028515 | 0,00517285 | SCG5 |
| 2524,94949 | -0,531684 | 0,14456478 | 0,00023523 | 0,00446419 | H4C14 |
| 58,2512852 | -0,5318831 | 0,26798774 | 0,04717463 | 0,19518721 | SVBP |
| 1551,02306 | -0,5324903 | 0,1550371 | 0,00059343 | 0,00905509 | HMGB2 |
| 97,8710412 | -0,5327919 | 0,25403946 | 0,0359688 | 0,16223259 | RPL39L |
| 347,343102 | -0,5331496 | 0,14533145 | 0,00024397 | 0,00458407 | PURPL |
| 1483,77922 | -0,5331902 | 0,15298374 | 0,00049163 | 0,00791005 | IFITM1 |
| 131,072181 | -0,5332899 | 0,25666573 | 0,03773144 | 0,16805947 | ING2 |
| 781,973506 | -0,5334787 | 0,13049503 | 4,35E-05 | 0,00115247 | SNRPD1 |
| 678,14705 | -0,5340568 | 0,16352 | 0,00109077 | 0,01436219 | PPIL4 |
| 1695,25607 | -0,5342431 | 0,09962271 | 8,20E-08 | 5,81E-06 | NUDC |
| 10093,0908 | -0,5346887 | 0,10771189 | 6,90E-07 | 3,60E-05 | CD163 |
| 847,911353 | -0,5354129 | 0,11873798 | 6,51E-06 | 0,00024285 | HNRNPA1P10 |
| 450,970088 | -0,5356246 | 0,17197595 | 0,00184237 | 0,02128175 | SVIP |
| 733,807661 | -0,536025 | 0,12002569 | 7,97E-06 | 0,00028954 | C4orf3 |
| 493,885963 | -0,5373383 | 0,1149776 | 2,96E-06 | 0,00012211 | EXOSC7 |
| 75,9438911 | -0,5379116 | 0,27233014 | 0,04824332 | 0,1980069 | LOC157273 |
| 3802,59052 | -0,5380091 | 0,10942132 | 8,79E-07 | 4,45E-05 | RPS25 |
| 1054,56566 | -0,5382397 | 0,13173823 | 4,39E-05 | 0,00116139 | CDCA7L |

| | | | | | |
|---|---|---|---|---|---|
| 168,166485 | -0,5391437 | 0,1926155 | 0,00512505 | 0,04419828 | WASHC3 |
| 274,411678 | -0,5400203 | 0,1389092 | 0,00010125 | 0,0023165 | CCDC92 |
| 655,954599 | -0,5401316 | 0,1348584 | 6,20E-05 | 0,00152876 | TYW3 |
| 282,108116 | -0,54167 | 0,14685542 | 0,00022562 | 0,00433392 | LETM2 |
| 1036,58776 | -0,5429754 | 0,11364249 | 1,77E-06 | 8,09E-05 | HSD17B4 |
| 81,5930238 | -0,5430579 | 0,21859523 | 0,01298035 | 0,08255503 | ANKRD34A |
| 6352,32362 | -0,5432729 | 0,11941624 | 5,38E-06 | 0,00020399 | RPL35A |
| 187,921146 | -0,5438577 | 0,22597834 | 0,01609824 | 0,09584478 | RNASEL |
| 76,0043689 | -0,5459656 | 0,2524802 | 0,03058661 | 0,1459822 | UQCC5 |
| 138,821884 | -0,5469475 | 0,22365904 | 0,01446707 | 0,08913428 | NECAB1 |
| 3013,01315 | -0,5470446 | 0,0897578 | 1,10E-09 | 1,39E-07 | EIF4G3 |
| 171,582751 | -0,5473273 | 0,17469732 | 0,00173028 | 0,02030244 | GPKOW |
| 740,976253 | -0,5483117 | 0,13854297 | 7,57E-05 | 0,00181936 | ATP5PD |
| 1050,24728 | -0,5489844 | 0,12466115 | 1,06E-05 | 0,00036725 | EEA1 |
| 186,790093 | -0,5520106 | 0,17792708 | 0,00191923 | 0,02183632 | MED7 |
| 773,44205 | -0,554051 | 0,11742549 | 2,38E-06 | 0,00010344 | ATP5MF |
| 546,417425 | -0,5557931 | 0,13307535 | 2,96E-05 | 0,00084984 | SEC61G |
| 321,80231 | -0,5559365 | 0,17805298 | 0,00179438 | 0,02089323 | FAM241A |
| 84,6021739 | -0,556626 | 0,22865516 | 0,01491868 | 0,09119843 | SPA17 |
| 2531,63243 | -0,5567134 | 0,14185688 | 8,69E-05 | 0,00203541 | H4C15 |
| 308,977712 | -0,5568085 | 0,15435406 | 0,00030935 | 0,00550105 | IFT25 |
| 143,263503 | -0,5571595 | 0,19595296 | 0,00446444 | 0,04003396 | ELFN1-AS1 |
| 72,3935113 | -0,5585109 | 0,2502935 | 0,02565307 | 0,13003289 | DPYD-AS1 |
| 147,845132 | -0,5586289 | 0,17460584 | 0,00137728 | 0,0171229 | TCEAL8 |
| 584,593868 | -0,5594253 | 0,13506889 | 3,45E-05 | 0,00095751 | ZNF330 |
| 539,826512 | -0,5599545 | 0,12276772 | 5,09E-06 | 0,00019571 | GPATCH4 |
| 1251,0285 | -0,5612803 | 0,16191814 | 0,00052739 | 0,0083295 | GADD45A |
| 914,713379 | -0,564223 | 0,13281691 | 2,16E-05 | 0,00065892 | LSM5 |
| 2266,89949 | -0,5645436 | 0,12607639 | 7,54E-06 | 0,00027549 | COX7C |
| 155,593709 | -0,5647452 | 0,18989936 | 0,00294023 | 0,02968629 | NUDT1 |
| 267,224582 | -0,5662326 | 0,16438663 | 0,00057208 | 0,00879312 | CMSS1 |
| 417,851662 | -0,5664087 | 0,15017932 | 0,00016224 | 0,00333662 | SYF2 |
| 7174,38492 | -0,5664107 | 0,12018605 | 2,44E-06 | 0,00010508 | RPS21 |
| 2377,4068 | -0,567596 | 0,15889837 | 0,00035417 | 0,0061328 | NPIPB13 |
| 153,547134 | -0,5677873 | 0,18932405 | 0,00270846 | 0,02809586 | C7orf25 |
| 6460,5438 | -0,567823 | 0,12854647 | 1,00E-05 | 0,00034896 | BTF3 |
| 765,053077 | -0,5691534 | 0,15300977 | 0,00019944 | 0,00392684 | ELOC |
| 302,907991 | -0,5697065 | 0,14495104 | 8,48E-05 | 0,00199628 | ABRACL |
| 393,432492 | -0,5704277 | 0,1417514 | 5,72E-05 | 0,00143879 | RXYLT1 |
| 1107,12944 | -0,5709382 | 0,11301748 | 4,38E-07 | 2,47E-05 | PSMA5 |
| 1500,43611 | -0,5710629 | 0,26465427 | 0,03094578 | 0,14709951 | H2AC16 |
| 131,965531 | -0,5740765 | 0,28196967 | 0,04175525 | 0,17931255 | DPM3 |
| 1587,13329 | -0,5765991 | 0,13081472 | 1,04E-05 | 0,00036328 | UBE2V2 |
| 652,128947 | -0,5772905 | 0,13310344 | 1,44E-05 | 0,00047005 | DYNLRB1 |
| 264,697799 | -0,5775021 | 0,19660693 | 0,00331037 | 0,03225309 | LRRCC1 |
| 1087,43604 | -0,5789851 | 0,13254872 | 1,25E-05 | 0,00042567 | HNRNPA1L2 |
| 320,508422 | -0,5808247 | 0,15522844 | 0,00018275 | 0,00368635 | POLR2F |
| 5189,87928 | -0,5830051 | 0,15389552 | 0,00015167 | 0,00315399 | RPL24 |

| | | | | | |
|---|---|---|---|---|---|
| 1191,93782 | -0,5831677 | 0,10913371 | 9,11E-08 | 6,36E-06 | ZNF280C |
| 370,530523 | -0,5858494 | 0,18409655 | 0,00146113 | 0,0179364 | NENF |
| 56,2153627 | -0,5859283 | 0,2753113 | 0,03331724 | 0,15405561 | TMEM178A |
| 154,19929 | -0,5861927 | 0,18864751 | 0,00188777 | 0,0216373 | LOC101929380 |
| 1923,94264 | -0,5874792 | 0,09467826 | 5,47E-10 | 7,55E-08 | SNRPD3 |
| 13610,1712 | -0,5881258 | 0,09598129 | 8,93E-10 | 1,16E-07 | NCL |
| 2809,39965 | -0,589922 | 0,15874769 | 0,00020232 | 0,0039547 | NME1 |
| 1000,56286 | -0,593805 | 0,12456404 | 1,87E-06 | 8,48E-05 | PSMB6 |
| 875,632885 | -0,5939334 | 0,14533679 | 4,38E-05 | 0,00115839 | ATP5MG |
| 643,761986 | -0,5946452 | 0,09777867 | 1,19E-09 | 1,50E-07 | HTATSF1 |
| 464,68944 | -0,5947169 | 0,15332525 | 0,00010498 | 0,00237306 | CCL20 |
| 67,9132961 | -0,5953378 | 0,27543539 | 0,03066117 | 0,14616781 | MTHFS |
| 286,362139 | -0,5963157 | 0,15279586 | 9,51E-05 | 0,00219233 | LOC101929185 |
| 3255,94957 | -0,5970285 | 0,13268807 | 6,81E-06 | 0,00025371 | RPLP2 |
| 124,219248 | -0,5982055 | 0,22668807 | 0,00831773 | 0,06138059 | ZCWPW1 |
| 922,913183 | -0,5994035 | 0,16126326 | 0,00020166 | 0,00394995 | PPIB |
| 62,7088364 | -0,6002765 | 0,29890005 | 0,04461301 | 0,1879169 | FAM151B |
| 1862,37738 | -0,6004961 | 0,10910118 | 3,71E-08 | 2,94E-06 | SMARCE1 |
| 147,36024 | -0,6006656 | 0,17229662 | 0,00048988 | 0,00790197 | ZNF830 |
| 70,5372163 | -0,601193 | 0,26128098 | 0,02139478 | 0,11526271 | PIN4 |
| 214,730109 | -0,6029351 | 0,22355302 | 0,00699554 | 0,05474054 | RPL26L1 |
| 1320,69184 | -0,6031571 | 0,10909892 | 3,23E-08 | 2,63E-06 | SLTM |
| 2918,01354 | -0,6048949 | 0,12921067 | 2,85E-06 | 0,00011793 | G0S2 |
| 331,93601 | -0,6062757 | 0,19650519 | 0,00203353 | 0,02271336 | FLRT3 |
| 5988,10446 | -0,608443 | 0,13327992 | 4,99E-06 | 0,00019274 | HRNR |
| 931,589365 | -0,6094454 | 0,18240692 | 0,00083438 | 0,01179915 | H4C4 |
| 887,962741 | -0,6098769 | 0,12868741 | 2,15E-06 | 9,57E-05 | SRP14 |
| 1965,08549 | -0,6114133 | 0,08081925 | 3,87E-14 | 1,88E-11 | BMS1 |
| 19916,2497 | -0,6134654 | 0,12518267 | 9,56E-07 | 4,75E-05 | TIMP1 |
| 43,7660009 | -0,6135266 | 0,29235201 | 0,0358532 | 0,16205885 | EGOT |
| 3545,41672 | -0,6150321 | 0,1163542 | 1,25E-07 | 8,39E-06 | RPL22 |
| 875,317231 | -0,6154061 | 0,14938012 | 3,79E-05 | 0,00102855 | NSA2 |
| 133,409485 | -0,6166416 | 0,24101163 | 0,01051081 | 0,07156625 | LINC03033 |
| 340,041078 | -0,6168758 | 0,17860691 | 0,00055271 | 0,00864283 | DCTN6 |
| 748,175923 | -0,6169625 | 0,11168843 | 3,31E-08 | 2,68E-06 | DAD1 |
| 1086,733 | -0,6174416 | 0,10825496 | 1,17E-08 | 1,06E-06 | IWS1 |
| 8633,80897 | -0,6183354 | 0,09868633 | 3,71E-10 | 5,57E-08 | RPL23A |
| 169,926994 | -0,6186902 | 0,17229003 | 0,00032944 | 0,00580377 | LINC03006 |
| 736,281423 | -0,619426 | 0,15624081 | 7,35E-05 | 0,00177687 | MPHOSPH10 |
| 147,168281 | -0,6196023 | 0,23804012 | 0,00924302 | 0,06580486 | ZMAT1 |
| 1336,84472 | -0,6198901 | 0,13963639 | 9,02E-06 | 0,00032218 | DNTTIP2 |
| 582,947549 | -0,6198969 | 0,13174644 | 2,54E-06 | 0,00010782 | MRPL22 |
| 8437,80681 | -0,6222411 | 0,11194316 | 2,72E-08 | 2,26E-06 | RPL41 |
| 969,041132 | -0,6231496 | 0,12143904 | 2,88E-07 | 1,77E-05 | ATP5MK |
| 2977,61736 | -0,6242897 | 0,10343066 | 1,58E-09 | 1,88E-07 | FXYD5 |
| 124,743634 | -0,6251804 | 0,21195372 | 0,00318177 | 0,03133905 | RNASEH1-DT |
| 690,685251 | -0,6260847 | 0,14407646 | 1,39E-05 | 0,00045999 | LYAR |
| 12566,6605 | -0,6270045 | 0,08779792 | 9,24E-13 | 3,30E-10 | HNRNPA1 |

| | | | | | |
|---|---|---|---|---|---|
| 856,21858 | -0,6276131 | 0,11954727 | 1,52E-07 | 1,00E-05 | ATP5MJ |
| 2301,17294 | -0,6282057 | 0,10887894 | 7,94E-09 | 7,74E-07 | RAB7A |
| 48,6866318 | -0,6285498 | 0,28996347 | 0,03018258 | 0,14447805 | SNORD14B |
| 7704,49438 | -0,6286842 | 0,30849111 | 0,04155663 | 0,17878421 | RPL35 |
| 311,102103 | -0,6286983 | 0,15206565 | 3,56E-05 | 0,00097655 | LOC100506076 |
| 58,7772482 | -0,6289306 | 0,30490545 | 0,03914049 | 0,17249805 | GLRA2 |
| 218,875537 | -0,6303215 | 0,1491898 | 2,39E-05 | 0,00071773 | TIMM8B |
| 2204,99664 | -0,630689 | 0,11883747 | 1,11E-07 | 7,61E-06 | DNAJA1 |
| 954,170845 | -0,6310894 | 0,1474504 | 1,87E-05 | 0,00057926 | TIMM23 |
| 216,444427 | -0,6315768 | 0,15462295 | 4,41E-05 | 0,00116503 | F8A1 |
| 283,978059 | -0,6321198 | 0,15357826 | 3,86E-05 | 0,00104078 | NSRP1 |
| 306,61732 | -0,6334572 | 0,16009004 | 7,59E-05 | 0,0018233 | CCDST |
| 60,4766039 | -0,6339576 | 0,28267499 | 0,02491562 | 0,12731904 | SND1-IT1 |
| 639,662617 | -0,6357221 | 0,12749601 | 6,16E-07 | 3,31E-05 | LTV1 |
| 134,791479 | -0,6369136 | 0,2040409 | 0,00179933 | 0,02093791 | ATP6AP1L |
| 85,6665022 | -0,6378406 | 0,24001555 | 0,00787233 | 0,05915506 | ZNF724 |
| 86,9480956 | -0,6386573 | 0,28327334 | 0,02416084 | 0,12476728 | GOLGA2P6 |
| 1396,5524 | -0,6396867 | 0,29237295 | 0,02867592 | 0,1399918 | H2BC10 |
| 94,8621989 | -0,6408998 | 0,22052869 | 0,00365851 | 0,03468072 | FKBP2 |
| 916,353962 | -0,6414174 | 0,14186133 | 6,14E-06 | 0,00023104 | IK |
| 226,925943 | -0,643719 | 0,15462316 | 3,14E-05 | 0,00089042 | LLPH |
| 462,444825 | -0,6441871 | 0,16760489 | 0,0001213 | 0,00266252 | MRPL24 |
| 1736,9529 | -0,649488 | 0,13005697 | 5,92E-07 | 3,20E-05 | GTPBP4 |
| 1379,48308 | -0,6503072 | 0,1099318 | 3,31E-09 | 3,66E-07 | SAFB |
| 179,922994 | -0,6512224 | 0,27815799 | 0,01922207 | 0,10734972 | LINC00987 |
| 9084,69096 | -0,6523091 | 0,10225656 | 1,78E-10 | 3,12E-08 | MKI67 |
| 718,384936 | -0,6529697 | 0,14606572 | 7,81E-06 | 0,00028465 | CWC27 |
| 535,057471 | -0,6530505 | 0,16320859 | 6,30E-05 | 0,00155184 | DNAJB11 |
| 1444,58051 | -0,6545842 | 0,13590944 | 1,46E-06 | 6,97E-05 | TAF15 |
| 2728,77307 | -0,6547138 | 0,23868522 | 0,00608805 | 0,04993149 | H3C3 |
| 185,227596 | -0,6561562 | 0,17880948 | 0,00024295 | 0,00457392 | IFT74 |
| 68,5432141 | -0,656774 | 0,25237535 | 0,00925819 | 0,06585498 | COPG2IT1 |
| 1136,92954 | -0,658129 | 0,15541953 | 2,29E-05 | 0,00069265 | COX5A |
| 578,518522 | -0,6593079 | 0,15757472 | 2,86E-05 | 0,00082831 | NDUFA5 |
| 48,3420756 | -0,6596491 | 0,31372821 | 0,03549959 | 0,16085444 | SNHG9 |
| 765,962119 | -0,6618856 | 0,15813613 | 2,84E-05 | 0,00082557 | SSBP1 |
| 1100,78095 | -0,6622197 | 0,15399596 | 1,71E-05 | 0,00054164 | CLEC4D |
| 68,6508294 | -0,6624313 | 0,25836663 | 0,01034975 | 0,07092602 | ANKRD20A1 |
| 4305,79312 | -0,6630284 | 0,15272545 | 1,42E-05 | 0,00046445 | RPL27A |
| 310,480397 | -0,6635359 | 0,1339056 | 7,22E-07 | 3,75E-05 | LINC01347 |
| 785,916496 | -0,6637839 | 0,14377291 | 3,90E-06 | 0,00015582 | MRPL51 |
| 448,906182 | -0,6649744 | 0,13891325 | 1,69E-06 | 7,87E-05 | CKS2 |
| 531,70041 | -0,665744 | 0,15643055 | 2,08E-05 | 0,00063866 | IFT57 |
| 142,268392 | -0,6665197 | 0,2045737 | 0,00112164 | 0,0146352 | CNTLN |
| 690,256596 | -0,6679703 | 0,19468685 | 0,00060136 | 0,00911731 | LOC102724594 |
| 385,486939 | -0,6706566 | 0,23236985 | 0,00389974 | 0,03626072 | HPF1 |
| 369,171913 | -0,6716468 | 0,14584414 | 4,12E-06 | 0,00016308 | CD163L1 |
| 43,6080543 | -0,6722389 | 0,34269803 | 0,04980823 | 0,20196902 | MIR1206 |

| | | | | | |
|---|---|---|---|---|---|
| 179,344635 | -0,6724517 | 0,23843289 | 0,00479791 | 0,04232229 | H2BC3 |
| 58,8835661 | -0,6727869 | 0,27208825 | 0,01341046 | 0,08457975 | SYCP3 |
| 560,993737 | -0,6729201 | 0,18586761 | 0,00029412 | 0,00530404 | LSM3 |
| 852,671791 | -0,6729941 | 0,12499158 | 7,27E-08 | 5,18E-06 | U2AF1 |
| 402,72437 | -0,673242 | 0,17496644 | 0,00011917 | 0,00263097 | GAR1 |
| 82,3499568 | -0,6735161 | 0,26274726 | 0,01036642 | 0,07096347 | LINC02325 |
| 1981,70956 | -0,6739084 | 0,10764354 | 3,84E-10 | 5,67E-08 | SMC3 |
| 4037,80183 | -0,6761887 | 0,10893707 | 5,40E-10 | 7,51E-08 | RPS12 |
| 277,400363 | -0,6788907 | 0,1883617 | 0,00031313 | 0,00554411 | MRPL33 |
| 666,103117 | -0,6794058 | 0,11240385 | 1,50E-09 | 1,82E-07 | UBLCP1 |
| 186,740077 | -0,6794427 | 0,19836771 | 0,00061442 | 0,00925593 | SNHG6 |
| 3172,16907 | -0,6806366 | 0,13462388 | 4,29E-07 | 2,43E-05 | HNRNPD |
| 7603,5671 | -0,6813124 | 0,2476709 | 0,00594358 | 0,04908709 | YBX1 |
| 142,478569 | -0,68263 | 0,20878913 | 0,00107749 | 0,01425676 | FAIM |
| 490,859956 | -0,6842776 | 0,17395625 | 8,37E-05 | 0,00197658 | UBE2T |
| 402,316913 | -0,6848921 | 0,15062002 | 5,44E-06 | 0,00020577 | ESF1 |
| 786,157858 | -0,6881892 | 0,29062962 | 0,01788815 | 0,10237798 | SLIRP |
| 553,85042 | -0,6890354 | 0,11021671 | 4,06E-10 | 5,96E-08 | GGCT |
| 554,200323 | -0,6906127 | 0,13901318 | 6,77E-07 | 3,57E-05 | MEA1 |
| 185,180024 | -0,691399 | 0,16135476 | 1,83E-05 | 0,00057066 | CFAP68 |
| 146,394412 | -0,6914251 | 0,18770938 | 0,00023006 | 0,00439259 | WBP4 |
| 238,299666 | -0,6916156 | 0,18538968 | 0,00019102 | 0,00380453 | BRME1 |
| 89,2553566 | -0,6929787 | 0,26440143 | 0,00876892 | 0,06365452 | LINC01094 |
| 298,149541 | -0,6945025 | 0,14456306 | 1,55E-06 | 7,37E-05 | KRT10 |
| 728,368921 | -0,6948089 | 0,12904694 | 7,28E-08 | 5,18E-06 | H2BC15 |
| 585,701011 | -0,6962568 | 0,13515364 | 2,58E-07 | 1,62E-05 | NEDD8 |
| 228,441955 | -0,696557 | 0,18509856 | 0,00016777 | 0,00343928 | SNRPF |
| 1507,85968 | -0,6974923 | 0,10180747 | 7,33E-12 | 1,90E-09 | SYN1 |
| 646,561593 | -0,6985123 | 0,12756405 | 4,36E-08 | 3,39E-06 | UFC1 |
| 1097,26048 | -0,7006309 | 0,1170642 | 2,16E-09 | 2,54E-07 | H2BC13 |
| 978,33893 | -0,7025794 | 0,30709297 | 0,02214687 | 0,11747716 | SPP1 |
| 1170,38324 | -0,7028918 | 0,13802293 | 3,53E-07 | 2,11E-05 | PSMB1 |
| 600,547114 | -0,7052069 | 0,14077943 | 5,46E-07 | 2,98E-05 | NDUFB10 |
| 170,631579 | -0,7059074 | 0,17549748 | 5,76E-05 | 0,00144611 | SERPINA9 |
| 401,554816 | -0,7060601 | 0,15519789 | 5,38E-06 | 0,00020399 | PPID |
| 1013,23878 | -0,7067459 | 0,11286896 | 3,81E-10 | 5,67E-08 | MZT2B |
| 442,062947 | -0,7083862 | 0,12940785 | 4,40E-08 | 3,41E-06 | CWC15 |
| 845,301554 | -0,7099529 | 0,16039816 | 9,59E-06 | 0,00033857 | H2BC26 |
| 172,5499 | -0,7139291 | 0,2114001 | 0,00073242 | 0,01062739 | MND1 |
| 253,414143 | -0,7148963 | 0,19329172 | 0,00021685 | 0,0041867 | MIX23 |
| 226,395748 | -0,7157474 | 0,19109468 | 0,00018003 | 0,00363919 | CETN3 |
| 43,8368422 | -0,7157846 | 0,36403424 | 0,04926902 | 0,20064983 | SLAMF8 |
| 205,417072 | -0,7174337 | 0,21676278 | 0,00093375 | 0,01283968 | RIDA |
| 3309,72807 | -0,719251 | 0,1564906 | 4,30E-06 | 0,00016966 | S100A11 |
| 349,843561 | -0,722812 | 0,15089346 | 1,67E-06 | 7,78E-05 | NDUFA12 |
| 1443,97728 | -0,7245062 | 0,10608254 | 8,51E-12 | 2,09E-09 | PNN |
| 419,723581 | -0,7289836 | 0,12443291 | 4,67E-09 | 4,91E-07 | CCDC34 |
| 1578,83603 | -0,7321637 | 0,12093209 | 1,41E-09 | 1,74E-07 | NASP |

| | | | | | |
|---|---|---|---|---|---|
| 558,088673 | -0,7341672 | 0,13275733 | 3,20E-08 | 2,62E-06 | POLR2I |
| 479,783883 | -0,7350538 | 0,13530513 | 5,55E-08 | 4,14E-06 | EXOSC3 |
| 38,0086905 | -0,7379439 | 0,34110779 | 0,03051238 | 0,14578908 | AK7 |
| 350,46553 | -0,739077 | 0,21001146 | 0,00043281 | 0,00717099 | NDUFB1 |
| 1145,04724 | -0,7391563 | 0,15661519 | 2,36E-06 | 0,00010304 | H2BC8 |
| 48,6139309 | -0,7415979 | 0,30448184 | 0,01486685 | 0,09103422 | RNU1-3 |
| 67,1545593 | -0,7417846 | 0,25213 | 0,00326024 | 0,03186314 | RNU1-1 |
| 9225,07395 | -0,742803 | 0,12282494 | 1,47E-09 | 1,79E-07 | RPL19 |
| 217,971488 | -0,744749 | 0,25616481 | 0,00364558 | 0,03462756 | MAS1L |
| 186,519042 | -0,745322 | 0,1819272 | 4,19E-05 | 0,00111785 | LSM6 |
| 11288,5375 | -0,7476004 | 0,13248146 | 1,67E-08 | 1,46E-06 | RPL6 |
| 47,5899929 | -0,7483047 | 0,34607725 | 0,03059909 | 0,1459822 | C5 |
| 70,2210744 | -0,7509353 | 0,31498533 | 0,01712408 | 0,09945292 | TFAP2A |
| 158,717425 | -0,7518936 | 0,17422991 | 1,59E-05 | 0,00051323 | RPL17 |
| 1475,13132 | -0,753975 | 0,15524558 | 1,19E-06 | 5,81E-05 | TBCA |
| 3057,01925 | -0,7583287 | 0,11088999 | 8,00E-12 | 2,05E-09 | FABP5 |
| 1556,79966 | -0,7607257 | 0,10483277 | 3,97E-13 | 1,57E-10 | H2BC5 |
| 64,9804072 | -0,7649042 | 0,25538753 | 0,0027438 | 0,02838457 | GIHCG |
| 326,435244 | -0,7669765 | 0,18568839 | 3,62E-05 | 0,00098852 | RWDD1 |
| 131,397878 | -0,7672027 | 0,20597152 | 0,00019547 | 0,00387277 | SPMIP4 |
| 1455,0516 | -0,7686347 | 0,09851444 | 6,08E-15 | 3,97E-12 | HNRNPH3 |
| 68,8165354 | -0,7697364 | 0,34623757 | 0,02620601 | 0,13173323 | LY96 |
| 2771,28441 | -0,7708073 | 0,27347685 | 0,00482416 | 0,0424154 | H2AC13 |
| 365,422282 | -0,774153 | 0,16064719 | 1,44E-06 | 6,89E-05 | MAGOH |
| 785,335729 | -0,7752277 | 0,14093608 | 3,79E-08 | 2,98E-06 | GNG5 |
| 144,917903 | -0,7805043 | 0,19068017 | 4,25E-05 | 0,0011332 | ABCA17P |
| 2551,72735 | -0,7815155 | 0,1357568 | 8,58E-09 | 8,32E-07 | H2BC4 |
| 325,766447 | -0,792428 | 0,17843683 | 8,96E-06 | 0,00032157 | LINC02067 |
| 45,1264928 | -0,7968305 | 0,29668917 | 0,0072369 | 0,05604966 | TMEM14A |
| 184,665817 | -0,8012666 | 0,18810732 | 2,05E-05 | 0,00062897 | SEC11C |
| 80,1524542 | -0,8074048 | 0,292535 | 0,00577964 | 0,04811112 | VCF2 |
| 73,5442826 | -0,8116507 | 0,23955603 | 0,00070367 | 0,01031303 | ADAM20P1 |
| 83,873928 | -0,813517 | 0,31587447 | 0,01001115 | 0,06946129 | RNU11 |
| 336,963545 | -0,8186798 | 0,16719919 | 9,76E-07 | 4,82E-05 | LEO1 |
| 321,335959 | -0,8207757 | 0,16757234 | 9,68E-07 | 4,79E-05 | NDUFB9 |
| 141,165562 | -0,8211876 | 0,21757096 | 0,00016043 | 0,00330869 | GLRX2 |
| 1251,08051 | -0,8219816 | 0,35020281 | 0,01891719 | 0,10621133 | H3C12 |
| 693,206322 | -0,8228955 | 0,1164723 | 1,60E-12 | 5,33E-10 | UQCRH |
| 136,118918 | -0,824489 | 0,21566828 | 0,00013186 | 0,00281932 | LINC02604 |
| 5261,96776 | -0,8308489 | 0,1646785 | 4,53E-07 | 2,54E-05 | RPL31 |
| 4089,53118 | -0,8314069 | 0,10870823 | 2,04E-14 | 1,14E-11 | NACA |
| 7349,40555 | -0,8321979 | 0,11812499 | 1,85E-12 | 5,94E-10 | RPL14 |
| 9534,58168 | -0,8322171 | 0,21769027 | 0,00013187 | 0,00281932 | RPS17 |
| 47,6446296 | -0,8369819 | 0,33357574 | 0,0121032 | 0,07915819 | KANK1 |
| 7939,47608 | -0,8383729 | 0,2426574 | 0,00055036 | 0,00862038 | RPS24 |
| 110,025944 | -0,8434134 | 0,28466261 | 0,00304802 | 0,0303695 | STAG3 |
| 392,944834 | -0,8443748 | 0,17996297 | 2,71E-06 | 0,00011354 | ATP5IF1 |
| 225,562387 | -0,8447654 | 0,19450541 | 1,40E-05 | 0,00046157 | NDUFAF2 |

| | | | | | |
|---|---|---|---|---|---|
| 72,9943477 | -0,8454778 | 0,27479899 | 0,00209296 | 0,02314491 | SCARNA17 |
| 7824,09382 | -0,8455705 | 0,14759066 | 1,01E-08 | 9,36E-07 | RPL12 |
| 327,112761 | -0,8491319 | 0,15981032 | 1,08E-07 | 7,43E-06 | RPS27L |
| 41,6231097 | -0,8536439 | 0,31649443 | 0,00699285 | 0,05474054 | C2orf74 |
| 34,5996766 | -0,8571045 | 0,39634003 | 0,03057592 | 0,1459822 | POPDC1 |
| 20,4637981 | -0,8601441 | 0,43303377 | 0,04699769 | 0,19462538 | ST7-OT3 |
| 56,4304106 | -0,8615137 | 0,3186823 | 0,00686419 | 0,0540519 | DDX11L16 |
| 915,979679 | -0,8619806 | 0,10983769 | 4,24E-15 | 2,97E-12 | PDCD5 |
| 139,568727 | -0,8629867 | 0,20086569 | 1,74E-05 | 0,00054877 | CHCHD1 |
| 1219,53336 | -0,8645513 | 0,16143988 | 8,54E-08 | 6,03E-06 | H2BC7 |
| 135,721266 | -0,8681493 | 0,33934185 | 0,01051755 | 0,07156625 | COX17 |
| 22,0748425 | -0,8724722 | 0,43791223 | 0,04633323 | 0,19301434 | LOC101927919 |
| 242,572028 | -0,8727127 | 0,18182197 | 1,59E-06 | 7,51E-05 | COA6 |
| 1173,14776 | -0,8729255 | 0,16441262 | 1,10E-07 | 7,55E-06 | SSB |
| 328,986786 | -0,873638 | 0,18525949 | 2,41E-06 | 0,00010409 | DBI |
| 7145,30928 | -0,8774217 | 0,22365821 | 8,74E-05 | 0,00204497 | RPS27A |
| 204,711087 | -0,8791178 | 0,18232791 | 1,42E-06 | 6,82E-05 | SCARNA9L |
| 12312,4618 | -0,8812135 | 0,11075503 | 1,77E-15 | 1,68E-12 | RPS18 |
| 210,613424 | -0,8813121 | 0,18559728 | 2,05E-06 | 9,21E-05 | GEMIN6 |
| 85,5560892 | -0,8855689 | 0,26820236 | 0,00096043 | 0,01306421 | RAB5IF |
| 109,580869 | -0,8910678 | 0,19434749 | 4,54E-06 | 0,00017718 | TCEAL4 |
| 105,790749 | -0,8943942 | 0,25291112 | 0,00040562 | 0,00684029 | ZNF311 |
| 1451,6401 | -0,904846 | 0,11951248 | 3,70E-14 | 1,84E-11 | RPS29 |
| 11154,0543 | -0,9066354 | 0,15791164 | 9,39E-09 | 8,79E-07 | RPS11 |
| 646,141825 | -0,9075734 | 0,18989307 | 1,76E-06 | 8,06E-05 | NDUFA6 |
| 34,963307 | -0,9085748 | 0,40658713 | 0,02544116 | 0,12933162 | SAMD15 |
| 256,568411 | -0,9154807 | 0,18895442 | 1,27E-06 | 6,11E-05 | CSTB |
| 175,247511 | -0,9175814 | 0,19419399 | 2,30E-06 | 0,00010098 | ATP5ME |
| 13874,7501 | -0,9202198 | 0,14851988 | 5,79E-10 | 7,94E-08 | RPS23 |
| 1793,62529 | -0,9217641 | 0,24999416 | 0,00022679 | 0,00435195 | RPS26 |
| 122,229151 | -0,9278952 | 0,2243311 | 3,53E-05 | 0,0009736 | MRPS18C |
| 493,233697 | -0,9391968 | 0,20891653 | 6,94E-06 | 0,00025692 | NDUFB3 |
| 19,4473197 | -0,9458147 | 0,44116598 | 0,03204109 | 0,15043931 | CTNND1 |
| 97,9731587 | -0,9496879 | 0,28432671 | 0,0008374 | 0,01182211 | MORN2 |
| 162,972768 | -0,9595825 | 0,26455613 | 0,00028658 | 0,00518726 | SCARNA21 |
| 612,528728 | -0,9634743 | 0,14303582 | 1,63E-11 | 3,89E-09 | TOMM5 |
| 377,574257 | -0,9645994 | 0,15960463 | 1,51E-09 | 1,82E-07 | MRFAP1L1 |
| 495,335194 | -0,9687635 | 0,14822575 | 6,33E-11 | 1,29E-08 | UBL5 |
| 200,148466 | -0,9687851 | 0,17446848 | 2,81E-08 | 2,32E-06 | PET100 |
| 400,587852 | -0,9695124 | 0,22475535 | 1,61E-05 | 0,00051558 | H4C13 |
| 24,1384751 | -0,9763697 | 0,44838787 | 0,02944243 | 0,14236654 | SILC1 |
| 969,784185 | -0,976817 | 0,27469201 | 0,00037648 | 0,00641746 | UQCRB |
| 244,96953 | -0,9910365 | 0,17756675 | 2,39E-08 | 2,01E-06 | NDUFA1 |
| 45,3334233 | -0,9956934 | 0,33549752 | 0,00299928 | 0,03001026 | RNVU1-19 |
| 40,0024319 | -0,9997254 | 0,38711432 | 0,00980854 | 0,06859635 | RANBP3L |
| 756,992807 | -1,0009599 | 0,17117819 | 4,99E-09 | 5,22E-07 | H2BC9 |
| 15,5410272 | -1,0024897 | 0,48642645 | 0,03930995 | 0,17281218 | PPIC |
| 210,659488 | -1,015612 | 0,18683198 | 5,45E-08 | 4,09E-06 | KGD4 |

| | | | | | |
|---|---|---|---|---|---|
| 266,63507 | -1,0161365 | 0,16350739 | 5,14E-10 | 7,26E-08 | PFDN2 |
| 24,2905743 | -1,0480591 | 0,42020341 | 0,01262518 | 0,08128719 | NR2F1-AS1 |
| 17,0883238 | -1,0498998 | 0,50913073 | 0,03919422 | 0,1725841 | TMEM30A-DT |
| 6276,38313 | -1,0529451 | 0,13494465 | 6,06E-15 | 3,97E-12 | RPL37A |
| 458,686272 | -1,0645614 | 0,19395494 | 4,05E-08 | 3,17E-06 | NDUFS5 |
| 19,8052805 | -1,0743956 | 0,48938234 | 0,0281341 | 0,13819454 | MGC2889 |
| 438,472746 | -1,0872333 | 0,14255726 | 2,41E-14 | 1,30E-11 | TMA7 |
| 874,125587 | -1,0873922 | 0,13756343 | 2,69E-15 | 2,20E-12 | SH3BGRL3 |
| 1352,32181 | -1,0921738 | 0,47686418 | 0,02200251 | 0,11720423 | SNORD3A |
| 491,753773 | -1,0953024 | 0,14116115 | 8,54E-15 | 5,39E-12 | RPL36AL |
| 530,636911 | -1,1087405 | 0,17854196 | 5,30E-10 | 7,43E-08 | SNRPE |
| 24,0395722 | -1,1169551 | 0,44227285 | 0,01155375 | 0,07660507 | TRK-CTT2-2 |
| 383,685979 | -1,120049 | 0,15602925 | 7,05E-13 | 2,62E-10 | NDUFA3 |
| 13,3688111 | -1,1279183 | 0,5750462 | 0,04982778 | 0,20196902 | TRT-AGT3-1 |
| 27,5224635 | -1,1379585 | 0,5387834 | 0,03467881 | 0,15822468 | DCDC1 |
| 36,2944486 | -1,1403387 | 0,38375806 | 0,00296345 | 0,02974609 | MIR3609 |
| 18,7716679 | -1,1557108 | 0,47181779 | 0,01430605 | 0,08841893 | MIR3134 |
| 6235,03348 | -1,1626912 | 0,31148361 | 0,0001894 | 0,00378417 | RPL37 |
| 14,0442834 | -1,1627224 | 0,59321802 | 0,04999281 | 0,20227277 | LINC01053 |
| 15,2109764 | -1,1673339 | 0,59454994 | 0,04960077 | 0,20152378 | OR7E2P |
| 38,5389085 | -1,1783369 | 0,49005844 | 0,01619538 | 0,09612071 | SNORD62B |
| 886,963609 | -1,1792907 | 0,16471471 | 8,09E-13 | 2,94E-10 | SCARNA2 |
| 16,3440571 | -1,1914904 | 0,58404055 | 0,04134219 | 0,17830764 | RNU5A-1 |
| 16,5330457 | -1,1964669 | 0,50880626 | 0,018697 | 0,10553877 | CCDC81 |
| 15,9136938 | -1,197386 | 0,58893044 | 0,04203702 | 0,18015457 | SLC26A7 |
| 173,299294 | -1,2038826 | 0,2092603 | 8,77E-09 | 8,46E-07 | LINC01061 |
| 14,4158599 | -1,2062797 | 0,54037549 | 0,0255952 | 0,12990526 | CAVIN4 |
| 171,494958 | -1,2117953 | 0,2003179 | 1,45E-09 | 1,79E-07 | CDC26 |
| 24,4306468 | -1,2227848 | 0,40094082 | 0,00229002 | 0,02473148 | SNORD30 |
| 4538,32684 | -1,2323105 | 0,11517504 | 1,02E-26 | 4,84E-23 | FCER1G |
| 2488,39214 | -1,2345954 | 0,35787097 | 0,00056093 | 0,00871378 | RPL27 |
| 336,098568 | -1,2369514 | 0,19366761 | 1,69E-10 | 2,99E-08 | H2BC14 |
| 43,1152036 | -1,2445615 | 0,36686838 | 0,00069285 | 0,01019396 | CADPS2 |
| 29,3964791 | -1,2468569 | 0,40286534 | 0,00196832 | 0,02223432 | LOC101928896 |
| 9,89138836 | -1,2479809 | 0,63141919 | 0,04810161 | 0,19755383 | EXTL3-AS1 |
| 62,0246565 | -1,2586192 | 0,36836528 | 0,00063369 | 0,0095159 | MT2A |
| 423,987763 | -1,2648457 | 0,18241063 | 4,09E-12 | 1,14E-09 | TOMM7 |
| 30,8311819 | -1,2740673 | 0,37842676 | 0,0007606 | 0,01096903 | VTRNA1-3 |
| 10,3200361 | -1,3347877 | 0,60621169 | 0,02767585 | 0,13655694 | TRU-TCA1-1 |
| 54,083436 | -1,3444461 | 0,38622424 | 0,00049955 | 0,00799102 | SCARNA9 |
| 34,2093581 | -1,3522838 | 0,37879957 | 0,00035709 | 0,00616668 | RNVU1-14 |
| 34,2762373 | -1,3528864 | 0,41299698 | 0,00105371 | 0,01407009 | SCARNA7 |
| 751,372313 | -1,357941 | 0,17140063 | 2,33E-15 | 2,10E-12 | HSPE1 |
| 12,6489279 | -1,3777109 | 0,62251477 | 0,02688815 | 0,1341996 | MIR1299 |
| 14,8753773 | -1,3827504 | 0,6045098 | 0,022173 | 0,11758275 | PRRT1 |
| 244,712002 | -1,3864504 | 0,16858125 | 1,96E-16 | 2,32E-13 | NDUFS6 |
| 4495,76517 | -1,4012241 | 0,13122835 | 1,29E-26 | 4,90E-23 | RPL11 |
| 13,174607 | -1,4090799 | 0,67863844 | 0,03786309 | 0,16840798 | MYO7A |

| | | | | | |
|---|---|---|---|---|---|
| 403,987744 | -1,4091533 | 0,19788532 | 1,07E-12 | 3,75E-10 | C19orf53 |
| 8,60808032 | -1,4447179 | 0,68226543 | 0,03421484 | 0,15690234 | CDIPTOSP |
| 2210,89273 | -1,4582749 | 0,13366742 | 1,04E-27 | 6,53E-24 | RPS27 |
| 14,5286926 | -1,495181 | 0,62877625 | 0,0174105 | 0,10040356 | SNORD54 |
| 17,6978612 | -1,5004325 | 0,47658023 | 0,00164205 | 0,01957834 | RNVU1-15 |
| 53,7402221 | -1,5557499 | 0,31316223 | 6,77E-07 | 3,57E-05 | SEM1 |
| 13,3079292 | -1,5942778 | 0,64350489 | 0,01323096 | 0,08369878 | LOC101929420 |
| 7,49380435 | -1,7181507 | 0,78471629 | 0,02855918 | 0,13973838 | CLCA2 |
| 14,3988344 | -1,7491736 | 0,59181548 | 0,00312055 | 0,03086766 | BCYRN1 |
| 15,0528694 | -1,7948558 | 0,72379621 | 0,01314632 | 0,08333052 | RNU5D-1 |
| 780,171974 | -1,8288806 | 0,16688166 | 6,01E-28 | 6,53E-24 | H4C5 |
| 14,3034657 | -1,9348514 | 0,59585005 | 0,00116542 | 0,01507234 | HOXA7 |
| 366,199532 | -2,0402373 | 0,68237226 | 0,00279052 | 0,02867977 | SNORD3C |
| 16,7294778 | -2,131475 | 0,57822928 | 0,00022762 | 0,00435907 | FAM111B |
| 3,82894945 | -2,2324476 | 1,07307737 | 0,03748737 | 0,16724793 | PRELP |
| 4,997742 | -2,3393851 | 1,10197021 | 0,03376128 | 0,15550078 | RNVU1-2 |
| 6,45142435 | -2,3437563 | 0,97926387 | 0,01669368 | 0,09805887 | SLC22A18AS |
| 7,31428907 | -2,3580794 | 0,9705972 | 0,01511909 | 0,09204255 | BHMT |
| 6,06573487 | -2,4884101 | 0,92195786 | 0,00695378 | 0,05458337 | FALEC |
| 8,52739371 | -2,8141616 | 0,80740463 | 0,00049133 | 0,00791005 | MIR378E |
| 3,46022194 | -2,8842159 | 1,27089058 | 0,0232413 | 0,12182235 | LINC02608 |
| 3,73473784 | -2,9786022 | 1,29214906 | 0,02115797 | 0,11451083 | SPEM1 |
| 5,26698418 | -3,0014347 | 1,37224913 | 0,0287252 | 0,14017744 | HTRA3 |
| 2,22918991 | -3,1889875 | 1,56276728 | 0,04129031 | 0,17819492 | PIK3C2G |
| 3,25045131 | -3,3585864 | 1,41194862 | 0,01737434 | 0,10033566 | MIR3143 |
| 2,51549641 | -3,4850712 | 1,52016532 | 0,02187331 | 0,11674609 | COMETT |
| 2,84500301 | -3,5406109 | 1,48098395 | 0,01681559 | 0,09850395 | MIR5690 |
| 1,58262582 | -3,5732375 | 1,80904228 | 0,04824438 | NA | ADH1A |
| 1,43749018 | -3,6291293 | 1,78414499 | 0,04194152 | NA | LURAP1L |
| 1,69567259 | -3,7571229 | 1,91424568 | 0,04967905 | 0,20166858 | CAP2 |
| 1,28661617 | -3,7705958 | 1,88996493 | 0,0460362 | NA | KLHL33 |
| 15,265598 | -3,994822 | 0,63705069 | 3,59E-10 | 5,44E-08 | NBPF18P |
| 1,43904634 | -4,0278759 | 1,97283105 | 0,04118398 | NA | LINC01247 |
| 1,52700601 | -4,0705852 | 1,7798592 | 0,02219427 | NA | NUP133-DT |
| 1,24653457 | -4,3606348 | 2,05746515 | 0,0340549 | NA | MIR6876 |
| 2,52415532 | -4,4101787 | 1,58009978 | 0,00525331 | 0,04496241 | TRS-AGA2-2 |
| 1,94840678 | -4,4688739 | 1,64888532 | 0,00672347 | 0,05318341 | RNF148 |
| 2,18742604 | -4,6518641 | 1,57738239 | 0,00318697 | 0,03137392 | HOXB5 |
| 1,80216826 | -4,6915241 | 2,21270301 | 0,03398342 | 0,15625767 | TRA-AGC3-1 |
| 1,8082121 | -4,9337858 | 1,79475247 | 0,00597762 | 0,04928219 | ZPBP |
| 2,85213614 | -5,6235311 | 1,60107025 | 0,00044415 | 0,00730762 | HPSE2 |
| 4,02122458 | -6,163344 | 2,88553857 | 0,03268411 | 0,1521774 | TRDN |

| A23N2_vs_CTL | | | | | |
|---|---|---|---|---|---|
| baseMean | log2FoldChange | lfcSE | pvalue | padj | gene_name |
| 30,0983051 | 5,69153398 | 1,32851733 | 8,56E-07 | 0,00029494 | CXCR2P1 |
| 22,7936235 | 5,325089799 | 1,35994307 | 4,57E-06 | 0,0010304 | DEFA1B |
| 23,4057792 | 5,288432335 | 1,37046352 | 5,76E-06 | 0,00123516 | DEFA1 |
| 10,947005 | 4,853555116 | 1,37086632 | 2,21E-05 | 0,00360271 | ARRDC4 |
| 45,6605087 | 4,521906344 | 0,77504045 | 1,10E-09 | 1,61E-06 | TYROBP |
| 10,9044664 | 3,805531571 | 1,31608904 | 0,00036012 | 0,02452699 | IL24 |
| 61,5673077 | 3,509286599 | 0,69063721 | 2,71E-08 | 1,84E-05 | PLXND1 |
| 60,1545178 | 3,496098523 | 0,57051291 | 8,88E-11 | 1,95E-07 | NRIP1 |
| 30,0876616 | 3,337632039 | 0,75417692 | 7,65E-07 | 0,0002744 | IGHA1 |
| 25,0509663 | 3,302431694 | 0,88783833 | 1,58E-05 | 0,00286188 | SIGLEC14 |
| 10,6307109 | 3,21284486 | 0,99048877 | 8,82E-05 | 0,00956838 | ADM |
| 279,112569 | 3,200012889 | 0,63787825 | 2,09E-08 | 1,83E-05 | XIST |
| 127,839999 | 3,199858994 | 0,69960285 | 2,02E-07 | 9,84E-05 | CYBB |
| 37,1761933 | 3,198619547 | 0,75653008 | 1,35E-06 | 0,00042206 | P2RX1 |
| 65,3099555 | 3,097193278 | 0,54962898 | 1,33E-09 | 1,79E-06 | CLEC7A |
| 49,6929676 | 3,092919596 | 0,72241632 | 1,01E-06 | 0,00034079 | CD69 |
| 30,157293 | 3,085725519 | 0,83985407 | 1,23E-05 | 0,00237409 | EPB41L3 |
| 54,5422274 | 3,039606878 | 0,65742187 | 2,10E-07 | 9,99E-05 | IRS2 |
| 12,4580708 | 3,019084421 | 0,9547223 | 0,00012907 | 0,01187438 | KIAA0040 |
| 69,5909647 | 2,929476179 | 0,48915588 | 1,46E-10 | 2,57E-07 | LRRK2 |
| 30,5255803 | 2,909432024 | 0,82757375 | 2,26E-05 | 0,00361794 | TES |
| 193,161192 | 2,850859747 | 0,8548842 | 2,66E-05 | 0,00401894 | TNFAIP2 |
| 185,243184 | 2,818941869 | 0,70421126 | 2,29E-06 | 0,00062834 | COTL1 |
| 50,6366336 | 2,765144685 | 0,62999014 | 6,06E-07 | 0,00023106 | TC2N |
| 90,6657446 | 2,757141638 | 0,49238496 | 1,09E-09 | 1,61E-06 | RIPOR2 |
| 167,137796 | 2,657590486 | 0,78497438 | 2,32E-05 | 0,00366346 | F13A1 |
| 10,6123084 | 2,65708488 | 0,98062099 | 0,00044548 | 0,02816972 | FCRL1 |
| 62,0192045 | 2,634462517 | 0,54292101 | 6,63E-08 | 3,64E-05 | LFNG |
| 139,829307 | 2,594226854 | 0,41751431 | 2,60E-11 | 6,54E-08 | ANKRD44 |
| 43,1216451 | 2,588688421 | 0,58781097 | 6,18E-07 | 0,00023106 | CYRIA |
| 80,6933948 | 2,564358343 | 0,52832566 | 5,67E-08 | 3,32E-05 | DMTN |
| 25,1287419 | 2,556158281 | 0,82109613 | 8,21E-05 | 0,00896019 | LCK |
| 67,5363835 | 2,529823699 | 0,91656827 | 0,00017111 | 0,01481843 | ELF4 |
| 75,7226034 | 2,49745646 | 0,55068079 | 2,69E-07 | 0,00012423 | PLEK |
| 37,6501806 | 2,482756727 | 0,71509852 | 2,34E-05 | 0,00366346 | ZNF331 |
| 79,5391881 | 2,468989205 | 0,49243564 | 2,50E-08 | 1,84E-05 | SESN3 |
| 367,577413 | 2,454744973 | 0,74966049 | 3,35E-05 | 0,00470331 | LYZ |
| 63,8898494 | 2,443085068 | 0,85816345 | 0,00014016 | 0,0126952 | GYPC |
| 42,475832 | 2,423439787 | 0,7296466 | 3,74E-05 | 0,00517536 | RCSD1 |
| 211,652708 | 2,396765847 | 0,78551029 | 6,91E-05 | 0,00814354 | VSIR |
| 50,9764519 | 2,364999693 | 0,70273806 | 3,03E-05 | 0,00438614 | IGHM |
| 63,5426922 | 2,364318945 | 0,58938547 | 2,65E-06 | 0,00067385 | IRAK3 |
| 118,130137 | 2,359752819 | 0,67215079 | 1,58E-05 | 0,00286188 | CAVIN2 |
| 117,615809 | 2,35928666 | 0,7973217 | 9,39E-05 | 0,00982436 | SLC4A1 |
| 60,8740772 | 2,331105814 | 0,60612068 | 5,35E-06 | 0,00115976 | ABCG1 |
| 198,037879 | 2,326129039 | 0,91196294 | 0,00027681 | 0,02061053 | PPBP |

| | | | | | |
|---|---|---|---|---|---|
| 168,743955 | 2,293112685 | 0,72341673 | 4,92E-05 | 0,00631375 | DOCK8 |
| 209,343295 | 2,286122281 | 0,64245878 | 1,28E-05 | 0,00245096 | CXCR4 |
| 145,540802 | 2,249583927 | 0,67317643 | 2,78E-05 | 0,00410223 | INPP5D |
| 111,458816 | 2,236945735 | 0,83755005 | 0,00021107 | 0,01693571 | PIK3R5 |
| 56,6690527 | 2,232376233 | 0,60283216 | 8,39E-06 | 0,00169367 | HCK |
| 136,202007 | 2,22244553 | 0,73259548 | 7,54E-05 | 0,00860706 | PECAM1 |
| 56,172082 | 2,216596592 | 0,55168791 | 2,57E-06 | 0,00067385 | RAB31 |
| 70,480477 | 2,213901059 | 0,49483167 | 3,47E-07 | 0,00015233 | ARRB1 |
| 68,5193962 | 2,204126349 | 0,54610027 | 2,45E-06 | 0,00066106 | SYK |
| 66,8884048 | 2,19913205 | 0,54999117 | 2,64E-06 | 0,00067385 | PDE3B |
| 146,28344 | 2,189589961 | 0,6710911 | 3,71E-05 | 0,0051743 | FGR |
| 60,2876389 | 2,16599925 | 0,58639666 | 8,96E-06 | 0,00178945 | RASGRP2 |
| 160,810889 | 2,116256498 | 0,80585737 | 0,00024015 | 0,01883899 | RASA3 |
| 63,2809588 | 2,095102417 | 0,53354204 | 3,56E-06 | 0,0008392 | AHR |
| 79,9410222 | 2,074752667 | 0,33195297 | 1,89E-11 | 5,53E-08 | SAXO1 |
| 287,336218 | 2,071754628 | 0,65093541 | 4,70E-05 | 0,00611735 | SPOCK2 |
| 93,9684607 | 2,066339265 | 0,80866556 | 0,00029824 | 0,0216556 | CSF2RB |
| 88,2367721 | 2,06539678 | 0,50971618 | 2,08E-06 | 0,00061284 | TMC8 |
| 42,2928455 | 2,044315955 | 0,66715586 | 8,11E-05 | 0,00894254 | SCML1 |
| 87,5732272 | 2,037915958 | 0,80674809 | 0,000325 | 0,02284331 | CHST15 |
| 57,2608858 | 2,030351964 | 0,58558202 | 2,00E-05 | 0,00340548 | SELL |
| 22,4050526 | 2,019196719 | 0,81743442 | 0,00044622 | 0,02816972 | MEFV |
| 55,7792318 | 2,005848279 | 0,5887193 | 2,41E-05 | 0,00374887 | ADAM28 |
| 50,2051284 | 1,960534112 | 0,52140316 | 7,30E-06 | 0,00150959 | TLE4 |
| 58,4645522 | 1,957197996 | 0,66338846 | 0,00011213 | 0,01092842 | CD84 |
| 31,1479913 | 1,952155796 | 0,7337312 | 0,00027643 | 0,02061053 | TESPA1 |
| 101,765496 | 1,935629081 | 0,90937516 | 0,00076209 | 0,04045768 | CXCL8 |
| 66,4747399 | 1,921022313 | 0,49617374 | 4,72E-06 | 0,00105041 | SNCA |
| 20,4593984 | 1,919262831 | 0,68566874 | 0,00020863 | 0,01681667 | SKAP1 |
| 64,5955909 | 1,914333676 | 0,62077667 | 7,21E-05 | 0,00833263 | NFAM1 |
| 89,1829241 | 1,909151276 | 0,51219372 | 7,54E-06 | 0,0015405 | SASH3 |
| 400,504886 | 1,907031173 | 0,69385345 | 0,00017203 | 0,01481843 | ALAS2 |
| 157,29656 | 1,904688406 | 0,77241477 | 0,00036433 | 0,02462331 | RASSF2 |
| 47,4215034 | 1,875443798 | 0,59034306 | 5,68E-05 | 0,00702307 | SRC |
| 81,4174201 | 1,823032423 | 0,90013709 | 0,00097885 | 0,04526397 | PTGS1 |
| 64,1957572 | 1,812478187 | 0,616491 | 0,00011416 | 0,01102249 | SH3BGRL2 |
| 43,1481364 | 1,806995846 | 0,64225882 | 0,0001699 | 0,01481843 | PRKAG2 |
| 66,2140258 | 1,806023689 | 0,55261418 | 3,90E-05 | 0,00530574 | ANPEP |
| 67,2899039 | 1,805342641 | 0,92104691 | 0,0011068 | 0,04898894 | MPEG1 |
| 58,4477524 | 1,798973546 | 0,60601845 | 0,00010469 | 0,01033917 | GNLY |
| 56,1966535 | 1,742470837 | 0,62817565 | 0,00019587 | 0,01615879 | ADA2 |
| 13,3409299 | 1,684433521 | 0,82395836 | 0,00108438 | 0,04851989 | ROS1 |
| 12,1765706 | 1,676857073 | 0,82700034 | 0,00111605 | 0,04902801 | RELN |
| 23,6892666 | 1,622980034 | 0,69311691 | 0,00053634 | 0,03238654 | ADAMTS2 |
| 78,1367349 | 1,545588666 | 0,3010895 | 1,23E-08 | 1,20E-05 | ADAMTSL1 |
| 70,8122624 | 1,540742692 | 0,58751936 | 0,00027005 | 0,02059691 | RAP1GAP2 |
| 191,295189 | 1,519944205 | 0,68524341 | 0,00068517 | 0,03770455 | NRGN |
| 54,9058585 | 1,500538977 | 0,68670145 | 0,00078644 | 0,04066247 | KCNA3 |

| | | | | | |
|---|---|---|---|---|---|
| 56,2059457 | 1,494515808 | 0,61286344 | 0,00044839 | 0,02816972 | C16orf54 |
| 52,1519531 | 1,481869887 | 0,49539144 | 9,63E-05 | 0,00995667 | FPR1 |
| 58,795756 | 1,462117234 | 0,62453078 | 0,00055193 | 0,03287612 | NCF1 |
| 251,185517 | 1,458801322 | 0,65135342 | 0,00064361 | 0,03601747 | CTSS |
| 482,784499 | 1,438514528 | 0,70581595 | 0,0009622 | 0,04516319 | LAPTM5 |
| 22,2947573 | 1,436491689 | 0,61629811 | 0,00056317 | 0,03332011 | AACSP1 |
| 111,436925 | 1,405683227 | 0,46986469 | 9,37E-05 | 0,00982436 | CCL5 |
| 46,2066515 | 1,403852401 | 0,59575388 | 0,00054701 | 0,03269393 | SPN |
| 69,5088187 | 1,356219256 | 0,55139348 | 0,00041016 | 0,02689279 | ARHGAP9 |
| 45,373572 | 1,338759528 | 0,59044777 | 0,00067901 | 0,03763913 | NCF1B |
| 63,3690869 | 1,327928829 | 0,63075638 | 0,00093403 | 0,04447896 | ALOX5 |
| 77,7713999 | 1,24773497 | 0,57073486 | 0,00077409 | 0,04066247 | BACH2 |
| 95,1453972 | 1,215200888 | 0,50247297 | 0,00044887 | 0,02816972 | PRKCB |
| 76,0930887 | 1,152615123 | 0,42951789 | 0,00023582 | 0,01858204 | BMF |
| 51,5482529 | 1,148304759 | 0,49593168 | 0,00059888 | 0,0343906 | LOC400499 |
| 264,118223 | 1,103309271 | 0,25086268 | 4,58E-07 | 0,00017894 | GPR146 |
| 340,291589 | 1,079615564 | 0,27009085 | 2,57E-06 | 0,00067385 | DEPP1 |
| 74,6792095 | 0,969160846 | 0,36888804 | 0,00027428 | 0,02059691 | LAMC3 |
| 37,4561031 | 0,96834808 | 0,43987747 | 0,00077874 | 0,04066247 | OPCML |
| 79,0544287 | 0,961600089 | 0,43937096 | 0,00079556 | 0,04099083 | SCARF1 |
| 123,288475 | 0,861976726 | 0,21268944 | 2,12E-06 | 0,00061284 | DACH1 |
| 430,338967 | 0,814081295 | 0,28417901 | 0,00014531 | 0,01302737 | CSF3R |
| 138,688835 | 0,808159692 | 0,30670087 | 0,0002791 | 0,0206934 | TNS1 |
| 87,070066 | 0,779283689 | 0,27264591 | 0,00015049 | 0,01335512 | FER1L4 |
| 59,8041488 | 0,769842426 | 0,3663592 | 0,00099927 | 0,04608697 | ZXDA |
| 398,993471 | 0,719193452 | 0,29504065 | 0,00047203 | 0,02900164 | FOS |
| 1032,34437 | 0,676807368 | 0,20126907 | 3,07E-05 | 0,00438614 | TNFAIP3 |
| 684,286112 | 0,665903729 | 0,16814753 | 3,23E-06 | 0,00077684 | ARRDC3 |
| 139,880623 | 0,642693761 | 0,28785045 | 0,00078646 | 0,04066247 | MYLIP |
| 178,195248 | 0,636835519 | 0,27346346 | 0,00062554 | 0,03511825 | HCLS1 |
| 351,307913 | 0,630296227 | 0,28111073 | 0,00076969 | 0,04066247 | LOC613266 |
| 683,633446 | 0,587527526 | 0,22189614 | 0,00028779 | 0,02115911 | SLC30A4 |
| 167,79604 | 0,554417299 | 0,19415038 | 0,00016433 | 0,01443808 | GOLGA8H |
| 471,737953 | 0,549819127 | 0,17740188 | 7,80E-05 | 0,00878399 | ABCA1 |
| 238,299666 | 0,501150629 | 0,22232205 | 0,0008119 | 0,04111454 | BRME1 |
| 211,820041 | 0,498628939 | 0,19997738 | 0,00046405 | 0,0286115 | TREX1 |
| 434,699666 | 0,488151503 | 0,138514 | 1,93E-05 | 0,00332165 | DEF6 |
| 495,881541 | 0,487776978 | 0,1633732 | 0,00011917 | 0,01125822 | HAPLN3 |
| 383,047249 | 0,482214921 | 0,17694414 | 0,00025209 | 0,01960045 | H2AB1 |
| 1492,90696 | 0,458752202 | 0,12384304 | 1,09E-05 | 0,002125 | ABLIM1 |
| 3414,26862 | 0,454540146 | 0,13391514 | 3,19E-05 | 0,00452009 | PIK3AP1 |
| 1464,85166 | 0,452874152 | 0,12304705 | 1,05E-05 | 0,00207823 | TENT5C |
| 816,741447 | 0,446539844 | 0,12647295 | 2,02E-05 | 0,00340548 | MID1IP1 |
| 355,129961 | 0,444948912 | 0,17532563 | 0,00043301 | 0,02776957 | TLE2 |
| 704,507743 | 0,439306432 | 0,14279133 | 9,47E-05 | 0,00984686 | C8orf82 |
| 1088,75567 | 0,429138538 | 0,19836894 | 0,00107191 | 0,04829634 | PNRC1 |
| 417,345663 | 0,427324418 | 0,18860032 | 0,00084925 | 0,04227498 | DMAC1 |
| 1534,74897 | 0,415742646 | 0,14692042 | 0,00020112 | 0,01641342 | KIFC2 |

| | | | | | |
|---|---|---|---|---|---|
| 790,09876 | 0,415182833 | 0,11643646 | 1,79E-05 | 0,00311259 | CORO1B |
| 776,018597 | 0,407314419 | 0,17920313 | 0,00085566 | 0,04235403 | MXD4 |
| 814,721664 | 0,405181402 | 0,11280141 | 1,65E-05 | 0,00296747 | PMF1 |
| 745,188 | 0,403188284 | 0,18130347 | 0,00097795 | 0,04526397 | H2AB3 |
| 401,09768 | 0,403181367 | 0,16989647 | 0,00068663 | 0,03770455 | TEDC2 |
| 5347,25967 | 0,397998849 | 0,1338714 | 0,00013784 | 0,01254998 | TMEM259 |
| 470,327116 | 0,392243806 | 0,14818109 | 0,00035223 | 0,02408321 | ZFYVE19 |
| 1167,04162 | 0,390574156 | 0,12261482 | 7,16E-05 | 0,00833174 | SH3GLB2 |
| 1669,36128 | 0,382078895 | 0,1392147 | 0,00027342 | 0,02059691 | BTG1 |
| 1593,31504 | 0,379176541 | 0,17351841 | 0,00113962 | 0,04944524 | PGGHG |
| 1084,01481 | 0,369944541 | 0,10146126 | 1,45E-05 | 0,0027381 | MET |
| 832,882218 | 0,368477282 | 0,13576667 | 0,00030365 | 0,02195739 | SEMA6B |
| 1618,02401 | 0,363168069 | 0,12616245 | 0,00018893 | 0,01580865 | ZBED10P |
| 3199,84501 | 0,361181509 | 0,15160448 | 0,00072455 | 0,0390543 | RORA |
| 1021,25302 | 0,360678911 | 0,12848032 | 0,00024507 | 0,01913956 | KATNB1 |
| 14740,271 | 0,357828766 | 0,16118689 | 0,00095343 | 0,0449469 | HS6ST1 |
| 990,069452 | 0,353840083 | 0,1321342 | 0,00034821 | 0,0239951 | ECSIT |
| 836,289905 | 0,342671106 | 0,14870281 | 0,00091705 | 0,0441491 | CCDC85B |
| 2790,89297 | 0,341775907 | 0,12242743 | 0,00026233 | 0,02025071 | SIPA1 |
| 2059,75435 | 0,339238703 | 0,1164313 | 0,00019088 | 0,01589631 | ABHD17A |
| 4450,48036 | 0,334170862 | 0,12242698 | 0,0003174 | 0,02248752 | POLD1 |
| 468,159463 | 0,331057891 | 0,14940228 | 0,00113673 | 0,04944212 | ADAMTS17 |
| 730,47654 | 0,323557139 | 0,11353325 | 0,00022802 | 0,01813029 | PCSK6 |
| 3556,19396 | 0,318483622 | 0,1321449 | 0,000847 | 0,04227498 | PKN1 |
| 955,527913 | 0,316235054 | 0,14019198 | 0,00109299 | 0,04857826 | SCAMP5 |
| 6053,13646 | 0,311642807 | 0,09222579 | 4,36E-05 | 0,00576133 | MAZ |
| 1781,16215 | 0,311109562 | 0,10693466 | 0,00020176 | 0,01641342 | CDT1 |
| 1004,22954 | 0,30531595 | 0,09939231 | 0,0001229 | 0,01148676 | IRS1 |
| 4510,03526 | 0,303754038 | 0,09825389 | 0,00011899 | 0,01125822 | IRAK1 |
| 1413,9713 | 0,291162088 | 0,10751963 | 0,00037859 | 0,02546639 | ZNF787 |
| 1912,78573 | 0,2891471 | 0,12477407 | 0,00111336 | 0,04902801 | PTPRJ |
| 853,912321 | 0,287511868 | 0,12531369 | 0,00108515 | 0,04851989 | PGAM4 |
| 3075,10884 | 0,285195665 | 0,11251419 | 0,0006216 | 0,03511825 | RELB |
| 1299,41043 | 0,285084773 | 0,10234538 | 0,00030854 | 0,0221295 | PTGES2 |
| 1510,19641 | 0,28204239 | 0,11777199 | 0,00088639 | 0,04326585 | FPGS |
| 4082,31389 | 0,278444185 | 0,10380044 | 0,00042623 | 0,02753569 | GRAMD1A |
| 2325,36894 | 0,277201654 | 0,11682773 | 0,00092494 | 0,04419875 | CDCA4 |
| 6465,64865 | 0,26973757 | 0,10281902 | 0,00056201 | 0,03332011 | PCED1A |
| 2238,73012 | 0,259410911 | 0,09189873 | 0,00031197 | 0,02228461 | ACADVL |
| 1663,83508 | 0,253917481 | 0,10171472 | 0,00080128 | 0,04099083 | RPS6KB2 |
| 1404,22486 | 0,240376645 | 0,10122506 | 0,00112237 | 0,04918295 | TBC1D10B |
| 34,8332919 | 0,038512789 | 0,08824036 | 4,11E-06 | 0,00093749 | PF4 |
| 26,6028342 | 0,037575678 | 0,0872922 | 4,92E-05 | 0,00631375 | ANKRD44-IT1 |
| 24,433054 | 0,035837322 | 0,08578444 | 1,79E-05 | 0,00311259 | CD28 |
| 26,5421438 | 0,035660236 | 0,08509687 | 0,00085172 | 0,04227824 | CD14 |
| 23,6424069 | 0,034625189 | 0,08478453 | 0,00010001 | 0,01009966 | LILRA5 |
| 30,0616263 | 0,032511851 | 0,08305509 | 6,43E-05 | 0,00779332 | SNX19 |
| 24,7476276 | 0,032102413 | 0,08268499 | 0,00017164 | 0,01481843 | IRAG2 |

| | | | | | |
|---|---|---|---|---|---|
| 22,2113289 | 0,031811643 | 0,08254181 | 0,00012477 | 0,0116006 | LTF |
| 27,2456179 | 0,030773206 | 0,08167328 | 0,00036297 | 0,02462331 | TENM1 |
| 32,4297327 | 0,030695116 | 0,08150323 | 0,00093848 | 0,04457002 | CHFR |
| 29,1951395 | 0,030157532 | 0,08144999 | 4,63E-05 | 0,00607019 | ABCB1 |
| 10,6711953 | 0,029471878 | 0,08099971 | 0,0001173 | 0,01124499 | LPCAT2 |
| 19,8190953 | 0,02883972 | 0,08049143 | 0,00021912 | 0,0175017 | FGD4 |
| 37,7851513 | 0,028406806 | 0,08022579 | 0,00018236 | 0,01548042 | LILRB2 |
| 18,1409653 | 0,028111487 | 0,08011699 | 0,00010473 | 0,01033917 | TLR4 |
| 21,2170782 | 0,027850195 | 0,08000795 | 0,00012771 | 0,01181119 | CD52 |
| 16,8853312 | 0,027585855 | 0,0797921 | 0,00029178 | 0,0212742 | CSF2RA |
| 29,1878039 | 0,027220106 | 0,07955393 | 0,000153 | 0,01350996 | NELL2 |
| 24,585879 | 0,027181313 | 0,07938143 | 0,00069342 | 0,03772374 | LOC102723996 |
| 33,6397644 | 0,026803519 | 0,07921288 | 0,0004325 | 0,02776957 | ARHGAP25 |
| 25,0049782 | 0,026667817 | 0,07920704 | 0,00018716 | 0,01573568 | MS4A6A |
| 14,9186564 | 0,02605035 | 0,07892805 | 0,00011025 | 0,01082291 | FAM20C |
| 32,3424812 | 0,026028869 | 0,07881331 | 0,0002723 | 0,02059691 | FCGR3B |
| 18,2056015 | 0,025515751 | 0,07857996 | 0,00041294 | 0,02697456 | P2RY2 |
| 29,3464939 | 0,025441053 | 0,07853559 | 0,00014931 | 0,01331804 | FCMR |
| 18,7954339 | 0,025208712 | 0,07831117 | 0,00065084 | 0,03630627 | SH2D1A |
| 35,3108527 | 0,025157284 | 0,07824788 | 0,0009236 | 0,04419875 | CD7 |
| 20,6983161 | 0,024615736 | 0,0779544 | 0,00113231 | 0,04944212 | SLC15A3 |
| 24,1420852 | 0,023881954 | 0,07772942 | 0,00020576 | 0,01666181 | MX1 |
| 16,8080044 | 0,023684858 | 0,07757107 | 0,00038116 | 0,02546639 | ABCA5 |
| 12,8705838 | 0,023260728 | 0,07731988 | 0,00109361 | 0,04857826 | JCHAIN |
| 30,1688632 | 0,023007821 | 0,07721858 | 0,00053136 | 0,03230827 | NUAK2 |
| 27,291533 | 0,022838325 | 0,07713475 | 0,00057363 | 0,0334719 | ZNF91 |
| 20,9332379 | 0,022443513 | 0,07694469 | 0,00061546 | 0,03488688 | CKAP4 |
| 26,6745615 | 0,02120933 | 0,07637095 | 0,00089475 | 0,04355296 | MNDA |
| 14,7430945 | 0,021119491 | 0,07633843 | 0,0010569 | 0,04798932 | MME |
| 17,58282 | 0,020866293 | 0,07631597 | 0,00017564 | 0,01498263 | MIR3667HG |
| 21,9477883 | 0,020643648 | 0,07617023 | 0,00039138 | 0,02585427 | RGCC |
| 14,2707555 | 0,020549938 | 0,07607327 | 0,00111472 | 0,04902801 | PTGER2 |
| 20,2030734 | 0,020193574 | 0,07597571 | 0,00046298 | 0,0286115 | XK |
| 17,0683891 | 0,019365799 | 0,07561757 | 0,00081706 | 0,04125684 | ZNF841 |
| 21,2351808 | 0,019148946 | 0,07551418 | 0,00101367 | 0,04662884 | MS4A1 |
| 17,966136 | 0,018605221 | 0,07531777 | 0,00087628 | 0,04289152 | LILRA1 |
| 12,4843674 | 0,01852582 | 0,07534351 | 3,84E-05 | 0,0052753 | ZNF14 |
| 11,0476049 | 0,018453407 | 0,0752626 | 0,00083956 | 0,04203063 | TRAT1 |
| 14,8186717 | 0,018193478 | 0,07518946 | 0,00081157 | 0,04111454 | ZNF831 |
| 10,2670072 | 0,017993099 | 0,07510609 | 0,00011257 | 0,01092842 | ZSCAN18 |
| 9,59478011 | 0,017663082 | 0,07496318 | 0,00102748 | 0,04714051 | FFAR2 |
| 21,5222585 | 0,017626726 | 0,07495512 | 0,00096786 | 0,04516319 | WWC3 |
| 43,8498002 | 0,016099529 | 0,07445561 | 0,00048427 | 0,02964995 | SNX9 |
| 43,7305298 | 0,015894653 | 0,07435847 | 0,00113629 | 0,04944212 | LRRC25 |
| 10,8863141 | 0,014844146 | 0,07402753 | 0,000338 | 0,02347576 | CXCR1 |
| 5,19290026 | 0,013570792 | 0,07368649 | 0,00017441 | 0,01494969 | LOC388436 |
| 9,58527933 | 0,013438942 | 0,07362853 | 0,0004145 | 0,0269765 | RIN2 |
| 7,36730332 | 0,013250809 | 0,07356752 | 0,00061403 | 0,03488688 | RIPK3 |

| | | | | | |
|---|---|---|---|---|---|
| 5,88688433 | 0,013214031 | 0,07356651 | 0,00046027 | 0,0286115 | LOC102723604 |
| 5,87892624 | 0,012688341 | 0,07341843 | 0,00075598 | 0,04025496 | ARHGEF5 |
| 3,94426266 | 0,010847433 | 0,07299405 | 0,00086121 | 0,04239003 | GRAPL-AS1 |
| 366,199532 | -0,022133317 | 0,07690108 | 9,13E-05 | 0,00966136 | SNORD3C |
| 9,00319899 | -0,024444913 | 0,07793986 | 0,00054313 | 0,03257286 | PLCE1-AS2 |
| 54,083436 | -0,046792357 | 0,09576016 | 0,00083864 | 0,04203063 | SCARNA9 |
| 30,8311819 | -0,048099134 | 0,09714115 | 0,00104365 | 0,04775798 | VTRNA1-3 |
| 34,2093581 | -0,051036778 | 0,10107207 | 0,00057412 | 0,0334719 | RNVU1-14 |
| 39,6215856 | -0,059567195 | 0,11518129 | 9,11E-05 | 0,00966136 | STARD8 |
| 75,414397 | -0,080836901 | 0,16798498 | 0,0005673 | 0,03343234 | IGF1 |
| 1362,539 | -0,238220198 | 0,09759677 | 0,00091586 | 0,0441491 | UBE2Q1 |
| 1036,13101 | -0,253020603 | 0,10473293 | 0,00091264 | 0,0441491 | RBM28 |
| 6108,67539 | -0,256886169 | 0,10220827 | 0,00072292 | 0,0390543 | FUS |
| 3469,26672 | -0,262623887 | 0,08185602 | 9,07E-05 | 0,00966136 | HNRNPM |
| 2220,15628 | -0,265299643 | 0,10401775 | 0,00062396 | 0,03511825 | ZC3H13 |
| 1282,99718 | -0,266601358 | 0,11477584 | 0,00109475 | 0,04857826 | MDC1 |
| 1386,52816 | -0,266668964 | 0,10589523 | 0,00069082 | 0,03772374 | PRR13 |
| 4269,86227 | -0,267256405 | 0,10593396 | 0,00071192 | 0,03861048 | DST |
| 4746,99626 | -0,279669031 | 0,11343613 | 0,00073364 | 0,0394234 | TM4SF1 |
| 1044,97386 | -0,28457382 | 0,11755029 | 0,00080479 | 0,04099083 | GOLIM4 |
| 594,352962 | -0,2929778 | 0,12556903 | 0,00096531 | 0,04516319 | DDRGK1 |
| 882,769499 | -0,295157156 | 0,12420689 | 0,00086056 | 0,04239003 | FTSJ3 |
| 2679,16919 | -0,299563494 | 0,10584377 | 0,00030705 | 0,02211256 | EIF5B |
| 6098,72052 | -0,300558405 | 0,08991034 | 5,19E-05 | 0,00651285 | HMGCR |
| 926,950163 | -0,301008785 | 0,12518415 | 0,00078678 | 0,04066247 | H4C3 |
| 3013,01315 | -0,301837614 | 0,1036324 | 0,00019992 | 0,01641342 | EIF4G3 |
| 3870,54833 | -0,310055308 | 0,12802679 | 0,00073914 | 0,03959796 | FDPS |
| 1040,48869 | -0,310061178 | 0,11582668 | 0,00038286 | 0,02548089 | NDUFA9 |
| 865,627493 | -0,310698708 | 0,12966499 | 0,00078257 | 0,04066247 | RBM34 |
| 493,885963 | -0,313043877 | 0,13927463 | 0,00105634 | 0,04798932 | EXOSC7 |
| 2155,20533 | -0,313850405 | 0,13116416 | 0,00074798 | 0,03994991 | CD63 |
| 1008,82042 | -0,3148915 | 0,133867 | 0,0007824 | 0,04066247 | AIFM1 |
| 647,012099 | -0,316693606 | 0,13859253 | 0,00106252 | 0,04812027 | NDUFAB1 |
| 1862,37738 | -0,319977314 | 0,12999484 | 0,00061506 | 0,03488688 | SMARCE1 |
| 8633,80897 | -0,322293218 | 0,11506716 | 0,00026276 | 0,02025071 | RPL23A |
| 794,703462 | -0,322396293 | 0,12735938 | 0,00056888 | 0,03343234 | TIMM13 |
| 1695,25607 | -0,322962655 | 0,11622726 | 0,00027297 | 0,02059691 | NUDC |
| 14237,2942 | -0,323083591 | 0,10194029 | 8,01E-05 | 0,00894254 | HNRNPU |
| 1050,10481 | -0,340986882 | 0,14386927 | 0,00077642 | 0,04066247 | OXCT1 |
| 16328,1794 | -0,342300266 | 0,09763676 | 2,63E-05 | 0,00401267 | HNRNPA2B1 |
| 336,536313 | -0,343630664 | 0,15155633 | 0,00096915 | 0,04516319 | CASC15 |
| 737,712759 | -0,345374131 | 0,12743028 | 0,00031503 | 0,02241187 | TTC1 |
| 7015,69147 | -0,349445882 | 0,12049361 | 0,00018701 | 0,01573568 | CTSL |
| 19916,2497 | -0,349633172 | 0,15248082 | 0,00092563 | 0,04419875 | TIMP1 |
| 1923,94264 | -0,355244116 | 0,1081979 | 5,65E-05 | 0,00702307 | SNRPD3 |
| 306,031856 | -0,355929308 | 0,16008581 | 0,00104898 | 0,04787692 | COPRS |
| 2149,2199 | -0,355964939 | 0,10070117 | 2,06E-05 | 0,0034551 | ANP32B |
| 13610,1712 | -0,356805873 | 0,10999621 | 6,18E-05 | 0,00759872 | NCL |

| | | | | | |
|---|---|---|---|---|---|
| 9225,07395 | -0,359266518 | 0,14773906 | 0,00060804 | 0,03480269 | RPL19 |
| 501,726171 | -0,365266524 | 0,14872121 | 0,00058382 | 0,03375418 | ZNF354B |
| 3255,94957 | -0,365720617 | 0,16322908 | 0,00094389 | 0,04470611 | RPLP2 |
| 1578,83603 | -0,368301452 | 0,14396134 | 0,00045059 | 0,02817738 | NASP |
| 1556,79966 | -0,369012606 | 0,12089491 | 9,96E-05 | 0,01009966 | H2BC5 |
| 969,041132 | -0,371158372 | 0,1445876 | 0,00044246 | 0,02816972 | ATP5MK |
| 1921,83098 | -0,374100811 | 0,09095836 | 2,22E-06 | 0,00062152 | MRFAP1 |
| 457,389163 | -0,378383936 | 0,14561864 | 0,00040611 | 0,02672715 | HABP4 |
| 856,21858 | -0,383103361 | 0,14093546 | 0,0002874 | 0,02115911 | ATP5MJ |
| 612,528728 | -0,383263959 | 0,17593714 | 0,00107554 | 0,04833591 | TOMM5 |
| 1170,38324 | -0,384434015 | 0,1689945 | 0,00087593 | 0,04289152 | PSMB1 |
| 1000,56286 | -0,388755899 | 0,14795279 | 0,00038427 | 0,02548089 | PSMB6 |
| 494,605597 | -0,389547856 | 0,13662254 | 0,00019506 | 0,01615879 | SDCCAG8 |
| 7824,09382 | -0,391874525 | 0,18383122 | 0,00097153 | 0,04516319 | RPL12 |
| 748,175923 | -0,395019655 | 0,12924939 | 0,00010352 | 0,0103355 | DAD1 |
| 10093,0908 | -0,395213537 | 0,12412263 | 8,14E-05 | 0,00894254 | CD163 |
| 1831,53489 | -0,398349478 | 0,12985849 | 9,83E-05 | 0,01009966 | MGA |
| 704,660413 | -0,4007484 | 0,11431276 | 2,25E-05 | 0,00361794 | TPMT |
| 479,783883 | -0,401007416 | 0,162487 | 0,00054216 | 0,03257286 | EXOSC3 |
| 9084,69096 | -0,401045833 | 0,11677079 | 2,53E-05 | 0,00390246 | MKI67 |
| 875,632885 | -0,401727152 | 0,179186 | 0,0008037 | 0,04099083 | ATP5MG |
| 1518,16074 | -0,403806795 | 0,18583625 | 0,00107111 | 0,04829634 | PRKXP1 |
| 479,272444 | -0,410933344 | 0,12017875 | 2,92E-05 | 0,00427259 | TRMT6 |
| 1107,12944 | -0,414580655 | 0,13043209 | 6,78E-05 | 0,00804907 | PSMA5 |
| 736,095247 | -0,415108839 | 0,1577694 | 0,00035169 | 0,02408321 | MANF |
| 12566,6605 | -0,415726351 | 0,09811981 | 1,17E-06 | 0,00037249 | HNRNPA1 |
| 732,892902 | -0,420892286 | 0,13291309 | 7,00E-05 | 0,00819968 | NOP16 |
| 666,103117 | -0,424266463 | 0,12879278 | 4,35E-05 | 0,00576133 | UBLCP1 |
| 539,826512 | -0,425923342 | 0,14332981 | 0,00012978 | 0,01187792 | GPATCH4 |
| 1013,23878 | -0,439181117 | 0,12893512 | 3,06E-05 | 0,00438614 | MZT2B |
| 1146,1791 | -0,441305663 | 0,12387288 | 1,73E-05 | 0,00306251 | CCDC47 |
| 1086,733 | -0,443725685 | 0,12305715 | 1,46E-05 | 0,0027381 | IWS1 |
| 1965,08549 | -0,445165496 | 0,08923233 | 3,22E-08 | 2,10E-05 | BMS1 |
| 4037,80183 | -0,445269737 | 0,12392435 | 1,53E-05 | 0,00282619 | RPS12 |
| 894,849397 | -0,447999299 | 0,19134495 | 0,00069333 | 0,03772374 | NALCN |
| 4305,79312 | -0,448918361 | 0,18587095 | 0,00057526 | 0,0334719 | RPL27A |
| 643,761986 | -0,44914739 | 0,10965602 | 2,13E-06 | 0,00061284 | HTATSF1 |
| 535,057471 | -0,449259994 | 0,20298774 | 0,00090965 | 0,0441491 | DNAJB11 |
| 419,723581 | -0,454983125 | 0,14340432 | 6,59E-05 | 0,00792602 | CCDC34 |
| 13874,7501 | -0,464879811 | 0,1782658 | 0,00031865 | 0,02248752 | RPS23 |
| 321,335959 | -0,465359153 | 0,20698024 | 0,00079796 | 0,04099083 | NDUFB9 |
| 383,685979 | -0,468961815 | 0,18617218 | 0,00044663 | 0,02816972 | NDUFA3 |
| 1507,85968 | -0,474424205 | 0,11403754 | 1,55E-06 | 0,00046831 | SYN1 |
| 283,978059 | -0,475165189 | 0,18489765 | 0,00037988 | 0,02546639 | NSRP1 |
| 554,200323 | -0,476475175 | 0,1630209 | 0,00014152 | 0,01275312 | MEA1 |
| 1443,97728 | -0,478006895 | 0,11923529 | 2,92E-06 | 0,0007333 | PNN |
| 11154,0543 | -0,478024863 | 0,19144736 | 0,00033348 | 0,02325331 | RPS11 |
| 669,493227 | -0,478029433 | 0,12605611 | 6,73E-06 | 0,00142403 | ALYREF |

| | | | | | |
|---|---|---|---|---|---|
| 339,378717 | -0,480320582 | 0,16163883 | 0,00012112 | 0,01138117 | MEIS2 |
| 585,701011 | -0,482437232 | 0,15727848 | 8,97E-05 | 0,00966136 | NEDD8 |
| 365,422282 | -0,483508052 | 0,19420129 | 0,00046337 | 0,0286115 | MAGOH |
| 447,462074 | -0,484157423 | 0,15450495 | 7,42E-05 | 0,00852438 | OSTF1 |
| 402,72437 | -0,48762013 | 0,21774906 | 0,00083396 | 0,04198966 | GAR1 |
| 4089,53119 | -0,490086954 | 0,12226536 | 2,99E-06 | 0,00073713 | NACA |
| 1736,9529 | -0,493505895 | 0,14998509 | 4,28E-05 | 0,00574642 | GTPBP4 |
| 553,85042 | -0,502410692 | 0,12376234 | 2,23E-06 | 0,00062152 | GGCT |
| 785,916496 | -0,507163159 | 0,16809443 | 0,00010319 | 0,0103355 | MRPL51 |
| 2524,9495 | -0,513640896 | 0,16907193 | 8,10E-05 | 0,00894254 | H4C14 |
| 1455,0516 | -0,513819423 | 0,10926463 | 1,26E-07 | 6,53E-05 | HNRNPH3 |
| 7349,40555 | -0,514461845 | 0,13357217 | 5,04E-06 | 0,00110663 | RPL14 |
| 462,444825 | -0,52355317 | 0,20199251 | 0,0003438 | 0,0237847 | MRPL24 |
| 2204,99664 | -0,528895224 | 0,13419176 | 3,75E-06 | 0,00086619 | DNAJA1 |
| 242,572028 | -0,52912977 | 0,22235979 | 0,00058396 | 0,03375418 | COA6 |
| 1526,31217 | -0,529352731 | 0,16072227 | 4,12E-05 | 0,00557234 | ZFP36 |
| 546,417425 | -0,532900397 | 0,15286417 | 2,21E-05 | 0,00360271 | SEC61G |
| 2977,61736 | -0,533251763 | 0,11509933 | 1,87E-07 | 9,41E-05 | FXYD5 |
| 2531,63243 | -0,53489276 | 0,16391086 | 5,06E-05 | 0,00644514 | H4C15 |
| 852,671791 | -0,534894446 | 0,14187845 | 7,29E-06 | 0,00150959 | U2AF1 |
| 11939,6468 | -0,535639725 | 0,15406758 | 2,17E-05 | 0,00359051 | TNC |
| 915,979679 | -0,544684153 | 0,1221812 | 3,91E-07 | 0,00016752 | PDCD5 |
| 1379,48309 | -0,546795146 | 0,12282233 | 4,01E-07 | 0,00016794 | SAFB |
| 756,992807 | -0,547674005 | 0,20424303 | 0,00026558 | 0,02037902 | H2BC9 |
| 377,574257 | -0,552417647 | 0,18634164 | 0,00011775 | 0,01124499 | MRFAP1L1 |
| 458,686272 | -0,553123046 | 0,2390864 | 0,00068189 | 0,0376795 | NDUFS5 |
| 256,568411 | -0,556892126 | 0,23059134 | 0,00052329 | 0,03192782 | CSTB |
| 1045,94304 | -0,566329715 | 0,13140362 | 7,53E-07 | 0,0002744 | RRBP1 |
| 216,444427 | -0,567950025 | 0,18086291 | 6,74E-05 | 0,00804907 | F8A1 |
| 186,519042 | -0,569792373 | 0,21966258 | 0,00033214 | 0,02325229 | LSM6 |
| 3057,01925 | -0,58355752 | 0,12328076 | 9,68E-08 | 5,15E-05 | FABP5 |
| 384,304387 | -0,592233067 | 0,13814026 | 8,21E-07 | 0,00028869 | ZDHHC21 |
| 179,364611 | -0,595584468 | 0,25878612 | 0,00067865 | 0,03763913 | LMO2 |
| 12312,4618 | -0,618045233 | 0,12266047 | 2,55E-08 | 1,84E-05 | RPS18 |
| 818,28054 | -0,627698347 | 0,15074345 | 1,38E-06 | 0,00042432 | NRP1 |
| 693,206322 | -0,635045407 | 0,12903299 | 3,90E-08 | 2,45E-05 | UQCRH |
| 171,494958 | -0,641553414 | 0,23708863 | 0,00023538 | 0,01858204 | CDC26 |
| 141,165562 | -0,641995653 | 0,26852026 | 0,00053444 | 0,03238344 | GLRX2 |
| 151,651268 | -0,660585749 | 0,26566263 | 0,00041942 | 0,02719601 | DELEC1 |
| 1451,6401 | -0,663338726 | 0,13229392 | 2,39E-08 | 1,84E-05 | RPS29 |
| 184,665817 | -0,692146781 | 0,22048217 | 6,31E-05 | 0,00770265 | SEC11C |
| 495,335194 | -0,701316833 | 0,16623757 | 1,07E-06 | 0,0003542 | UBL5 |
| 231,521912 | -0,702389085 | 0,16716902 | 1,15E-06 | 0,00037249 | SCG5 |
| 491,753773 | -0,703139383 | 0,15695369 | 3,30E-07 | 0,00014886 | RPL36AL |
| 101,044764 | -0,709648425 | 0,27068355 | 0,00029131 | 0,0212742 | VANGL1 |
| 131,460269 | -0,713574938 | 0,23789876 | 1,00E-04 | 0,01009966 | SLC25A24 |
| 80,147372 | -0,731901211 | 0,34579487 | 0,00095409 | 0,0449469 | ZFP2 |
| 129,151449 | -0,747201535 | 0,22019001 | 2,68E-05 | 0,00401894 | TMEM216 |

| | | | | | |
|---|---|---|---|---|---|
| 175,247511 | -0,760498955 | 0,22474553 | 2,74E-05 | 0,00408382 | ATP5ME |
| 438,472746 | -0,764604492 | 0,15777354 | 5,67E-08 | 3,32E-05 | TMA7 |
| 6276,38313 | -0,767988267 | 0,14928371 | 1,15E-08 | 1,19E-05 | RPL37A |
| 85,5560892 | -0,782740135 | 0,33641725 | 0,00059509 | 0,03428468 | RAB5IF |
| 200,148466 | -0,78385076 | 0,1974163 | 3,02E-06 | 0,00073713 | PET100 |
| 336,098568 | -0,85842376 | 0,21885191 | 3,58E-06 | 0,0008392 | H2BC14 |
| 244,969531 | -0,879161007 | 0,19950814 | 4,54E-07 | 0,00017894 | NDUFA1 |
| 886,963609 | -0,919307443 | 0,18251657 | 2,09E-08 | 1,83E-05 | SCARNA2 |
| 874,125587 | -0,942193059 | 0,15064593 | 1,87E-11 | 5,53E-08 | SH3BGRL3 |
| 690,256596 | -0,968573118 | 0,21937939 | 4,22E-07 | 0,00017247 | LOC102724594 |
| 4538,32684 | -0,9798107 | 0,12499002 | 2,20E-16 | 3,86E-12 | FCER1G |
| 530,636911 | -0,996570647 | 0,19835548 | 2,19E-08 | 1,83E-05 | SNRPE |
| 266,63507 | -1,001845084 | 0,18140708 | 1,49E-09 | 1,87E-06 | PFDN2 |
| 4495,76517 | -1,010325358 | 0,14324345 | 3,37E-14 | 1,48E-10 | RPL11 |
| 72,9943477 | -1,010647737 | 0,32951705 | 7,63E-05 | 0,00865378 | SCARNA17 |
| 751,372313 | -1,032952871 | 0,18875845 | 1,97E-09 | 2,31E-06 | HSPE1 |
| 2210,89273 | -1,050063529 | 0,14528781 | 2,29E-14 | 1,34E-10 | RPS27 |
| 423,987763 | -1,058143547 | 0,20173434 | 6,87E-09 | 7,55E-06 | TOMM7 |
| 403,987744 | -1,09242855 | 0,21924265 | 2,72E-08 | 1,84E-05 | C19orf53 |
| 244,712002 | -1,097497618 | 0,18432187 | 1,20E-10 | 2,35E-07 | NDUFS6 |
| 780,171974 | -1,321510871 | 0,18083686 | 1,23E-14 | 1,08E-10 | H4C5 |
| 34,5996767 | -1,56435564 | 0,49162117 | 5,12E-05 | 0,00646992 | POPDC1 |
| 15,265598 | -3,301283754 | 0,67312446 | 6,29E-08 | 3,56E-05 | NBPF18P |
