## Supplemental Informations for "*NPM1::ALK* fusion-derived circular RNAs drive aggressiveness in ALK-positive anaplastic large cell lymphoma"

**Running head: Circular RNAs in ALCL**

Loelia Babin,[1,2,3*] Lola Colras,[1,23] Sandra Dailhau,[1,2,3] Steffen Fuchs,[1,2,3] Stéphane Pyronnet,[1,2] Laurence Lamant,[1,2,3] Fabienne Meggetto.[1,2,3*]

**Supplemental Table S1.** Antibodies and primers used.

Antibodies

| Target | Provider | Ref |
|---|---|---|
| ALK | Cell signaling technology | 3633 |
| STAT3 | Cell signaling technology | 9132 |
| pALK | Cell signaling technology | 3341 |
| pSTAT3 | Cell signaling technology | 9131 |
| GAPDH | Santa Cruz | sc-25778 |
| CXCL5 | Abcam | ab305100 |
| TIMP2 | Cell signaling technology | 5738 |
| IL-1alpha | Cell signaling technology | 34819 |

Primers

| Primers | Sequence |
|---|---|
| mRNA-NPM-ALK-F | ACCACCAGTGGTCTTAAGGTTG |
| mRNA-NPM-ALK-B | CGGAGCTTGCTCAGCTTGTA |
| circ-A26N3-F | GAGACATCTACAGGGCGAGC |
| circ-A26N3-B | CAGCCCCTAAACTGACCATG |
| circ-A23N2-F | CATTGTTCGCTGCATTGGGG |
| circ-A23N2-B | GCCTTTAGTTCACAACCGGG |
| MLN51-F | TAATCCCAGTTACCCTTATGCTCCA |
| MLN51-B | GTTATAGTAGGTCACTCCTCCATATACCTGT |
| circNPM-ALK-div-F | GCAGAGCCCTGAGTACAAGC |
| circNPM-ALK-div-B | AATATGCACTGGCCCTGAAC |

**Supplemental figure legends:**

**Supplemental Figure S1: (A)** Schematic representation of the divergent primer design used to detect *NPM1::ALK* fusion-derived circular RNAs, with binding sites located in exon 4 of NPM1 and exon 20 of ALK. **(B)** Schematic representation of *NPM1::ALK* fusion-derived f-circ-A23N2 and f-circ-A26N3. **(C)** Subcellular localization of f-circ-A23N2 and f-circ-A26N3 determined by cellular fractionation followed by RT-qPCR. *MALAT1* long noncoding RNA (lincMALAT1) and *GAPDH* mRNAs served as nuclear and cytoplasmic controls, respectively**.**

**Supplemental Figure S2: *In vitro* effects of f-circA23N2 and f-circA26N3 overexpression on cell proliferation and cell-cycle distribution in crizotinib-treated SU-DHL-1 cells.** SU-DHL-1 cells were transduced with either an empty vector (empty) or expression vectors encoding f-circ-A23N2 or f-circ-A26N3 and treated or not with crizotinib (200 or 350 nM). (**A**) Cell proliferation monitored by cell trace dilution assay over time (from day 0 to day 7), showing no modification. (**B**) Cell cycle distribution analysis showing the absence of alteration.

**Supplemental Figure S3. Transcriptomic alterations induced by *NPM::ALK*-derived f-circRNA overexpression in SU-DHL-1 cells.** (**A**) Volcano plots showing differentially expressed genes in SU-DHL-1 cells overexpressing the *NPM::ALK*-derived f-circ-A23N2 and f-circ-A26N3 compared with cells transduced with an empty vector used as control (empty). Differential expression analysis was performed relative to the control condition. (**B**) Gene Set Enrichment Analysis (GSEA) of the differentially expressed genes identified in A.

**Supplemental Figure S4.** ImmuCC deconvolution analysis of the relative abundance of immune sub-populations in NSG tumor models obtained using RNAseq murine reads on xenograft tumo
rs (N = 3 per group) generated from control empty vector (empty) or f-circA23N2 (OE-A23N2) and f-circA26N3 (OE-A26N3)-overexpressing SU-DHL-1 cells.

### Supplemental Figure S1

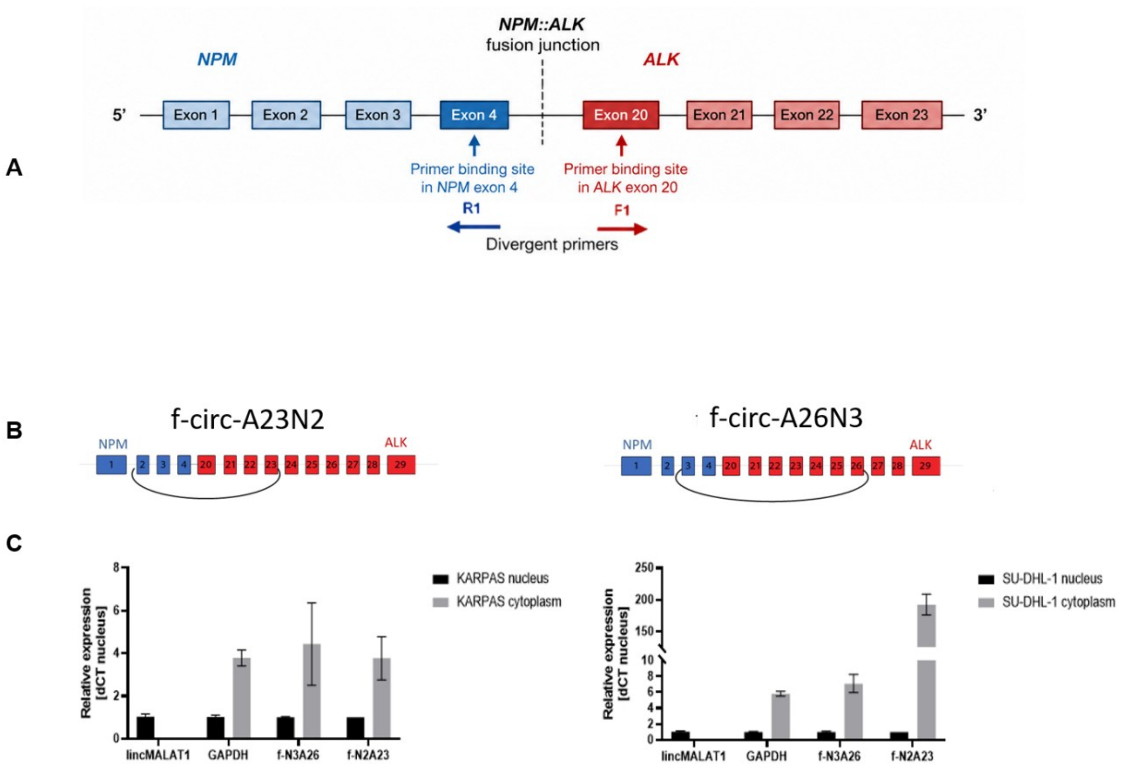


### Supplemental Figure S2

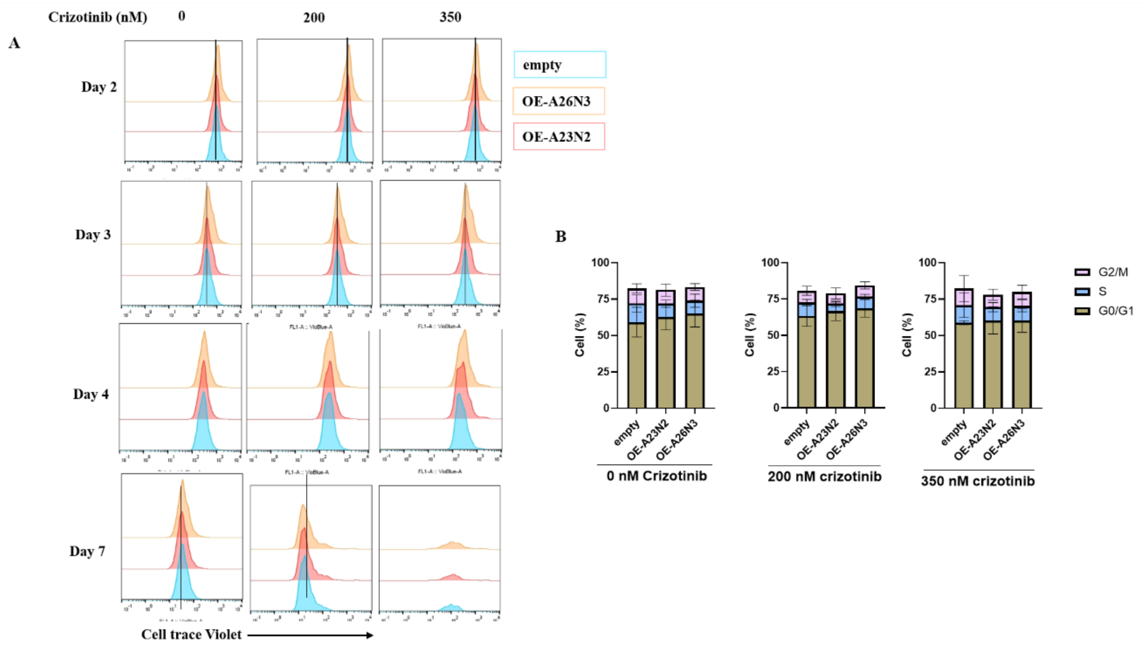

### Supplemental Figure S3

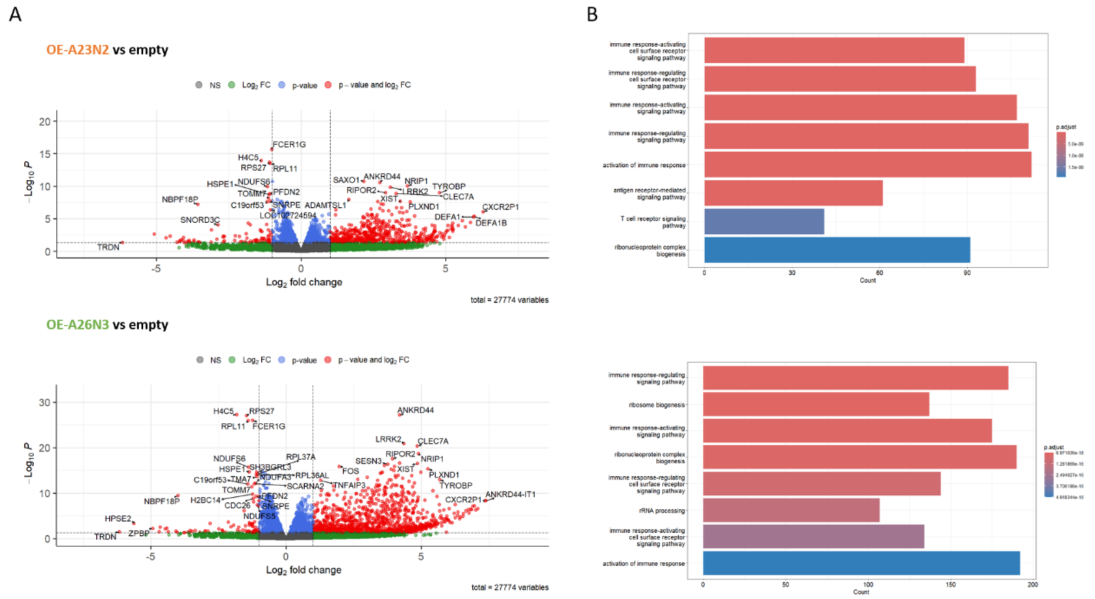


### Supplemental Figure S4

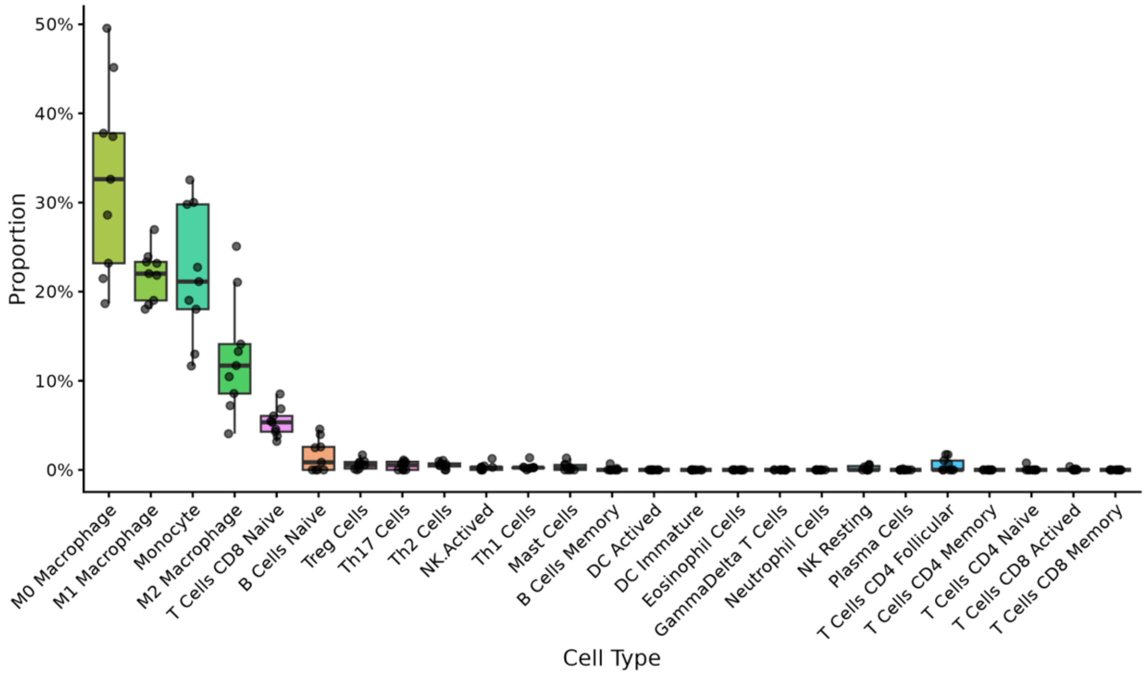
